# Prolonged morphological expansion of spiny-rayed fishes following the end-Cretaceous

**DOI:** 10.1101/2021.07.12.452083

**Authors:** Ava Ghezelayagh, Richard C. Harrington, Edward D. Burress, Matthew A. Campbell, Janet C. Buckner, Prosanta Chakrabarty, Jessica R. Glass, W. Tyler McCraney, Peter J. Unmack, Christine E. Thacker, Michael E. Alfaro, Sarah T. Friedman, William B. Ludt, Peter F. Cowman, Matt Friedman, Samantha A. Price, Alex Dornburg, Brant C. Faircloth, Peter C. Wainwright, Thomas J. Near

**Author notes:** These authors contributed equally to this work (Ava Ghezelayagh, Richard C. Harrington).

## Abstract

Spiny-rayed fishes (Acanthomorpha) dominate modern marine habitats and comprise more than a quarter of all living vertebrate species^1–3^. It is believed that this dominance resulted from explosive lineage and phenotypic diversification coincident with the Cretaceous-Paleogene (K-Pg) mass-extinction event^4^. It remains unclear, however, if living acanthomorph diversity is the result of a punctuated burst or gradual accumulation of diversity following the K-Pg. We assess these hypotheses with a time-calibrated phylogeny inferred using ultraconserved elements from a sampling of species that represent over 91% of all acanthomorph families, as well as an extensive body shape dataset of extant species. Our results indicate that several million years after the end-Cretaceous, acanthomorphs underwent a prolonged and significant expansion of morphological disparity primarily driven by changes in body elongation, and that acanthomorph lineages containing the bulk of the living species diversity originated throughout the Cenozoic. These acanthomorph lineages radiated into distinct regions of morphospace and retained their iconic phenotypes, including a large group of laterally compressed reef fishes, fast-swimming open-ocean predators, bottom-dwelling flatfishes, seahorses, and pufferfishes. The evolutionary success of spiny-rayed fishes is the culmination of a post K-Pg adaptive radiation in which rates of lineage diversification were decoupled from periods of high phenotypic disparity.

## Main

The Cretaceous-Paleogene (K-Pg) mass extinction fundamentally affected the evolutionary trajectory of terrestrial vertebrates, laying the foundation for spectacular radiations of eutherian mammals, neoavian birds, amphibians, and squamates^5–12^. In contrast, the impact of the K-Pg on marine vertebrate lineages is less clear. Most marine vertebrates are spiny-rayed fishes (Acanthomorpha), which collectively represent over a quarter of all living vertebrates^3^. It is hypothesized that the mass extinction of dominant Mesozoic plankton feeders following the K-Pg relieved acanthomorph fishes from previous predation and competitive pressures, enabling their occupation of vacated niches^13^. Paleogene fossils suggest a corresponding increase in acanthomorph taxonomic diversity and morphological disparity^14^, and phylogenomic analyses have prompted the complementary proposal that the origins of many major acanthomorph lineages coincide with the K-Pg^4, 15^. However, the acanthomorph fossil record is sparse in the 20 million years around the end-Cretaceous^16^, and phylogenomic efforts to date have been limited by sampling designs that inadequately represents the group’s staggering taxonomic richness^4, 15^. These factors have hindered the resolution of the timing and patterns of acanthomorph diversification near the K-Pg; it remains uncertain if acanthomorph diversification in the Cenozoic was gradual or punctuated, and whether there is a coupling of phenotypic and lineage diversification that is considered a hallmark of adaptive radiation^17^. A well-resolved, time-calibrated, phylogeny that includes all major lineages is critical to understanding the evolutionary dynamics of spiny-rayed fishes in the wake of the K-Pg.

The largest challenge to acanthomorph evolutionary studies is inferring a phylogeny of its more than 19,450 species^3^. The resolution of relationships within the subclade Percomorpha, which contains more than 95% of all acanthomorph species, has been particularly difficult. During most of the twentieth century, inferences of acanthomorph and percomorph relationships relied on anatomical characters that resulted in largely unresolved phylogenetic hypotheses^18^. Although these early morphological investigations defined major groups of acanthomorphs, their conclusions were dramatically upended by the introduction of phylogenies inferred from a relatively small number of Sanger-sequenced mitochondrial and nuclear genes^1, 19–22^. For instance, molecular phylogenies placed the anglerfishes—long classified with the group of non-percomorph acanthomorphs that includes the economically-important cods (Gadiformes)—well within Percomorpha as the sister lineage of the pufferfishes^19^. In addition, molecular phylogenies have identified several major lineages of percomorphs that each encompass a large number of species and taxonomic families^1, 21^. As an example, one of these lineages discovered in molecular phylogenetic studies contains such disparate lineages as cichlids, blennies, guppies, flyingfishes, surfperches, and mullets^22^. Despite this progress, the interrelationships among and within the major lineages of percomorphs and acanthomorphs remain unresolved due to limited informativeness in Sanger DNA sequence datasets and limited taxonomic sampling of previous phylogenomic analyses^1, 4, 15, 19–23^.

Here, we present the results of comprehensive phylogenomic analyses and estimates of divergence times of 1,084 species representing 308 of the 337 (91.4%) acanthomorph taxonomic families. We additionally sampled nine species from Aulopiformes (lizardfishes), Myctophidae (lanternfishes), and Neoscopelidae (blackchins) to serve as outgroups. Our phylogenomic inferences are based on a DNA sequence alignment of 989 ultraconserved element (UCE) loci and our divergence time estimates were calibrated with 43 fossil constraints. We combine this new acanthomorph time tree with phenotypic data for 680 living acanthomorph species^24^ to explore the patterns of body shape diversification in spiny-rayed fishes across the K-Pg boundary.

The maximum likelihood analysis of the concatenated UCE dataset departs from previous efforts by providing confident phylogenetic resolution of nearly all sampled families of acanthomorphs and percomorphs (Fig. 1, 2, Supplementary Figs. 1-25). The UCE phylogeny differs from a phylogenomic analysis of exon capture data in the identification of Paracanthopterygii as monophyletic, the resolution of a clade containing beardfishes (*Polymixia*) and percopsiforms (troutperches, Pirate Perch, and amblyopsid cavefishes), and the placement of Beryciformes (containing Berycoidei and Holocentridae) as the sister lineage of the species-rich Percomorpha (Fig. 1, Extended Data Fig. 1)^15^. Our results are consistent with earlier molecular analyses in regard to the resolution of major percomorph clades that each include a large number of taxonomic families^1, 4, 15, 19–23^. For example, there is strong node support [Bootstrap support (BSS) = 100%] for the percomorph subclade Acanthuriformes^25^, a monophyletic group comprising more than 2,325 species, including anglerfishes, pufferfishes, butterflyfishes, and scores of other percomorph lineages that long evaded resolution in morphological and molecular analyses (Fig. 2). The relationships among the most inclusive lineages of acanthomorphs inferred from the concatenated datasets are largely corroborated in species tree estimates using coalescent methods from individual gene trees (Supplementary Fig. 26). Exceptions include the placement of codlets (*Bregmaceros*), Beryciformes, and several lineages of Eupercaria whose phylogenetic relationships have historically been difficult to resolve^1, 21^.

**Fig. 1:**
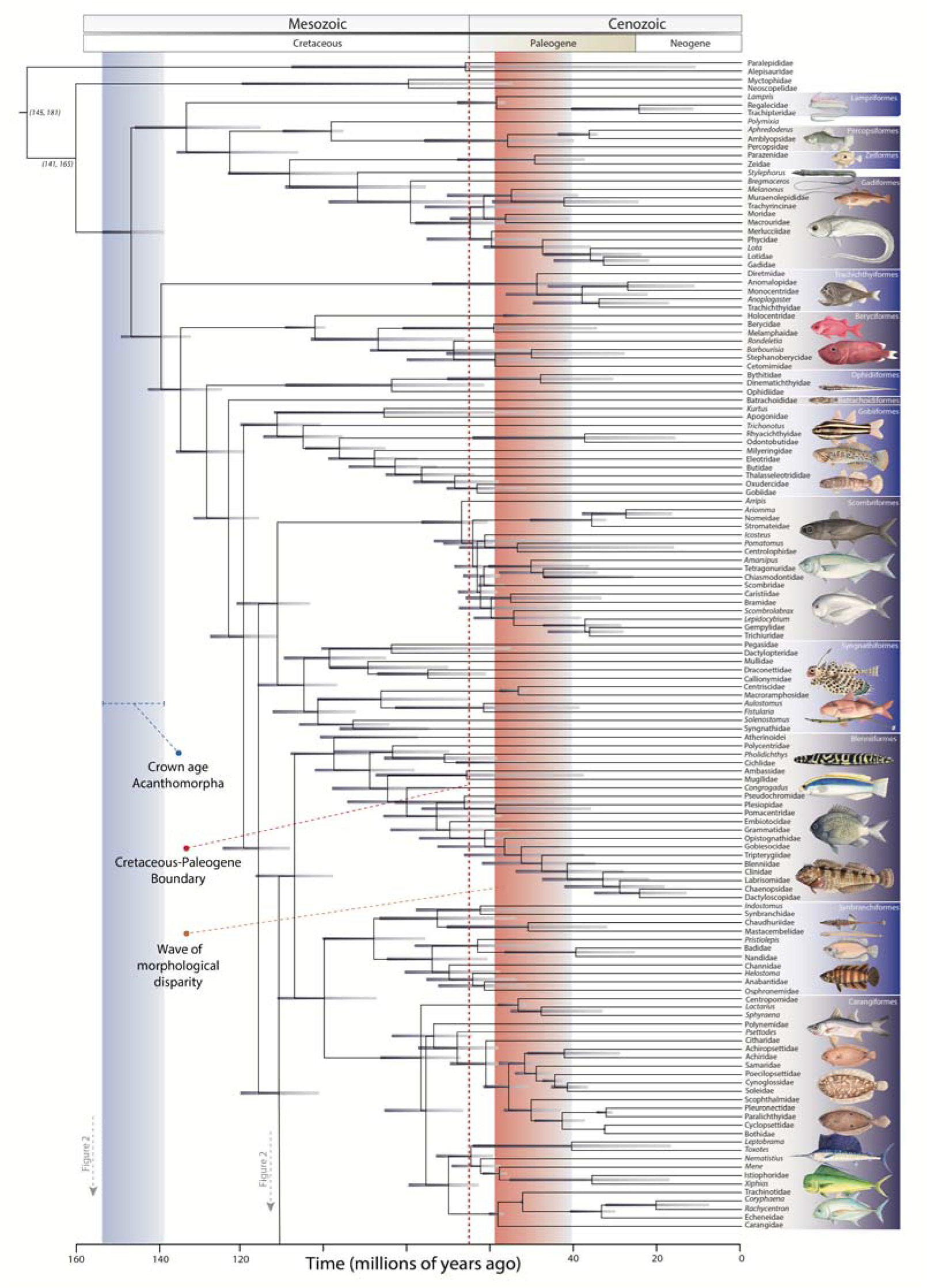
Time-calibrated phylogeny of Acanthomorpha. The phylogeny is condensed to represent taxonomic families at the tips. For families containing a single species or genus, the genus name is given at the tip. Shaded tabs to the right of taxon labels identify inclusive taxonomic orders. Horizontal gray bars at each node portray the 95% highest posterior density (HPD) of node age estimates. The blue shaded region reflects the 95% HPD of the crown age of Acanthomorpha. Maximum likelihood bootstrap support values for relationships are in Supplementary Figs. 1-25. The time-calibrated phylogeny continues in Fig. 2.

**Fig. 2:**
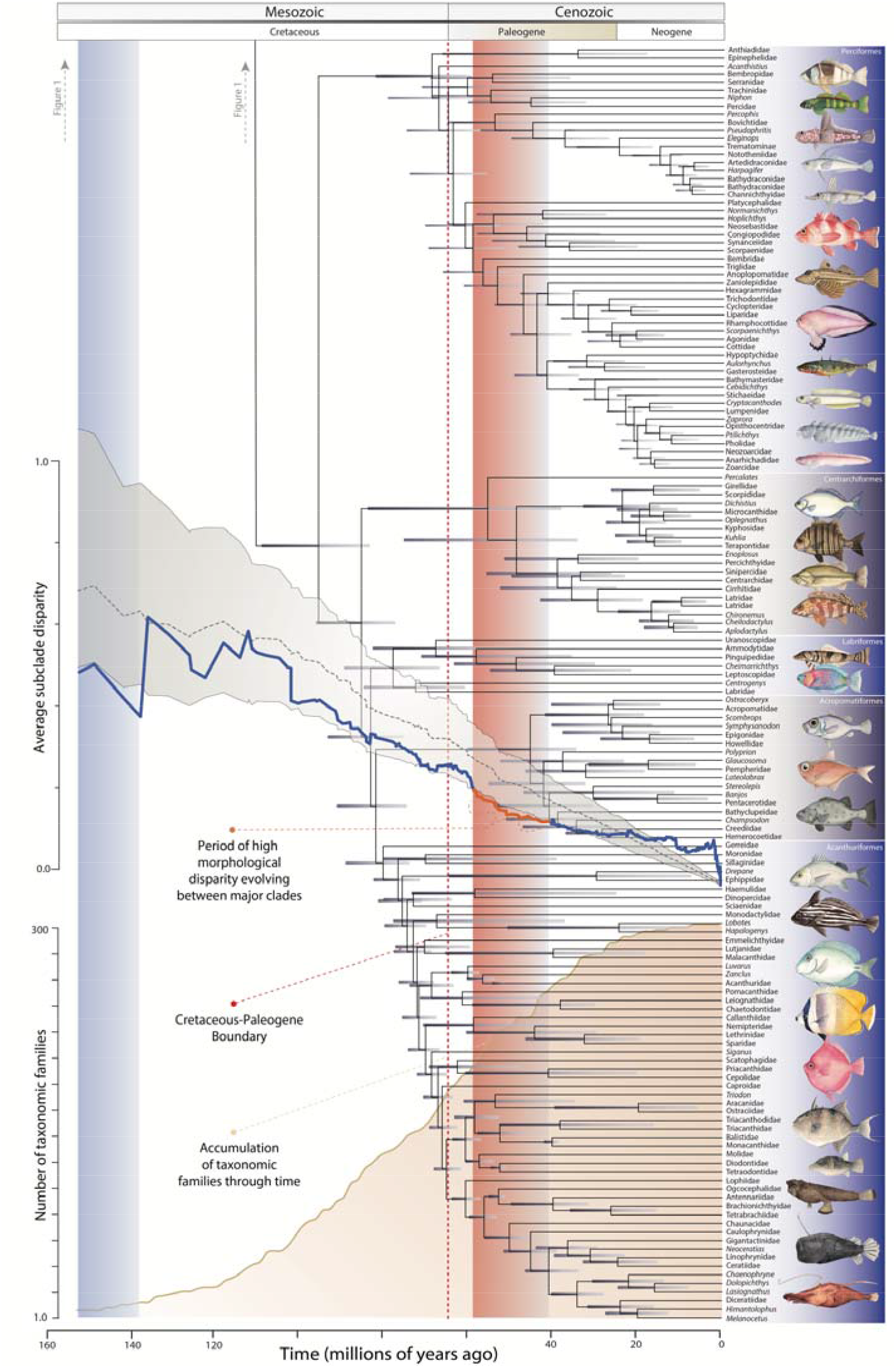
Time-calibrated phylogeny and disparity and lineage accumulation through time for Acanthomorpha. Time-calibrated phylogeny condensed to represent taxonomic families at the tips (continued from Fig. 1). The solid blue line (middle plot) shows observed morphological disparity through time (DTT) and the red portion of the line represents the significant increase in among-clade disparity in the early Eocene. The dashed black line and surrounding gray envelope represent the median DTT and 95% confidence interval under Brownian evolution, respectively. The lower plot estimates the cumulative number of taxonomic families through time. A vertical red dashed line marks the K-Pg.

In addition to yielding confident phylogenetic resolution among major lineages and taxonomic families, the maximum likelihood UCE phylogeny reveals novel relationships among some of the most scientifically interesting lineages of percomorph fishes (Figs. 1, 2, Supplementary Figs. 1-25). For example, Sanger sequencing studies led to the discovery that the enigmatic coral reef-dwelling engineer gobies (*Pholidichthys*) are the sister lineage of freshwater cichlids (Fig. 1, Supplementary Fig. 11)^1, 21, 22^. Whereas these Sanger analyses provided weak resolution beyond the monophyly of these two lineages, our UCE phylogeny resolves the freshwater tropical African and South American leaffishes (Polycentridae) as the sister lineage of the *Pholidichthys*-cichlid clade (BSS = 100%), providing an opportunity for insight into the evolution of the remarkable species richness and key morphological novelties found in cichlid fishes. We also confidently resolve the near-shore rocky reef dwelling False Scorpionfish (*Centrogenys vaigiensis*) as the sister lineage to a clade of more than 630 species of wrasses and parrotfishes (Labridae; BSS = 100%) (Fig. 2, Supplementary Fig. 21). This newly discovered relationship is not corroborated by the species tree analysis, but consistent with this discovery is that wrasses and *Centrogenys vaigiensis* share ancestral, highly modified components of the “labroid” pharyngeal jaw apparatus^22^. This result thereby reduces the number of independent evolutions of pharyngognathy, an advanced feeding mechanism that promotes trophic diversification by freeing the oral jaws from prey-processing functions^22^. The maximum likelihood phylogeny inferred from our UCE dataset provides an important framework for further exploration of the relationships among the most closely related lineages of acanthomorph and percomorph fishes.

Divergence time estimates for Acanthomorpha, inferred from relaxed molecular clock analyses calibrated with 43 well-justified fossil calibrations, allow for unprecedented resolution of the timing and tempo of family-level lineage diversification. The mean Bayesian posterior age estimates for stem lineages for 80% of living acanthomorph families post-date the K-Pg and are primarily distributed across the Paleocene through the early Miocene (∼65 – 15 Mya) (Extended Data Fig. 2). This pattern contrasts with earlier hypotheses of an explosive origin of acanthomorph familial diversity following the K-Pg mass extinction^4, 16^. Plotting the accumulation of acanthomorph family origination times reveals an extended period of lineage origination that persists for approximately 40 million years after the K-Pg (Fig. 2). Although most of the taxonomic families assigned to living acanthomorphs originated in the Paleogene, we detect no statistically supported mass extinctions or tree-wide shifts in lineage diversification rates, indicating an indiscernible effect of the K-Pg on acanthomorph lineage diversification (Extended Data Fig. 3). Moreover, stepping-stone simulations, used to evaluate the relative and absolute fit of competing diversification models to the observed time-calibrated phylogeny, strongly support a constant-rate birth-death model (Extended Data Table 1). Bayesian estimates of the evolutionary rates of individual acanthomorph lineages are similarly homogenous across the K-Pg and through the Cenozoic (Extended Data Fig. 4). We observe increases in speciation rates in only a few percomorph clades, such as branches in the phylogeny leading to Apogonidae (cardinalfishes), Pseudocrenilabrinae (most Middle Eastern and African cichlids), Chaetodontidae (butterflyfishes), the species-rich *Sebastes* (rockfishes), and Lycodinae (a lineage of eelpouts).

Although lineage diversification rate remained constant through the K-Pg boundary, acanthomorph body shapes began diversifying sharply at the start of the Paleogene into distinct regions of morphospace that correspond to iconic present-day ecomorphological types (Fig. 2, Extended Data Fig. 5, Supplementary Fig. 27). Our analyses of body shape disparity through time, repeated on a sample of 100 trees from the posterior distribution of BEAST time-trees, show that the sudden diversification of acanthomorph body shapes began an average of approximately five million years after the K-Pg (Supplementary Fig. 28). This period of high between-clade phenotypic variation persisted until the early to mid-Eocene (∼45 – 40 Mya) (Fig. 2). This pattern was not driven by a single clade, but rather occurred across all of Acanthomorpha coincident with the origin of several major lineages (Figs. 1,2, Extended Data Fig. 6a). In contrast, subsequent radiations of acanthomorphs primarily occurred within specific percomorph lineages (Extended Data Fig. 5b, Supplementary Fig. 27). These clades expanded into well recognized ecomorphological types, such as pufferfishes and allies (Tetraodontoidei), the charismatic seahorses and relatives (Syngnathiformes), a group of open ocean predators (Scombriformes), bottom-dwelling flatfishes, a large clade of deep-bodied laterally compressed reef fishes including surgeonfishes, butterflyfishes and rabbitfishes, and Perciformes, which includes many large-mouthed predators that repeatedly invaded benthic habitats (Extended Data Fig. 5, Supplementary Fig. 27).

Our estimate of the timing of acanthomorph body shape diversification at the onset of the Paleogene refines inferences from the fossil record. Acanthomorph taxonomic diversity and morphological disparity was low in the Late Cretaceous prior to the K-Pg event, after which body shape disparity and taxonomic diversity became greatly elevated^14, 16^. The timing of this diversification remained unclear due to a scarcity of deposits yielding abundant and well-preserved teleost skeletons between the Campanian-Maastricthian (∼72 Mya) in the Late Cretaceous to the Paleocene-Eocene boundary (∼56 Mya)^14, 16^. Though isolated acanthomorph otoliths are known throughout this interval, they remain little studied relative to those from Eocene and younger deposits^26^. Our analyses provide the precision lacking in the fossil record, revealing that the disparity observed in early Cenozoic fossil acanthomorphs was likely the product of an extended period of phenotypic diversification that began approximately 60 Mya and lasted approximately 15-20 Myr. The initiation of a steady accumulation of living acanthomorph families and morphological disparity following the K-Pg extends through much of the acanthomorph fossil gap (Fig. 2), demonstrating that the rise of acanthomorph diversity in the early Eocene was not punctuated^16^.

The prolonged expansion of morphological variation following the K-Pg was dominated by divergences in body elongation between clades (Extended Data Fig. 5a, Extended Data Fig. 6b,c), a phenotypic trait that has adaptive consequences. Body elongation is a primary axis of body shape variation in freshwater and marine fishes^24, 27^ that is often coupled with habitat transitions between pelagic, demersal and fully benthic habitats^28^, including in the aftermath of the K-Pg extinction^23^. Our observations of diversification in body elongation shortly after the K-Pg is therefore likely correlated with niche divergence along the benthic-pelagic axis. This pattern of morphological diversification is consistent with the classic trend in phenotypic evolution during adaptive radiation^17^, although it was not accompanied by changes in lineage diversification rate. Nonetheless, our results imply that conditions in the aftermath of the global K-Pg event promoted exceptional morphological diversification perhaps in response to the novel landscape of ecological opportunity that took shape during the Paleogene^13^.

Our results provide a new perspective to the role of the K-Pg as a catalyst of vertebrate diversification by demonstrating that its effects on acanthomorph lineage diversification rates and disparity were decoupled. Although there is no indication that acanthomorph lineage diversification rates increased near the K-Pg, much less explosively (Extended Data Fig. 3), the time following the K-Pg undoubtedly played a critical role in increasing taxonomic diversity and shaping the ecomorphological disparity of the vast majority of living spiny-rayed fishes (Fig. 2 and Extended Data Fig. 2). The approximately 15-20 Myr period following the K-Pg brought about the origin of ecomorphological types that acted as a reservoir of diversity, setting the stage for more recent, phylogenetically or geographically localized radiations, including endothermic open-ocean tunas, Antarctic notothenioids, parrotfishes on modern coral reefs, and African Rift Lake cichlids^29–32^. Though the decoupling of lineage and phenotypic diversification in Acanthomorpha contradicts commonly invoked expectations of adaptive radiation^17^, this pattern is evident in the evolutionary dynamics of other vertebrate adaptive radiations following the K-Pg, including birds and placental mammals^5, 8, 33, 34^. Our findings illuminate the origin of phenotypic diversity that characterizes the dominance of acanthomorph fishes in modern marine habitats and broadens our understanding of adaptive radiation in species-rich vertebrate lineages following the K-Pg mass extinction event.

## Methods

Detailed descriptions of all procedures are available in the Supplementary Information.

### Taxon sampling and procurement of sequence data

This study incorporates ultraconserved element (UCE) sequence data from 1,109 specimens, including nine outgroup taxa and 1,075 acanthomorph species spanning 308 recognized, taxonomic families in Acanthomorpha (Supplementary Table 1). We generated new sequence data from 647 specimens representing 628 species of acanthomorphs and six outgroups. We extracted DNA from muscle or fin tissue following the standard protocol for Qiagen’s DNeasy Blood and Tissue kits. We prepared dual-indexed DNA libraries and targeted the 1,314 UCE loci enriched by the myBaits UCE Acanthomorph kit from Arbor Biosciences^4^. Enriched libraries were sequenced on Illumina HiSeq platforms in 2 × 150 bp paired-end runs. UCE data for 360 specimens came from five previous phylogenomic studies (Supplementary Table 1), and UCE sequences for 96 species were extracted from whole genome shotgun sequence data published in the NCBI GenBank^35, 36^

### Data processing and phylogenetic analyses

We used the PHYLUCE^35, 36^ computer package to process raw read data, conduct de novo assembly and construct alignments of UCE loci. We generated two separate alignments of UCE loci present in at least 75% of the samples: one consisting of all 1,109 specimens (1,084 species) and another including only a subset of the 702 species that overlapped with the taxon sampling of a previously published morphological dataset. The 1,084-and 702-taxon alignments consisted of 987 loci (383,250 bp) and 989 loci (499,957 bp), respectively.

Using these two 75% complete data matrices, we conducted multiple phylogenetic analyses using maximum likelihood methods in IQ-TREE v. 1.7^37^ and RAxML-ng v. 0.9.0^38^. We topologically constrained the divergence time analyses represented in Figure 1 with phylogenies inferred in IQ-TREE using single-partition alignments of the 702 taxa and 1,084 taxa datasets. Both of these tree searches assumed the GTR+Gamma model of molecular evolution and used ultrafast bootstrap approximation to generate 1,000 bootstrap replicates and 1,000 replicates of the Shimodaira-Hasegawa approximate likelihood ratio test (SH-aLRT). We used the program TOPD_v4.6^39^ to ensure the phylogenies inferred using different partitioning schemes and maximum likelihood programs presented similar tree topologies.

We also used IQ-TREE’s ModelFinder Plus and ultrafast bootstrap approximation options to infer maximum likelihood gene trees for each UCE locus. In TreeShrink^40^, we used a false positive tolerance rate parameter (alpha) of 0.05 to identify and remove potentially aberrant sequences from the single-gene alignments. We re-aligned filtered alignments using MAFFTv7.130b^41^ and used them to repeat inference of gene trees in IQ-TREE. We employed the resulting gene trees to generate a summary species tree in ASTRAL-III v5.6.3^42^.

We applied Bayesian methods to understand the degree of topological discordance along the backbone of the Acanthomorph tree. We inferred gene trees for UCE loci with Bayesian methods for an 82-taxon alignment that represented the major acanthomorph subclades and was generated using MAFFT. We ran these tree searches in MrBayes v. 3.2.7^43^ for 2 million generations, assuming a GTR+Gamma model of molecular evolution. Using an alpha value of 1.0, we estimated genome-wide concordance factors in BUCKy^44^ as a measure of the amount of topological discordance between these Bayesian gene trees.

### Divergence time estimation

We estimate divergence times for the 702-and 1,084-taxa phylogenies in BEAST v.2.5^45–47^ using reduced datasets of randomly subsampled UCE loci^29, 48^. For both phylogenies, we repeated dating analyses on three different alignments of 30 loci. For each of these alignments, we accounted for site-specific variation in evolutionary patterns by selecting the best-fit partitioning schemes using PartitionFinder2 v. 2.1.1^49^. We performed at least three replicate analyses for each 30-locus dataset. We ran all BEAST analyses under a relaxed lognormal clock model and a birth-death tree model, using the IQ-TREE phylogenies inferred from the single-partition, 702-taxa and 1,084-taxa alignments as topological constraints. We used forty-three fossil constraints (detailed in the Supplementary Information) to assign minimum age priors and ran BEAST analyses for a minimum of 200 million generations after discarding a burnin of 200 million iterations. We used Tracer v.1.7.1^50^ to assess convergence of parameters across replicate MCMC chains and ensure there were no directional trends in parameter estimates. For each set of random loci, we combined replicate analyses in LogCombiner and constructed maximum clade credibility (MCC) trees using TreeAnnotator in BEAST v1.8.4^51^, with summarized node heights rescaled to reflect the posterior median heights. MCC trees for the 702-taxon datasets were summarized from 10,000 post-burnin, randomly sampled trees, while MCC trees for the 1,084-taxon datasets were summarized from 1,200 such trees.

### Diversification rate analyses

We removed all outgroup and duplicate taxa from the 1,084 taxa maximum clade credibility tree displaying the highest effective sample size (ESS) and used this pruned tree for all diversification rate analyses. We used TESS^52^ to conduct stepping-stone simulations that estimated the marginal likelihoods of eight birth-death models, allowing us to calculate Bayes Factors and assess the competing models’ relative and absolute fits to the time-calibrated phylogeny of Acanthomorpha. We examined tree-wide speciation, extinction, and net-diversification rates using TESS’s CoMET model. For this analysis, we accounted for our incomplete sampling, specified a uniform sampling strategy, and ran three replicate reversible-jump MCMC chains until ESS values were ≥200. CoMET runs were checked for within-and between-analysis convergence of the diversification rate parameters as described in the Supplementary Information. Using the same time-calibrated phylogeny, we also inferred rate heterogeneity across lineages using BAMM v.2.5.0^53^. BAMM analyses accounted for the incomplete taxon sampling of taxonomic families and other major representative clades and used empirically determined rate priors identified by the R package BAMMtools v.2.1.6^54^. We ran 23 MCMC chains with different combinations of parameters in BAMM, each for 100 million generations, and we identified the rate shifts along specific branches that were predicted by at least sixteen of these analyses.

### Phylogenetic comparative methods

We pruned a body trait dataset of teleost fishes^24^ to include maximum body depth, maximum fish width, head depth, lower-jaw length, mouth width, minimum caudal-peduncle depth, and minimum caudal-peduncle width for the 680 species that matched the species represented in the 702-taxon UCE phylogeny. To correct for body size, we regressed log-transformed trait values against log-transformed standard fish lengths and calculated phylogenetic residuals in PHYTOOLS^55^. We assessed disparity through time using GEIGER^56^ and the time-calibrated IQ-TREE phylogeny. We repeated disparity through time analyses on 100 randomly sampled time trees from the posterior distribution generated in BEAST, and summarized the mean and 95% confidence interval for the 1.0 My time interval after the K-Pg during which average subclade disparity first dropped below the value simulated under Brownian evolution, as well as the proportion of trees through time that strayed from the Brownian prediction. We tested the sensitivity of disparity through time analyses to the exclusion of body shape traits and of major clades that arose immediately around the K-Pg boundary. Finally, we used PHYTOOLS to visualize body shape morphospace using principal component analysis (PCA) and to generate phenograms that visualize the evolutionary histories of the following seven major lineages: Carangiformes, Lophioidei, Perciformes, Scombriformes, “Squamipinnes” (the clade in Supplemental Fig. 23 defined by *Chaetodon kleinii* and *Luvarus imperialis*), Syngnathiformes and Tetraodontoidei.

### Code availability

Computer code for analyses can be obtained from the authors upon request.

### Data availability

Alignments used to perform the analyses in this manuscript are available upon request. NCBI BioSample Accession numbers corresponding to sequence data are listed in Supplementary Table 1.

## Acknowledgements

We thank Julie Johnson for the fish illustrations in Figs. 1,2 and Supplementary Fig. 29, and the numerous undergraduate and graduate researchers from the University of California, Davis and Clemson University that helped collect morphological data. Portions of this research were conducted with high performance computational resources provided by Louisiana State University (http://www.hpc.lsu.edu). We are grateful to the ichthyology curators and staff of the following collections for granting access to the tissues and specimens that made this study possible: Smithsonian National Museum of Natural History (Washington, D.C.), University of Florida Museum of Natural History (Gainesville), Scripps Institution of Oceanography (La Jolla), South African Institute for Aquatic Biodiversity (Grahamstown), Southeastern Louisiana University Museum of Biology (Hammond), American Museum of Natural History (New York), Australian Museum (Sydney), Academy of Natural Sciences (Philadelphia), Field Museum of Natural History (Chicago), California Academy of Sciences (San Francisco), Cornell University Museum of Vertebrates (Ithaca), University of Tennessee David A. Etnier Ichthyological Collection (Knoxville), Burke Museum of Natural History and Culture (Seattle), Academia Sinica, Biodiversity Research Museum (Taipei), Natural History Museum and Institute (Chiba), Australian National Fish Collection (Hobart), Kyoto University Museum, Mie University Fish Collection of the Fisheries Research Laboratory (Shima), Hokkaido University Museum (Sapporo), Illinois Natural History Survey (Champaign), Kagoshima University Museum (Korimoto), University of Kansas Biodiversity Institute (Lawrence), Natural History Museum of Los Angeles County, Universidade Estadual Paulista (São Paulo), Louisiana Museum of Natural History (Baton Rouge), Harvard Museum of Comparative Zoology (Cambridge), Museo Nacional de Ciencias Naturales (Madrid), Museo Nacional de Historia Natural (Santiago), North Carolina Museum of Natural Sciences (Raleigh), National Museum of Natural History (New Delhi), Museum of New Zealand Te Papa Tongarewa (Wellington), Museum Victoria (Melbourne), National Museum of Nature and Science (Tokyo), Museums and Art Galleries of the Northern Territory (Darwin), University of Tokyo Ocean Research Institute, Queensland Museum (Brisbane), Royal Ontario Museum (Toronto), Seikai National Fisheries Research Institute (Nagasaki), Universitetsmuseet i Bergen (Hordaland), University of Copenhagen Zoological Museum, and Peabody Museum of Natural History (New Haven).

## Author contributions

The project was conceived and designed by TJN, AG, RCH, JRG, BCF, and PCW. AG, RCH, JRG, MAC, JCB, WTM, WBL, BCF, PFC and TJN organized and executed the collection of UCE sequence data. SAP, SFT, and PCW coordinated and collected morphological data. AG, RCH, BCF, and AD carried out phylogenetic analyses. AG and RCH performed phylogenetic divergence dating analyses, and AG performed analyses of lineage diversification rates. EDB and PCW performed morphological comparative analyses. AG, RCH, AD, and TJN wrote the first draft of the manuscript. All authors contributed to the writing of the final draft of the manuscript.

## Additional information

**Supplementary Information** is available for this paper.

Correspondence and requests for materials should be addressed to. Reprints and permissions information is available at www.nature.com/reprints.

## Extended data table and figure legends

The Extended Data table and figures (with legends) are available in the document labelled ExtendedDataTableAndFigs.pdf, but legends are also provided below.

**Extended Data Table 1:**
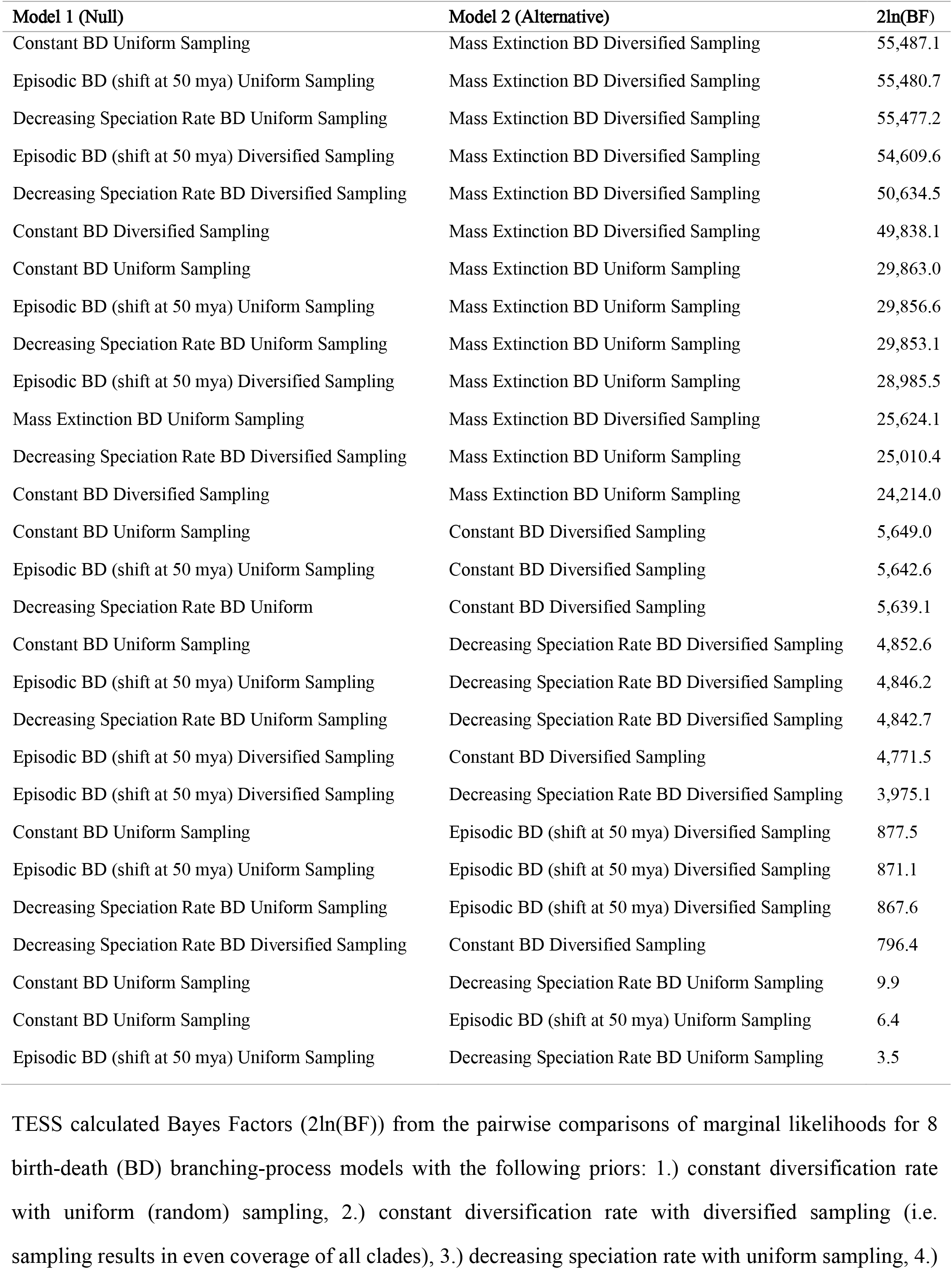
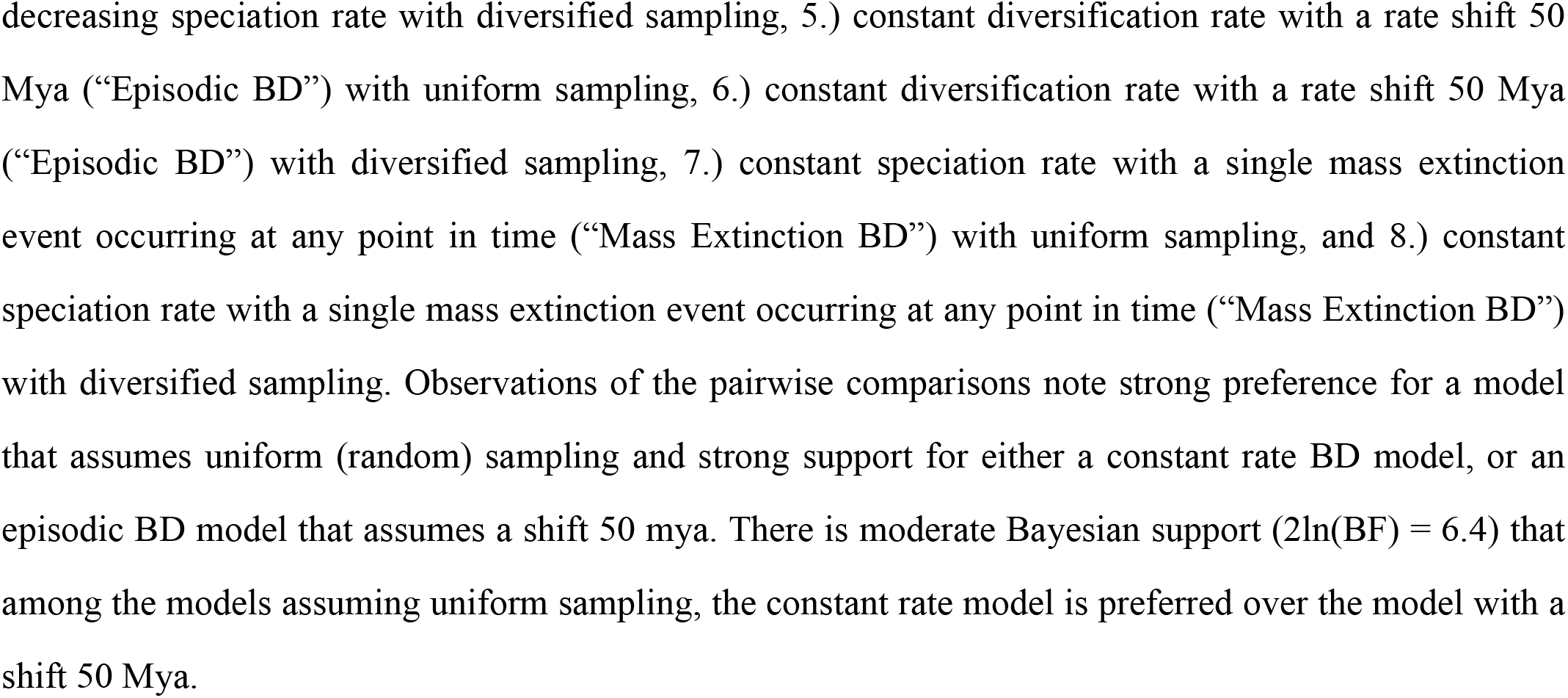
Pairwise comparisons of 8 birth-death (BD) branching process models. TESS calculated Bayes Factors (2ln(BF)) from the pairwise comparisons of marginal likelihoods for 8 birth-death (BD) branching-process models with the following priors: 1.) constant diversification rate with uniform (random) sampling, 2.) constant diversification rate with diversified sampling (i.e. sampling results in even coverage of all clades), 3.) decreasing speciation rate with uniform sampling, 4.) decreasing speciation rate with diversified sampling, 5.) constant diversification rate with a rate shift 50 Mya (“Episodic BD”) with uniform sampling, 6.) constant diversification rate with a rate shift 50 Mya (“Episodic BD”) with diversified sampling, 7.) constant speciation rate with a single mass extinction event occurring at any point in time (“Mass Extinction BD”) with uniform sampling, and 8.) constant speciation rate with a single mass extinction event occurring at any point in time (“Mass Extinction BD”) with diversified sampling. Observations of the pairwise comparisons note strong preference for a model that assumes uniform (random) sampling and strong support for either a constant rate BD model, or an episodic BD model that assumes a shift 50 mya. There is moderate Bayesian support (2ln(BF) = 6.4) that among the models assuming uniform sampling, the constant rate model is preferred over the model with a shift 50 Mya.

**Extended Data Fig. 1:**
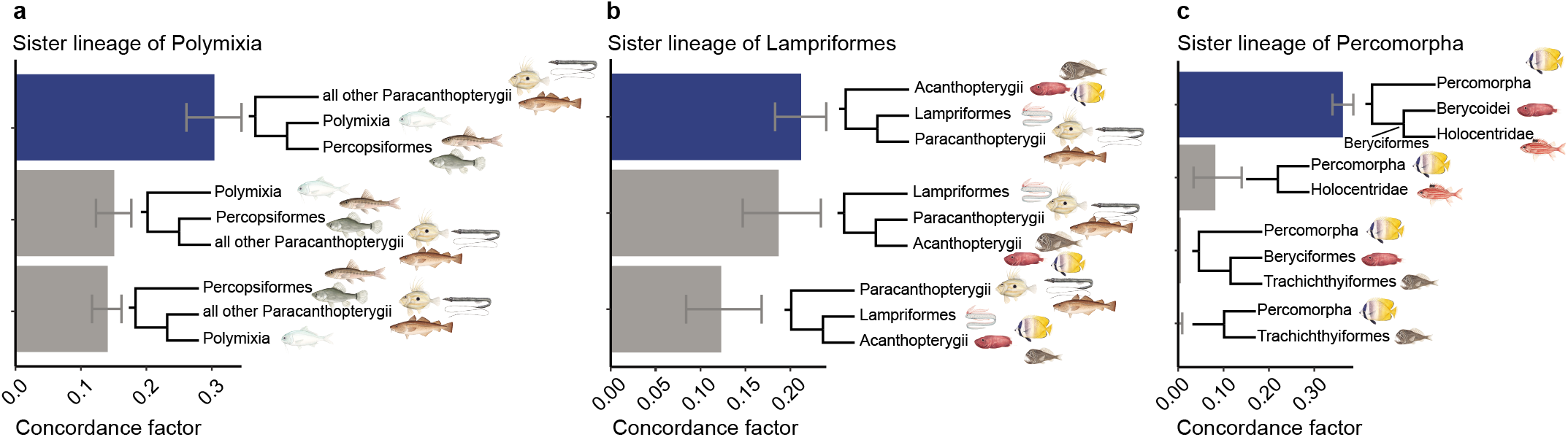
Bayesian concordance analyses used to compare alternative phylogenetic hypotheses concerning the sister taxa of 3 major acanthomorph lineages. BUCKy-inferred genome-wide concordance factor estimates, with 95% confidence intervals, representing the proportion of gene trees exhibiting alternative topologies among major acanthomorph lineages. Blue bars reflect the concordance factor for the topology inferred in our concatenated analyses (see Figs. 1,2) and gray bars represent concordance factors of alternative topologies not observed in the concatenated phylogeny. **a,** Gene tree concordance for Percopsiformes, Polymixia, and all other Paracanthopterygii; **b,** Lampriformes, Paracanthopterygii, and Acanthopterygii; **c,** Alternative sister lineages of Percomorpha.

**Extended Data Fig. 2:**
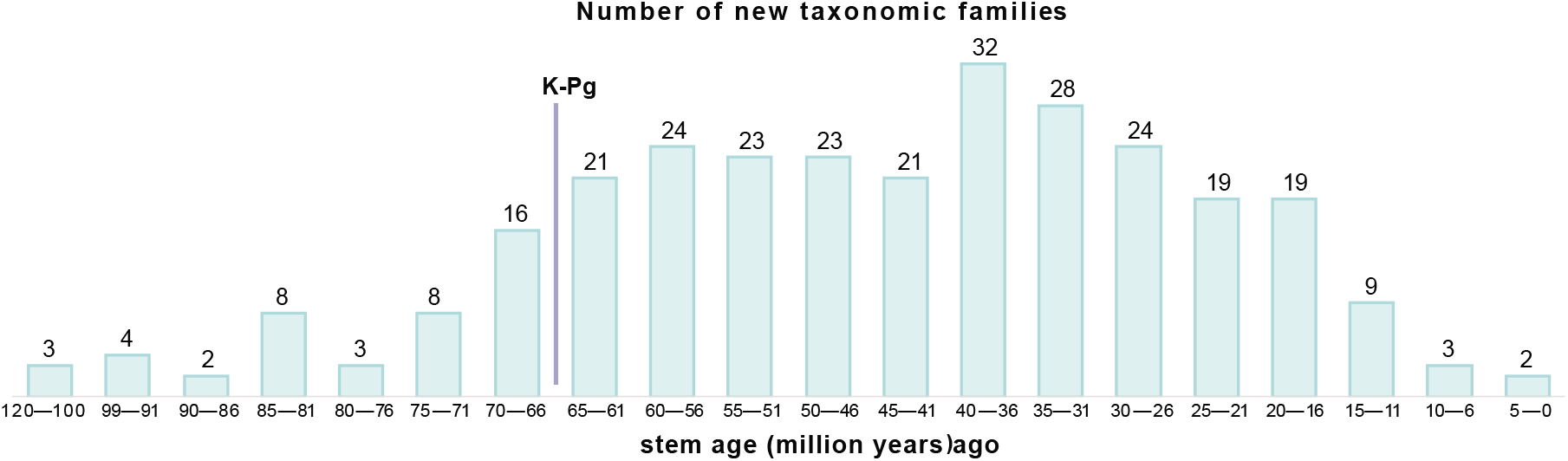
Acanthomorphs underwent a long, steady period of increased taxonomic family origination beginning at the K-Pg boundary. The bar plot of the estimated number of new taxonomic families originating during 5 million year bins. Note that the amounts of time represented by the first two bars are greater than the 5 million years represented by all other bars.

**Extended Data Fig. 3:**
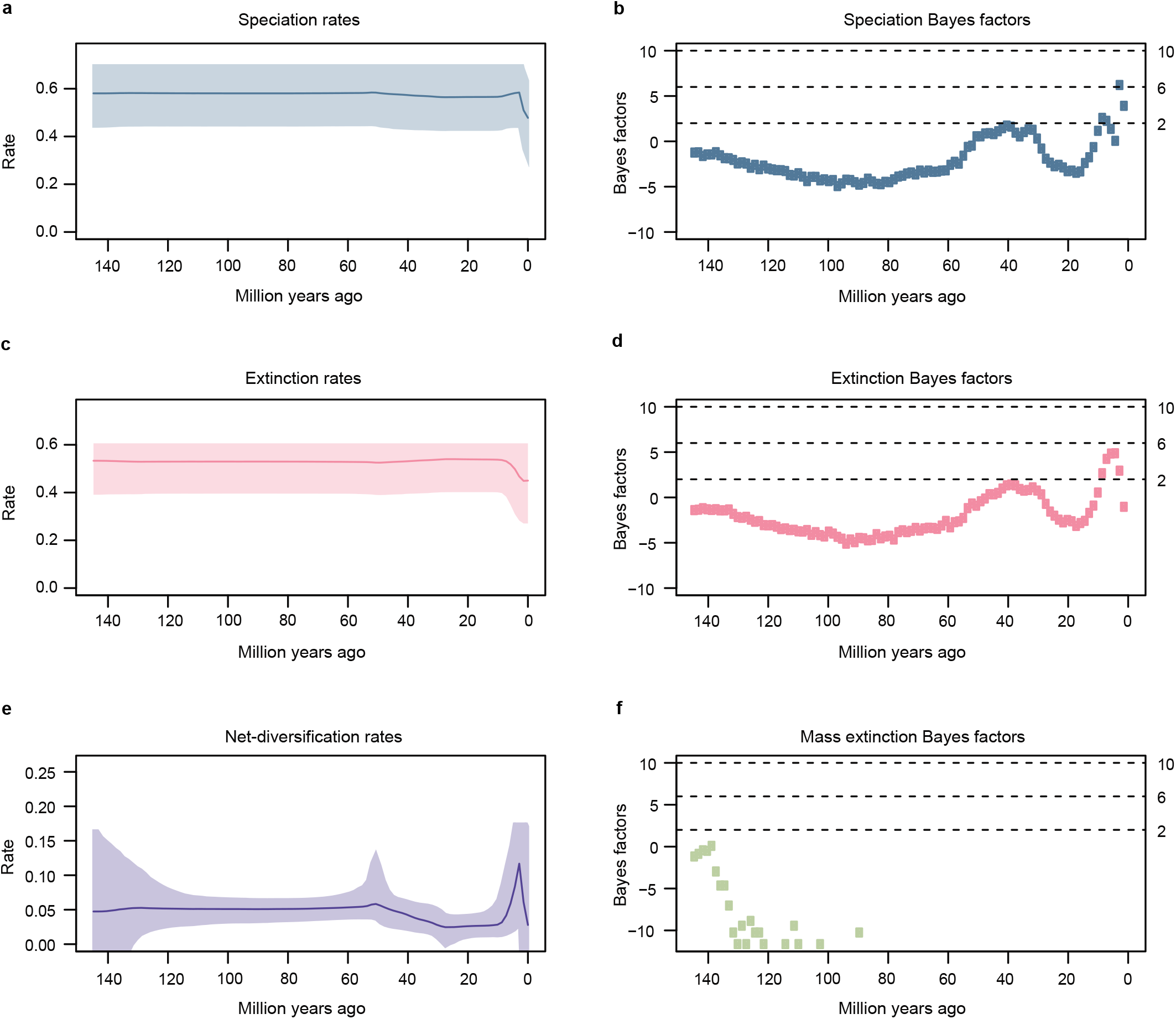
TESS-CoMET analyses indicate constant tree-wide diversification rates through most of the history of Acanthomorpha. In **b, d,** and **e,** horizontal dashed lines and the right-hand y-axis mark statistical support cutoffs for rate shifts, with 2 ≤ BF < 6 considered to be low support, 6 ≤ BF < 10 considered to be moderate support and ≥10 considered to be high support. The rate shifts observed in **a, c,** and **e** over the last 10 million years are likely an artifact of the CoMET model (see **Supplementary Information** for further discussion). **a,** Speciation rate estimates through time for Acanthomorpha. **b,** Bayes factors (BF) support for a shift in speciation rate at every 1 Myr time period. **c,** Extinction rates through time for Acanthomorpha. **d,** Bayesian support (in Bayes factors) for a shift in extinction rate at every 1 Myr time period. **e,** Net-diversification (speciation minus extinction) rates through time for Acanthomorpha. Though this plot suggests that there is a small shift in net-diversification rate ∼50 Mya, there is no statistical support for such a shift (see **b** and **d**). **f,** There is no statistical support (in Bayes factors) for a mass extinction event in Acanthomorpha at any 1 Myr time period.

**Extended Data Fig. 4:**
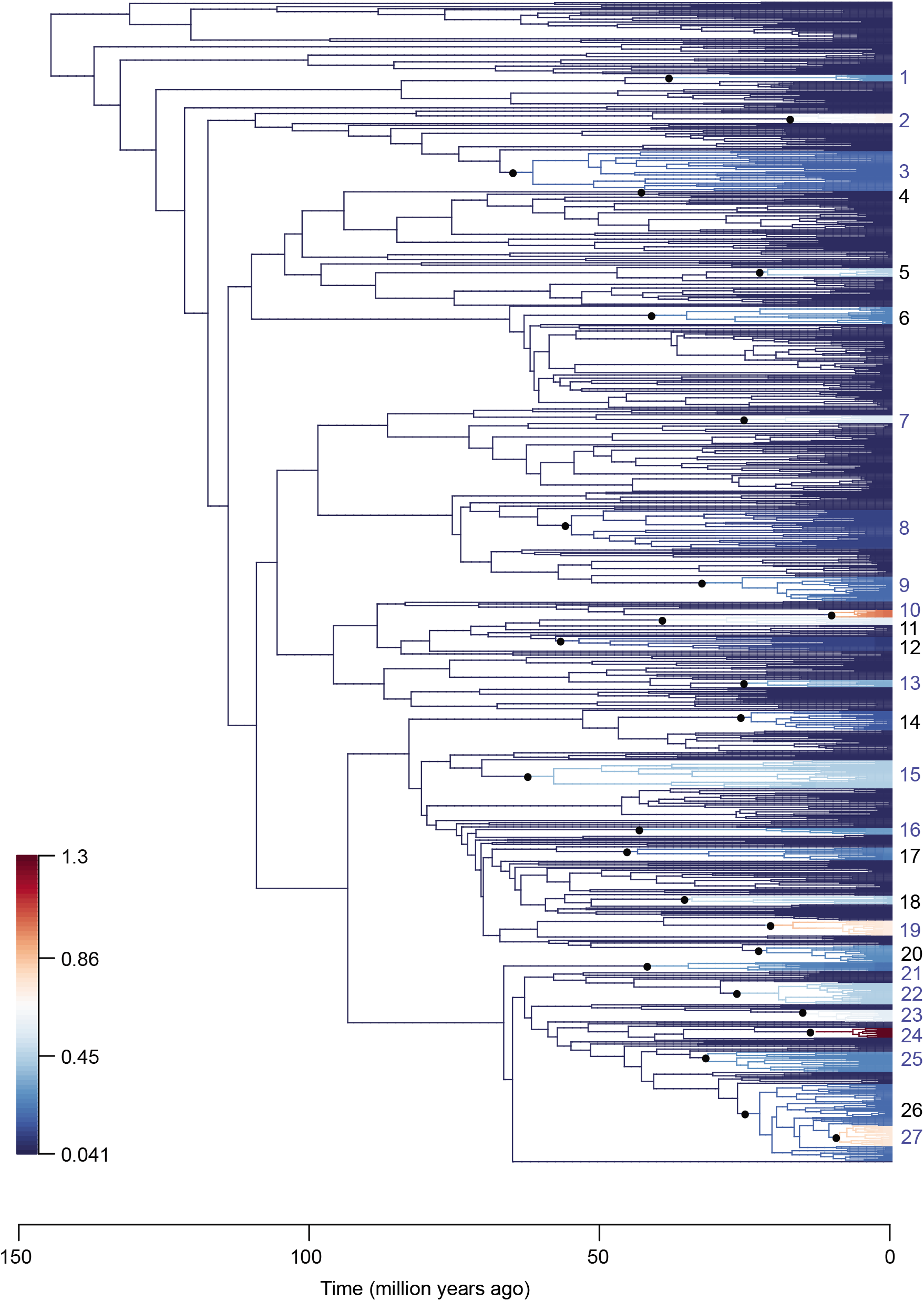
Visual summary of shifts in speciation rates inferred using BAMM. The configuration presented here is the maximum shift credibility (MSC) configuration for an analysis that expected 15 rate shifts under the prior. Darker pink colors depict relatively fast rates, while darker green colors depict relatively slow rates. Rate shifts along branches are denoted with filled, black circles and assigned identifying numbers. Shifts in speciation rate are estimated to have occurred on the branches leading to the following 27 clades: 1.) Dinematichthyidae, 2.) Apogonidae (to the exclusion of Pseudamia), 3.) Gobiidae and Oxudercidae, 4.) Solenostomus, 5.) Parupeneus and Pseudopeneus (in Mulllidae), 6.) Ariomma, Nomeidae and Stromateidae, 7.) Mastacembelidae, 8.) the clade defined by Scophthalmidae and Soleidae, 9.) Carangidae (to the exclusion of Seriola), 10.) Pseudocrenilabrinae (in Cichlidae), 11.) Pomacentridae, 12.) the clade defined by Gobiesocidae and Dactyloscopidae, 13.) Poeciliidae, 14.) the clade defined by Scorpididae and Terapontidae, 15.) Labridae, 16.) Sciaenidae, 17.) the clade defined by Nemipteridae and Sparidae, 18.) Tetraodontidae, 19.) Chaetodontidae, 20.) Acanthuridae (to the exclusion of Prionurus and Naso), 21.) Anthiadinae and Epinephelidae, 22.) darters (Etheostomatinae), 23.) the clade defined by “Nototheniidae” and Channichthyidae, 24.) Sebastes, 25.) the clade defined by Trichodontidae and Psychrolutidae, 26.) the clade defined by Stichaeidae and Zoarcidae and 27.) the clade defined by Bothrocara and Lycodes concolor (in Zoarcidae). Shifts labelled with blue rather than black numbers are estimated to have occurred by 16 of the 23 BAMM analyses conducted in this study. Although shift number 20 and shift number 26 are labelled in black, the vast majority of BAMM analyses predicted a rate shift in branches leading to slightly more inclusive clades (specifically Acanthuridae minus Naso, and all of Lycodinae, respectively).

**Extended Data Fig. 5:**
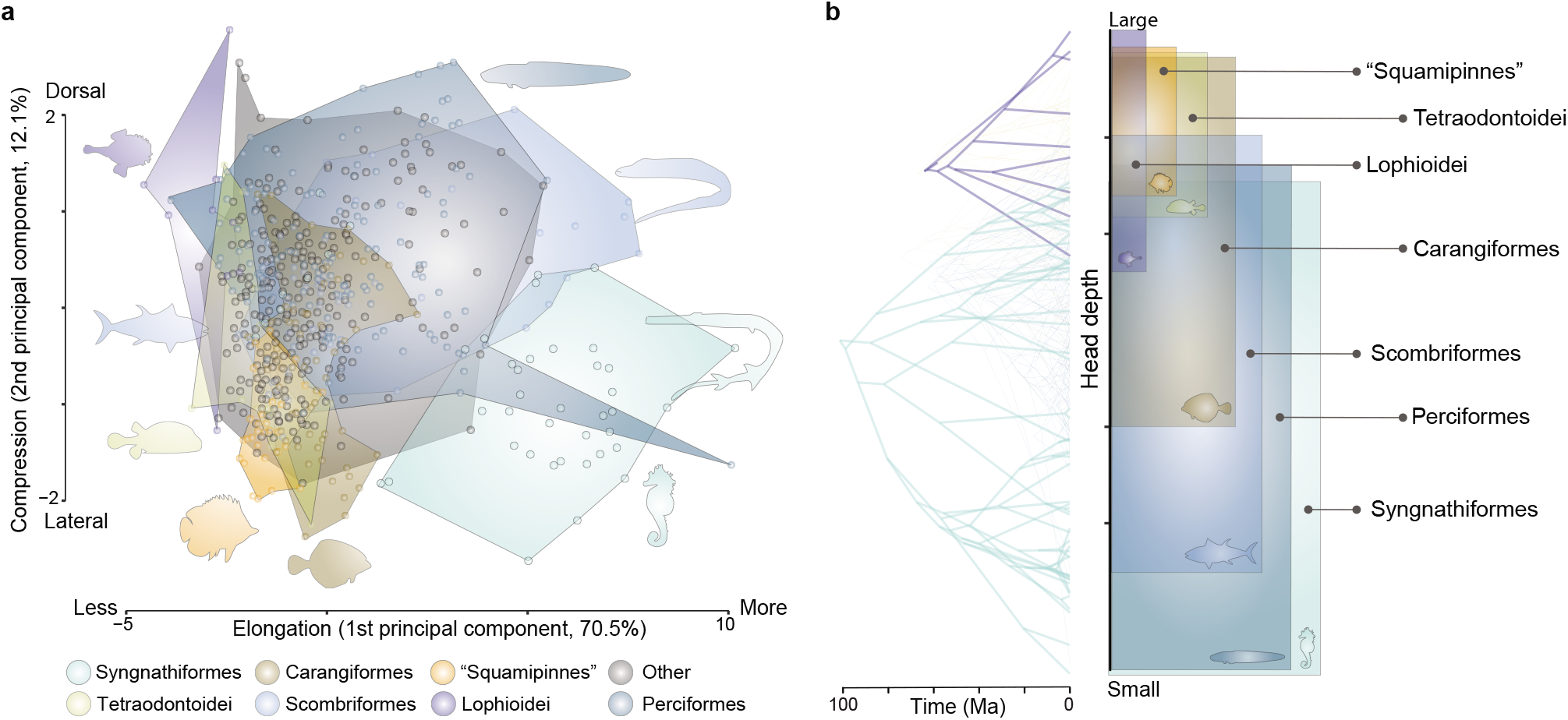
Changes in body elongation and compression. Patterns of acanthomorph morphological disparification. **a,** The first two principal components of morphospace (elongation and compression), with seven major acanthomorph lineages color-coded by taxon. The proportion of variance explained by PC1 is 70.5% and by PC2 is 12.1%. **b,** Phenogram depicting the evolutionary history of head depth—which often reflects body elongation—across seven major lineages that arose around the K-Pg. Note that all major lineages except Syngnathiformes originate near the K-Pg boundary, but “Squamipinnes”, Lophioidei and Tetraodontoidei have markedly different ancestral trait values from Carangiformes, Scombriformes and Perciformes.

**Extended Data Fig. 6:**
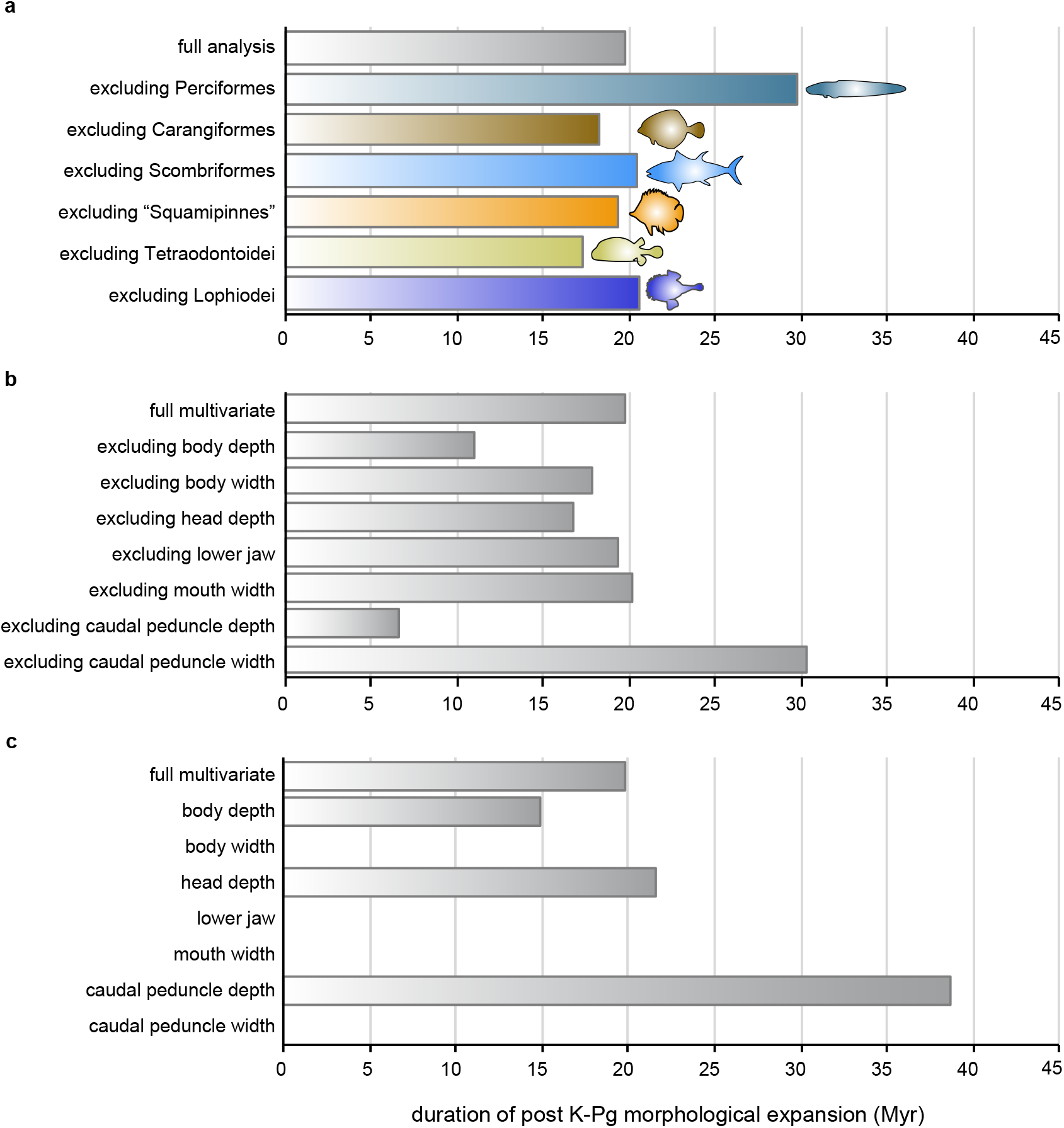
Robustness of DTT analyses. Effect of individual lineages or body shape traits on the length of the estimated period of between-clade morphological expansion after the K-Pg (i.e., the period in which the observed disparity falls below that expected from a Brownian motion process). There is no single major lineage driving the extended duration of this period of phenotypic diversification, but the body shape traits related to elongation (body depth, head depth, and caudal peduncle depth) have more pronounced contributions to this pattern than the other body shape traits. **a,** Duration of the period of morphological expansion when a single major lineage is excluded from the analysis. **b,** Duration of the period of morphological expansion when data for one of the seven body shape traits in the original analysis is excluded. **c,** Duration of the period of morphological expansion when the DTT analysis is based on a single body shape trait.

## 1. Supplementary Methods

### a. Taxon sampling

The phylogenetic analyses included UCE sequence data sampled from 1,109 specimens that include 1,084 species. For this study, we collected new UCE sequence data from 653 specimens. We retrieved UCE loci from the remaining 456 specimens at NCBI from previous studies of acanthomorph phylogeny^4, 29, 48, 57, 58^ or from whole genomic data (Supplementary Table 1). This study included representatives of all currently recognized acanthomorph taxonomic orders, 307 of the 331 (92.7%) described taxonomic families of Acanthomorpha and 276 of the 290 (95.2%) families of Percomorpha^3^. Based on previous phylogenetic analyses^15, 20^, we sampled two species of Aulopiformes and seven species of Myctophiformes as outgroups for the phylogenetic analyses: *Notolepis coatsi*, *Alepisauraus ferox*, *Scopelengys tristis*, *Neoscopelus macrolepidotus*, *Neoscopelus microchir*, *Dasyscopelus selenops*, *Benthosema glaciale*, *Notoscopelus caudispinosus*, and *Ceratoscopelus warmingii*. We sampled phenotypic data from 680 of the 1,075-acanthomorph species sampled in the UCE inferred phylogeny. We inferred phylogenies from the UCE dataset for an inclusive taxon sampling of 1,084 species and for a set of 702 species that included the 680 for which corresponding morphological data was obtained. We refer to these UCE datasets and resulting phylogenies as the 1,084 and 702 species datasets and trees, respectively.

### b. Library preparation, target enrichment, and sequencing

We isolated DNA from tissue biopsies using Qiagen DNeasy Blood and Tissue kits following the manufacturer’s protocol. We used a Qubit fluorometer (Life Technologies) to quantify 1-2 uL of all DNA extractions. We sheared approximately 500 ng of genomic DNA from each sampled specimen using a QSonica Q800R3 sonicator to obtain fragment sizes between 300-600 bp. For library preparation and target enrichment of UCE loci, we followed protocols described in a previous study^4^ using Kapa HyperPrep kits (Kapa Biosystems) and Illumina TruSeq iTru5 and iTru7 adapters for dual indexing of genomic libraries^59^. We used a bait set from Arbor Biosciences designed to target 1,314 UCE loci in acanthomorph fishes for target-enrichment of UCE loci^4^. Enriched libraries were sequenced using 150 bp paired-end sequencing on Illumina HiSeq platforms.

### c. Assembly and alignment construction

To process raw read data, construct assemblies, identify UCE sequences, and construct alignments for each UCE locus, we used the PHYLUCE computer package^35, 36^, which uses parallel wrapper functions around external programs to perform tasks required to process high-throughput DNA sequence data. We used the illumiprocessor parallel wrapper program (https://github.com/faircloth-lab/illumiprocessor) to implement Trimmomatic^60^, which removes adapter index sequences and low-quality bases from demultiplexed raw reads. We constructed assemblies for each sample using *trinity* v.r2013-02-25^61^ and used the PHYLUCE program phyluce_assembly_match_contigs_to_loci.py to identify which assembled contigs represented UCE sequences by matching them to a FASTA file containing the UCE bait sequences.

To supplement these sequence data, we extracted UCE loci from existing genomes of 96 species (Supplementary Table 1). We downloaded genomes from NCBI and converted to 2bit format using faToTwoBit from the UCSC Genome Browser Downloads (https://hgdownload.cse.ucsc.edu/admin/exe/). Then, we located UCE loci in each genome assembly by running the PHYLUCE program run_multiple_lastzs_sqlite.py with default parameters and the FASTA file containing the UCE bait sequences. We extracted from each assembly identified UCE loci along with ± 500 bp of sequence from each flank using phyluce_probe_slice_sequence_from_genomes.py with default parameters. We used PHYLUCE to run match_contigs_to_loci.py against the extracted loci, before the UCE data harvested from genome assemblies were integrated with the UCE loci enriched from libraries.

We integrated these two data sets and used PHYLUCE programs to infer a separate set of alignments and phylogenies for the 702 species including the sampling of morphological data and the full 1,109 specimen sampling using the same protocols. We generated alignments for each UCE locus using MAFFT v7.130b^41^ and trimmed with trimAL v1.4.rev^62^. For further downstream phylogenetic analyses, we retained UCE loci that included at least 75% of the total specimens in both the 1,084 and 702 species datasets.

### d. Phylogenetic analyses

We inferred phylogenies for both the 1,084 and 702 species datasets using maximum likelihood methods as implemented in IQ-TREE v. 1.7^37^ and RAxML-ng 0.9.0^38^. In IQ-TREE, we ran tree searches using ultrafast bootstrap approximation until 1,000 bootstrap replicates and 1,000 replicates of the Shimodaira-Hasegawa approximate likelihood ratio test (SH-aLRT) were generated^63^. Phylogenies inferred in IQ-TREE using the unpartitioned alignments described above served as the topological constraints for subsequent divergence time analyses. Maximum likelihood tree searches conducted in IQ-TREE used the ModelFinderPlus^64^ option, which applies a relaxed hierarchical clustering search algorithm to determine partitioning schemes and models of molecular evolution.

Before conducting analyses in RAxML-ng v0.9.0^38^, we determined partitioning schemes using PartitionFinder2 v2.1.1^49^, with units of partitioning based on individual UCE loci. We identified these partitioning schemes using a relaxed hierarchical clustering search algorithm with a maximum cluster size set to 1000 data blocks, the Bayesian Information Criterion method for partition model comparison, and GTR+Gamma as the model of molecular evolution. We removed all fully undetermined columns from our partitioned, concatenated alignment using RAxML-ng, compressed the alignment, and converted to binary. We performed maximum likelihood tree searches on these alignments using the extended majority rule criterion method implemented with the autoMRE command to continue bootstrapping until convergence.

We compared the tree topologies inferred in the concatenated analyses described above using the program TOPD_v4.6^39^. Phylogenies based on the 1,084 species dataset (IQ-TREE single-partition, IQ-TREE multiple partitions, and RAxML-ng multiple partitions) were compared to one another, and separate analyses compared the three phylogenies inferred from the 702 species dataset. We utilized TOPD to calculate partition metrics, i.e., TOPD’s “Split” method^65^, and path length metrics, i.e., TOPD’s “Nodal” method^66^. We produced lists of taxa with different phylogenetic positions among the compared trees using TOPD’s “Disagree” method, which removes a single taxon at each iteration and calculates the reduction in trees’ split distance.

We inferred a summary species tree in ASTRAL-III v5.6.3^42^, using maximum likelihood gene trees for each UCE locus generated with IQ-TREE v 1.6.12. For individual gene tree inference, we applied the ModelFinder Plus^64^ option to determine the optimal model of molecular evolution as well as the ultrafast bootstrap approximation with a maximum of 10,000 bootstrap iterations. We used TreeShrink^40^ to identify abnormally long branch lengths in the individual UCE gene trees and to minimize the impact of misspecified sequences. We used a false positive tolerance rate parameter (alpha) of 0.05 in our TreeShrink filtering to automate the removal of potentially aberrant sequences from the UCE alignment. We re-aligned the TreeShrink-pruned alignments using MAFFT v7.130b^41^ and re-inferred gene trees using IQ-TREE, as above. We used these gene trees to infer a species tree in ASTRAL-III.

### e. Concordance factor analyses

We used BUCKy^44^ to estimate genome-wide concordance factors, which allows the evaluation of phylogenetic discordance among individual UCE gene trees. We compared alternative phylogenetic hypotheses among the major lineages of Acanthomorpha using these concordance factors. BUCKy estimates the frequency of topological partitions that occur in a sample of Bayesian gene tree posterior distributions and provides an estimate for their genome-wide frequency and 95% confidence intervals. Because the number of possible topological relationships among taxa increases non-linearly as the number of species included in the analysis increases, large taxonomic datasets are intractable for thorough concordance factor analyses with BUCKy. We therefore used a reduced taxonomic dataset of 82 species that represent each of the major subclades of Acanthomorpha to reduce computational burden. For this reduced dataset, we generated alignments of the 82 species using MAFFT (**Supplementary Table 1**). We used MrBayes v. 3.2.7^43^ to infer a Bayesian gene tree distributions for each UCE locus. We ran MrBayes analyses with a GTR+Gamma molecular evolutionary model for 2 million generations, with trees sampled every 2,000 generations. Using the BUCKy program mbsum, we discarded the first 75% of sampled trees from the posterior sample of each UCE locus as burnin, summarized branching patterns in the remaining post-burnin posterior distribution, and used an alpha value of 1.0 for concordance factor estimation. These gene tree posterior distributions were then used for subsequent BUCKy estimation of concordance factors.

### f. Divergence-time estimation

We estimated divergence times for both the 702 and 1,084 species phylogenies using BEAST v.2.5^45^ following a protocol used in other UCE studies where BEAST analyses were run using reduced datasets of 30 UCE loci^29, 48, 67^. We constrained the phylogeny used in the BEAST runs using the IQ-TREE analyses of the 1,084 and 702 species datasets. For both the 1,084 and 702 species datasets, we performed BEAST analyses on three different sets of 30 loci, each randomly drawn without replacement from the entire set of ∼1,000 UCE loci. Exploratory BEAST analyses using sets of 25 and 50 loci yielded similar divergence time estimates, justifying the moderately computationally expensive analyses using the 30 locus datasets. For each subset of 30 loci, we determined the optimum partitioning scheme using the relaxed hierarchical clustering algorithm in PartitionFinder2 v. 2.1.1^49^. For divergence dating analyses performed in BEAST, we employed a relaxed lognormal clock model and a birth-death tree model. We assigned minima for age priors using forty-three fossil occurrences, described below. For each 30-locus dataset, we ran at least three replicate analyses for a minimum of 200 million generations beyond a total burnin of 200 million generations. As MCMC chains progressed, we accepted suggestions provided by BEAST for improving operators’ tuning parameter values. We used Tracer v.1.7.1 to observe effective sample size (ESS) values for posterior parameter estimates, to assess convergence among replicate MCMC chains, and to check for directional trends in parameter estimates^50^. Though the large size of the phylogenies created computational limits that resulted in moderate ESS values for some parameters, most parameter estimates reached ESS values well above 200. Some analyses were run up to 900 million generations in length to obtain adequate ESS values and ensure no directional trends in and unimodal distributions of parameter estimates. We used LogCombiner in BEAST v.1.8.4^51, 68^ to remove a burnin, and trees from each replicate run were randomly thinned. We thinned the sets of 1,084-species trees such that a total of 1,200 trees would remain across all runs, with an equal number of trees from each replicate contributing to the final summary trees. We randomly thinned the sets of 702-species trees such that 10,000 total trees remained across replicate runs. For each set of loci, we used LogCombiner to combine trees from replicate runs and TreeAnnotator in BEAST v.1.8.4 (Drummond et al. 2012) to estimate maximum clade credibility trees.

### g. Diversification rate analyses

We performed diversification rate analyses on the time-calibrated 1,084-taxa phylogeny that was run for the most generations and therefore had the highest ESS values for posterior parameter estimates. Species diversity of acanthomorph lineages used to determine sampling fractions for all diversification rate analyses were based on numbers of species presented in Eschmeyer’s Catalog of Fishes^3^, which at the time of the analyses included 19,219 species of Acanthomorpha. We removed all outgroups and duplicate samples of the same species before performing diversification rate analyses, leaving a phylogeny consisting of 1,075 acanthomorph species and a sampling fraction of 0.056. We performed stepping-stone simulations within TESS^52^ to estimate the marginal likelihoods of eight birth-death models and evaluate their fit to the acanthomorph time-calibrated phylogeny. Competing models included those with constant diversification rates, with episodically changing rates, or with decreasing speciation rates. Variants of the constant rate model with and without a mass extinction event were also tested to assess the potential impact of the K-Pg mass extinction on acanthomorph diversification. We tested variants of the competing models that accommodated the effects of incomplete taxon sampling by correcting for uniform (random) or diversified sampling methods.

Using the simulated marginal likelihoods of candidate models, we calculated Bayes factors and compared the relative fits of models to the acanthomorph phylogeny (Extended Data Table 1). With the exception of the models assuming a decreasing speciation rate, we ran stepping-stone simulations for 10,000 iterations with 1,000 power posteriors (stepping stones) and a burn-in period of 3,000 generations. Due to limited computational resources, we ran the simulations of marginal likelihoods for the decreasing speciation rate (i.e., continuous) models for 2,000 iterations (a burn-in of 500 generations was removed) with 100 power posteriors. This test of relative model fit greatly favored models assuming uniform (random) sampling methods over diversified methods (Extended Data Table 1), informing our model selection in downstream TESS analyses.

We extracted the three branching process-models that were most favored in the tests of relative model fit and assessed their absolute fit to the acanthomorph phylogeny using posterior-predictive simulations. We compared the predictive distributions of the gamma statistic, number of taxa, and lineage-through-time (LTT) plots to the empirical estimates.

We inferred the global speciation, extinction and net-diversification rates through time using the CoMET model as employed by TESS. For CoMET analyses represented in Extended Data Fig. 2, we specified a uniform (random) sampling strategy (ρ = 0.056), conditioned the model on taxa survival, discarded a burn-in of 30,000 iterations, and ran three replicate reversible-jump MCMC chains until the effective sample size (ESS) reached 200 or greater. We assessed convergence within each CoMET run by calculating the effective sample sizes and Geweke diagnostics for the diversification rate, and we assessed convergence of multiple runs using the Rubin-Gelman test.

We inferred global diversification rates and examined rate heterogeneity across clades using BAMM v.2.5.0^53^. Our analyses corrected for incomplete sampling by accounting for the sampling fractions of 304 major representative clades, most of which are taxonomic families, in accordance with the number of species described by Eschmeyer’s Catalog of Fishes ^3^. We also used the information on species diversity to appropriately adjust these clade-specific sampling fractions for non-monophyletic taxonomic groups. As in the TESS analyses, we set the global sampling fraction for all BAMM analyses to 0.056. We identified appropriate rate priors using the function setBAMMpriors in the R package BAMMtools v.2.1.6^69^, accounting for incomplete taxon sampling.

We ran Markov chain Monte Carlo simulations for 100 million generations under the default settings. We also used default settings for MCMC scaling operators and move frequencies. We ran BAMM 23 times, each time under different conditions that might be appropriate for a medium-sized tree consisting of ∼1,000 taxa. Runs tested different combinations of numbers of MCMC chains (2 or 4), temperature increment parameters (deltaT = 0.05 or 0.1) and expected number of diversification rate shifts (1, 5, 10, 15, 20, 25, 30, 35, 40 and 45). We tested the effect of changing the number of expected shifts—despite the setBAMMpriors function in BAMMtools recommending an input of just one expected shift— because preliminary runs suggested that the number of diversification rate shifts is greater than 20. For each analysis, we compared Bayes factors for competing prior models with different numbers of expected rate shifts.

### h. Body shape trait data

We pruned a body trait dataset to the 680 species that matched those in the 702-species UCE phylogeny^24^. The body shape dataset included maximum body depth, maximum fish width, head depth, lower-jaw length, mouth width, minimum caudal-peduncle depth, and minimum caudal-peduncle width. We measured maximum body depth as the maximum linear distance from the dorsal to ventral margin of the body, anywhere in the region between the posterior edge of the operculum to the anterior portion of the caudal peduncle. We measured maximum fish width as the maximum width anywhere along the body. We measured head depth as the vertical distance from the dorsal to ventral margins of the head passing through the center of the eye. We measured lower jaw length as the linear distance from the anterior end of the lower jaw to the articular-quadrate joint. We measured mouth width as the linear distance between the left and right articular-quadrate joints. We measured minimum caudal peduncle depth as the minimum vertical distance from the dorsal to ventral margin of the caudal peduncle. We measured minimum caudal peduncle width as the narrowest point along the caudal peduncle. Lastly, we measured fish standard length as the linear distance from the anterior tip of the upper jaw to the posterior edge of the hypural plate or to the posterior end of the vertebral column (i.e., in species that lack a hypural plate). We accounted for size by regressing log-transformed lengths against log-transformed standard fish length and calculating phylogenetic residuals using the phyl.resid function implemented in PHYTOOLS^55^. We used this matrix of residuals for seven traits for phylogenetic comparative analyses using a multivariate framework.

### i. Disparity through time analyses

We assessed disparity through time (DTT) using the dtt function implemented in GEIGER^56^. We incorporated uncertainty in divergence times by repeating analyses across 100 trees randomly sampled from the posterior distribution. Since we were specifically interested in patterns around and following the K-Pg boundary, we summarized disparity through time in 1 My intervals as the proportion of trees in which the observed subclade disparity fell below that expected from a Brownian motion process (*i.e.*, the expected pattern if the clade is undergoing adaptive radiation) (results in Supplementary Fig. 28). Focusing on the time following the K-Pg boundary, we statistically summarized the 1 My time interval during which the observed subclade disparity first dropped below that expected from a Brownian motion process (results in Supplementary Fig. 28). We tested the sensitivity of disparity through time analyses to body shape traits and to the inclusion of major clades by repeating all analyses to the exclusion of clades that arose immediately around the K-Pg boundary and to the exclusion of each trait, as well as repeating the analyses for each trait separately in a univariate framework (results in Extended Data Fig. 6). We also visualized the evolutionary histories of the following focal lineages using the phenogram function implemented in PHYTOOLS^55^: Carangiformes, Lophioidei, Perciformes, Scombriformes, “Squamipinnes” (here defined as the clade containing *Chaetodon kleinii* and *Luvarus imperialis*), Syngnathiformes, and Tetraodontoidei. We visualized body shape morphospace using principal component analysis (PCA) with the prcomp function. Since the traits had different variances, we scaled all traits prior to performing PCA.

## 2. Fossil calibrations for relaxed clock BEAST analyses

**i. *Node*: Stem lineage Lampriformes**, dating the most recent common ancestor (MRCA) of *Lampris guttatus* and *Stylephorus chordatus*, which is the MRCA of Lampriformes and all other paracanthopterygians. *First occurrence*: †*Aipichthys minor*. Sannine Limestone, Hadjula, Lebanon. *Resolution in phylogenetic analyses*: †*Aipichthys minor* is resolved as a lampriform stem lineage in parsimony analyses of morphological characters^70, 71^. *Stratigraphy*: The Hadjula fish beds are located below reported occurrence of *Mantelliceras mantelli*, which defines the first complete ammonite zone of the Late Cretaceous. The top of the *Mantelliceras mantelli* Zone is dated at 98.0 Ma^72^, from which we derive a minimum age for *Aipichthys*. *Minimum age*: 98.0 Ma. *Prior setting*: lognormal prior, minimum age 98.0 Ma, mean=1.0, S.D.=1.25, upper 95% CI: 119.0 Ma^48^.

**ii. *Node*: Stem lineage Polymixiiformes**, dating the MRCA of *Polymixia lowei* and *Aphredoderus sayanus*, which subtends the MRCA of *Polymixia* and Percopsiformes. *First occurrence*: †*Homonotichthys dorsalis*. Lower Chalk of Sussex and Kent, UK^73^. *Resolution in phylogenetic analyses*: none. *Character states*: four full-sized branchiostegals; anterior branchiostegals reduced and forming support for chin barbel^16, 73^. *Stratigraphy*: middle-upper Cenomanian, zone of *Holoaster subglobosus*, corresponding to the late Cenomanian^73–75^. The minimal age for †*Homonotichthys dorsalis* is provided by the age of Cenomanian-Turonian boundary, which is dated as 93.9 Ma^72^. *Absolute age estimate*: 93.6 Ma (28). *Prior setting*: lognormal prior, minimum age 93.6 Ma, mean = 1.06, S.D.=1.25, upper 95% CI: 116.35 Ma^4^.

**iii. *Node*: Stem lineage Aphredoderidae**, dating the MRCA of *Aphredoderus sayanus* and *Typhlichthys subterraneus*, which subtends the MRCA of Aphredoderidae and Amblyopsidae. *First occurrence*: †*Trichophanes foliarum*. Florissant Formation, Colorado, USA^76^. *Resolution in phylogenetic analyses*: †*Trichophanes* and Aphredoderus form a clade in a maximum parsimony analysis of 47 morphological characters^77^. *Character states*: ventral margins of lachrymal and infraorbitals spiny; alveolar process of premaxilla divided into separate segments^76–78^. *Stratigraphy*: The upper Priabonian Florissant Roadcut is dated radiometrically using ^40^Ar/^39^Ar isotope ratios as 34.07 Ma^79^. *Absolute age estimate*: 34.1 Ma. *Prior setting*: lognormal prior, minimum age 34.1 Ma, mean=1.0, S.D.= 1.347, upper 95% CI: 59.0 Ma^20^.

**iv. *Node*: Stem lineage *Zenopsis***, dating the MRCA of *Zenopsis* and *Zeus*. *First occurrence*: †*Zenopsis clarus*, †*Z. tyleri*, and †*Z. hoernesi*. Lower Maikopian series, Psheka Horizon of the Belaya River, Caucasus^80^, and Lower Dysodylic shales, Strujinoasa-Drăguşina and Piatra Neamţ, Romania^81^; Lower Dysodylic shales, Piatra Neamţ, Romania^81^; Laško (Tüffer), Slovenia^81^. *Resolution in phylogenetic analyses*: maximum parsimony analysis of 45 morphological characters resulted in a phylogeny where †*Zenopsis clarus*, †*Z. tyleri*, and †*Z. hoernesi,* and *Z*. *oblongus* are resolved in a clade^82^. *Character states*: slender lachrymal; pelvic-fin spines absent; buckler-like plates present along ventral midline of abdomen, and along dorsal ridge from middle of spinous to end of soft dorsal fin^82^. *Stratigraphy*: lower Rupelian [P18], lower Khadumian regional stage^83^. *Absolute age estimate*: 32.0 Ma^84^. *Prior setting*: lognormal prior, minimum age 32.0 Ma, mean=1.0, S.D.= 0.33, upper 95% CI: 36.7 Ma^20^.

**v. *Node*: Stem lineage Lampridae**, dating the MRCA of *Lampris guttatus* and *Zu elongatus*, which is the MRCA of Lampriformes. *First occurrence*: †*Turkmene finitimus*. Danatinsk Suite, Uylya-Kushlyuk locality, Turkmenistan^85, 86^. *Resolution in phylogenetic analysis*: none. *Character states*: first dorsal fin pterygiophore strongly reclined posteriorly; enlarged pectoral fins inserting high on flank; shoulder girdle broad ventrally, with expanded coracoid; long parapophyses absent from abdominal vertebrate^86^. *Stratigraphy*: uppermost Thanetian-lowermost Ypresian^87^. *Absolute age estimate*: 55.8 Ma^84^. *Prior setting*: lognormal prior, minimum age 55.8 Ma, mean=1.0, S.D.= 1.411, upper 95% CI: 83.5 Ma^20^.

**vi. *Node:* Stem lineage Holocentridae,** dating the MRCA of *Myripristis violacea* and *Rondeletia loricata*, which subtends the MRCA of Holocentridae and Berycoidei. *First occurrence*: †*Stichocentrus liratus* from the Sannine Limestone, Hadjula, Lebanon^88^. *Resolution in phylogenetic analyses*: none. *Character states*: The penultimate anal-fin spine of †*Stichocentrus* is enlarged and represents a synapomorphy of holocentroids^16, 89^. *Stratigraphy*: See discussion for Calibration i, †*Aipichthys minor*. *Absolute age estimate*: 98.0 Ma. *Prior setting*: lognormal prior, minimum age 98.0 Ma, mean=1.0, S.D.= 0.8, upper 95% CI: 108.0 Ma^48^.

**vii. *Node*: Stem lineage Myripristinae**, dating the MRCA of *Myripristis violacea* and *Sargocentron coruscum*, which subtends the MRCA of Holocentridae. *First occurrence*: †*Eoholocentrum macrocephalum*, †*Berybolcensis leptacanthus*, and †*Tenuicentrum pattersoni*. Pesciara beds of ‘Calcari nummulitici’, Bolca, Italy^90–92^. *Resolution in phylogenetic analyses*: analysis of 72 morphological characters resolve †*Eoholocentrum*, †*Berybolcensis*, and †*Tenuicentrum* as stem-lineage Myripristinae^93^. *Character states*: tooth-bearing platform expanded and overhangs lateral side of dentary near symphysis; premaxillary tooth field curves dorsally toward ascending process at symphysis; edentulous ectopterygoid (†*Berybolcensis* and †*Tenuicentrum*); spinous procurrent caudal-fin rays reduced to four in the upper and three in the lower lobe (†*Berybolcensis* and †*Tenuicentrum*)^93^. *Stratigraphy*: upper Ypresian [NP14]^94, 95^. *Absolute age estimate*: 50 Ma^84^. *Prior setting*: lognormal prior, minimum age 50.0 Ma, mean=1.0, S.D.= 0.6, upper 95% CI: 57.3 Ma^20^.

**viii. *Node*: Crown lineage Syngnathiformes**, dating the MRCA of *Pegasus volitans* and *Syngnathus scovelli*, which subtends the MRCA of Syngnathiformes. *First occurrence*: †*Gasterorhamphosus zuppichinii*. “Calcari di Melissano”, Porto Selvaggio, Lecce province, Italy^96^. *Resolution in phylogenetic analyses*: none, but Orr^97^ argues that †*Gasterorhamphosus* is a stem lineage of a clade containing Macrorhamphosidae and Centriscidae. *Character states*: anal-fin spine absent; enlarged dorsal-fin spine with serrated posterior margin; elongated tubular snout; pleural ribs absent; cleithrum bears enlarged posterodorsal process; rod-like anteroventral process of coracoid; pectoral rays simple^97, 98^. *Stratigraphy*: uppermost Campanian-lowermost Maastrichtia^99^. *Absolute age estimate*: 69.71 Ma ^48^. *Prior setting*: lognormal prior, minimum age 69.71 Ma, mean=1.0, S.D.= 0.785, upper 95% CI: 79.6 Ma^48^.

**ix. *Node*: Stem lineage Centriscidae**, dating the MRCA of *Aeoliscus strigatus* and *Macroramphosus gracilis*, which subtends the MRCA of Centriscidae. *First occurrence*: †*Paramphisile weileri* and †*Paraeoliscus robinetae*. Pesciara beds of ‘Calcari nummulitici’, Bolca, Italy^100^. *Resolution in phylogenetic analyses*: none. *Character states*: caudal fin directed posteroventrally (†*Paraeoliscus*); dorsal spine jointed distally^97^. *Stratigraphy*: See discussion for Calibration viii †*Eoholocentrum macrocephalum*. *Absolute age estimate*: 50 Ma^84^. *Prior setting*: lognormal prior, minimum age 50.0 Ma, mean=1.0, S.D.= 0.6, upper 95% CI: 57.3 Ma^20^.

**x. *Node*: Crown lineage *Ariomma***, dating the MRCA of *Ariomma indica* and *A. melana*. *First occurrence*: †*Ariomma geslini* from the diatomites of St Eugène, Chelif Basin, Algeria^29^. *Resolution in phylogenetic analyses*: none. *Character states*: extension of pleural ribs onto caudal vertebrae^101, 102^, and a distinctive pattern of anal-fin skeleton shared with *A. bondi* and *A. melana*, consisting of anterior pterygiophores that bend posteriorly, overlapping the similarly curved shafts of successive pterygiophores^102, 103^. *Stratigraphy*: Details presented in Friedman et al.^29^. *Absolute age estimate*: 5.94 Ma. *Prior setting*: lognormal prior, minimum age 5.94 Ma, mean=1.0, S.D.= 1.299, upper 95% CI: 28.96 Ma^29^.

**xi. *Node*: stem lineage Stromateidae**, dating the MRCA of *Pampus argenteus* and *Ariomma indica*, which subtends the MRCA Stromateidae and the clade that comprises *Ariomma* and Nomeidae. *First occurrence*: †*Pinichthys pulcher* from the Pshekha Horizon †*Planorbella* Beds, Lower Maikop Caucasus^104^. *Resolution in phylogenetic analyses*: none. *Character states*: †*Pinichthys* has a stellate, bony base of papillae in the region expected to bear pharyngeal sacs^105^. Synapomorphies of stromateiod displayed by †*Pinichthys* include: a continuous dorsal fin; a ventral articulation between the coracoid and ventral tip of the cleithrum; and an elongated first anal fin pterygiophore that is strongly reclined^106^. †*Pinichthys* differs from extant lineages of Stromateidae by having well-developed pelvic fins in adults and lacking extension of the first anal fin pterygiophore posterior to the first haemal spine^29^. †*Pinichthys* is interpreted as a stem-lineages stromateiod, which is consistent with previous interpretations^104^ *Stratigraphy*: Details presented in Friedman et al.^29^. *Absolute age estimate*: 32.02 Ma. *Prior setting*: lognormal prior, minimum age 32.02 Ma, mean=1.0, S.D.= 1.2, upper 95% CI: 51.6 Ma^29^.

**xii. *Node*: stem lineage Chiasmodontidae**, dating the MRCA of *Kali kerberti* and *Tetragonurus atlanticus*, which subtends the MRCA of Chiasmodontidae and *Tetragonurus*. *First occurrence*: †*Bannikovichthys paelignus* from the fish-bearing laminates of Toricella Paligna, Italy^107^. *Resolution in phylogenetic analyses*: none. *Character states*: The hypothesis that †*Bannikovichthys* is related to living Chiasmodontidae is based on an absence of scales, elongated opercles, delicate ossification of the interopercle and subopercle, and enlarged hourglass-shaped radials^107^. The relationships of †*Bannikovichthys* among living chiasmodontids is uncertain, so it is conservatively treated as a stem-lineage taxon^29^. *Stratigraphy*: Details presented in Friedman et al.^29^. *Absolute age estimate*: 11.9 Ma. *Prior setting*: lognormal prior, minimum age 11.9 Ma, mean=1.0, S.D.= 1.454, upper 95% CI: 41.56 Ma^29^.

**xiii. *Node*: within crown of *Scomber***, dating the MRCA of *Scomber australasicus* and *Scomber japonicus*. *First occurrence*: *Scomber colias* from the Raz-el-Aïn, Oran, Algeria. *Resolution in phylogenetic analyses*: This fossil is attributed to the living species *Scomber colias*, which is resolved as the sister lineage of S. japonicus in a molecular phylogenetic analysis^108^. *Stratigraphy*: Details presented in Friedman et al.^29^. *Absolute age estimate*: 5.94 Ma. *Prior setting*: lognormal prior, minimum age 5.94 Ma, mean=1.0, S.D.= 1.255, upper 95% CI: 27.4 Ma^29^.

**xiv. *Node*: stem lineage *Scomber***, dating the MRCA of *Scomber australasicus* and *Rastrelliger kanagurta*. *First occurrence*: †*Scomber saadii* from the Padbeh Formation of Istehbanât, Iran^109^. *Resolution in phylogenetic analyses*: None. Character states: †*Scomber saadii* exhibits serrations on the teeth that is a synapomorphy of *Scomber*^108, 110^. *Stratigraphy*: Details presented in Friedman et al.^29^. *Absolute age estimate*: 33.9 Ma. *Prior setting*: lognormal prior, minimum age 33.9 Ma, mean=1.0, S.D.= 1.02, upper 95% CI: 48.45 Ma^29^.

**xv. *Node*: stem lineage *Acanthocybium***, dating the MRCA of *Acanthocybium solandri* and *Scomberomorus maculatus*. *First occurrence*: †*Palaeocybium proosti* from the London Clay Formation, Isle of Sheppey, UK^111^. *Resolution in phylogenetic analysis*: none. *Character states*: †*Palaeocybium* and *Acanthocybium* share closely spaced teeth with blunt tips and the holotype specimen shows an elongate lower jaw that is similar to *Acanthocybium* and *Scomberomorus*^112–, 114^. Stratigraphy: Details presented in Friedman et al.^29^. *Absolute age estimate*: 50.5 Ma. *Prior setting*: lognormal prior, minimum age 50.5 Ma, mean=1.0, S.D.= 0.5, upper 95% CI: 56.72 Ma^29^.

**xvi. *Node*: stem lineage Sardini + Thunnini**, dating the MRCA of *Scomberomorus maculatus* and *Euthynnus affinis*. *First occurrence*: †*Eocoelopoma portentosum* from the Danatinsk Formation, Turkmenistan^115^. *Resolution in phylogenetic analyses*: None. *Character states*: Details presented in Friedman et al.^29^. Stratigraphy: Details presented in Friedman et al.^29^. Absolute age estimate: 54.17 Ma. *Prior setting*: lognormal prior, minimum age 54.17 Ma, mean=1.0, S.D.= 0.698, upper 95% CI: 62.71 Ma^29^.

**xvii. *Node*: stem lineage *Thunnus***, dating the MRCA of *Thunnus orientalis* and *Auxis rochei*, which is the MRCA of Thunnini. *First occurrence*: †*Thunnus abchasicus* from the Kuma Horizon, Pshekha River, Caucasus^116^. *Resolution in phylogenetic* analyses: None. *Character states*: †*Thunnus abchasicus* exhibits several synapomorphies of *Thunnus* that include the presence of small oral teeth and the shortened preural vertebrae^116^. The presence of a spine in the second dorsal fin suggests that †*T. abchasicus* is a stem lineage *Thunnus*^115^. Stratigraphy: Details presented in Friedman et al.^29^. *Absolute age estimate*: 38.62 Ma. *Prior setting*: lognormal prior, minimum age 38.62 Ma, mean=1.0, S.D.= 0.91, upper 95% CI: 50.78 Ma^29^.

**xviii. *Node*: stem lineage *Promethichthys***, dating the MRCA of *Promethichthys prometheus* and *Diplospinus multistriatus*. *First occurrence*: †*Hemithyrsites maicopicus* from the Upper Maikop of the Kurdzhips River, Adygea, Caucasus^115^. *Resolution in phylogenetic analyses*: None. *Character states*: Derived traits shared by *Promethichthys* and †*Hemithyrsites maicopicus* include reduction of the pelvic fin to a single spine and the presence of two pairs of dorsal and anal finlets^115^. *Stratigraphy*: Details presented in Friedman et al.^29^. *Absolute age estimate*: 15.97 Ma. *Prior setting*: lognormal prior, minimum age 15.97 Ma, mean=1.0, S.D.= 0.986, upper 95% CI: 29.67 Ma^29^.

**xix. *Node*: Stem lineage Channidae**, dating the MRCA of *Channa micropeltes* and *Helostoma temminckii,* which subtends the MRCA of Channidae and a clade containing Helostoma, Anabantidae, and Osphronemidae. *First occurrence*: †*Anchichanna kuldanensis* from the Ganda Kas Area, Kuldana Formation, Kala Chitta Hills, Pakistan^117^. *Resolution in phylogenetic analyses*: None. *Character states*: †*Anchichanna kuldanensis* exhibits the Channidae synapomorphy of the outer wall of the auditory bulla formed mainly by the prootic, but is likely a stem-lineage taxon because it does not share synapomorphies with either *Channa* or *Parachanna*^117^. *Stratigraphy*: The Kuldana Formation and other vertebrate fossil localities of the Kala Chitta Hills straddle the Ypresian-Lutetian boundary^118^. *Absolute age estimate*: 41.3 Ma. *Prior setting*: lognormal prior, minimum age 41.3 Ma, mean=1.0, S.D.= 1.31, upper 95% CI: 64.7 Ma^119^.

**xx. *Node*: Stem lineage Sphyraenidae,** dating the MRCA of *Sphyraena jello* and *Lactarius lactarius*. *First occurrence*: †*Sphyraena bolcensis,* Pesciara beds of ‘Calcari nummulitici’, Bolca, Italy^120, 121^. *Resolution in phylogenetic analyses*: None. *Character states*: three ‘T’-shaped, sutured predorsals or spineless pterygiophores^122^; elongate gape; upper jaw non-protrusible; enlarged fangs on dentary, premaxilla, and palatine. *Stratigraphy*: upper Ypresian [NP14]^94, 95^*. Absolute age estimate*: 50.0 Ma^95^. *Prior setting*: lognormal prior, minimum age 50.0 Ma, mean=1.0, S.D.= 0.5999, upper 95% CI: 57.3 Ma^1^.

**xxi. *Node*: Stem lineage *Mene***, dating the MRCA of *Mene maculata* and *Xiphias gladius*. *First occurrence*: †*Mene purdyi*, ‘Mancora Formation’, Perú^123^; †*M. triangulum*, Danatinsk Suite, Uylya-Kushlyuk locality, Turkmenistan^85, 87^; and †*Mene* sp., ‘Stolle Klint clay’, Ølst Formation, Denmark^124^. *Resolution in phylogenetic analyses*: None. *Character states*: frontal vault; many small infraorbitals; all anal fin rays short, with anteriormost rays plate like; anal fin spines absent; postcleithrum broad and contacts first anal fin pterygiophore; disc-like body with deep ventral keel; greatly elongated second anal fin ray^123^. *Stratigraphy*: uppermost Thanetian-lowermost Ypresian [NP14]^87, 124^*. Absolute age estimate*: 55.2 Ma^125^. *Prior setting*: lognormal prior, minimum age 55.2 Ma, mean=1.0, S.D.= 0.92, upper 95% CI: 67.5 Ma^1, 48^.

**xxii. *Node*: Stem lineage Carangidae**, dating the MRCA of *Caranx caninus* and *Trachinotus goodei*, which subtends the MRCA of the clade comprising Carangidae, *Coryphaena*, *Rachycentron*, Scomberoidinae, and Trachinotinae. *First occurrence*: †*Archaeus oblongus*. Danatinsk Suite, Uylya-Kushlyuk locality, Turkmenistan^85^. *Resolution in phylogenetic analyses*: None. *Character states*: broad gap between second and third anal-fin spines^126^. *Stratigraphy*: uppermost Thanetian-lowermost Ypresian^87^. *Absolute age estimate*: 55.8 Ma^84^. *Prior setting*: lognormal prior, minimum age 55.8 Ma, mean=1.0, S.D.= 0.661, upper 95% CI: 63.9 Ma^1^.

**xxiii. *Node*: Stem lineage Caranginae,** dating the MRCA of *Caranx caninus* and *Seriola dumerili*, which subtends the MRCA of Carangidae. *First occurrence*: †*Eastmanalepes primaevus,* Pesciara beds of ‘Calcari nummulitici’, Bolca, Italy^127^. *Resolution in phylogenetic analyses*: None. *Character states*: lateral line scales modified as thick scutes^126, 128^. *Stratigraphy*: upper Ypresian [NP14]^94^*. Absolute age estimate*: 49.0 Ma^48^. *Prior setting*: lognormal prior, minimum age 49.0 Ma, mean=1.0, S.D.= 0.1, upper 95% CI: 52.2 Ma^1, 48^.

**xxiv. *Node*: Stem lineage Echeneioidei**, dating the MRCA of *Echeneis naucrates* and *Scomberoides lysan*, which subtends the MRCA of the clade comprising Echeneidae, *Coryphaena*, *Rachycentron*, Scomberoidinae, and Trachinotinae. *First occurrence*: †*Ductor vestenae* from the Pesciara locality of Bolca, Italy^120, 121^. *Resolution in phylogenetic analyses*: parsimony and Bayesian analyses of morphology and combined molecular and morphological datasets resolve †*Ductor vestenae* as either a stem echeneioid or as the sister the clade containing *Rachycentron* and *Coryphaena*^129^. We follow the conservative consideration of †*Ductor vestenae* as a stem-lineage taxon, as used in Harrington et al.^48^. *Character states*: Discussed in Friedman et al.^129^. *Stratigraphy*: upper Ypresian [between NP14 and SPZ11]^94^*. Absolute age estimate*: 49.0 Ma^48^. *Prior setting*: lognormal prior, minimum age 49.0 Ma, mean=1.0, S.D.= 0.1, upper 95% CI: 52.2 Ma^48^.

**xxv. *Node*: Stem lineage Echeneidae**, dating the MRCA of *Echeneis naucrates* and *Coryphaena hippurus,* which subtends the MRCA of clade containing Echeneidae, *Coryphaena*, *and Rachycentron*. *First occurrence*: unnamed Echeneidae undet. from the Frauenweiler clay pit, Germany^130^. *Resolution in phylogenetic analyses*: None. *Character states*: no supraneurals; multiple anal fin pterygiophores insert anterior to first haemal spine; spinous dorsal fin modified as adhesion disc^129, 131^. *Stratigraphy*: See discussion in Harrington et al.^48^. *Absolute age estimate*: 29.62 Ma^48^. *Prior setting*: lognormal prior, minimum age 29.62 Ma, mean=1.0, S.D.= 0.87, upper 95% CI: 41.0 Ma^48^.

**xxvi. *Node*: Stem lineage *Scomberoides***, dating the MRCA of *Scomberoides lysan* and *Trachinotus goodei*, which subtends the MRCA of Scomberoidinae and Trachinotinae. *First occurrence*: *Scomberoides spinosus* from the Upper Maikop at Chernaya Rechka, Caucasus^87^. *Resolution in phylogenetic analyses*: None. *Character states*: *Scomberoides spinosus* exhibits two synapomorphies of Scomberoidinae: 26 versus 24 vertebrae and posterior rays of the dorsal and anal fin developed as finlets ^126, 128, 132^. *Stratigraphy*: See discussion in Harrington et al.^48^. *Absolute age estimate*: 19.3 Ma^48^. *Prior setting*: lognormal prior, minimum age 19.3 Ma, mean=1.0, S.D.= 1.095, upper 95% CI: 35.8 Ma^48^.

**xxvii. *Node*: Stem lineage Citharidae**, dating the MRCA of *Gymnachirus melas* and *Scophthalmus* rhombus, which subtends the MRCA of Citharidae and all other pleuronectoids except *Psettodes*. *First occurrence*: †*Eobothus minimus* from the Pesciara beds of ‘Calcari nummulitici’, Bolca, Italy^133, 134^. *Resolution in phylogenetic analyses*: †*Eobothus* is the sister lineage of *Citharus*^133^. *Character states*: complete orbital asymmetry; dorsal fin extends above orbit; hook-shaped urohyal; parahypural not in articulation with pural centrum 1; long neural spine on preural centrum 2^133^. *Stratigraphy*: See discussion in Harrington et al. ^48^. *Absolute age estimate*: 49.0 Ma^48^. *Prior setting*: lognormal prior, minimum age 49.0 Ma, mean=1.0, S.D.= 0.68, upper 95% CI: 57.3 Ma^48^.

**xxviii. *Node*: Stem lineage of the clade containing Soleidae and Cynoglossidae**, dating the MRCA of *Aseraggodes xenicus* and *Poecilopsetta plinthus*. *First occurrence*: †*Eobuglossus eocenicus* from the Mokkatam Formation, Djebel Turah, Egypt^135^. *Resolution in phylogenetic analyses*: none. *Character states*: †*Eobuglossus* shows derived cranial features limited to Soleidae, including blind side preopercular canal terminating on ventral margin of preopercular and convex portion of blind side dentary anterior to angulo-articular^135, 136^. *Stratigraphy*: See discussion in Harrington et al.^48^. *Absolute age estimate*: 41.2 Ma^48^. *Prior setting*: lognormal prior, minimum age 41.9 Ma, mean=1.0, S.D.= 0.4, upper 95% CI: 46.4 Ma^48^.

**xxix. *Node:* Stem lineage Bothidae**, dating the MRCA of *Bothus pantherinus* and *Cyclopsetta frimbriata*, which subtends the MRCA of the clade containing Bothidae and Cyclopsettidae. *First occurrence*: †*Oligobothus pristinus* from the Lower Dysodilic shales, Piatra Neamţ, Romania^137^. *Resolution in phylogenetic analyses*: None. *Character states*: myorhabdoi present^137^. *Stratigraphy*: See discussion in Harrington et al.^48^. *Absolute age estimate*: 29.62 Ma^48^. *Prior setting*: lognormal prior, minimum age 29.62 Ma, mean=1.0, S.D.= 0.06, upper 95% CI: 32.6 Ma^48^.

**xxx. *Node:* Stem lineage Pleuronectidae**, dating the MRCA of *Pleuronichthys cornutus* and *Paralichthys albigutta*, which subtends the MRCA of the clade containing Pleuronectidae and Paralichthyidae. *First occurrence*: †*Oligopleuronectes germanicus* from the Frauenweiler fossil site, Germany^138^. *Resolution in phylogenetic analyses*: None. *Character states*: Suggested as a member of Pleuronectidae based on †*Oligopleuronectes* being right-eyed and having a lateral process on the eye-side frontal^138, 139^. *Stratigraphy*: See discussion in Harrington et al.^48^. *Absolute age estimate*: 29.62 Ma^48^. *Prior setting*: lognormal prior, minimum age 29.62 Ma, mean=1.0, S.D.= 0.45, upper 95% CI: 35.3 Ma^48^.

**xxxi. *Node:* Stem lineage *Bothus***, dating the MRCA of *Bothus pantherinus* and *Crossorhombus azureus*. *First occurrence*: *Bothus sp.* from the Middle Tsurevsky Member of the Tsurevsky Formation along the bank of the Psheka River, Caucasus^140^. *Resolution in phylogenetic analyses*: None. *Character states*: A specific affinity to *Bothus* is supported by the presence of robust rectangular haemal spines^140, 141^. *Stratigraphy*: See discussion in Harrington et al.^48^. *Absolute age estimate*: 11.056 Ma^48^. *Prior setting*: lognormal prior, minimum age 29.62 Ma, mean=1.0, S.D.= 0.84, upper 95% CI: 39.9 Ma^48^.

**xxxii. *Node:* Stem lineage Pseudocrenilabrinae,** dating the MRCA of *Bathybates minor* and *Amphilophus citrinellus*, which subtends the MRCA of the clade that contains Pseudocrenilabrinae and Cichlinae. *First occurrence*: †*Mahengechromis plethos* from Mahenge, Tanzania^142^. *Resolution in phylogenetic analyses*: Several phylogenetic resolutions of †*Mahengechromis* place the taxon as either a crown lineage Cichlidae or crown lineage Pseudocrenilabrinae^142, 143^. We follow Rabosky et al.^119^ in conservatively treating †*Mahengechromis* as a stem-lineage Pseudocrenilabrinae. *Character states*: The identification of †*Mahengechromis* as a cichlid is supported by the morphology of the lower pharyngeal jaw, squamation, and meristics of vertebrae and median fins^142–144^. *Stratigraphy*: The Mahenge fossil deposits accumulated in a crater formed by intrusion of kimberlite and U-Pb of a zircon crystal dates the formation of the kimberlite at 45.83 +/-0.17 Ma^145^. Because the lacustrine sediments predate the kimberlite emplacement it was suggested the age of the fossil deposits are 0.2 to 0.1 Ma younger^145, 146^. We use the conservative date of 45 Ma used in Rabosky et al.^119^. Absolute age estimate: 45.0 Ma. *Prior setting*: lognormal prior, minimum age 45.0 Ma, mean=1.0, S.D.= 1.29, upper 95% CI: 67.3 Ma.

**xxxiii. *Node*: Crown lineage Exocetoidei**, dating the MRCA of *Xenentodon cancila* and *Hemiramphus far*, which subtends the MRCA of Belonidae and Hemiramphidae. *First occurrence*: †*Rhamphexocoetus volans* from Pesciara beds of ‘Calcari nummulitici’, Bolca, Italy^147^. *Resolution in phylogenetic analyses*: None. *Character states*: lower jaw symphysis extended; ventral lobe of caudal fin enlarged relative to dorsal lobe; pectoral fins greatly expanded^148–150^. *Stratigraphy*: upper Ypresian [NP14]^94, 95^. *Absolute age estimate*: 49.0 Ma^4^. *Prior setting*: lognormal prior, minimum age 49.0 Ma, mean=1.0, S.D.= 1.49, upper 95% CI: 80.52 Ma^4^.

**xxxiv. *Node*: Stem lineage *Calotomus* (Labridae)**, dating the MRCA of *Calotomus spinidens* and *Sparisoma radians*. *First occurrence*: †*Calotomus preisli* from the Leitha Limestone, St. Margarethen, Austria^151^. Resolution in phylogenetic analyses: None. *Character states*: Presence of upper pharyngeal bones bearing 1 to 3 rows of teeth and presence of a lateral canine on the premaxilla are synapomorphies for Scarinae. A conical tooth on the medial face of the premaxilla is a synapomorphy of *Calotomus*^151^. *Stratigraphy*: The Leitha Limestone is constrained by calcareous nannofossils to the standard biozone NN6^151^, which is entirely in the Serravallian stage of the Miocene. The top of NN6 is dated at 11.9 Ma^4^. Absolute age estimate: 11.9 Ma^4^. *Prior setting*: lognormal prior, minimum age 11.9 Ma, mean=1.0, S.D.= 1.501, upper 95% CI: 43.95 Ma^4^.

**xxxv. *Node*: Stem lineage Luvaridae**, dating the MRCA of *Luvarus imperialis* and *Acanthurus bahianus*, which subtends the MRCA of *Luvarus* and the clade containing *Zanclus cornutus* and Acanthuridae. *First occurrence*: †*Avitoluvarus dianae*, †*A. mariannae*, †*Kushlukia permira*, and †*Luvarus necopinatus*. Danatinsk Suite, Uylya-Kushlyuk locality, Turkmenistan^152^. *Resolution in phylogenetic analyses*: a parsimony analysis of morphological characters resolves a clade that contains †*Avitoluvarus,* †*Kushlukia*, †*Luvarus necopinatus*, and *L*. *imperialis*, which is sister to a clade comprising *Zanclus* and Acanthuridae^152^. *Character states*: median pterygial truss surrounding most of body; two or fewer dorsal-fin spines; no anal-fin spines; distal end of first anal-fin pterygiophore greatly elongated anteriorly; hypurals 1-4 fused; caudal fins broadly overlap hypural(s); pelvic fin rudimentary in adults; teeth absent or greatly reduced^152^. *Stratigraphy*: uppermost Thanetian-lowermost Ypresian^87^. *Absolute age estimate*: 55.8 Ma^84^. *Prior setting*: lognormal prior, minimum age 55.8 Ma, mean=1.0, S.D.= 0.661, upper 95% CI: 63.9 Ma^20^.

**xxxvi. *Node*: Stem lineage Acanthurinae**, dating the MRCA of *Acanthurus bahianus* and *Naso unicornis*, which subtends the MRCA of Acanthuridae. *First occurrence*: †*Proacanthurus tenius* from the ‘Pesciara beds of ‘Calcari nummulitici’, Bolca, Italy^153^. *Resolution in phylogenetic analyses*: None. *Character states*: caudal peduncle bears folding spine^153^. *Stratigraphy*: upper Ypresian [NP14]^94^*. Absolute age estimate*: 50 Ma^94^. *Prior setting*: lognormal prior, minimum age 50.0 Ma, mean=1.0, S.D.= 0.6, upper 95% CI: 57.3 Ma^1^.

**xxxvii. *Node*: Stem lineage Siganidae**, dating the MRCA of *Siganus spinus* and *Scatophagus argus*. *First occurrence*: †*Siganopygaeus rarus*. Danatinsk Suite, Uylya-Kushlyuk locality, Turkmenistan^154^. *Resolution in phylogenetic analyses*: parsimony analysis of morphological characters resolves four Eocene and Oligocene taxa, including †*Siganopygaeus*, as stem-lineage Siganidae^154^. *Character states*: two pelvic-fin spines; seven or more anal-fin spines; ten or fewer anal-fin rays^154^. *Stratigraphy*: uppermost Thanetian-lowermost Ypresian^87^. *Absolute age estimate*: 55.8 Ma^84^. *Prior setting*: lognormal prior, minimum age 55.8 Ma, mean=1.0, S.D.= 0.661, upper 95% CI: 63.9 Ma^20^.

**xxxviii. *Node*: Stem lineage of Chaetodontidae**, dating the MRCA of *Chaetodon kleinii* and *Leiognathus equula*, which subtends the MRCA of the clade that contains Chaetodontidae and Leiognathidae. *First occurrence*: Chaetodontidae cf. *Chaetodon* (tholichthys-stage larvae) from the ‘Fish shales’, Frauenweiler clay pit, Germany^130, 155^. *Resolution in phylogenetic analyses*: None. *Character states*: overlapping, sequential articulation between first dorsal fin pterygiophores, supraneurals, and supraoccipital crest; second infraorbital excluded from orbital margin; two sets of lateral processes on each side of first dorsal fin pterygiophore define a clear groove; distal head of second supraneural longer than that of first supraneural^155–157^. *Stratigraphy*: The ‘Fish shales’ lie within NP23^138^. *Absolute age estimate*: 29.62 Ma^4^. *Prior setting*: lognormal prior, minimum age 29.62 Ma, mean=1.0, S.D.= 1.498, upper 95% CI: 61.6 Ma^4^.

**xxxix. *Node*: Stem lineage *Chaetodon***, dating the MRCA of *Chaetodon kleinii* and *Prognathodes marcellae*. *First occurrence*: †*Chaetodon ficheuri* from the Saint-Denis du Sig, Raz-el-Aïn, Les Planteurs, and Eugène, Algeria^156^. *Resolution in phylogenetic analyses*: None. *Character states*: overlapping, sequential articulation between first dorsal fin pterygiophores, supraneurals, and supraoccipital crest; second infraorbital excluded from orbital margin; two sets of lateral processes on each side of first dorsal fin pterygiophore define a clear groove; distal head of second supraneural longer than that of first supraneural^156, 157^. *Stratigraphy*: Messinian^158–160^. *Absolute age estimate*: 7.1 Ma. *Prior setting*: lognormal prior, minimum age 7.1 Ma, mean=0.26, S.D.= 0.2, upper 95% CI: 8.9 Ma^20^.

**xl. *Node*: Stem lineage *Gazza***, dating the MRCA of *Gazza minuta* and *Leiognathus equula*. *First occurrence*: †*Euleiognathus tottori* from the Iwami Formation, Tottori Group, Japan^161, 162^. *Resolution in phylogenetic analyses*: none. *Character states*: long ascending processes of premaxillae; paddle-like expansions of neural and haemal spine of preural centrum 4; single supraneural; serrated anterior margins of fin spines; caniniform teeth^163^. The final character is unique to *Gazza* within leiognathids^161, 164^. The nesting of *Gazza* within the phylogeny of Leiognathidae indicates caniniform teeth are derived within the clade^163, 165, 166^. See Gill and Michalski^167^ who are sceptical that †*Euleiognathus is a* leiognathid*. Stratigraphy*: middle Miocene^161, 162^. *Absolute age estimate*: 11.6 Ma^168^. *Prior setting*: lognormal prior, minimum age 11.6 Ma, mean=0.8, S.D.= 1.0, upper 95% CI: 23.1 Ma^20^.

**xli. *Node*: Stem lineage Diodontidae**, dating the MRCA of *Diodon liturosus* and *Canthigaster rostrata*, which subtends the MRCA of Diodontidae and Tetraodontidae. *First occurrence*: †*Prodiodon tenuispinus*, †*P. erinaceus*, †*Heptadiodon echinus*, and †*Zignodon fornasieroae* from the Pesciara beds of ‘Calcari nummulitici’, Bolca, Italy^169^. *Resolution in phylogenetic analyses*: parsimony analysis of morphological characters resolves a clade containing †*Prodiodon tenuispinus*, †*P. erinaceus*, †*Heptadiodon echinus*, †*Zignodon fornasieroae*, *Diodon holocanthus*, and *Chilomycterus schoepfi*^169^. *Character states*: premaxillae fused along midline; dentaries fused along midline; jaws massive^169^. *Stratigraphy*: upper Ypresian [NP14]^94^. *Absolute age estimate*: 50 Ma^84^. *Prior setting*: lognormal prior, minimum age 50.0 Ma, mean=1.0, S.D.= 0.6, upper 95% CI: 57.3 Ma^20^.

**xlii. *Node*: Stem lineage of Balistidae**, dating the MRCA of *Balistes capriscus* and *Aluterus monoceros*, which subtends the MRCA of Balistidae and Monacanthidae. *First occurrence*: †*Gornylistes prodigiosus* from the Kuma Horizon, Krasnodar Region, Caucasus^170^. *Resolution in phylogenetic analyses*: None. *Character states*: ventral shaft of second spine-bearing dorsal pterygiophore absent; supraneural strut present between abdominal neural spine and final spine-bearing dorsal pterygiophore; four anal-fin pterygiophores anterior to the haemal spine of the third caudal vertebra^169^. *Stratigraphy*: Bartonian [NP17] Kumian regional stage^83^. *Absolute age estimate*: 37.2 Ma^84^. *Prior setting*: lognormal prior, minimum age 37.2 Ma, mean=1.0, S.D.= 0.42, upper 95% CI: 42.6Ma^20^.

**xliii. *Node*: Stem lineage Antennarioidei,** dating the MRCA of *Antennarius striatus* and *Ogcocephalus radiatus*. *First occurrence*: †*Eophryne bartutii* from the Pesciara beds of ‘Calcari nummulitici’, Bolca, Italy^171^. *Resolution in phylogenetic analyses*: None. †*Eophryne* was hypothesized to share common ancestry with Antennariidae based on overall morphological similarity^171^. The phylogenetic placement used for this calibration is conservatively applied to the more inclusive Antennarioidei. *Character states*: triradiate ectopterygoid; spatulate postmaxillary process of premaxilla^171–173^. *Stratigraphy*: upper Ypresian [NP14]^94^*. Absolute age estimate*: 50 Ma^84^. *Prior setting*: lognormal prior, minimum age 50.0 Ma, mean=1.0, S.D.= 0.6, upper 95% CI: 57.3 Ma^1^.

## 3. Supplementary results and discussion

### a. Phylogeny and classification of Acanthomorpha, and a new delimitation of taxonomic orders in Percomorpha

The maximum likelihood analyses of the 702-taxon and 1,084-taxon UCE datasets confidently resolve the phylogenetic relationships of all sampled acanthomorph families. The phylogeny inferred from the IQ-TREE analysis of the multiple-partition, concatenated 1084-taxon dataset is represented in Supplementary Figs. 1-25. The results of the five other phylogenetic analyses are available on Dryad. Bootstrap support values across the entire IQ-TREE phylogeny are high, with most nodes being supported by 100% bootstrap support and only 27 of 1,107 nodes (2.4%) having values ≤95%. Tree topologies inferred using the different methods described in *Section 1d* are largely consistent according to comparisons conducted in TOPD-FMTS v.4.6^39^ (see Supplementary Table 2). Robinson-Foulds metrics across all comparisons were low (≤0.033), as were nodal distances (≤1.046). Supplementary Table 2 lists the 19 species that differ in their phylogenetic relationships across the different analyses. Most of these species are found in areas of the tree that have been historically difficult to resolve, such as Anthiadidae, Carangidae and Apogonidae. Tree topologies inferred in IQ-TREE and RAxML using multiple-partition, 700-taxon alignments were identical.

The non-percomorph acanthomorphs are classified into eight taxonomic orders: Lampriformes, Percopsiformes, Polymixiiformes (a single genus, *Polymixia*) Zeiformes, Gadiformes, Stylephoriformes (a single species, *Stylephorus chordatus*), Trachichthyiformes, and Beryciformes (Fig. 1).

The new UCE-inferred phylogeny provides an opportunity to reevaluate the classification of Percomorpha to identify inclusive taxonomic groupings that reflect phylogenetic relationships. An initial effort at building classifications based on molecular phylogenies attempted to preserve the ordinal ranks of the clades such as the Pleuronectiformes, Tetraodontiformes, Atheriniformes, and Synbranchiformes^174^. This classification proposed a delimitation of 33 taxonomic orders in Percomorpha that each contain an average of only 7.4 families and 492.9 species, and resulted in 10% of all percomorph families unassigned to a taxonomic order^174^. In Figs. 1 and 2, Supplementary Figs. 1-25, and Supplementary Table 3 we offer an alternative delimitation of 13 clades that we treat as taxonomic orders to accommodate the classification of all 290 percomorph families. In the proposed classification, each taxonomic order comprises an average 2,226.8 species and 18.2 families.

The names of the orders in the proposed percomorph classification are consistent with conventions in ichthyological systematics by establishing a link with one of the constituent taxonomic families and using the “–iformes” suffix^175, 176^. For example, this delimitation of Blenniiformes includes the Blenniidae and 48 other families (e.g., cichlids, livebearers, and mullets) that were previously identified as a monophyletic group in molecular phylogenetic studies^1, 4, 15, 21, 22, 177^. Previously named Ovalentaria^22^, this diverse clade of percomorphs had been divided among seven orders^174^, one of which, Mugiliformes, included a single family, Mugilidae (mullets).

### b. Phylogeny of non-percomorph acanthomorphs

#### i. Monophyly and relationships of Paracanthopterygii

Similar to other phylogenomic studies, this UCE inferred phylogeny resolves a monophyletic and strongly supported Paracanthopterygii that includes *Polymixia* (beardfishes), Percopsiformes, Zeiformes, *Stylephorus chordatus* (Tube-eye), and Gadiformes (Fig. 1, Supplementary Fig. 1)^4, 15, 178^. While relationships within Paracanthopterygii differ among phylogenomic studies, the resolution of a clade containing *Polymixia* and Percopsiformes is the best-supported topology in the concordance factor analysis, with the 95% credible interval of the two alternative resolutions of *Polymixia* not overlapping with the optimal tree topology (Extended Date Fig. 1a, Supplementary Fig. 1).

#### ii. Phylogenetic relationships of Lampriformes

The new UCE phylogeny of Acanthomorpha resolves Lampriformes as the sister lineage of Paracanthopterygii (Fig. 1), which agrees with a previous phylogenetic analysis of UCE loci^4^. Other studies offer conflicting phylogenetic positions for Lampriformes; a phylogenomic analysis of ∼1,000 exons resolve lampriforms as the sister lineage of the Acanthopterygii^15^, while morphological phylogenies place the lampriforms as the sister lineage of all other Acanthomorpha^18^. The Bayesian concordance factor analyses result in overlapping 95% credible intervals for the phylogeny that places lampriforms as sister to Paracanthopterygii and the phylogeny that resolves lampriforms as the sister lineage of all other acanthomorphs (Extended Data Fig. 1b). The phylogeny that resolves lampriforms and Acanthopterygii as sister lineages scored the lowest concordance factors and the 95% credible interval of its concordance factors do not overlap with the credible interval of the phylogeny that resolves Lampriformes as the sister lineage of Paracanthopterygii (Extended Data Fig. 1b).

#### iii. Phylogenetic relationships of Gadiformes

The new UCE phylogeny of Gadiformes is similar to a recent study that utilized DNA sequences from more than 14,000 portions of protein coding genes ^179^. Results from the two analyses are congruent in resolving *Bregmaceros* (codlets) as the sister lineage of all other Gadiformes, but differ in three specific relationships. The UCE phylogeny resolves Muraenolepididae (eel cods) and Trachyrincidae (armored grenadiers) as sister lineages, but the exon phylogeny resolves *Melanonus* (pelagic cods) as the sister lineage of Muraenolepididae. The UCE phylogeny places Merlucciidae (merluccid hakes) as the sister lineage of an inclusive clade containing Phycidae (phycid hakes), Lotidae (hakes and burbots), and Gadidae (cods and haddocks), but the exon phylogeny resolves Merlucciidae as the sister lineage of a clade containing *Eulichthys polynemus* (Eucla cod), Muraenolepididae, *Melanonus*, Trachyrincidae, Moridae, *Macruronus* (blue grenadiers), *Lyconus* (Atlantic hakes), Bathygadidae (rattails), *Steindachneria argenta*, and Macrouridae. Lotidae is resolved as monophyletic in the exon phylogeny^179^, but this lineage is paraphyletic in the UCE phylogeny with *Lota lota* (Burbot) resolved as the sister lineage of a clade containing *Brosme brosme* (Cusk), *Molva* (lings), and Gadidae (true cods) (Fig. 1, Supplementary Fig. 1).

#### iv. Phylogenetic relationships of Acanthopterygii and resolution of the sister lineage of the hyper-diverse Percomorpha

Acanthopterygii includes Trachichthyiformes, Beryciformes, and Percomorpha (Fig. 1). The earliest molecular studies identified the sister lineage of percomorphs as either a monophyletic group consisting of Trachichthyiformes and Beryciformes^1, 20^ or the beryciform subclade Holocentridae (squirrelfishes)^21^. The UCE inferred phylogeny presented in this study and previous phylogenomic analyses resolve Trachichthyiformes as the sister lineage to all other acanthopterygians and Beryciformes, which includes Holocentridae, as the percomorph sister lineage^4, 15, 178, 180^ (Fig. 1, Supplementary Fig. 2). This topology is also the most supported hypothesis in the concordance factor analyses (Extended Data Fig. 1c). The alternative hypotheses, including the phylogeny that places Holocentridae as sister to Percomorpha, produced substantially lower Bayesian concordance factor scores with 95% credible intervals that do not overlap with the scores of the optimal phylogeny (Extended Data Fig. 1c). The UCE data presented here reject the hypothesis that Holocentridae is the sister lineage of Percomorpha.

### c. Phylogenetic relationships of Percomorpha

Percomorphs are classified in an astounding 290 taxonomic families^181^, which is more than living birds (252 families) and many more than mammals (167 families), squamates (58 families), amphibians (67 families), and turtles (11 families)^182–184^. By the middle of the 20th century the vast majority of percomorph families were delimited and considered monophyletic groups^175, 185^, but relationships among these delimited taxonomic families were woefully unresolved at the start of the 21st century^186, 187^. This lack of phylogenetic resolution among the nearly 300 taxonomic families led to the now famous analogy that Percomorpha was the “bush at the top” of the teleost phylogeny^188^. Since the start of the 21st century, however, molecular studies have offered dramatic resolution of percomorph relationships^1, 4, 15, 21^. The UCE phylogeny inferred in this study resolves the species sampled from 276 taxonomic families into one of twelve taxonomic orders that contain more than one family. All 12 of these inclusive lineages are monophyletic and strongly supported in the UCE phylogeny (Figs. 1,2, Supplementary Figs. 3-25). Relationships are consistent with previous efforts using Sanger sequenced loci and phylogenomic analyses, but no prior analysis included a high fraction (276 of 290) of the percomorph families.

The UCE phylogeny has strong node support along the backbone of Percomorpha including Eupercaria, which is an inclusive clade containing Perciformes, Centrarchiformes, Labriformes, Acropomatiformes, and Acanthuriformes (Fig. 2, Supplementary Figs. 16-25). Previous phylogenetic analyses strongly support monophyly of Eupercaria but provide little resolution and node support for the interrelationships of its constituent lineages^1, 15, 21^. The only node in the UCE percomorph phylogeny shared among the delimited 13 taxonomic orders that is not supported with a bootstrap score of 100% is the resolution of Acropomatiformes and Acanthuriformes as a clade, which is supported with a bootstrap score of 99% (Supplementary Fig. 22).

#### i. The deepest node in the phylogeny of percomorphs: the phylogenetic relationships of Ophidiiformes

Ophidiiformes (cusk eels) is resolved as the sister lineage to all other percomorphs (Fig. 1). Relationships within this species-rich lineage of mostly marine fishes are not well known and the delimitation of taxonomic families is in flux^189^. The UCE phylogeny resolves two clades in Ophidiiformes that correspond to the suborders Bythitoidei and Ophidioidei (Fig. 1, Supplementary Fig. 3). Within Bythitoidei, the families Dinematichthyidae (pygmy brotulas) and Bythitidae (livebearing brotulas) are both monophyletic (Supplementary Fig. 3). As discovered in previous studies^189–191^, the formerly recognized taxonomic families Parabrotulidae and Aphyonidae are phylogenetically nested in Bythitidae (Supplementary Fig. 3). Historically, Ophidioidei contained Ophidiidae and Carapidae, but the UCE phylogeny (Fig. 1, Supplementary Fig. 3) as well as previous studies using whole mtDNA genome sequences and Sanger sequenced nuclear genes resolve Carapidae as phylogenetically nested within Ophidiidae^1, 21, 119, 192^, prompting the synonymization of the family Carapidae with Ophidiidae^174^.

#### ii. Phylogenetic relationships of Batrachoididae

There is limited taxon sampling of Batrachoididae in the UCE phylogeny, but the subfamily Batrachoidinae (sampled with *Batrachoides pacifici* and *Opsanus tau*) is paraphyletic (Supplementary Fig. 3). Porichthyinae is paraphyletic in a phylogeny inferred from Sanger sequenced mitochondrial and nuclear genes^119^.

#### iii. Phylogeny of Gobiiformes

One of the most surprising results from molecular analyses of percomorph phylogeny was the discovery that gobies, cardinalfishes (Apogonidae), and nurseryfishes (Kurtidae) resolve in a strongly supported monophyletic group^20, 193, 194^. Subsequent molecular studies resolved two competing hypotheses of relationships among the major lineages of Gobiiformes: 1) *Kurtus* and Apogonidae resolved as a clade, or 2) Apogonidae as the sister lineage of a clade containing *Trichonotus* (sand divers) and Gobioidei^1, 4, 21, 194–198^. The UCE phylogeny in this study resolves two major lineages in Gobiiformes: the Apogonoidei that contains *Kurtus* and Apogonidae, and a clade containing *Trichonotus* and Gobioidei (Fig. 1, Supplementary Figs. 3,4). The relationships of the eight taxonomic families classified in Gobioidei are identical to phylogenies inferred using Sanger sequenced mtDNA and nuclear genes^197, 198^ (Fig. 1, Supplementary Fig. 4).

#### iv. Phylogenetic relationships of Scombriformes and Syngnathiformes

Scombriformes (e.g., tunas, cutlassfishes and butterfishes) and Syngnathiformes (e.g., seahorses, flying gurnards and dragonets) are resolved as sister lineages in the UCE phylogeny (Fig. 1), a result that is consistent with other molecular phylogenetic analyses^1, 4, 15^. This particular result has the surprising implication that relative to all other percomorph teleosts, tunas and seahorses are closely related.

Tunas, barracudas, cutlassfishes and swordfishes were historically classified in Scombroidei. Early phylogenetic analyses of Sanger sequenced mtDNA and nuclear genes resolved species classified in the traditional delimitation of Scombroidei into two distantly related percomorph lineages^199^. Xiphiidae (swordfishes), Istiophoridae (billfishes) and Sphyraenidae (barracudas) were resolved in the clade Carangiformes along with flatfishes, jacks and archerfishes (Fig. 1)^48^. The remaining lineages of the historical Scombroidei— Scombridae (tunas), Gempylinae (snake mackerels), Trichiurinae (cultassfishes) and *Pomatomus saltatrix* (Bluefish)—are resolved in the clade Scombriformes that also includes the stromatioids, e.g., Stromateidae (butterfishes) and Chiasmodontidae (swallowers)^29^. Other lineages in Scombriformes include the enigmatic *Icosteus aenigmaticus* (Ragfish) and *Amarsipus carlsbergi* (Bagless Glassfish), both of which are deep-branching, monotypic lineages that long evaded phylogenetic resolution among percomorphs (Fig. 1, Supplementary Fig. 5). The UCE phylogeny resolves *Lepidocybium flavobrunneum* (Escolar) as the sister lineage of a clade containing Gempylidae and Trichiuridae (Fig. 1, Supplementary Fig. 6). *Lepidocybium flavobrunneum* is traditionally classified in Gempylidae, which is paraphyletic in the UCE phylogeny as the remaining gempylids are more closely related to species of Trichiuridae (Fig. 1, Supplementary Fig. 6). This relationship has previously been resolved in a phylogenetic analysis using morphological data^122^.

Syngnathiformes as consisting of seahorses and pipefishes, flying gurnards, goatfishes, and dragonets was first resolved as a clade in phylogenetic analyses of Sanger sequenced mtDNA and nuclear genes^21, 200^. Prior to the application of molecular phylogenetics, Callionymidae (dragonets) and Draconettidae (draconettids) were classified as closely related to Gobiesocidae (clingfishes)^201–204^. As reflected in molecular phylogenies, Pietsch^98^ proposed that Dactylopteridae (helmet gurnards) were closely related to other syngnathiform lineages, but they were classified by others with the sculpins, rockfishes and scorpionfishes in Scorpaeniformes. Mullidae (goatfishes) were long classified in the wastebasket taxon Perciformes^175, 202, 205^ but thought to be closely related to Haemulidae (grunts) and Sparidae (porgies)^206^. The UCE phylogeny resolves two major lineages of Syngnathiformes: 1) a clade we delimit as Mulloidei that contains Pegasidae (seamoths), Dactylopteridae, Mullidae, Callionymidae, and Draconettidae and 2) a lineage we delimit as Syngnathoidei, consisting of Syngnathidae (pipefishes and seahorses), Solenostomidae (ghost pipefishes), Aulostomidae (trumpetfishes), Fistularidae (cornetfishes), Centriscidae (shrimpfishes) and Macroramphosidae (snipefishes) (Fig. 1, Supplementary Figs. 7-9).

#### v. Relationships among Blenniiformes, Synbranchiformes and Carangiformes

The UCE phylogeny resolves a strongly supported monophyletic group where Blenniiformes (e.g., blennies, cichlids, livebearers, and mullets) is the sister lineage of a clade containing Synbranchiformes (e.g., swamp eels, gouramies, snakeheads, and leaffishes) and Carangiformes (e.g., flatfishes, jacks and pompanos, billfishes, and barracudas). The discovery of this clade is one of the many dramatic results from molecular phylogenetics of percomorphs^1^, as this large and inclusive group contains more than 7,300 species that are classified into 92 taxonomic families.

#### vi. Blenniiformes: Phylogenetic relationships of Atherinoidei (Cyprinodontoidea, Belonoidea and Atherinoidea)

The monophyly of Blenniiformes—originally named Ovalentaria—resulted from a phylogenetic analysis of Sanger sequenced nuclear genes^22^. In the UCE phylogeny, Blenniiformes is monophyletic and contains two major clades: Atherinoidei (formerly Atherinomorpha) and all other lineages of Blenniiformes. Atherinoidei contains three major lineages: 1) Belonoidea (formerly Beloniformes, which includes needlefishes, medakas, halfbeaks, and flying fishes), 2) Cyprinodontoidea (formerly Cyprinodontiformes, which includes killifishes, rivulines, pupfishes, and livebearers), and 3) Atherinoidea (formerly Atheriniformes, which includes silversides and rainbowfishes). Phylogenomic analysis of our UCE dataset results in a well-resolved phylogeny of Atherinoidei, where every node along the backbone of the atherinoid tree is supported with a 100% bootstrap score (Fig. 1, Supplementary Fig. 10). Morphological phylogenies resolve Atherinoidea as the sister lineage of a clade containing Belonoidea and Cyprinodontoidea^149, 207^, but the UCE phylogeny resolves Atherinoidea and Belonoidea as sister lineages (Fig. 1, Supplementary Fig. 10).

Over the past 15 years, the number of taxonomic families recognized in Cyprinodontoidea (killifishes, rivulines, pupfishes, and livebearers) increased from 10 to 16^181, 186^. The recognition of these six additional families resulted from phylogenetic analyses of Sanger sequenced mtDNA and nuclear genes that overturned the results of morphological studies^208–210^ and demonstrated the non-monophyly of Cyprinodontidae (pupfishes) and Poeciliidae (livebearers). The UCE phylogeny (Fig. 1, Supplementary Fig. 10) is consistent with morphological phylogenetic analyses in the resolution of Rivulidae (New World rivulines), Aplocheilidae (Asian rivulines), and Nothobranchiidae (African rivulines) as a monophyletic group, as well as a clade containing Poeciliidae, Anablepidae (four-eyed fishes), *Valencia* (toothcarps), Procatopodidae (African lampeyes), Fundulidae (topminnows), Cyprinodontidae, *Profundulus* (Middle American killifishes), Goodeidae (goodeids), and *Cubanichthys* (Caribbean killifishes)^211, 212^. Morphological and molecular phylogenetic studies show congruence on specific relationships within Cyprinodontoidea. For example, the morphological analysis by Costa^212^ identified *Profundulus* and Goodeidae as a clade. This result is supported in the UCE phylogeny as well as studies from Sanger sequenced mtDNA and nuclear genes^208–210^. Among the notable differences between molecular and morphological phylogenies of cyprinodontoids is the resolution of Cyprinodontidae and Fundulidae as sister lineages (Fig. 1, Supplementary Fig. 10).

Belonoidea is monophyletic in the UCE phylogeny, but despite limited taxon sampling the families Zenarchopteridae and Hemiramphidae are found to be paraphyletic (Fig. 1, Supplementary Fig. 10). Five families of Belonoidea are recognized in Eschmeyer’s Catalog of Fishes^181^. Phylogenies resulting from analyses of morphological characters and Sanger sequenced mtDNA nest Scomberesocidae (sauries) well within Belonidae (needlefishes) and resolve Hemiramphidae (halfbeaks) as paraphyletic relative to Exocoetidae (flyingfishes)^213, 214^. In the UCE phylogeny, the hemiramphid *Oxyporhamphus micropterus* is more closely related to the sampled species of Exocoetidae than the other two sampled hemiramphids (Fig. 1, Supplementary Fig. 10), as previously inferred using morphological data^215^. The new phylogeny also resolves *Zenarchopterus dispar* as more closely related to the sampled belonid species *Xenentodon cancila* than to the other two sampled zenarchopterid species (Fig. 1, Supplementary Fig. 10).

There are 11 taxonomic families of Atherinoidea recognized in Eschmeyer’s Catalog of Fishes^181^, but Notocheiridae is well nested in Atherinopsidae in phylogenetic analyses of Sanger sequenced mtDNA and nuclear genes^216, 217^. Of the ten other families, eight are sampled in the UCE phylogeny. Missing from the analysis is the monotypic Dentatherinidae (Mercer’s tusked silverside) and Telmatherinidae (Celebes rainbowfishes). The UCE phylogeny is consistent with current classifications and previous molecular and morphological phylogenetic analyses in resolving Atherinopsidae (New World silversides) as the sister lineage of all other atherinoids^1, 21, 216, 218^ (Fig. 1, Supplementary Fig. 10). Another area of agreement in the UCE phylogeny and previous molecular analyses is the resolution of a clade containing the monogeneric Atherionidae (pricklenose silversides) and Phallostethidae (priapiumfishes)^217^ (Fig. 1, Supplementary Fig. 10).

There are differences in the recognition of taxonomic families of Atherinoidea in Eschmeyer’s Catalog of Fishes^181^ and Nelson et al.^176^ that involve lineages associated with Melanotaeniidae (rainbowfishes). Following Dyer and Chernoff^218^, Nelson et al.^176^ treat Bedotiidae (Madagascar rainbowfishes), Pseudomugilidae (blue eyes), and Telmatherinidae as subfamilies of Melanotaeniidae. This delimitation of Melanotaeniidae is not supported in the UCE phylogeny as Bedotiidae is resolved as the sister lineage of Atherinidae (Old World silversides) (Fig. 1, Supplementary Fig. 10). In previous analyses of Sanger sequenced mtDNA and nuclear genes, Melanotaeniidae is paraphyletic because the species *Cairinsichthys rhombosomoides* is resolved as the sister lineage of a clade containing all sampled species of Telmatherinidae and Pseudomugilidae^216, 217^; however, in the UCE phylogeny Melanotaeniidae— sampled with *Cairinsichthys* and *Iriatherina werneri*—is monophyletic with strong node support (Fig. 1, Supplementary Fig. 10). The non-monophyly of Melanotaeniidae in previous phylogenetic analyses is likely driven by limited phylogenetic informativeness of Sanger sequenced genes to resolve relationships among Atherinoidea, as evidenced by the low bootstrap and Bayesian posterior node support values associated with these phylogenies^216, 217^.

#### vii. Blenniiformes: Phylogenetic relationships of Cichlidae, Pholidichthys and Polycentridae

Within Blenniiformes, the UCE phylogeny resolves a clade containing Cichlidae, *Pholidichthys* (engineer gobies) and Polycentridae (leaffishes) (Fig. 1, Supplementary Fig. 11). Before the advent of molecular phylogenetics, the morphology of the pharyngeal jaw apparatus was cited as evidence to group Cichlidae with Labridae (wrasses and parrotfishes), Embiotocidae (surfperches), and Pomacentridae (damselfishes) in a group named Labroidei^219–222^. Phylogenetic analyses of ten Sanger sequenced nuclear genes strongly resolved *Pholidichthys* (engineer gobies) and Cichlidae as sister lineages, but these analyses did not confidently identify a sister lineage of this clade within the Blenniiformes^1, 22^. As recently delimited^223^, Polycentridae contains five species classified among four genera distributed in freshwater habitats in Africa and South America: *Afronandus sheljuzhkoi* (Africa), *Polycentropsis abbreviata* (Africa), *Polycentrus* (two species in South America), and *Monocirrhus polyacanthus* (South America). The resolution of the freshwater Polycentridae as the sister lineage of the clade containing *Pholidichthys*-Cichlidae provides insight into the origin of freshwater blenniiform lineages and the biogeographic relationships of freshwater fishes in South America and Africa^224, 225^.

#### viii. Blenniiformes: Relationships of Mugilidae, Ambassidae, Pseudochromidae, Congrogadidae and Pomacentridae

The phylogenetic relationships of Mugilidae (mullets) have long vexed phylogenetic studies of acanthomorphs, leading Stiassny^226^ to declare in a morphological phylogenetic study that “without mullets our lives would be a lot simpler.” Morphological analysis of branchial musculature supported a hypothesis that Mugilidae and Atherinoidei (formerly Atherinomorpha) are sister lineages^227^, but analysis of pelvic girdle morphology supported an affinity with other percomorph lineages^226^. The heterogeneity of relationships inferred from these two different anatomical systems appears explained in the results of molecular phylogenetic analyses with Mugilidae, Atherinoidei, or other “higher” percomorph lineages sensu Johnson and Patterson^18^ resolved in Blenniiformes. In Sanger sequencing efforts, the sister lineage of Mugilidae within Blenniiformes varies between Plesiopidae (longfins) and Embiotocidae (surfperches), always with weak node support^1, 22, 177^. However, analysis of the UCE dataset and phylogenomic inferences based on exon data^15^ both resolve Mugilidae and Ambassidae as sister lineages with strong node support (Fig. 1, Supplementary Fig. 11).

In phylogenetic studies of Sanger sequenced mtDNA and nuclear genes, Pseudochromidae (dottybacks) is resolved as paraphyletic relative to Plesiopidae (roundheads) and Pomacentridae (damselfishes)^1, 21^. In the UCE phylogeny, Pseudochromidae is paraphyletic because Congrogadidae (eelblennies) is resolved as the sister lineage of a clade containing the remaining species of Pseudochromidae, Plesiopidae, Pomacentridae, Grammatidae (basslets), Opistognathidae (jawfsihes), Gobiesocidae (clingfishes), and Blennioidei (blennies) (Fig. 1, Supplementary Fig. 11).

#### ix. Blenniiformes: Phylogenetic relationships of Grammatidae, Opistognathidae, Gobiesocidae and Blennioidei

The UCE phylogeny is congruent with trees inferred using exon data and Sanger sequenced mtDNA and nuclear genes in resolving a clade containing Grammatidae, Opistognathidae, Gobiesocidae, and Blennioidei^1, 15, 21^, but differs in resolving Embiotocidae (surfperches) as the sister lineage of this clade (Fig. 1, Supplementary Fig. 11). Grammatidae and Opistognathidae are successive outgroups to a clade containing Gobiesocidae and Blennioidei (Fig. 1, Supplementary Fig. 11). In the UCE phylogeny, the relationships among the families of Blennioidei—Tripterygiidae (triplefin blennies), Dactyloscopidae (sand stargazers), Blenniidae (combtooth blennies), Clinidae (kelp blennies), Labrisomidae (labrisomid blennies), and Chaenopsidae (tube blennies)—are nearly identical to those inferred from Sanger sequenced mtDNA and nuclear genes^1, 21^ (Fig. 1, Supplementary Fig. 11). However, a study with the largest sampling of blennioid species indicates that Labrisomidae and Chaenopsidae are both paraphyletic^228^.

#### x. Phylogenetic relationships of Synbranchiformes

Previous analyses of Sanger sequenced genes and phylogenomic datasets consistently resolve Synbranchoidei (swamp eels, earthworm eels, armored sticklebacks, and freshwater spiny eels) and Anabantoidei (gouramies, snakeheads, and Asian leaffishes) as a clade^1, 15, 21, 22, 177^.

One of the many surprises to emerge from the molecular phylogenies of percomorphs was the resolution of *Indostomus* (armored sticklebacks) as nested within Synbranchoidei^1, 177, 192, 229, 230^. Traditionally, *Indostomus* was classified with seahorses, shrimpfish, and sticklebacks in the polyphyletic Gasterosteiformes based on the presence of dermal bony plates along the side of the body, a reduced cranial skeleton, and a small mouth size^186^. Analysis of developmental ontogeny of several morphological traits led to the conclusion that *Indostomus* is closely related to sticklebacks (Gasterosteidae)^231^. However, *Indostomus* and sticklebacks are distantly related in all molecular phylogenies and Gasterosteidae is classified here in Perciformes. Aside from the resolution of *Indostomus* as a synbranchoid, specifically as the sister lineage of Synbranchidae (swamp eels), the UCE phylogeny of Synbranchoidei is identical to relationships inferred from morphology where Synbranchidae is resolved as the sister lineage of a clade containing Chaudhuriidae (earthworm eels) and Mastacembelidae (freshwater spiny eels)^232^ (Fig. 1, Supplementary Fig. 12).

Anabantoidei—delimited here as including Nandidae (Asian leaffishes), Badidae (chameleonfishes), Pristolepididae (Malayan leaffishes), Channidae (snakeheads), Helostomatidae (Kissing Gourami), Anabantidae (climbing gouramies), and Osphronemidae (gouramies and fighting fishes)—is monophyletic in the UCE phylogeny and characterized by the unique possession of teeth on the parasphenoid^223, 233^:^9–12^. The UCE phylogeny resolves a clade containing Pristolepididae, Nandidae, and Badidae (Fig. 1, Supplementary Figs. 12, 13), which is supported with a unique morphology of the ventral gill arch musculature where the *rectus ventralis IV* has an additional insertion on the ceratobranchial 5^223, 234^. The clade containing Badidae and Nandidae is supported with two morphological traits: 1) a distally divided haemal spine on the second preural centrum and 2) the restriction of the attachments cells in eggs to the ventral side of the yolk sac^223^: Figs. 2d-f, ^4^. Previous classifications grouped Pristolepididae and Badidae in Nandidae^186^. Given that the current delimitation of Nandidae and Pristolepididae each contain a single genus and Badidae includes two genera, future classifications can reduce monogeneric families and limit redundant taxonomic group names by classifying all these lineages in Nandidae.

The UCE phylogeny resolves Channidae, Helostomatidae, Anabantidae, and Osphronemidae as a strongly supported monophyletic group (Fig. 1, Supplementary Figs. 12,13). This phylogeny differs from those inferred using Sanger sequenced nuclear genes in resolving Helostomatidae as the sister lineage of a clade containing Anabantidae and Osphronemidae versus Helostomatidae as the sister lineage of Anabantidae or Osphronemidae^1, 223^.

#### xi. Phylogenetic relationships of Carangiformes

The UCE phylogeny is broadly congruent with other studies using UCEs^48^, but differs in the relationships of *Lates*, *Centropomus* (snooks), and *Sphyraena* (barracudas) (Fig. 1, Supplementary Fig. 14). Analyses of Sanger sequenced nuclear genes consistently supported the monophyly of Carangiformes^1, 21, 235, 236^, but did not strongly support the relationships among the major carangiform clades. The UCE phylogeny includes a clade containing Centropomidae (sampled with *Lates* and *Centropomus*), *Lactarius lactarius* (False Trevally), and *Sphyraena*. Previous analyses using UCEs also resolved Centropomidae as monophyletic^48^, but placed *Sphyraena* as the sister lineage of all other Carangiformes. A phylogenetic analysis of Carangiformes based on a dataset that combines a smaller number of UCE loci with discretely coded morphological characters resolves Centropomidae as paraphyletic with *Lates* and *Psammoperca*, forming a clade that is sister to a lineage containing *Centropomus*, *Lactarius* and *Sphyraena*^237^.

The UCE phylogeny resolves three major lineages of Carangiformes: 1) an unnamed clade (discussed above) containing Centropomidae, *Lactarius*, and *Sphyraena*, 2) an unnamed clade containing Polynemidae (threadfins) and Pleuronectoidei (flatfishes), and 3) a clade delimited here as Carangoidei that contains *Leptobrama* (beachsalmons), *Toxotes* (archerfishes), *Nematistius pectoralis* (roosterfish), *Mene maculata* (moonfish), *Xiphias gladius* (swordfish), Istiophoridae (billfishes), and Carangoidea (Fig. 1, Supplementary Figs. 14,15).

As resolved in several analyses using Sanger sequenced loci and UCE-based phylogenomic analyses^1, 21, 25, 48, 119^, the traditional delimitation of Carangidae (jacks and pompanos) is paraphyletic. Carangidae, limited to the subfamilies Naucratinae and Caranginae, is the sister lineage of a clade containing the former carangid subfamilies Scomberoidinae and Trachinotinae, Echeneidae (remoras), *Rachycentron canadum* (cobia), and *Coryphaena* (dolphinfishes) (Fig. 1, Supplementary Fig. 15).

#### xii. Eupercaria: relationships among Perciformes, Centrarchiformes, Labriformes, Acropomatiformes and Acanthuriformes

Phylogenetic analyses of Sanger sequenced loci led to the resolution of Eupercaria, an inclusive clade that contains more than 37% of species and 53% of all taxonomic families in Percomorpha^174^. While earlier studies resolved a clade containing Perciformes, Centrarchiformes, Labriformes, Acropomatiformes, and Acanthuriformes, phylogenies inferred from Sanger sequenced genes provided little resolution for the relationships among the major lineages of Eupercaria^1, 21^. Analysis of the UCE dataset results in a phylogeny where Eupercaria is resolved as a clade with strong support for the relationships among the major lineages; Perciformes, Centrarchiformes, and Labriformes are found to be successive outgroups to a clade containing Acropomatiformes and Acanthuriformes. All of the nodes along the backbone of the eupercarian phylogeny are supported with bootstrap values of 100% except the clade containing Acropomatiformes and Acanthuriformes (BSS = 99%) (Fig. 2, Supplementary Fig. 22). The phylogenomic analyses of the UCE loci resolve this substantial issue of the teleost phylogeny, which was once identified as the new bush on the top of the percomorph phylogeny^21^.

#### xiii. Phylogenetic relationships of Perciformes

Prior to the application of molecular data to investigate the phylogenetic relationships of teleosts, the vast majority of percomorphs were classified in the catchall taxon Perciformes. In addition to perciforms, percomorphs included morphologically unique and disparate lineages such as Pleuronectiformes (flatfishes), Tetraodontiformes (pufferfishes), and Gasterosteiformes (sticklebacks and seahorses). Any lineage of percomorph that was not as morphologically distinctive as a flatfish or seahorse was classified in Perciformes. At the end of the 20th century Perciformes contained more than 10,000 species and 160 taxonomic families^186^. Phylogenetic analyses of molecular datasets revealed that the perciform wastebasket contained lineages that spanned much of the backbone of the Percomorpha phylogeny. The UCE phylogeny offers a confirmation for the disassembly of Perciformes that involves the migration of more than 100 taxonomic families into nearly all other major lineages of Percomorpha.

An entirely different concept and delimitation of Perciformes emerged from molecular phylogenetic studies^1, 19, 21, 177, 187, 193, 238–241^. No longer a taxonomic wastebasket, Perciformes (as delimited here) contains more than 3,200 species that are classified among at least 53 taxonomic families and comprises a diverse array of lineages that not only were previously classified as perciforms, but also as scorpaeniforms and gasterosteiforms^3, 176^. In addition to the namesake Percidae (perches, walleyes, and darters), the newer concept of Perciformes resulting from molecular phylogenetic analyses includes seabasses, scorpionfishes, rockfishes, sculpins, searobins, weaverfishes, notothenioids, eelpouts, sticklebacks, flatheads, and many others (Supplementary Table 3).

Molecular analyses consistently resolve the traditional delimitation of Serranidae (sea basses), which contains more than 60 genera and 450 species, as non-monophyletic in Perciformes^1, 187, 241^. In the UCE phylogeny, taxa comprising the traditional Serranidae resolve in four or five different lineages (Supplementary Fig. 16). Epinephelidae (groupers) and Anthiadidae (basslets and anthians) are resolved as successive sister lineages or as a monophyletic group that is the sister lineage of all other Perciformes. The less inclusive delimitation of Serranidae is strongly resolved as the sister lineage of Bembropidae (duckbills). *Acanthistius* is resolved as either the sister lineage of Anthiadidae, where it is traditionally classified, or as the sister lineage of all other perciforms except Epinephelidae and Anthiadidae. The enigmatic *Niphon spinosus*, which has long avoided a confident phylogenetic or taxonomic placement^242, 243^, strongly resolves as the sister lineage of Percidae (Fig. 2, Supplementary Fig. 16).

In addition to the non-monophyly of the traditional Serranidae, the molecular phylogeny of Perciformes provides several strongly supported relationships that were not anticipated from inferences using morphological characters^18, 205, 244^. The UCE phylogeny is congruent with earlier analyses using Sanger sequenced loci but provides greater resolution and node support than obtained in earlier efforts. The commercially and scientifically important freshwater lineage Percidae is resolved in a clade that contains the marine *Niphon spinosus* and Trachinidae (weaverfishes), a lineage we refer to as Percoidei (Fig. 2, Supplementary Fig. 16, Supplementary Table 3). Notothenioidei includes the well-known Antarctic adaptive radiation as well as *Percophis brasiliensis* (Brazilian Flathead), which is distributed in the Atlantic Ocean from the coast of southern Brazil to central Argentina^245^. Notothenioidei is strongly resolved as the sister lineage of a new delimitation of Scorpaenoidei that includes almost all the lineages traditionally classified in Scorpaeniformes such as Platycephalidae (flatheads), Scorpaenidae (scorpionfishes and rockfishes), Triglidae (searobins), Bembridae (deepwater flatheads), Platycephalidae (flatheads), Hoplichthyidae (ghost flatheads), Hexagrammidae (greenlings), Trichodontidae (sandfishes), Cyclopteridae (lumpfishes), Liparidae (snailfishes), and Cottidae (sculpins). In addition to these classic scorpaeniform lineages the new delimitation of Scorpaenoidei includes Zoarcoidea (eelpouts, ronquils, and wolffishes) and Gasterosteoidea (sand eel, tubesnouts, and sticklebacks) (Fig. 2, Supplementary Figs. 17, 18).

The UCE phylogeny of Scorpaenoidei is quite different from trees inferred using a dataset of morphological characters and Sanger sequenced mtDNA and nuclear genes^246^. For example, Smith et al.^246^ resolved Platycephalidae as the sister lineage of Triglidae, but the UCE phylogeny places Platycephalidae as the sister lineage of all other scorpaenoids (Fig. 2, Supplementary Fig. 17). In the UCE tree there are three major clades of Scorpaenoidei that contain multiple taxonomic families, but several families resolve along the backbone of the scorpaenoid phylogeny. Scorpaenoidea is here delimited as a strongly supported clade that includes *Normanichthys crockeri* (Mote Sculpin), Neosebastidae (gurnard scorpionfishes), Congiopodidae (horsefishes), Synanceiidae (stonefishes), and Scorpaenidae (Fig. 2, Supplementary Fig. 17). Bembridae, Triglidae, and Anoplopomatidae (sablefishes) resolve as successive sister lineages to a large clade containing all the lineages classified in Cottoidea (sculpins), Gasterosteoidea, and Zoarcoidea (Fig. 2, Supplementary Figs. 18, 19, Supplementary Table 3).

Gasterosteoidea and Zoarcoidea resolve as sister lineages with strong node support, which is a result supported in earlier phylogenetic analyses of Sanger sequenced mtDNA and nuclear genes^1, 21^. Within Zoarcoidea the Bathymasteridae (ronquils) are resolved as the sister lineage of all other zoarcoids. In an earlier study using Sanger sequenced loci, Bathymasteridae was resolved as paraphyletic^1^, but the UCE phylogenetic analyses sample all three bathymasterid genera and strongly support the lineage as a monophyletic group (Supplementary Fig. 18). *Scytalina cerdale* (Graveldiver) is nested well within Stichaeidae (Supplementary Fig. 18) while *Gymnoclinus cristulatus* (Trident Prickleback) is distantly related to other stichaeids (Supplementary Fig. 19), resolving as the sister lineage of Neozoarcidae. Previous phylogenetic analyses of morphology and Sanger sequenced mtDNA and nuclear genes resolve Stichaeidae as paraphyletic^247–249^. The zoarcoid lineages *Cryptacanthodes* (wrymouths), Lumpenidae (pricklebacks), *Zaprora silenus* (Prowfish), Opisthocentridae (ocellated blennies), *Ptilichthys goodei* (Quillfish), Pholidae (gunnels), *Gymnoclinus*, Neozoarcidae, Anarhichadidae (wolffishes), and Zoarcidae (eelpouts) resolve as a strongly supported clade that is the sister lineage of the paraphyletic Stichaeidae (Fig. 2, Supplementary Figs. 18,19).

#### xiii. Phylogenetic relationships of Centrarchiformes

Phylogenetic analyses of Sanger sequenced mtDNA and nuclear genes led to the discovery of Centrarchiformes, a clade of approximately 300 species classified in at least 16 taxonomic families^1, 21, 236, 250, 251^. Consistent with these earlier phylogenetic studies^1, 21, 250, 251^, the UCE phylogeny resolves *Percalates* as the sister lineage of all other Centrarchiformes (Fig. 2, Supplementary Fig. 20). Terapontoidei includes Girellidae (nibblers), Scorpididae (halfmoons), *Dichistius*, Microcanthidae, *Oplegnathus* (knifejaws), Kyphosidae (sea chubs), *Kuhlia* (flagtails), and Terapontidae (grunters) (Supplementary Fig. 20). Centrarchoidei is delimited here as a clade that includes *Enoplosus armatus* (Oldwife) and Percichthyidae (temperate perches), Centrarchoidea which includes Centrarchidae (sunfishes, blackbasses, and pygmy sunfishes) and Sinipercidae (Chinese perches), and Cirrhitoidea which includes Cirrhitidae (hawkfishes), Latridae (trumpeters), *Chironemus* (kelpfishes), *Cheilodactylus*, and *Aplodactylus* (marblefishes) (Fig. 2, Supplementary Fig. 20, Supplementary Table 3).

The classification of Centrarchiformes is dynamic and unsettled, reflected in part by a high proportion of families that contain a single genus. Molecular phylogenies consistently resolve two sets of traditionally delimited centrarchiform families as non-monophyletic. First, the two species of *Percalates* were traditionally classified as Percichthyidae^244^, but resolve as the sister lineage of all other centrachiforms^1, 21, 174, 236, 250^. There is no described taxonomic family to accommodate the classification of *Percalates.* Second, the classification of families within Cirrhitoidea was dramatically realigned as a result of molecular phylogenetic analyses. Traditionally, Cheilodactylidae (morwongs) contained three to five genera and approximately 20 species^202, 252^. Previous phylogenetic analyses of mtDNA genes, morphological characters, and UCE data resolved Cheilodactylidae as polyphyletic, with all but two of the species traditionally classified as cheilodactylids nested within a paraphyletic Latridae^253–255^. The findings of these phylogenetic analyses resulted in a transfer of these species to Latridae from Cheilodactylidae. The new UCE phylogeny is consistent with these previous results, but differs from the phylogeny presented in Ludt et al.^255^ in resolving *Aplodactylus* rather than *Chironemus* as the sister lineage of *Cheilodactylus* (Fig. 2, Supplementary Fig. 20).

#### xiv. Phylogenetic relationships of Labriformes

The UCE phylogeny strongly resolves a clade we delimit as Labriformes which contains two major lineages: Uranoscopoidei and Labroidei (Supplementary Fig. 21). As resolved in previous phylogenies inferred from Sanger sequenced nuclear genes^198^, Uranoscopoidei contains Uranoscopidae (stargazers), Ammodytidae (sandlances), and Pinguipedidae (sandperches) as successive sister lineages to a clade containing the enigmatic *Cheimarrichthys fosteri* (Torrentfish) and Leptoscopidae (southern sandfishes) (Fig. 2, Supplementary Fig. 21). Morphological studies place *Cheimarrichthys* as closely related to Pinguipedidae or as the sister lineage of all “trachinoids,” which include all the lineages delimited here as Uranoscopoidei^256–258^. The hypothesis that *Cheimarrichthys* and Pinguipedidae share common ancestry was rejected through the discovery that *Cheimarrichthys* shares more derived morphological character states with Leptoscopidae than any other “trachinoid” or uranoscopoid lineage^259^. Reflective of their shared common ancestry, *Cheimarrichthys* and Leptoscopidae have a similar geographic distribution; *Cheimarrichthys* is an anadromous species widely distributed among the rivers of New Zealand and leptoscopids are distributed along the Pacific and Indian coasts of Australia and New Zealand ^260, 261^.

Labroidei includes the sister lineages *Centrogenys vaigiensis* (False Scorpionfish) and Labridae (wrasses and parrotfishes) (Fig. 2, Supplementary Fig. 21). As discussed above regarding Blenniiformes, the species rich marine Labridae and the freshwater Cichlidae were hypothesized as closely related based on the morphology of the modified “labroid” pharyngeal jaw apparatus^219–221^, which is now known to have originated multiple times in Percomorpha^22^. Previous molecular phylogenetic analyses using Sanger sequenced nuclear genes demonstrated that labrids and cichlids are not closely related despite both having modified “labroid” pharyngeal jaws, but these early molecular studies resulted in an ambiguous and poorly supported resolution of the species rich Labridae^1, 22^. The resolution of Labridae and *Centrogenys vaigiensis* as sister lineages in the UCE phylogeny is interesting as both lineages have all three components of the modified labroid pharyngeal jaw apparatus^22^.

#### xv. Phylogenetic relationships of Acropomatiformes

The discovery of Acropomatiformes resulted entirely from phylogenetic analyses of Sanger sequenced mtDNA and nuclear and genes^1, 198, 236, 245^ and was never intimated through the study of morphology^205, 244^. The lineage is an odd assortment of deep sea and near shore percomorph lineages that were previously classified in the wastebasket version of Perciformes and the demonstrably polyphyletic Trachinoidei^1, 198, 245^. While molecular studies consistently resolve Acropomatiformes as a clade, its monophyly is not strongly supported with Bayesian posteriors or bootstrap values and relationships within the clade differ dramatically across studies^1, 198, 236, 245, 262^.

The UCE phylogeny resolves Acropomatiformes as monophyletic with 100% bootstrap support (Fig. 2, Supplementary Fig. 22). All but two nodes within Acropomatiformes are present in 100% of the bootstrap replicates. First, the clade containing *Bathyclupea* (deepsea herrings), *Champsodon* (gapers), Creediidae (sandburrowers), and Hemerocoetidae (signalfishes) is supported with a bootstrap score of 97%. Second, the clade containing *Scombrops* (gnomefishes), Symphysanodontidae (slopefishes), Epigonidae (deepwater cardinalfishes), and Howellidae (oceanic basslets) is supported with a 99% bootstrap score (Fig. 2, Supplementary Fig. 22). The relationships among acropomatiforms in the UCE tree are quite different from any previous study and include the non-monophyly of Polyprionidae (wreckfishes) as *Polyprion* is resolved as the sister lineage of a clade containing *Glaucosoma* (pearl perches), Pempheridae (sweepers), and *Lateolabrax* (Asian seaperches) while *Stereolepis* is the sister lineage of a clade containing *Banjos* (banjofishes) and Pentacerotidae (armorheads). The resolution of Howellidae and Epigonidae as sister lineages in the UCE phylogeny is interesting in the context of four osteological traits shared between these two lineages and the possibility of additional shared traits in Howellidae and other acropomatiform lineages^263^. Future phylogenetic studies will round out the sampling of Acropomatiformes by including *Dinolestes lewini* (Long-finned Pike), Malakichthyidae (temperate ocean-basses), and Synagropidae (splitfin ocean-basses) to take advantage of the strong phylogenetic resolution provided by the UCE loci for this clade.

#### xvi. Phylogenetic relationships of Acanthuriformes

The first wave of molecular phylogenetic analyses of percomorphs was based on whole mitochondrial genomes and small sets of Sanger sequenced mitochondrial and nuclear genes. The major lineages of percomorphs and their composition began to emerge because of these molecular phylogenetic analyses. The monophyly of the major percomorph lineages and their relationships to one another are well resolved in the UCE phylogeny and we delimit these inclusive clades as taxonomic orders (e.g., Gobiiformes, Scombriformes, and Perciformes). Hampering these earliest molecular phylogenetic studies of percomorphs was a lack of phylogenetic resolution for relationships within and between major lineages of Eupercaria which include Perciformes, Centrarchiformes, Acropomatiformes, and what Smith et al.^25^ delimit as Acanthuriformes. The lack of phylogenetic resolution was particularly acute in Acanthuriformes, a lineage that includes more than 2,320 species classified in 56 taxonomic families. The phylogenies resulting from earlier molecular studies resolved taxonomic families within Acanthuriformes as monophyletic with high bootstrap support [e.g., Chaetodontidae (butterflyfishes), Acanthuridae (surgeonfishes), and Sparidae (porgies)], but had very poor support for relationships among these lineages^1, 21, 25^.

Acanthuriformes is monophyletic in the UCE phylogeny with strong bootstrap node support; however, six nodes along the backbone of the acanthuriform phylogeny have bootstrap scores ranging from 70% to 94% (Fig. 2, Supplementary Figs. 22-24). Despite this lower support for a small portion of the acanthuriform phylogeny, the UCE tree provides insight into several issues not resolved in earlier molecular studies of percomorph phylogenetics. First, Gerreidae (morjarras), which long evaded phylogenetic resolution, is the sister lineage of all other Acanthuriformes (Fig. 2, Supplementary Fig. 22). Second, Moronidae (temperate basses) and Sillaginidae (whitings) are resolved here as sister lineages (Fig. 2, Supplementary Fig. 22). Though they were resolved as a closely related in earlier molecular studies^1^, Lutjanidae and Haemulidae are distantly related in the UCE phylogeny (Fig. 2, Supplementary Fig. 22,23). Finally, ten lineages identified as *incertae sedis* within Eupercaria by Betancur-R et al.^174^ are phylogenetically resolved with moderate to high bootstrap support in Acanthuriformes (Fig. 2, Supplementary Figs. 22-24).

One of the most surprising findings from molecular studies of the teleost phylogeny was the resolution of Lophioidei (anglerfishes, formerly Lophiiformes) and Tetraodontoidei (puffers and molas, formerly Tetraodontiformes) as sister lineages^1, 4, 15, 19–21, 192^. An analogous rearrangement within Mammalia in terms of divergence times might be recognition of a clade of marsupials as the sister lineage of primates. In morphology-based classifications, the lophioids were placed in Paracanthopterygii^186, 264^, phylogenetically distant from other percomorph lineages. This migration of lophioids as paracanthopterygians into a derived clade of percomorphs is among the most significant changes in 21st century vertebrate phylogenetics. While the discovery that Lophioidei and Tetraodontoidei are closely related was based on phylogenetic analyses of molecular data, subsequent investigation of their morphology identified several soft tissue characters that are likely synapomorphies of a lophioid-tetraodontoid clade^265^. It has also been discovered that the larvae of these two lineages exhibit unique morphology and pigmentation^266^. Congruent with earlier studies, the UCE tree resolves an inclusive lineage containing *Siganus* (rabbitfishes), Scatophagidae (scats), Priacanthidae (bigeyes), Cepolidae (bandfishes), and Caproidae (boarfishes) as the sister lineage of the Lophioidei-Tetraodontoidei clade. These nodes in the UCE phylogeny are all characterized by high bootstrap support values (Fig. 2, Supplementary Figs. 24,25).

#### xvii. Acanthuriformes: phylogenetic relationships of Tetraodontoidei

Some of the earliest phylogenetic analyses of ray-finned fishes focused on relationships within Tetraodontoidei, resulting in numerous phylogenetic analyses based on morphological and molecular datasets^21, 169, 267–274^. While there are important differences among nearly all the phylogenies of tetraodontoids, most analyses consistently resolve three to four sets of sister lineages that include Triacanthodidae (spikefishes)-Triacanthidae (triplespines), Diodontidae (porcupinefishes)-Tetraodontidae (puffers), Balistidae (triggerfishes)-Monacanthidae (filefishes), and Aracanidae (deepwater boxfishes)-Ostraciidae (boxfishes)^1, 21, 169, 267, 274^, but all these phylogenies differ in how these lineages relate to one another. They are also incongruent regarding the relationships of *Triodon macropterus* (Threetooth Puffer) and Molidae (molas and ocean sunfishes). The UCE phylogeny and most of the earlier molecular phylogenies do not resolve the subclade formerly named Tetraodontoidei, which contained Triodontidae, Molidae, Diodontidae, and Tetraodontidae^1, 21, 269–272^. The monophyly of this group was inferred, in part, based on the beak-like teeth in the upper and lower jaws and on a non-protractile upper jaw^169^.

The relationships of Tetraodontoidei in the UCE phylogeny are strongly supported with high bootstrap values at every node (Fig. 2, Supplementary Fig. 24), but differ from all previous topologies inferred using morphology, molecules, or combined morphological and molecular datasets. The UCE phylogeny resolves the relationships of *Triodon* and Molidae and contains three major lineages: 1) a clade containing *Triodon*, Aracanidae, and Ostraciidae, 2) a clade containing Triacanthodidae, Triacanthidae, Balistidae, and Monacanthidae, and 3) a clade containing Molidae, Diodontidae, and Tetraodontidae (Fig. 2, Supplementary Fig. 24).

#### xviii. Acanthuriformes: phylogenetic relationships of Lophioidei

The UCE phylogeny includes 16 of 18 recognized taxonomic families of Lophioidei; the monotypic Centrophrynidae and Lophichthyidae are not sampled. Phylogenies of Lophioidei inferred from morphological characters, whole mtDNA genome sequences, and Sanger sequenced mtDNA and nuclear genes all differ from one another^1, 21, 275, 276^. Congruent with all other phylogenetic analyses, the UCE phylogeny resolves the Lophiidae (goosefishes) as the sister lineage of all other Lophioidei (Fig. 2, Supplementary Fig. 25). It is also congruent with a previous phylogenetic analysis of lophioids in resolving Ogcocephalidae (batfishes) as the sister lineage of a clade previously named Antennarioidei that contains Antennariidae (frogfishes), Tetrabrachiidae (tetrabranchid frogfishes), and Brachionichthyidae (handfishes)^277^. All nodes along the backbone of the UCE lophioid phylogeny have high bootstrap support (Fig. 2, Supplementary Fig. 25).

The UCE phylogeny is congruent with previous molecular phylogenetic analyses in resolving Chaunacidae (coffinfishes) as the sister lineage of the eleven taxonomic families that comprise the deepsea anglerfishes^1, 21, 275^, delimited here as Ceratioidea (Fig. 2, Supplementary Fig. 25, Supplementary Table 3). The UCE phylogeny and previous molecular analyses using whole mitochondrial genomes resolve Thaumatichthyidae (wolftrap anglers) as paraphyletic with *Lasiognathus* nested within Oneirodidae (dreamers)^275^. Within the ceratioids, the UCE phylogeny resolves *Neoceratias spinifer* (Toothed Seadevil), Linophrynidae (leftvents), and Ceratiidae (seadevils) as a monophyletic group (Fig. 2, Supplementary Fig. 25). These three lineages all exhibit male obligate sexual parasitism and dramatically altered their immune systems through losing the capacity for somatic diversification of antigen receptor genes^278^. All previous morphological and molecular phylogenetic analyses of ceratioids resulted in the non-monophyly of the lineages exhibiting obligate male sexual parasitism, implying multiple origins of this unique reproductive mode^275, 278, 279^. In contrast, the UCE phylogeny implies a single evolutionary origin of this unique trait (Fig. 2, Supplementary Fig. 25).

### d. ASTRAL-III species tree analysis

The ASTRAL-III inferred summary species tree is similar to the phylogeny inferred from the concatenated data analyses (Figs. 1,2, Supplementary Figs. 1-25) in resolving Lampriformes as the sister lineage of a monophyletic Paracanthopterygii, monophyly of Acanthopterygii, and very similar relationships among the major lineages of Percomorpha (Supplementary Fig. 26). The non-collapsed species tree is available on Dryad. The ASTRAL-III species tree differs from the concatenated-dataset, maximum likelihood phylogeny in the inference of Trachichthyiformes and Beryciformes as a clade that is sister to Percomorpha, the gadiform *Bregmaceros* as the sister lineage of a clade containing *Stylephorus* and all other gadiforms, the placement of *Centrogenys*, and the non-monophyly of Acropomatiformes and Labriformes. The ASTRAL-III species tree also does not resolve Gerreidae as the sister lineage of all other Acanthuriformes. In general, the phylogenetic placement of major eupercarian clades in the tree estimated under the multi-species coalescent model differs from the topology of the concatenated-dataset, maximum likelihood trees, but nodal support values were significantly stronger for the backbone relationships inferred using maximum likelihood methods (Supplementary Figs. 1-26).

### e. Divergence-time estimates

For both the 702-and 1,084-taxon phylogenies, our relaxed-clock molecular dating analyses estimated similar stem lineage ages across the multiple subsamples of UCE loci, with overlapping 95% highest posterior densities for most nodes (Supplementary Table 4). There were also no observable differences in the median node heights reported in Maximum Clade Credibility trees built using post-burn-in trees that were randomly sampled (as in the tree represented in Figs. 1,2), or using post-burn-in trees that were systematically sampled after a certain number of MCMC iterations (Supplementary Table 4).

Our age estimates are largely in agreement with those presented in previous phylogenomic analyses of ∼1,000 UCE loci^4^ and ∼1,100 exons^15^ (Supplementary Fig. 29). For most major clades, our analyses inferred slightly older stem ages, but with 95% highest posterior densities that overlap with the dates reported in previous studies (Supplementary Fig. 29). However, our analyses inferred comparably younger stem lineage ages for *Polymixia*, Percopsiformes, and clades within Eupercaria^4, 15^. Our stem lineage age estimates are significantly older for Gobiiformes and the Syngnathiformes-Scombriformes clade^4^. Perhaps as a result of incongruent phylogenetic topologies, our analyses inferred younger stem age estimates for Acanthuriformes than estimated in the phylogenomic analysis of exon data^15^.

We similarly estimate crown lineage ages for major clades with 95% highest posterior densities that largely overlap with previous phylogenomic analyses^4, 15^. Exceptions include the crown age estimates of Acanthuriformes, Syngnathiformes and Scombriformes, Trachichthyiformes, Beryciformes, Gobiiformes, Syngnathiformes and Blenniiformes. Many of the discrepancies in crown ages likely result from differences among inferred phylogenetic trees (especially within Eupercaria), or from the relatively sparse taxon sampling in previous phylogenomic studies^4, 15^.

### f. Diversification rate analyses

Our results do not support a significant effect of the K-Pg on acanthomorph lineage diversification rates. TESS-CoMET results indicate constant tree-wide diversification rates through most of the history of Acanthomorpha, with no evidence of a mass extinction (Extended Data Fig. 3). Although Extended Data Fig. 3e suggest a brief increase in the global net-diversification (speciation minus extinction) rate at approximately 50 mya (followed by a decline), we observe very low Bayesian support for these rate shifts (Extended Data Fig. 3b,d). Parameter estimates for speciation and extinction rates converged in all TESS-CoMET runs (ESS values > 200 and Geweke statistics within the 95% confidence intervals for all time points). Convergence results for the CoMET run visualized in Extended Data Fig. 3 are available in Supplementary Fig. 30. All replicate analyses produced diversification plots that were virtually indistinguishable from one another. The results of the Gelman-Rubin test, which calculates the ratio of within-sample variance to between-sample variance, ensured that rate parameters and shift time parameters converged across the independent, replicate analyses (Rubin-Gelman statistic < 1.05) (Supplementary Fig. 31).

The CoMET results do report moderately supported shifts (2 ≤ 2ln Bayes Factors ≤ 6) in speciation and extinction rates in the last 10 million years (Extended Data Fig. 3a-e). We strongly suspect, however, that these shifts are an artefact of the CoMET model, perhaps due to how it accounts for incomplete taxon sampling. This suspicion is supported by diversification patterns reported in numerous other studies^10, 280–282^, which all demonstrate (often dramatic) rate shifts close to the present-day. It remains unclear how these recent patterns may affect the inference of older rate shifts. May et al.^283^ describe that more recent mass extinction events may cause a “shadow effect” that diminishes the signal of earlier mass extinctions. As it stands, we have no evidence that the rate shift we observe in the mid-to late-Neogene could be overshadowing or weakening a signal for an earlier shift in diversification rates.

Indeed, our tests of relative model fit in TESS corroborate CoMET’s estimate of constant diversification rates (Extended Data Table 1). Pairwise comparisons of marginal likelihoods using Bayes Factors suggested that a constant branching-process model best represents the pattern of diversification in Acanthomorpha. In general, this test of relative model fit strongly favored models assuming uniform sampling methods over diversified methods. Therefore, after the model assuming a constant birth-death process and uniform sampling, the next most-favored branching process models were an episodic model that accounted for a rate shift 50 mya (in accordance with the blip in net-diversification rates from the CoMET analysis) and a model with speciation rates that decreased through time, both of which assumed uniform sampling. The predictive distributions of the gamma statistic, number of taxa, and lineage-through-time (LTT) plots for the three most-favored branching process models are reported in Supplementary Fig. 32. The models assuming a tree-wide rate shift at 50 my or a decreasing speciation rate produce posterior-predictive distributions that are in accordance with the empirical estimates from the time-calibrated acanthomorph phylogeny (Supplementary Fig. 32b,c,e,f,h,i). However, the distribution under a simpler, constant rate model is also in agreement and appears to produce the most similar LTT plots to the empirical data (Supplementary Fig. 32a,d,g).

There is controversy surrounding the popular macroevolutionary modeling program BAMM^284, 285^, so indications of lineage diversification rate heterogeneity in the time-calibrated acanthomorph phylogeny are viewed with a degree of skepticism. Since the size of the phylogeny is so large, in every analysis BAMM estimated multiple shift configurations that were relatively equiprobable. We therefore chose not to examine configurations with the maximum a posteriori probability. Rather, for each analysis we chose to interrogate the maximum shift credibility (MSC) configuration, which considers only configurations sampled in the reversible-jump MCMC simulations and maximizes the product of the marginal branch-specific shift probabilities. In the 23 BAMM analyses that were conducted using the time-calibrated 1,084-taxon phylogeny, MSC configurations inferred anywhere between 22 and 33 well-supported rate shifts (Supplementary Fig. 33). The regression line in Supplementary Fig. 33 suggests that the prior number of shifts specified in BAMM does not dramatically affect the estimated number of rate shifts in the posterior prediction (slope = 0.1049). The average number of rate shifts among the MSC configurations was approximately 28. Bayes factor comparisons for each analysis, however, supported models with a number of diversification rate shifts in the upper 30s or low 40s, depending on the specified priors. This was true even for analyses with prior models that expected one rate shift in the phylogeny.

In Extended Data Fig. 4, we present the MSC configuration resulting from an analysis with a prior model that expected 15 rate shifts and ran a 2-chain MCMC simulation with a deltaT value of 0.5. Log-likelihood values across all analyses were similar, but the analyses with prior models assuming 15 or 40 expected shifts displayed the highest effective sample sizes (ESS) for their log-likelihoods and for the number of shift events. Since the estimated number and positions of the rate shifts were similar across analyses with different priors, we take note of the 19 clades with leading branches that were estimated to have undergone rate shifts by at least 16 (≥70%) of the 23 BAMM analyses. The 19 shared clades, several of which (e.g. zoarcoids, darters, notothenioids, labrids, cichlids and gobies) have previously been described as having undergone radiations, are labeled with blue numbers in Extended Data Fig. 4. They include: 1) Dinematichthyidae, 2) Apogoninae (Apogonidae to the exclusion of *Pseudamia*), 3) Gobiidae and Oxudercidae, 4) Mastacembelidae, 5) the clade defined by Scopthalmidae and Soleidae, 6) Carangidae (to the exclusion of *Seriola*), 7) Pseudocrenilabrinae (in Cichlidae), 8) Poeciliidae, 9) Labridae, 10) Sciaenidae, 11) Chaetodontidae, 12) Acanthuridae (to the exclusion of *Naso*), 13) Anthiadinae and Epinephelidae, 14.) darters (Etheostomatinae), 15) the clade defined by “Nototheniidae” and Channichthyidae (see Supplementary Fig. 17 for clarification), 16) *Sebastes*, 17) the clade defined by Trichodontidae and Psychrolutidae, 18) Lycodinae (in Zoarcidae), and 19) the clade defined by Stichaeidae and Zoarcidae. We chose to present the configuration in Extended Data Fig. 4 because the analysis assuming 40 rate shifts had more rate shifts beyond these 19 clades than the analysis assuming 15 expected rate shifts. Still, the MSC configuration presented in Extended Data Fig. 4 highlights eight additional rate shifts that are labelled with black numbers and are not present in this list of 19. Furthermore, while the MSC configuration in Extended Data Fig. 4 identifies rate shifts occurring within Acanthuridae and Zoarcidae, these shifts occur along a different-yet-nearby internal branch than in most of the other configurations. Specifically, the configuration in Extended Data Fig. 4 flags one rate shift in the branch leading to Acanthuridae to the exclusion of *Naso* and *Prionurus*, and another in the lineage defined by *Bothrocara* and *Lycodes concolor* (within Zoarcidae).

The fact that many of the rate shifts identified by BAMM occur along tipwards branches may indicate that the species richness of Acanthomorpha is driven by relatively recent, more phylogenetically or geographically localized radiations instead of a burst of diversification after the K-Pg. This pattern does not seem to be unique to acanthomorphs; numerous radiations have shown to have a similar pattern of constant diversification across the K-Pg boundary despite bursts in the origin of disparity in morphological and ecological traits^7^. Nevertheless, there is a chance that this pattern is the by-product of analytical corrections for incomplete sampling. Such corrections may reduce the statistical power to infer shifts in diversification rates and increase the probability of Type II error when rate variation is present^286^. It is also known that Bayesian methods of diversification rate estimation, including BAMM and TESS, penalize parameter-rich diversification models and favor less-complex models even under liberal priors. Given the infinite number of possible diversification scenarios that could result in the observed phylogeny^287^, it is therefore possible that TESS estimated a constant-rate model to best fit our data because of its simplicity rather than its accuracy.

---

**Supplementary Table 1:**
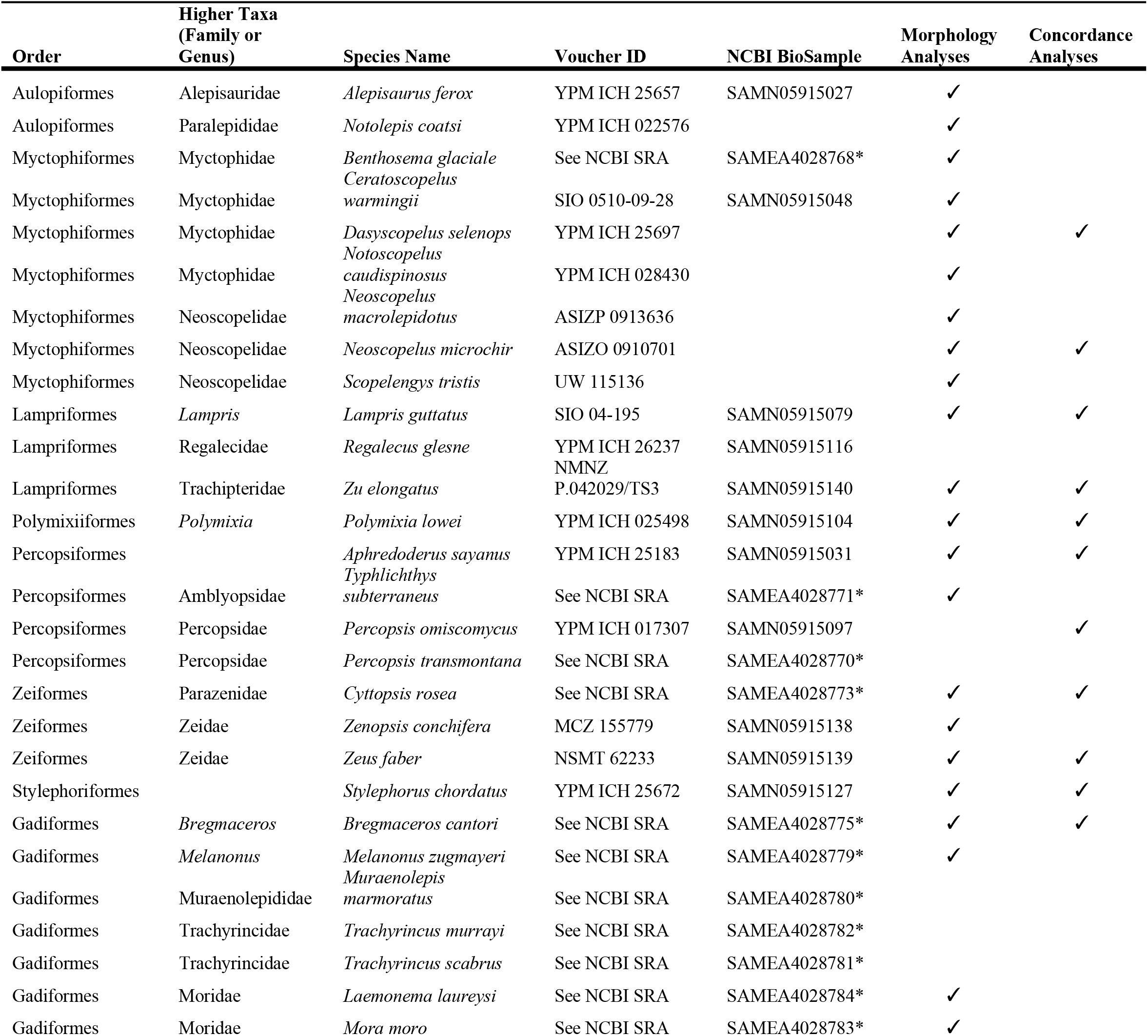
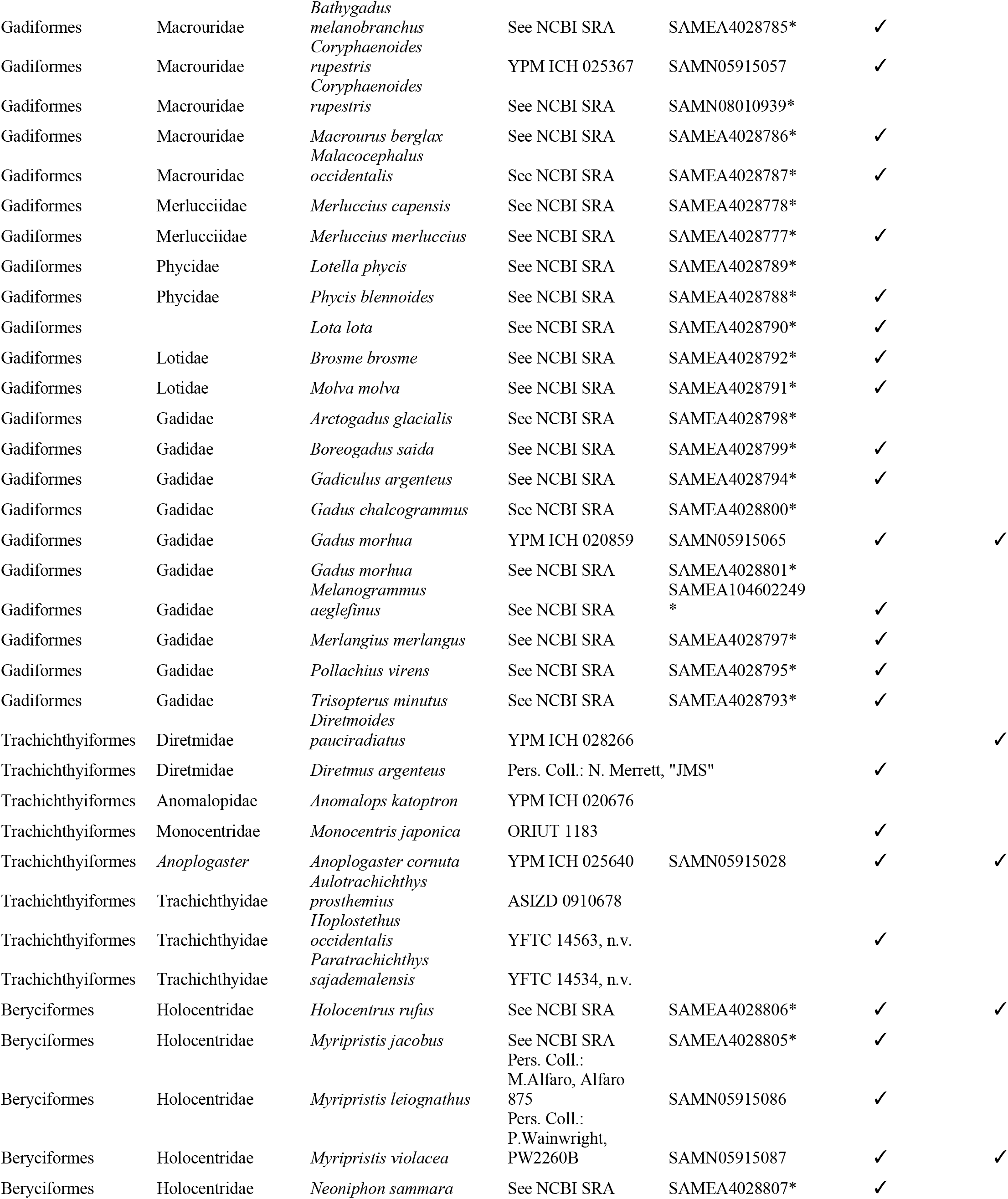
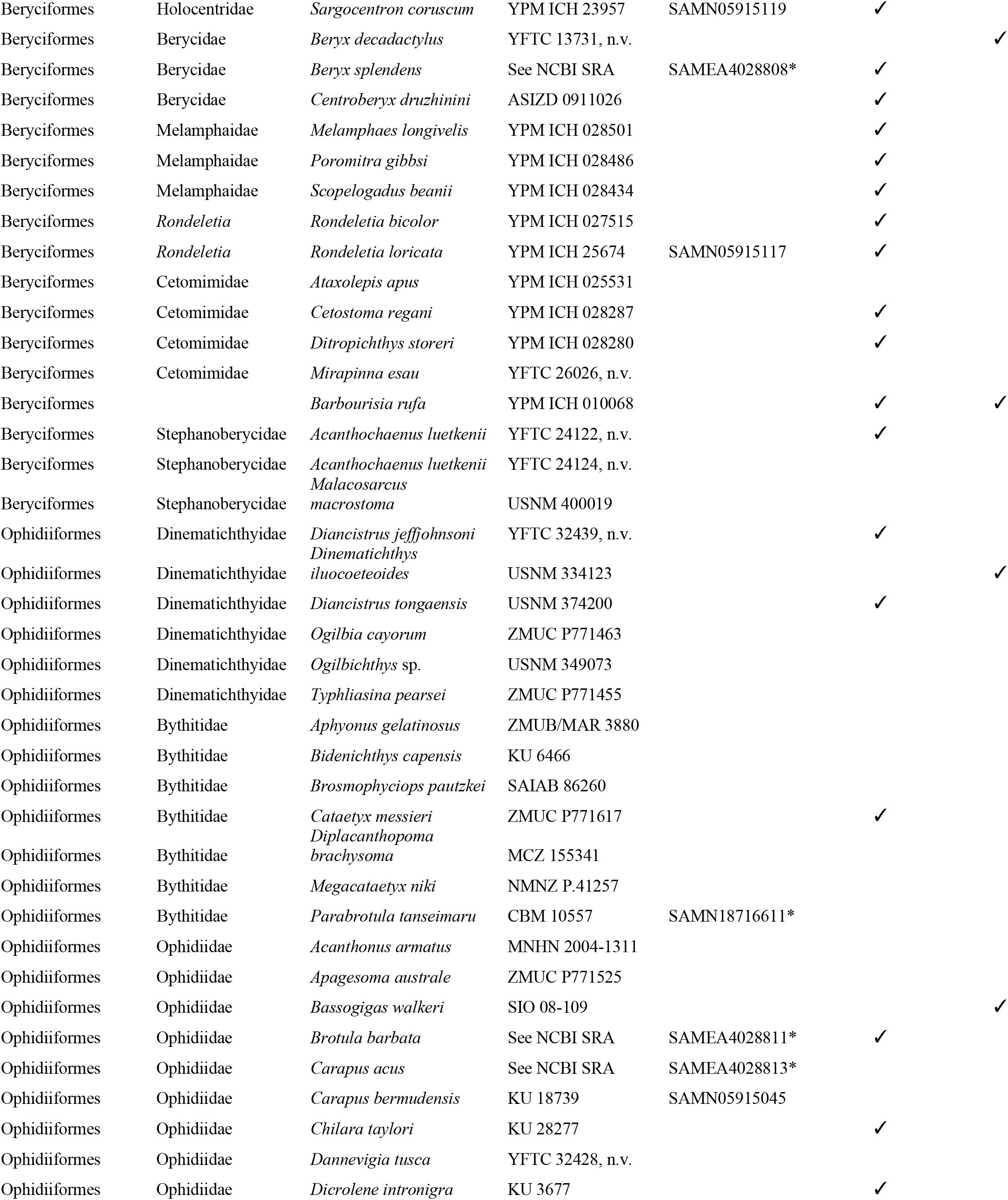
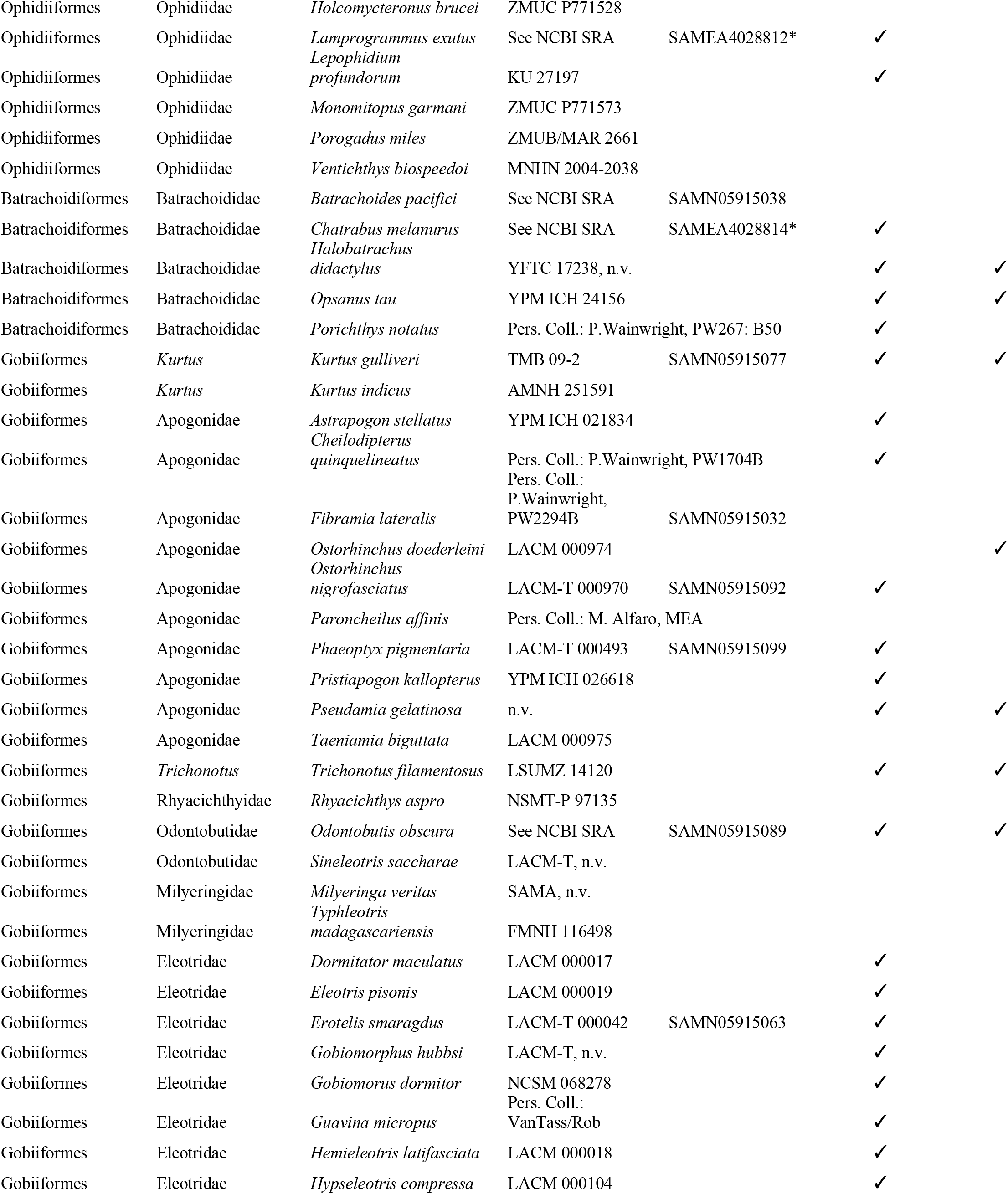
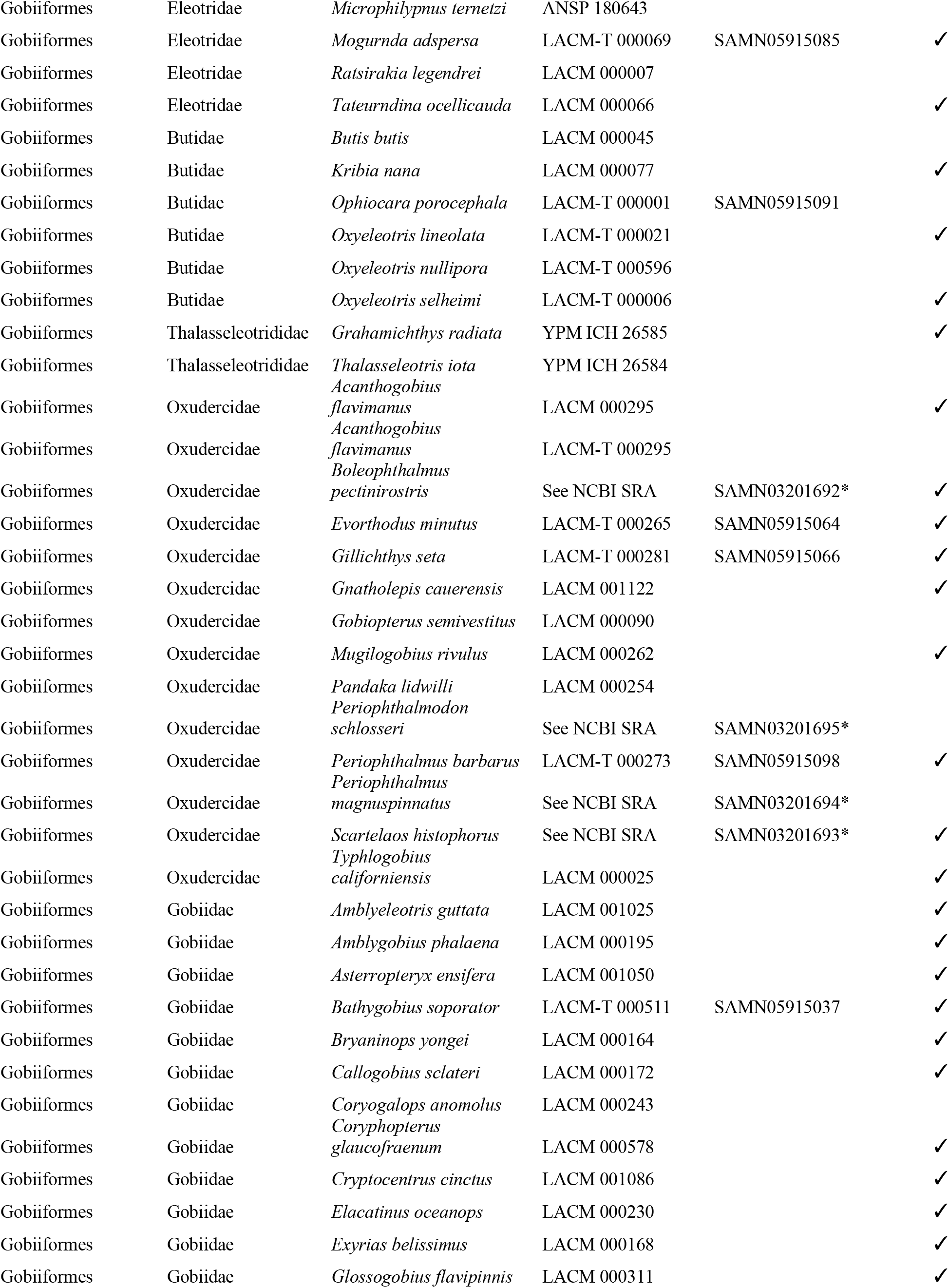
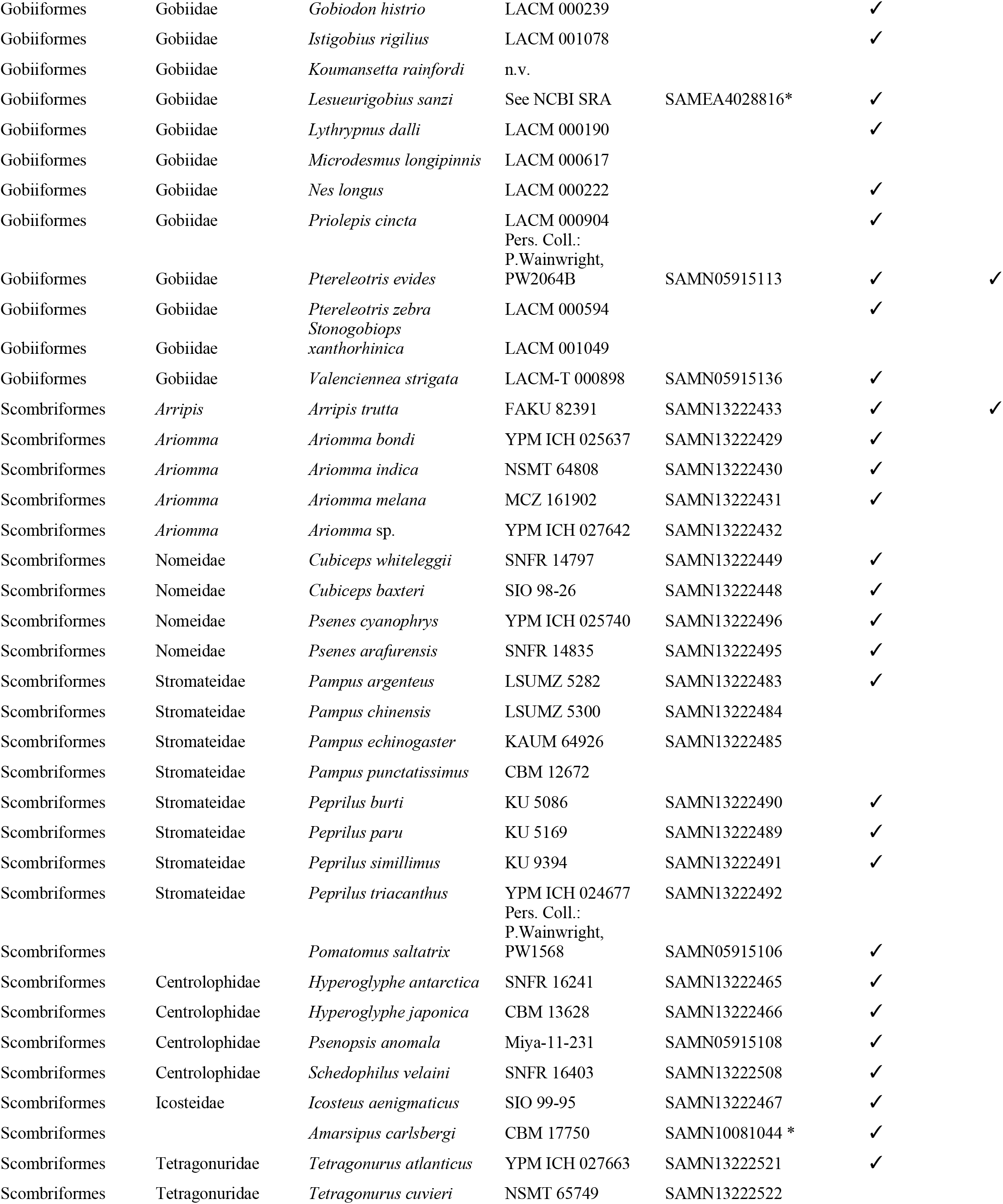
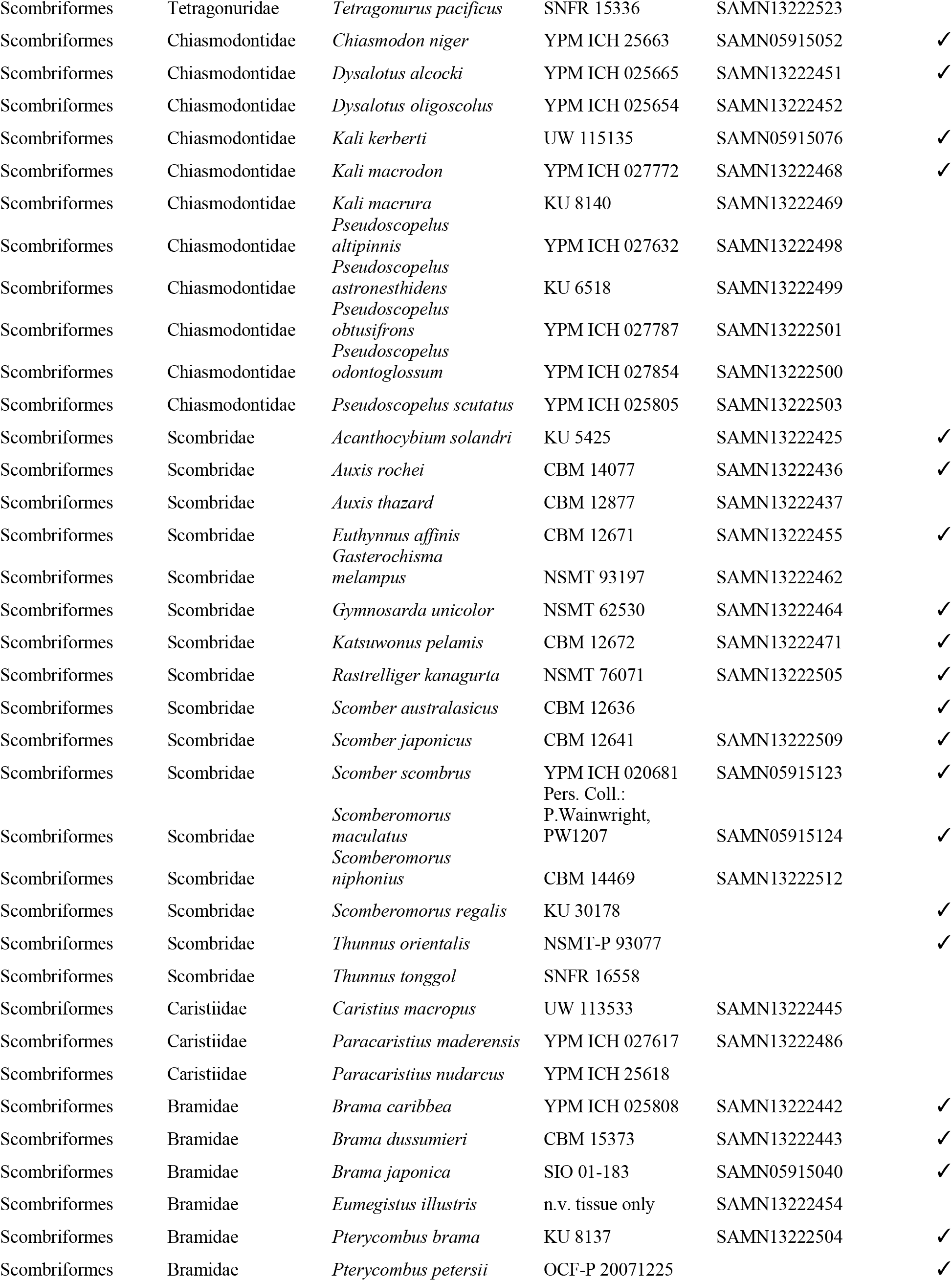
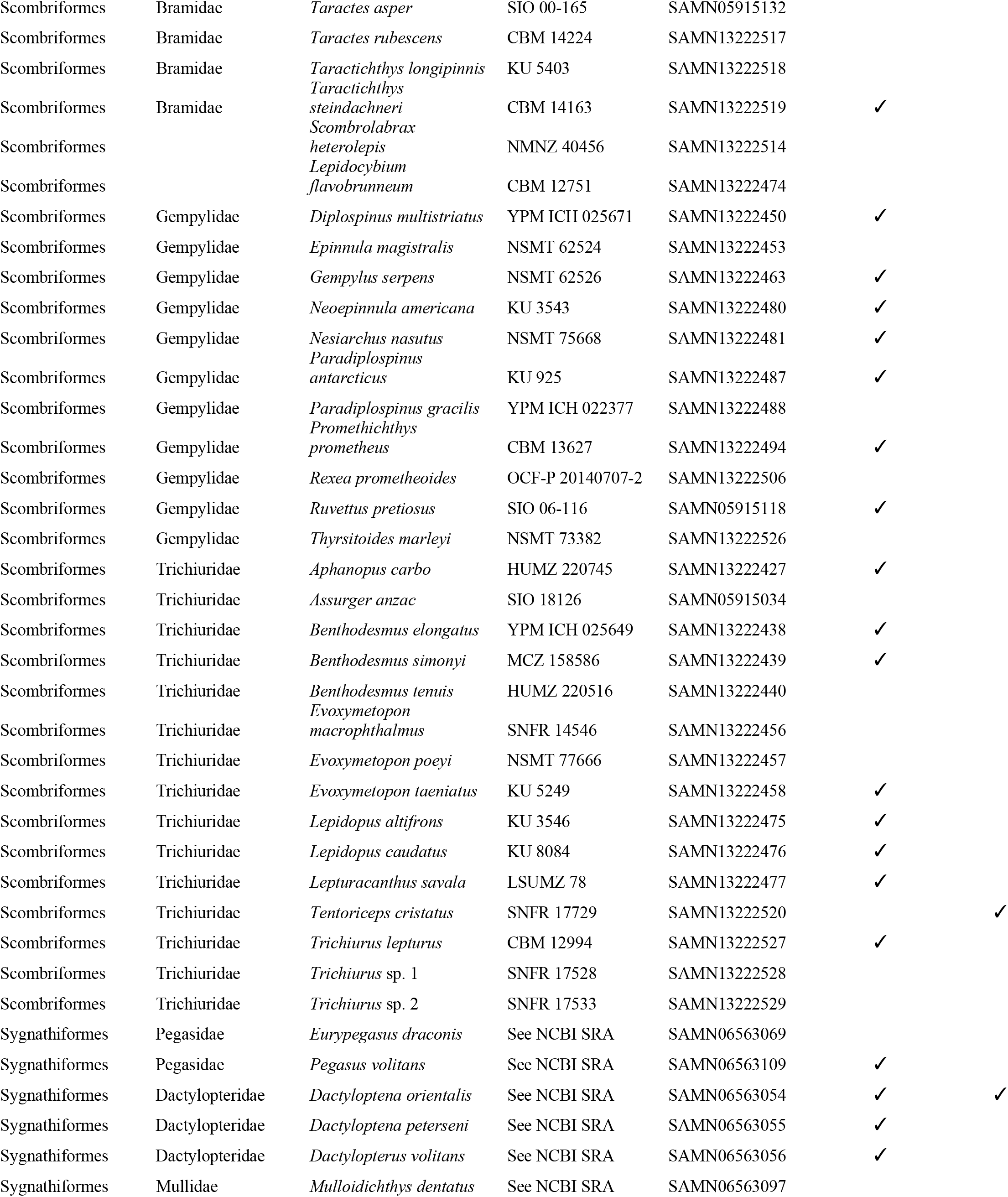
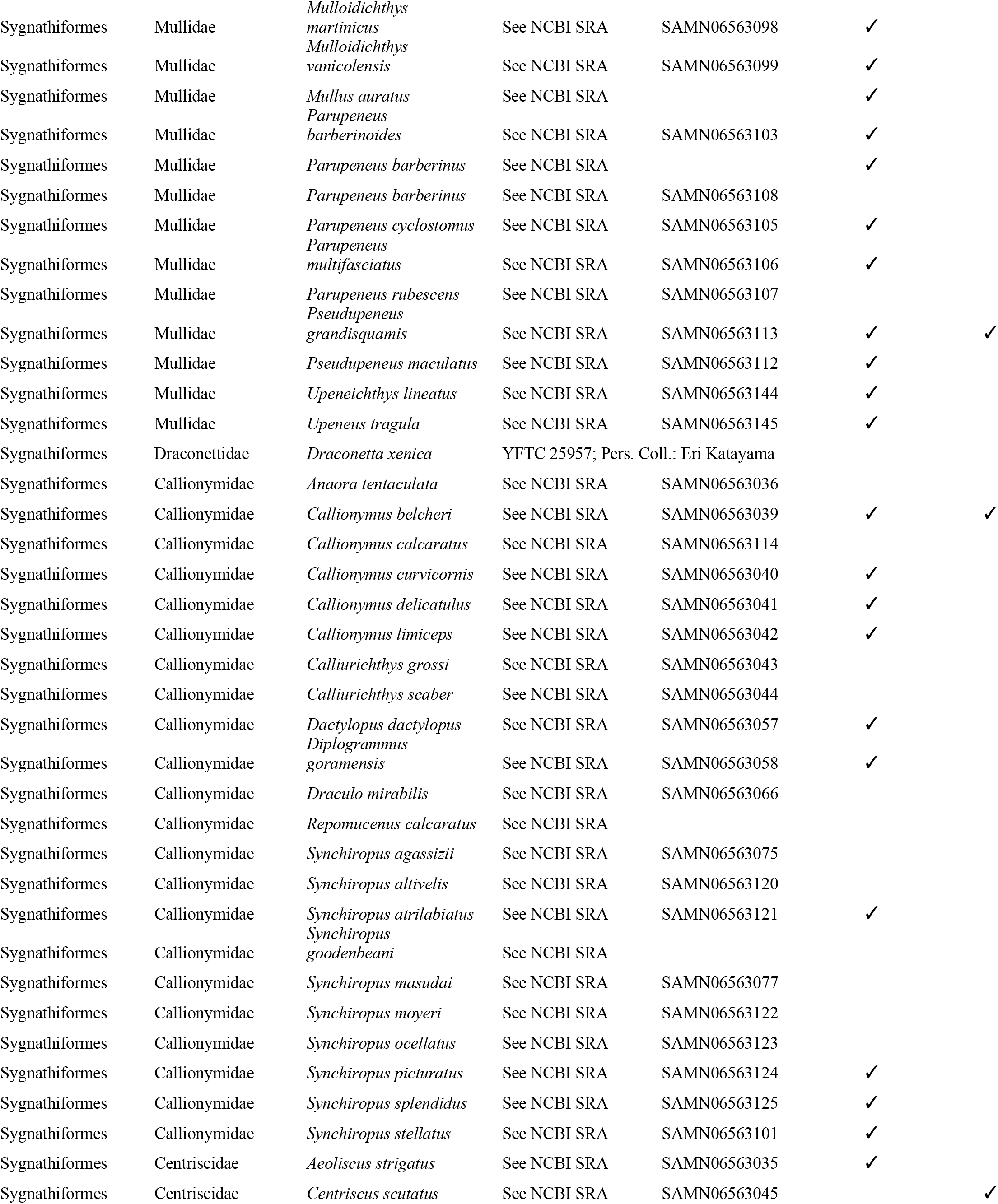
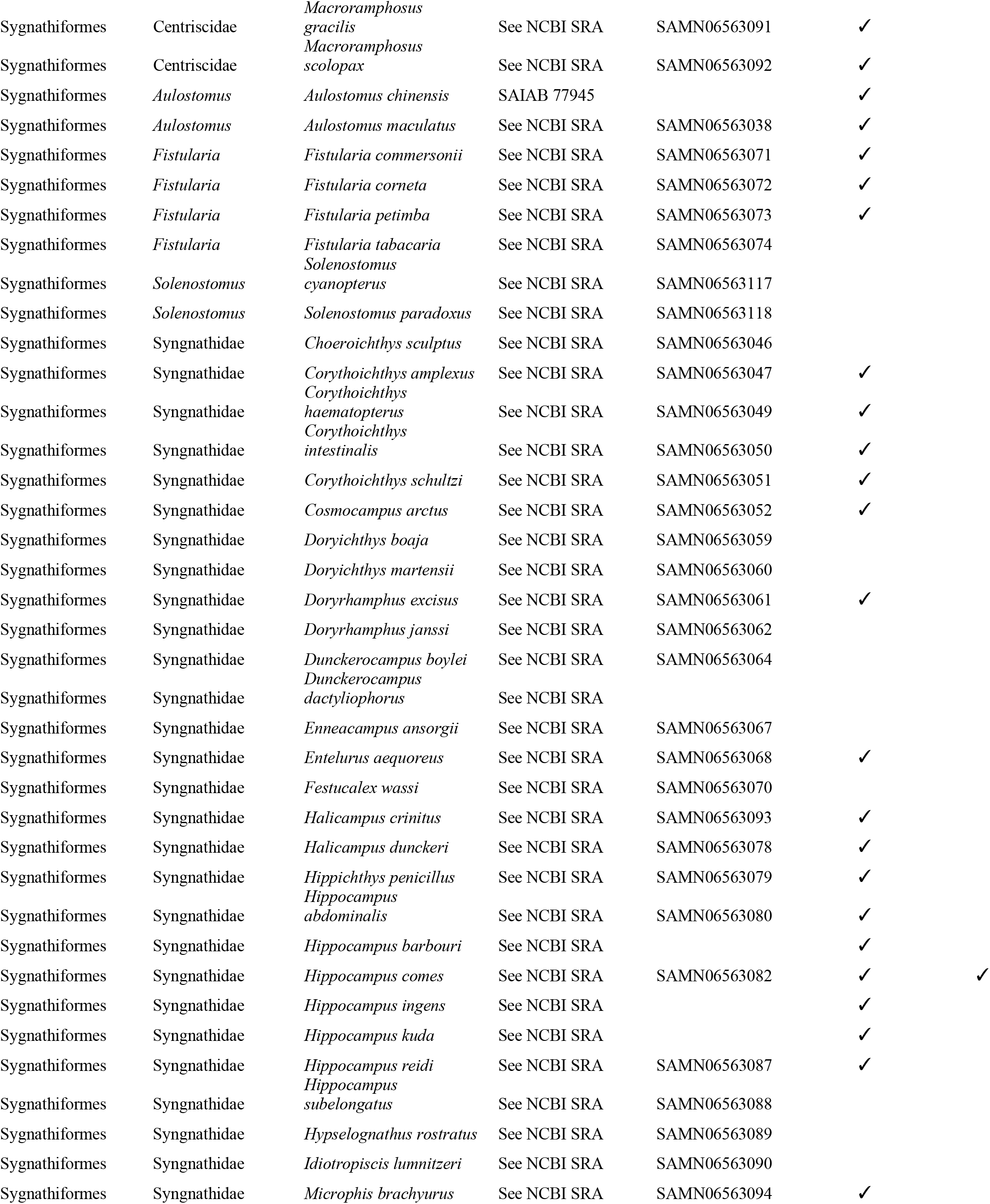
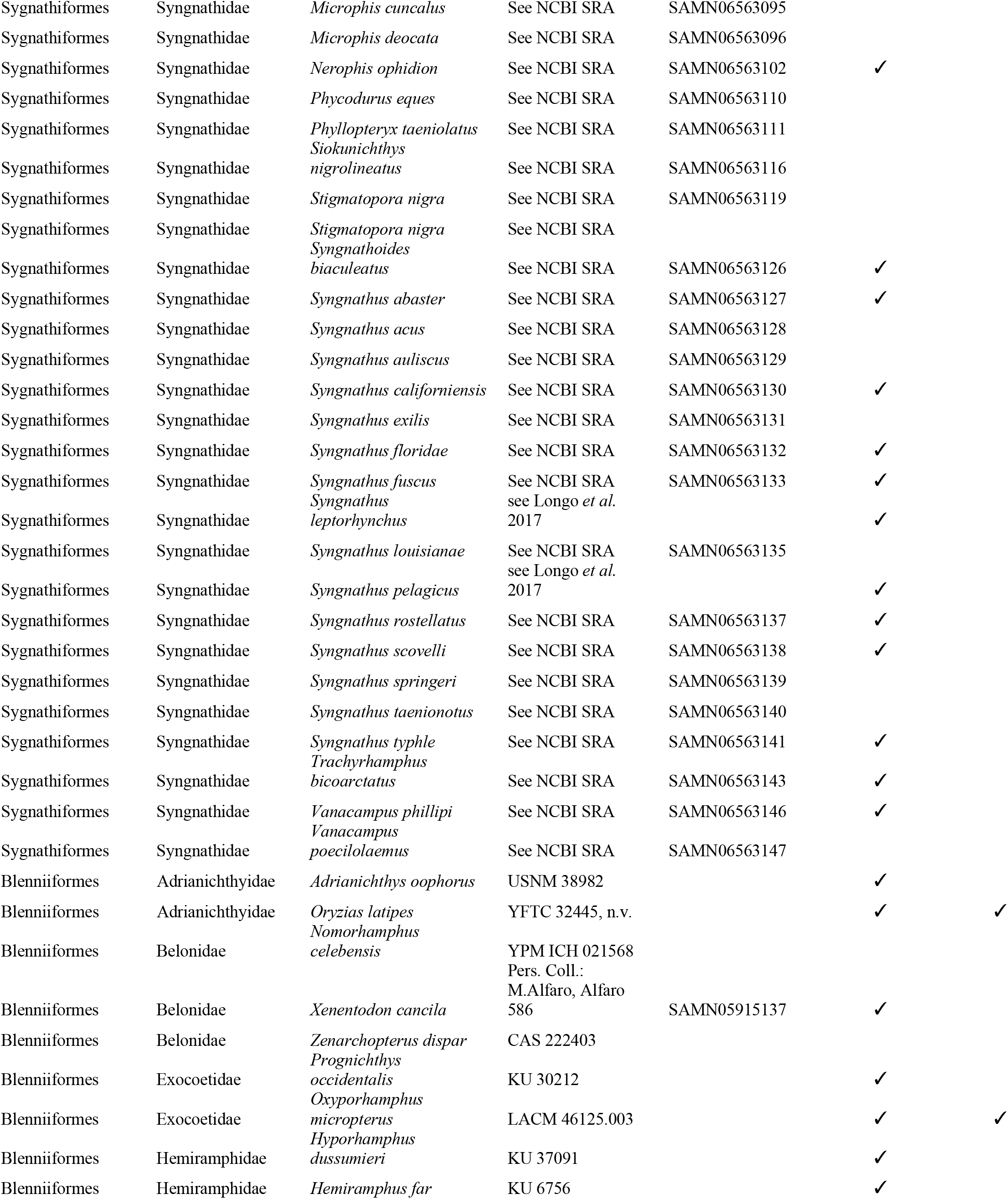
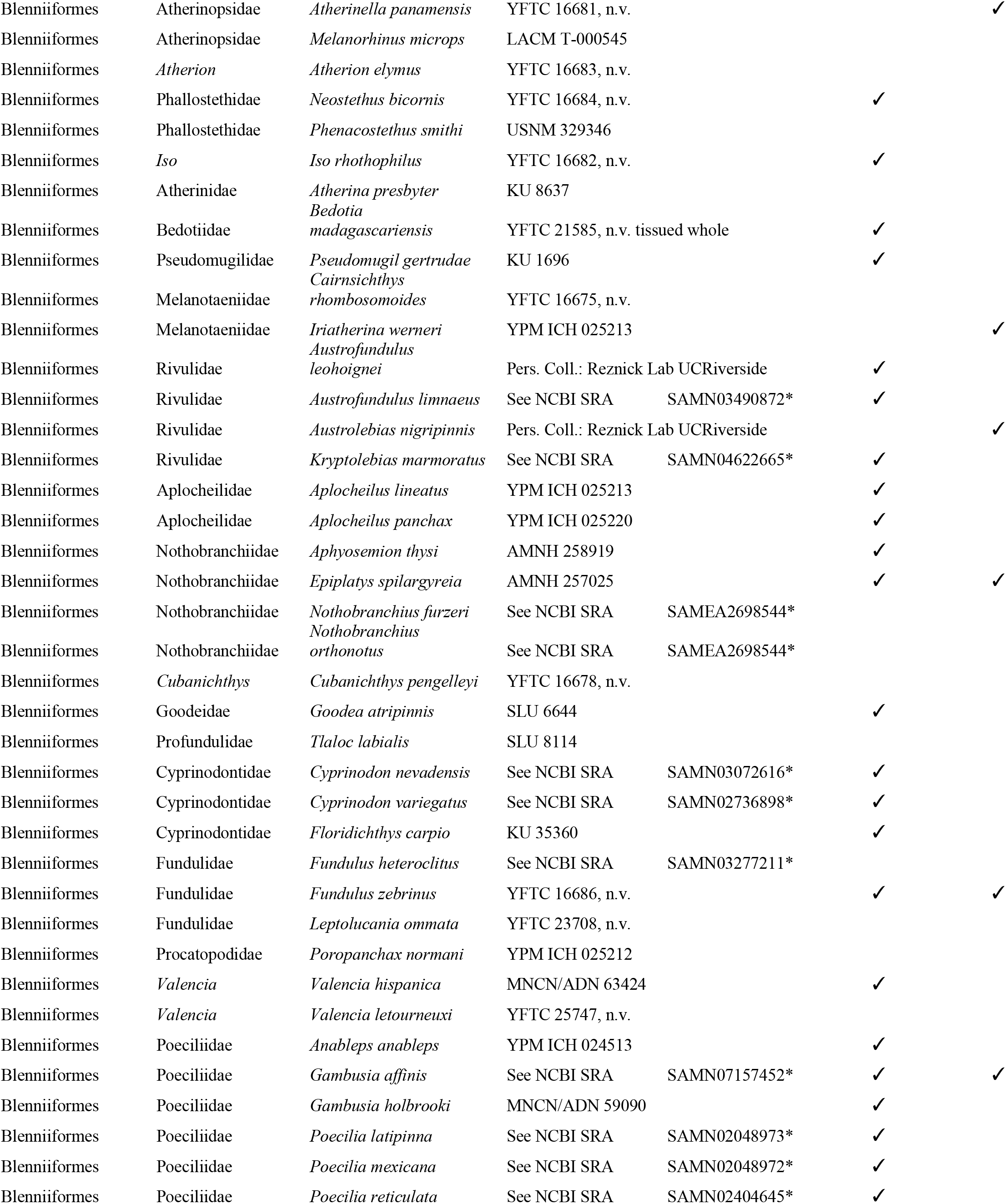
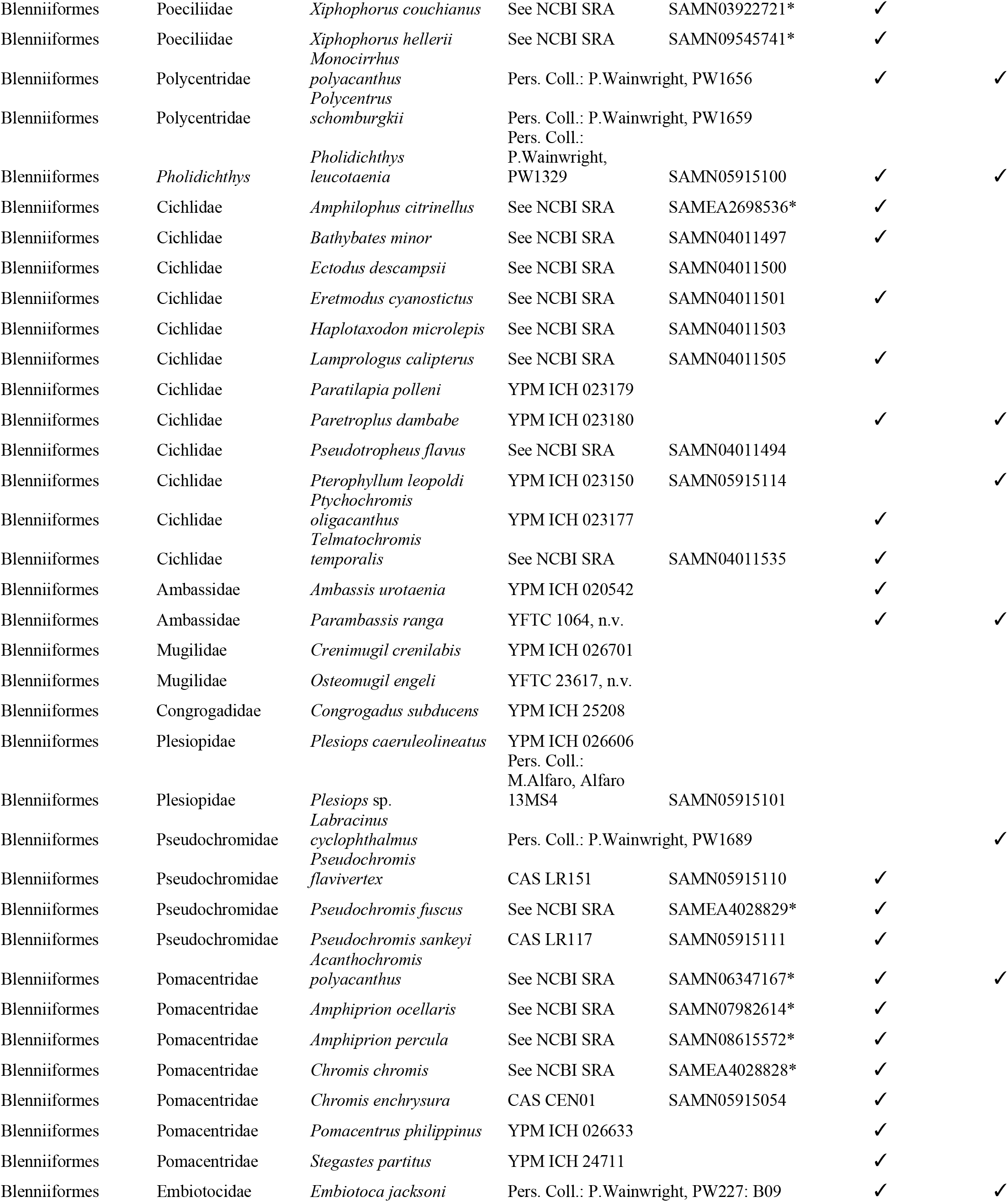
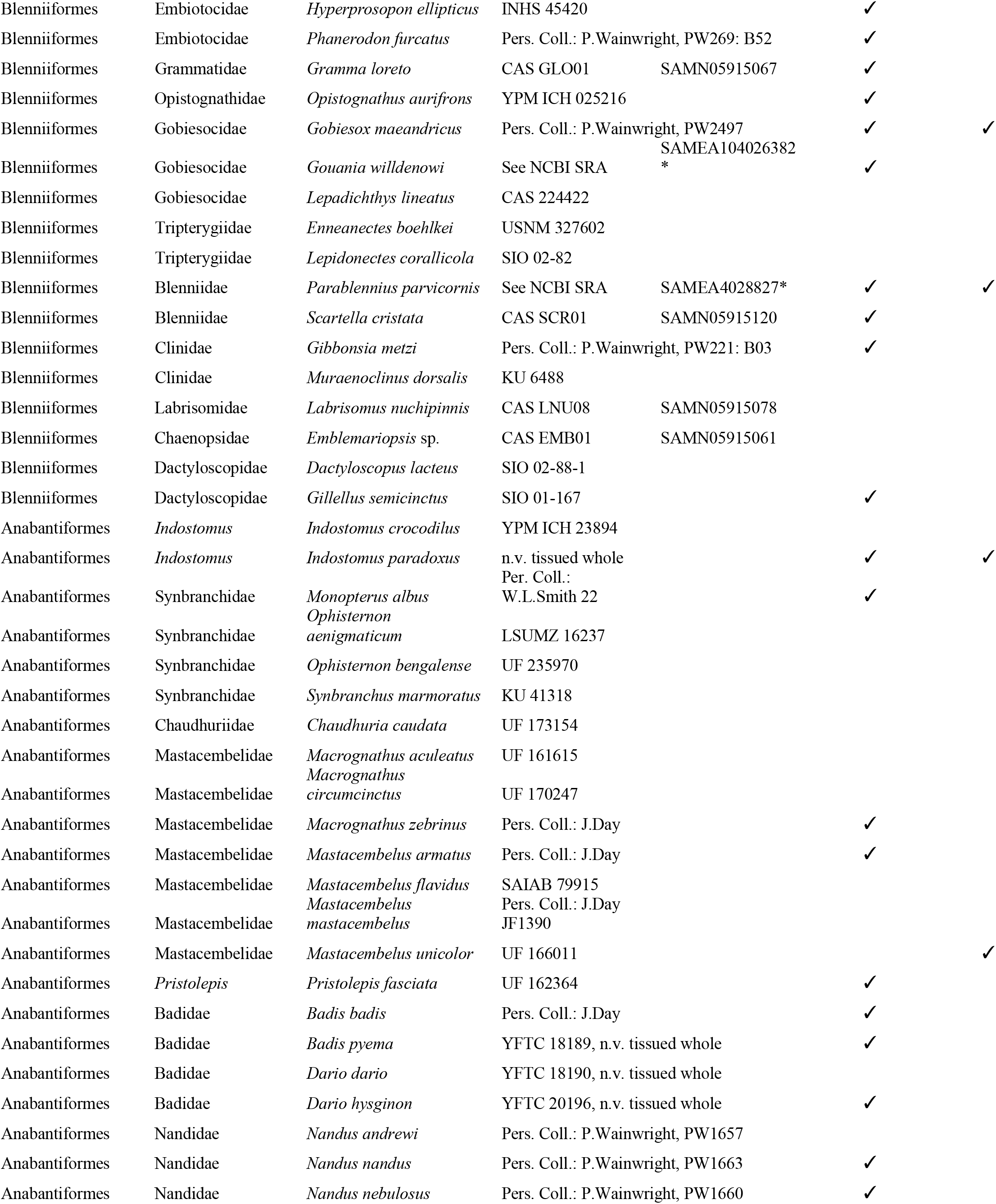
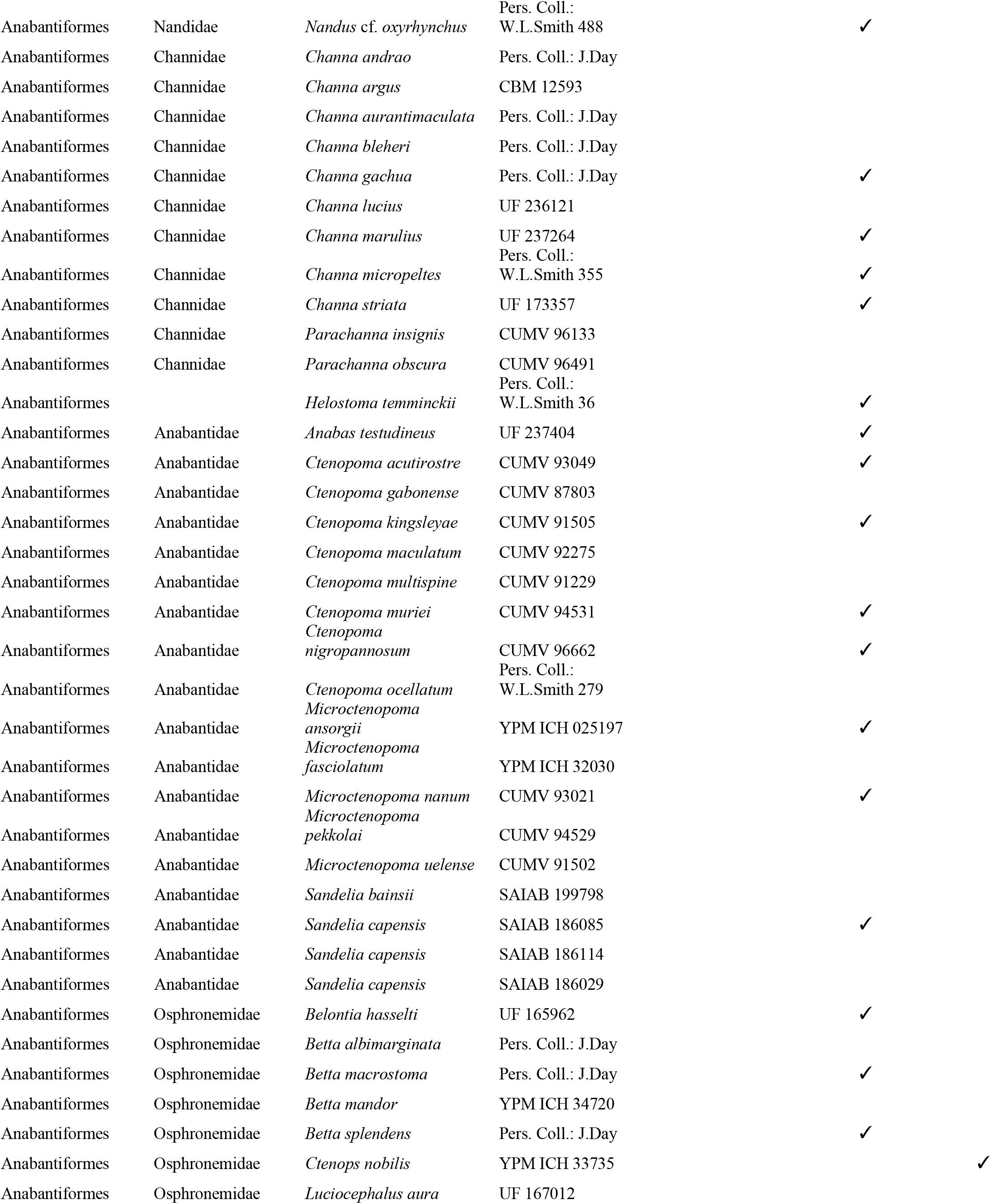
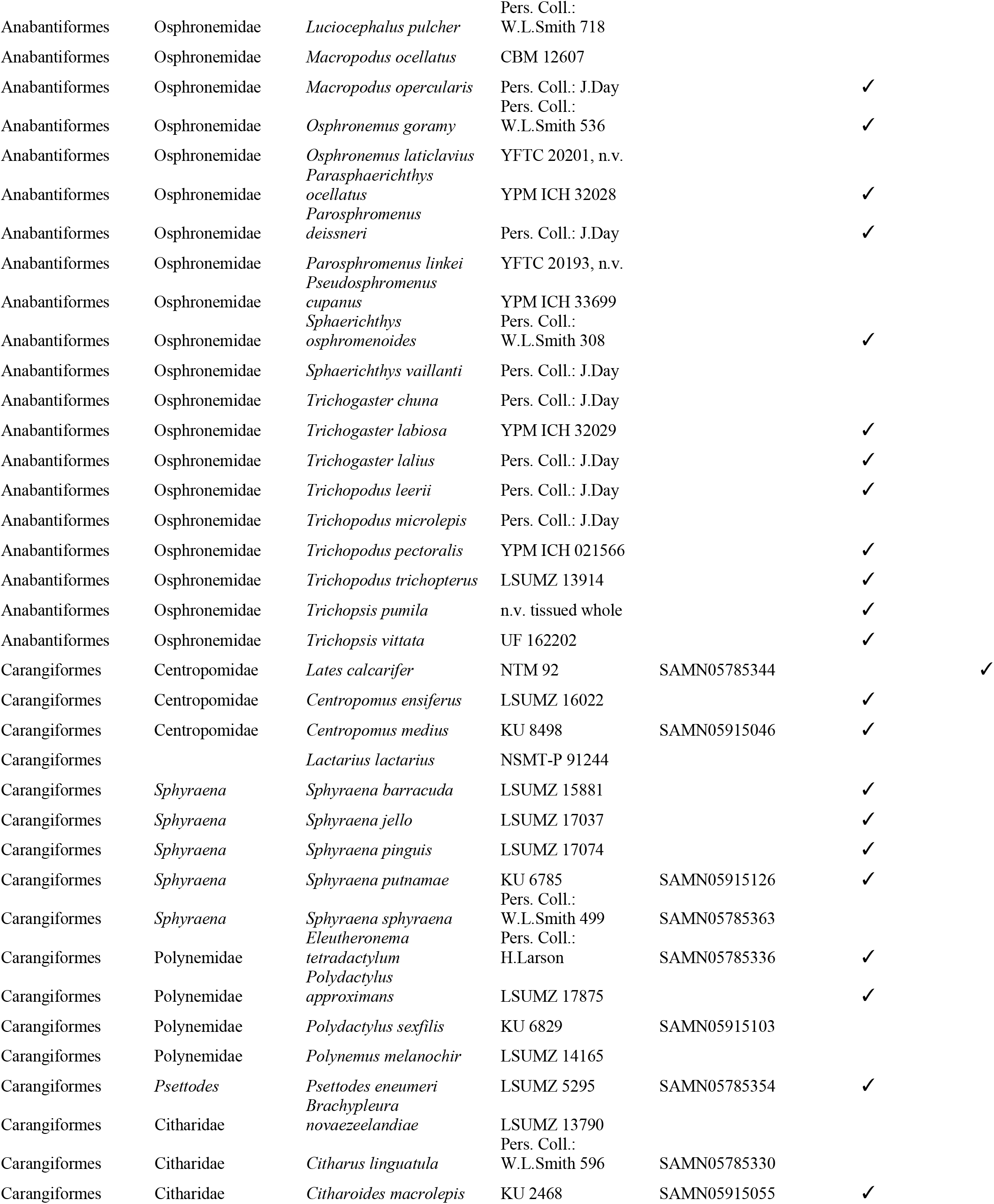
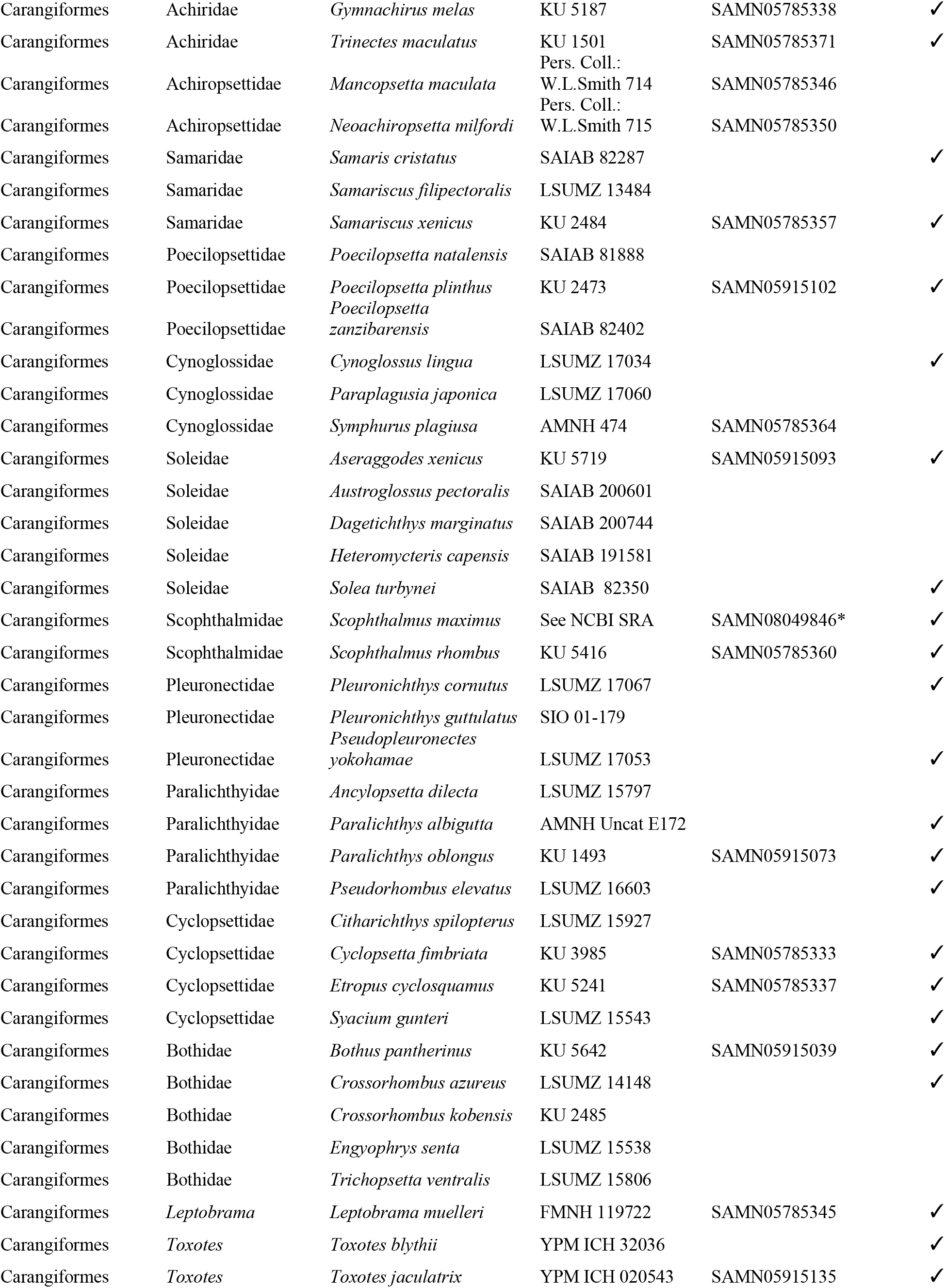
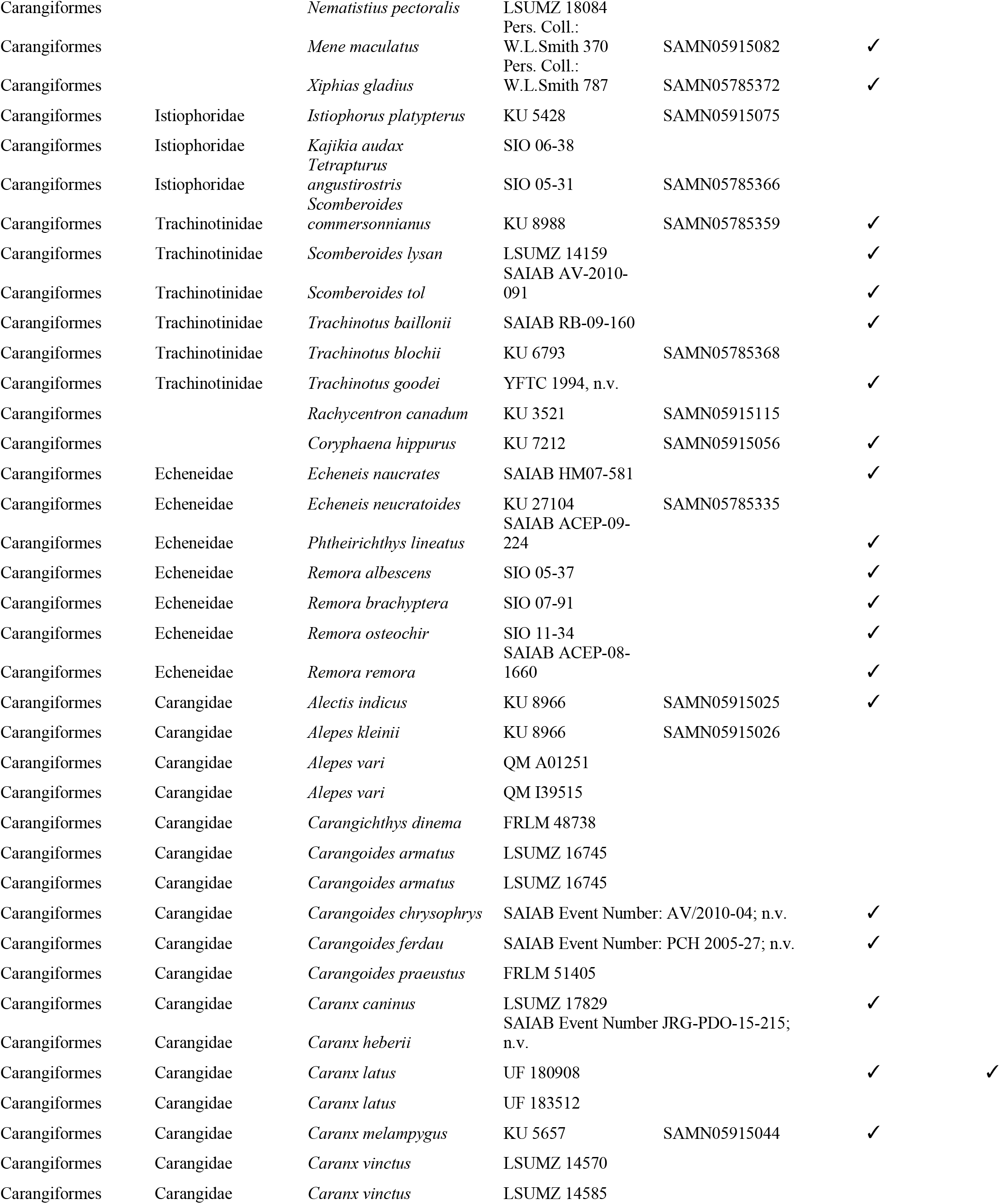
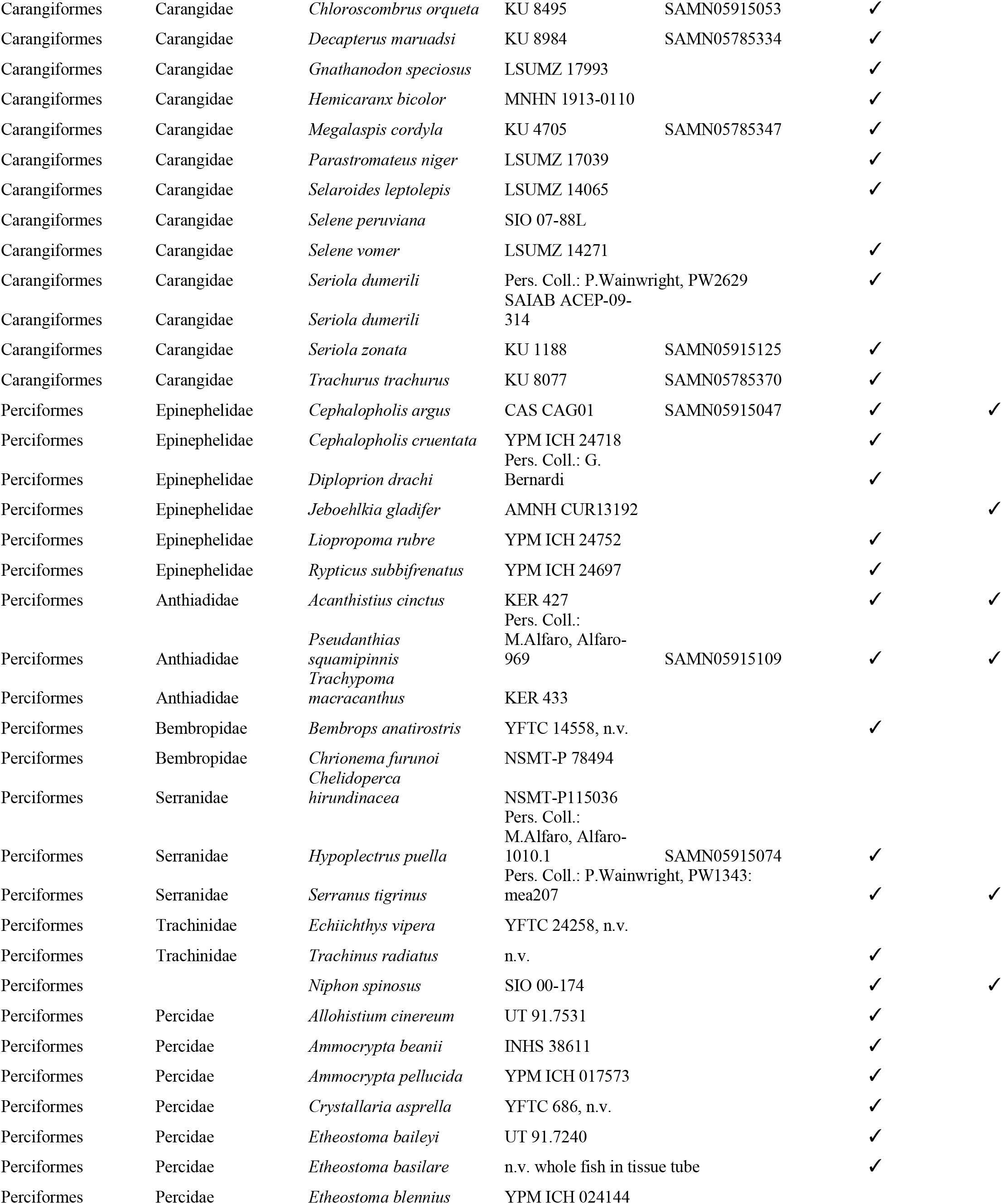
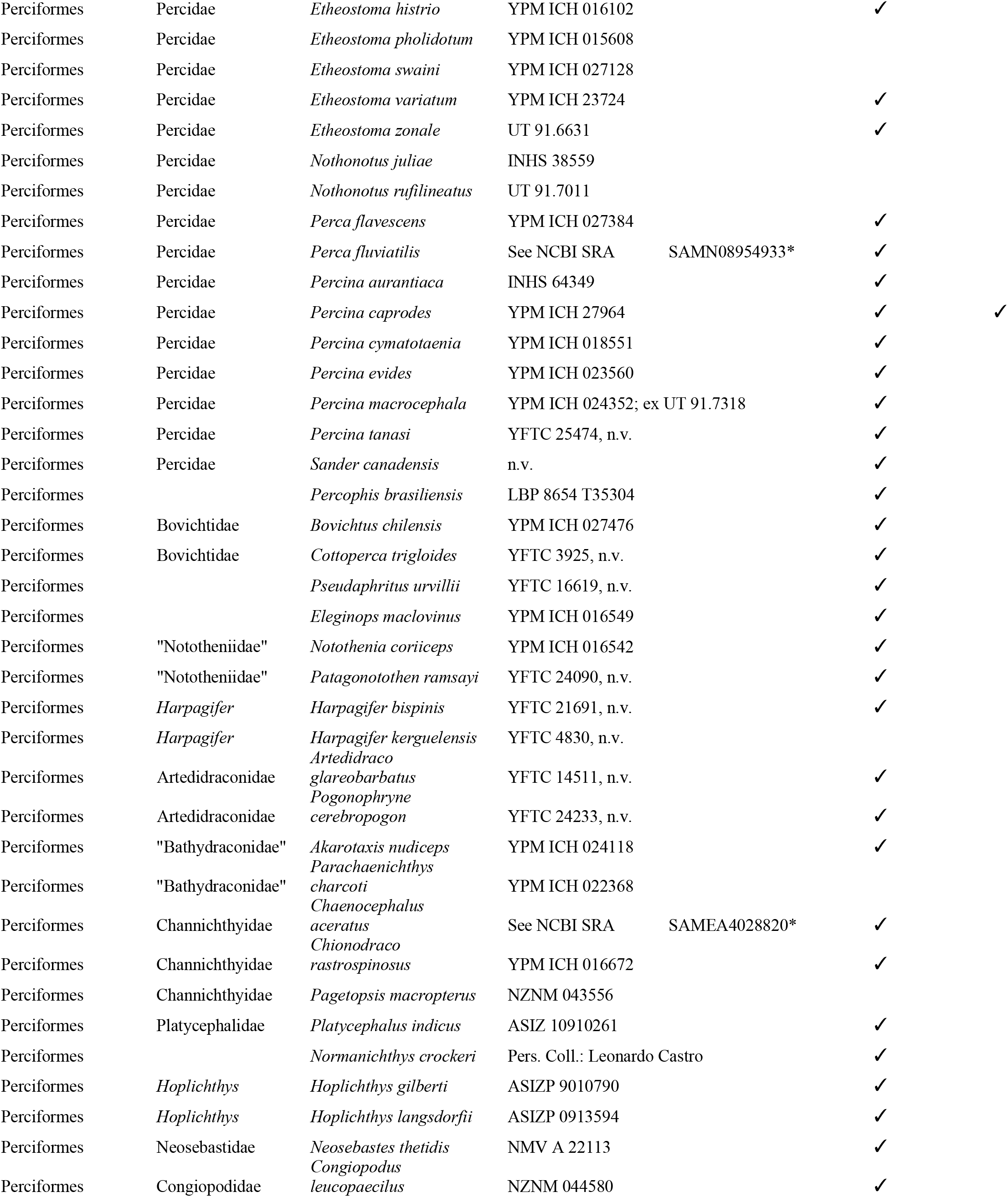
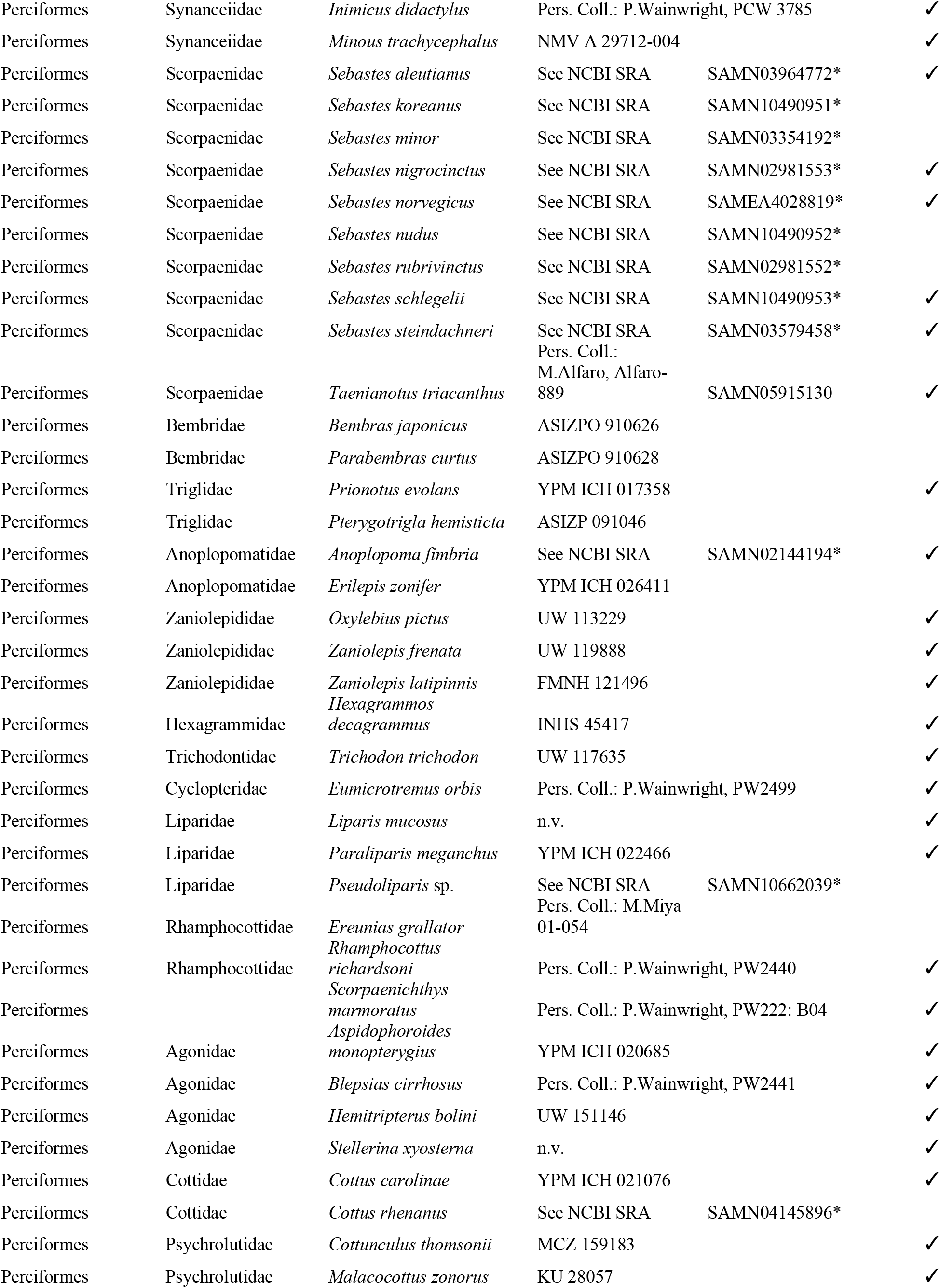
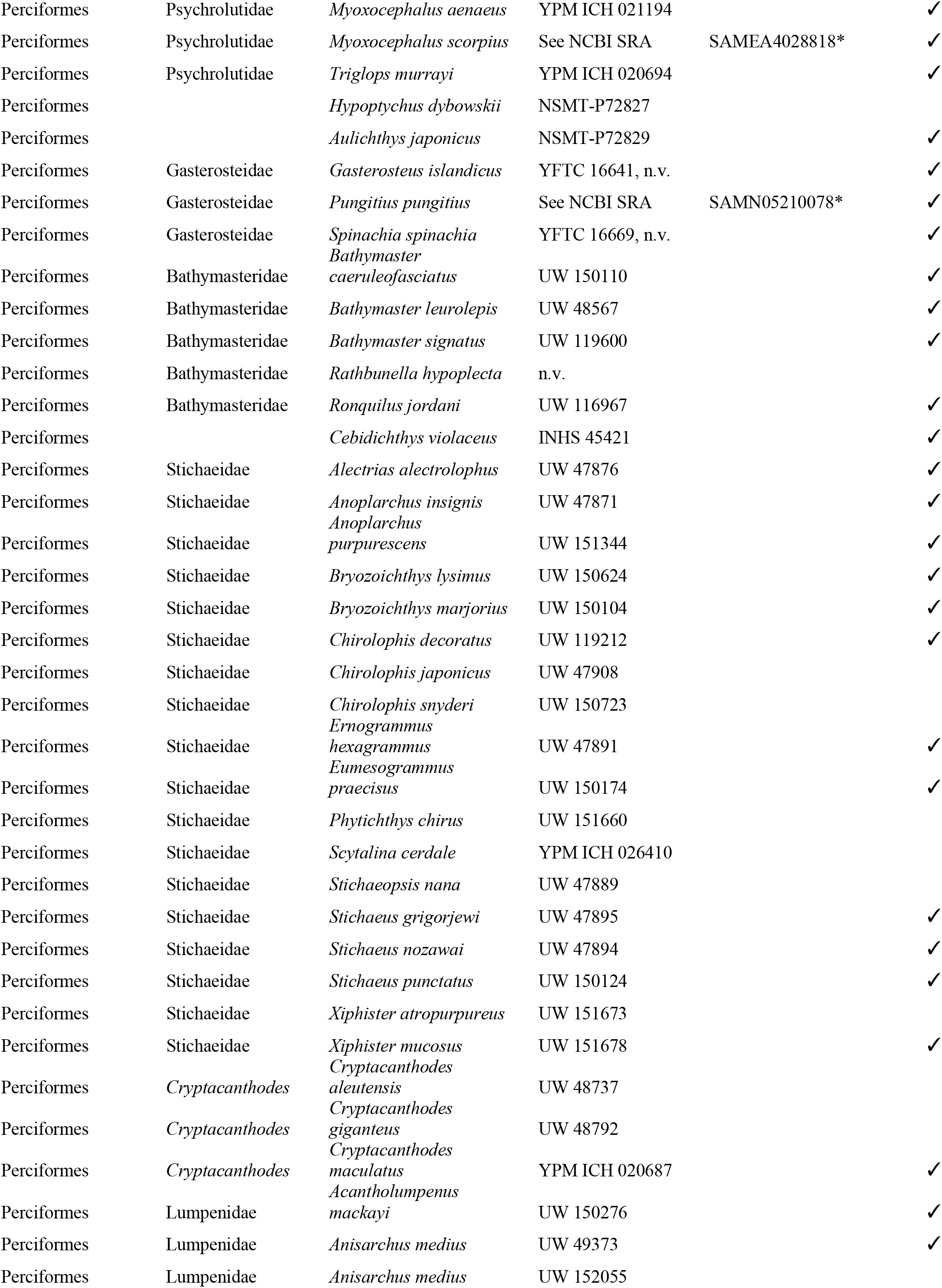
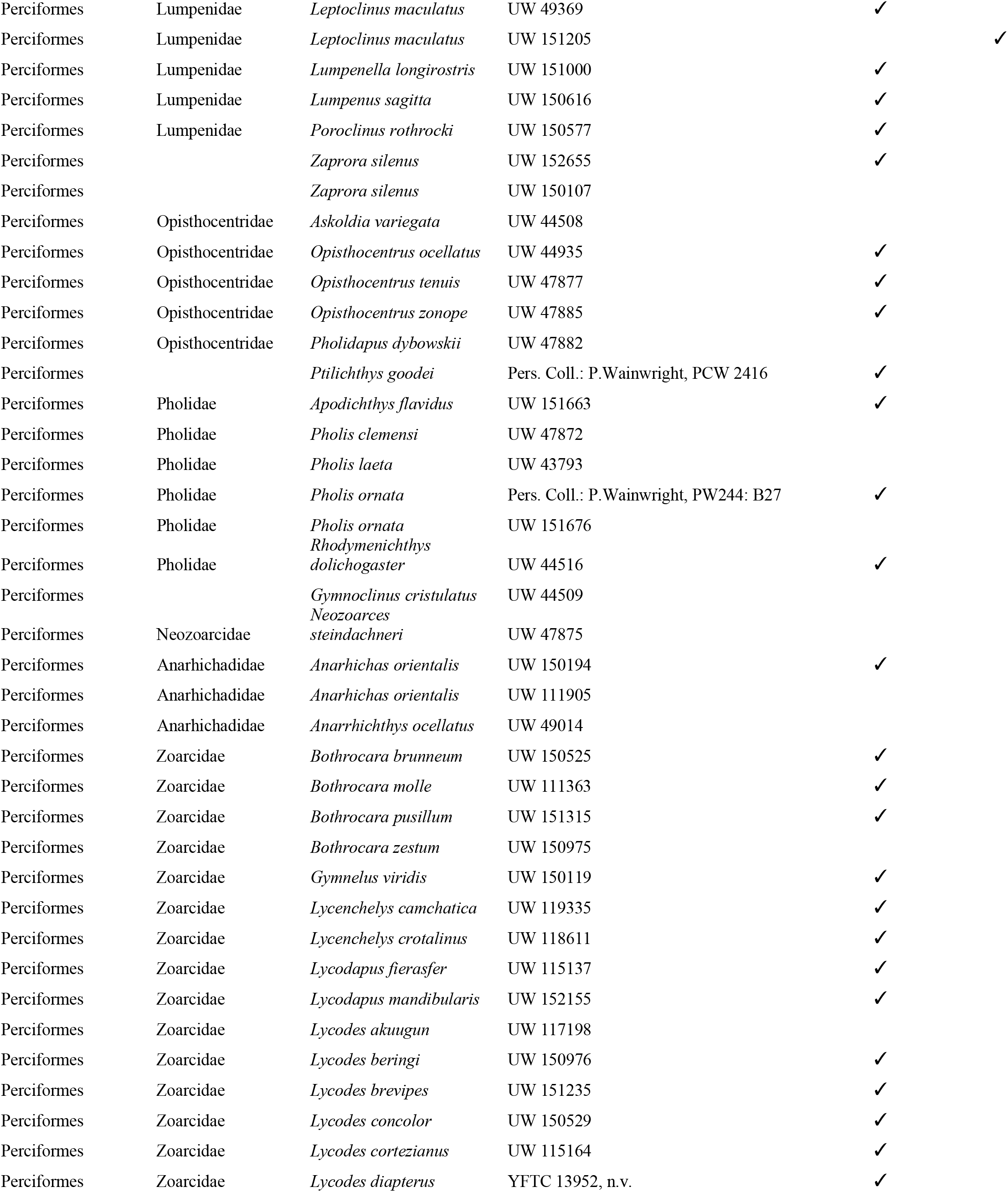
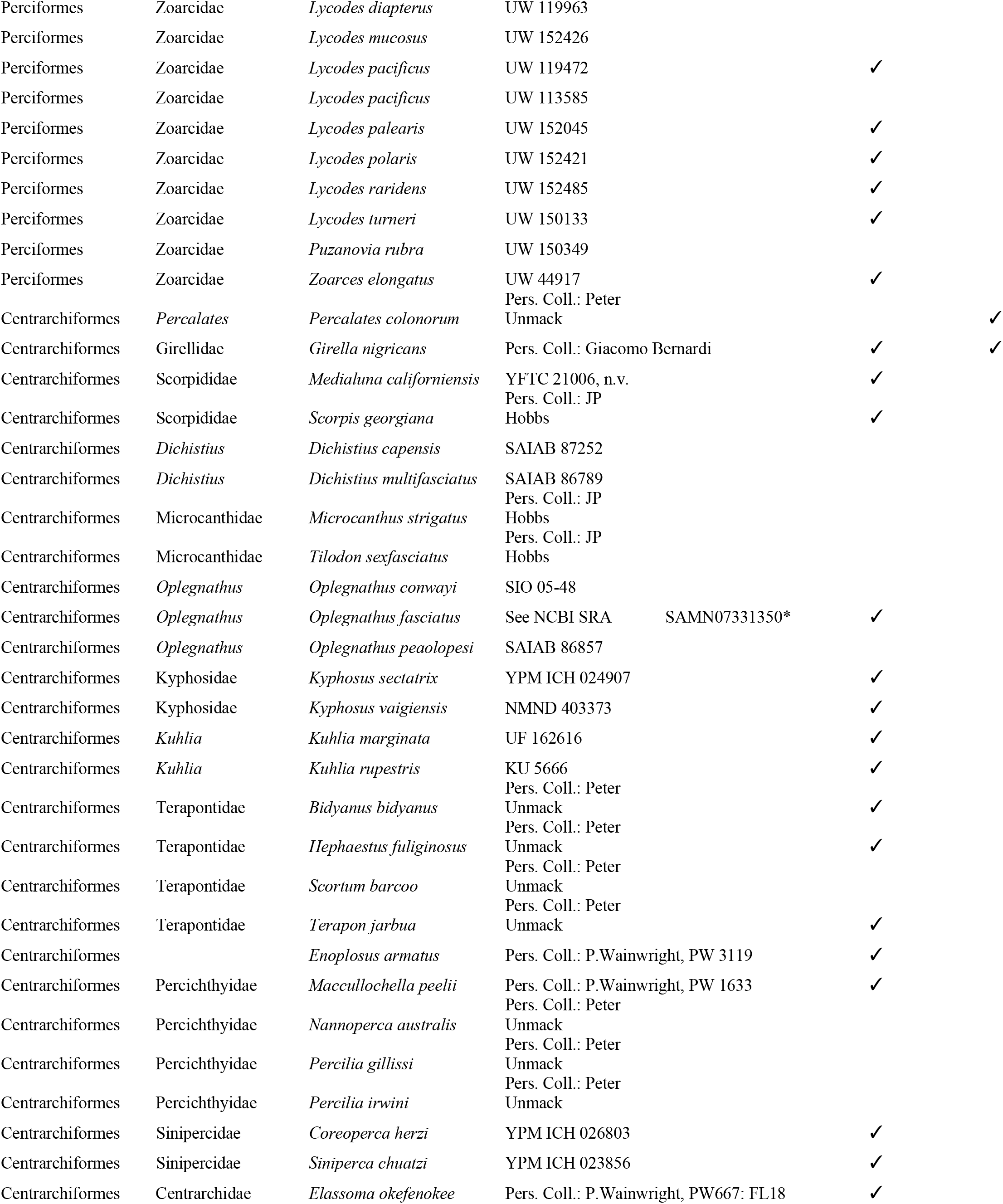
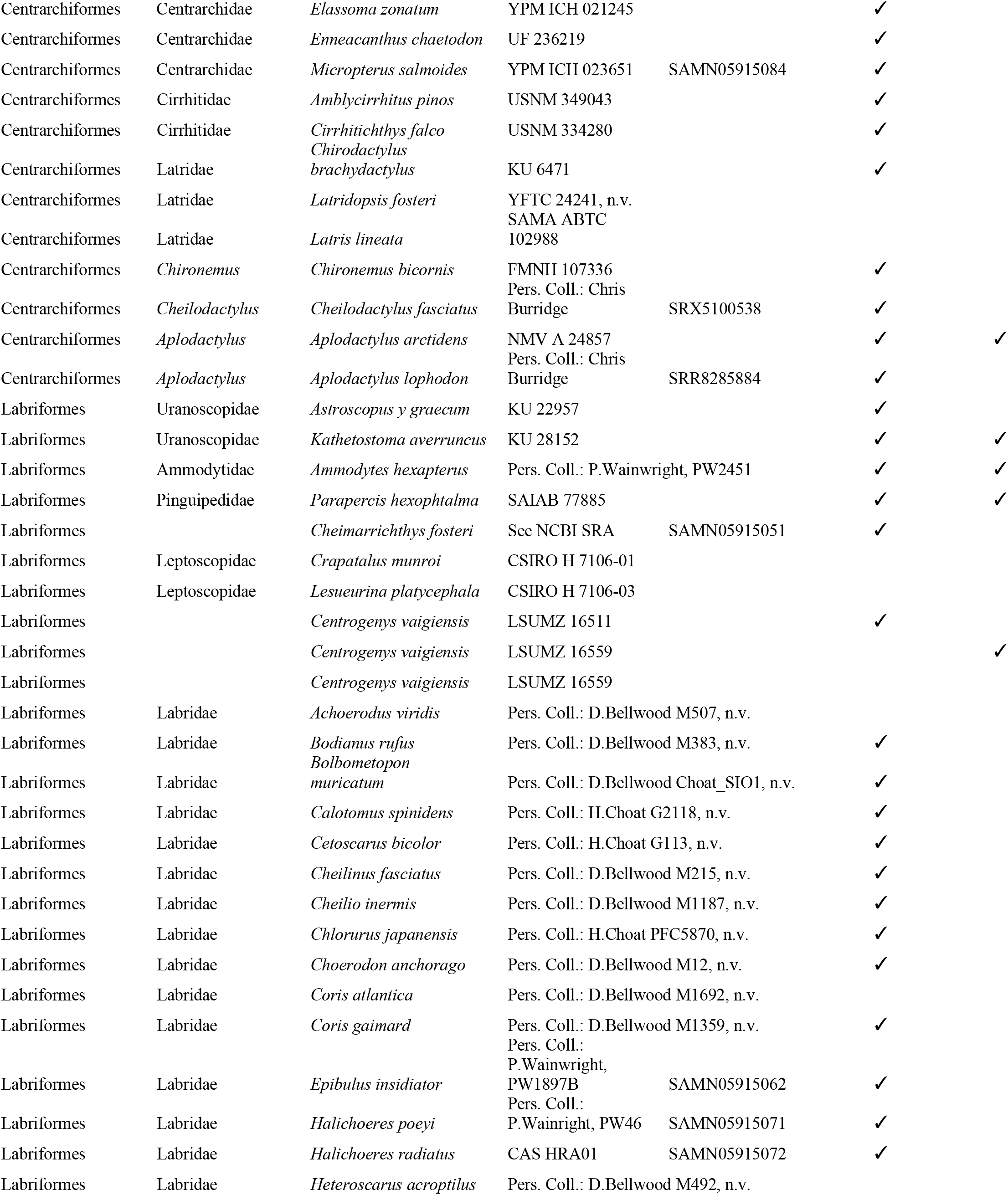
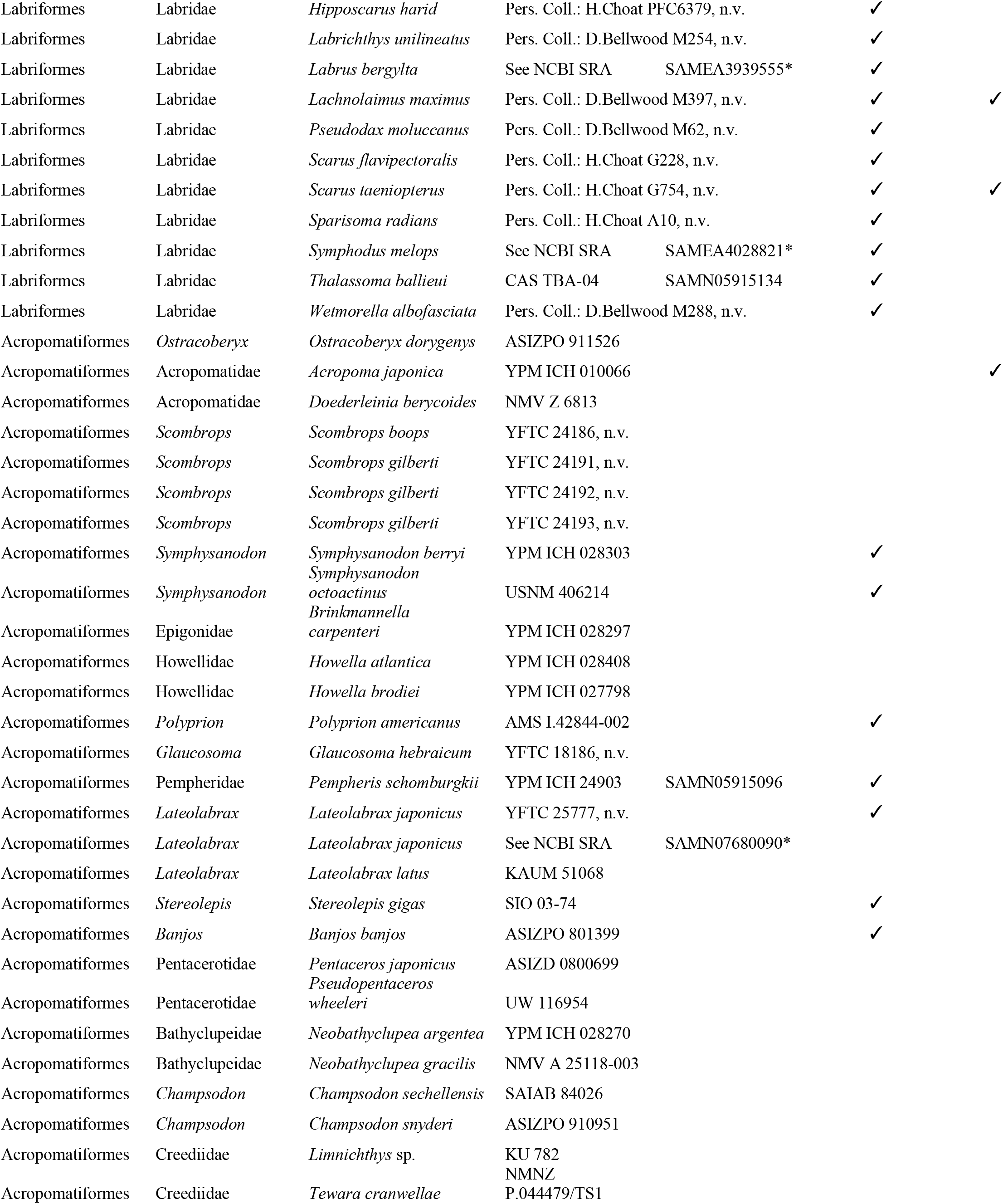
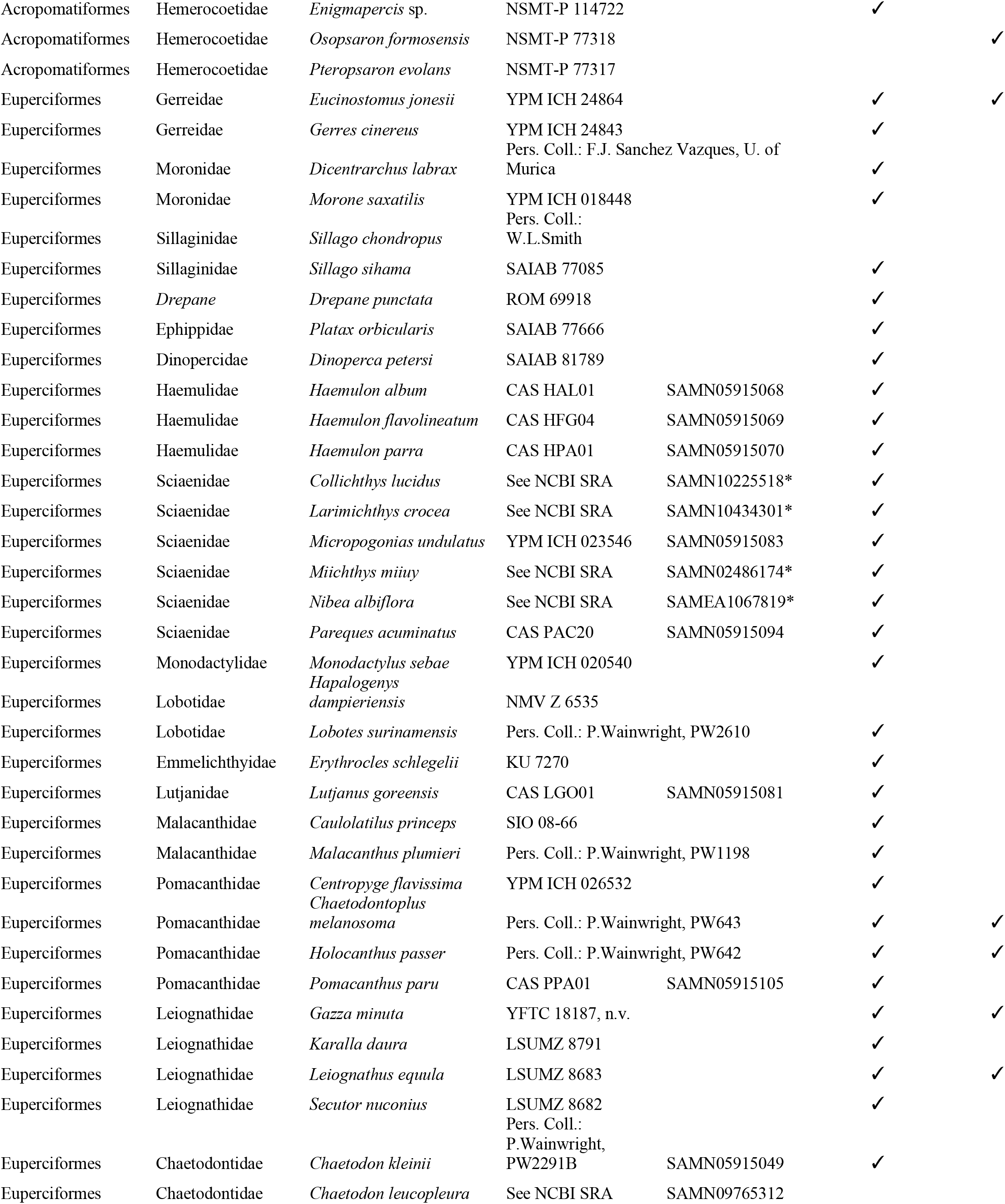
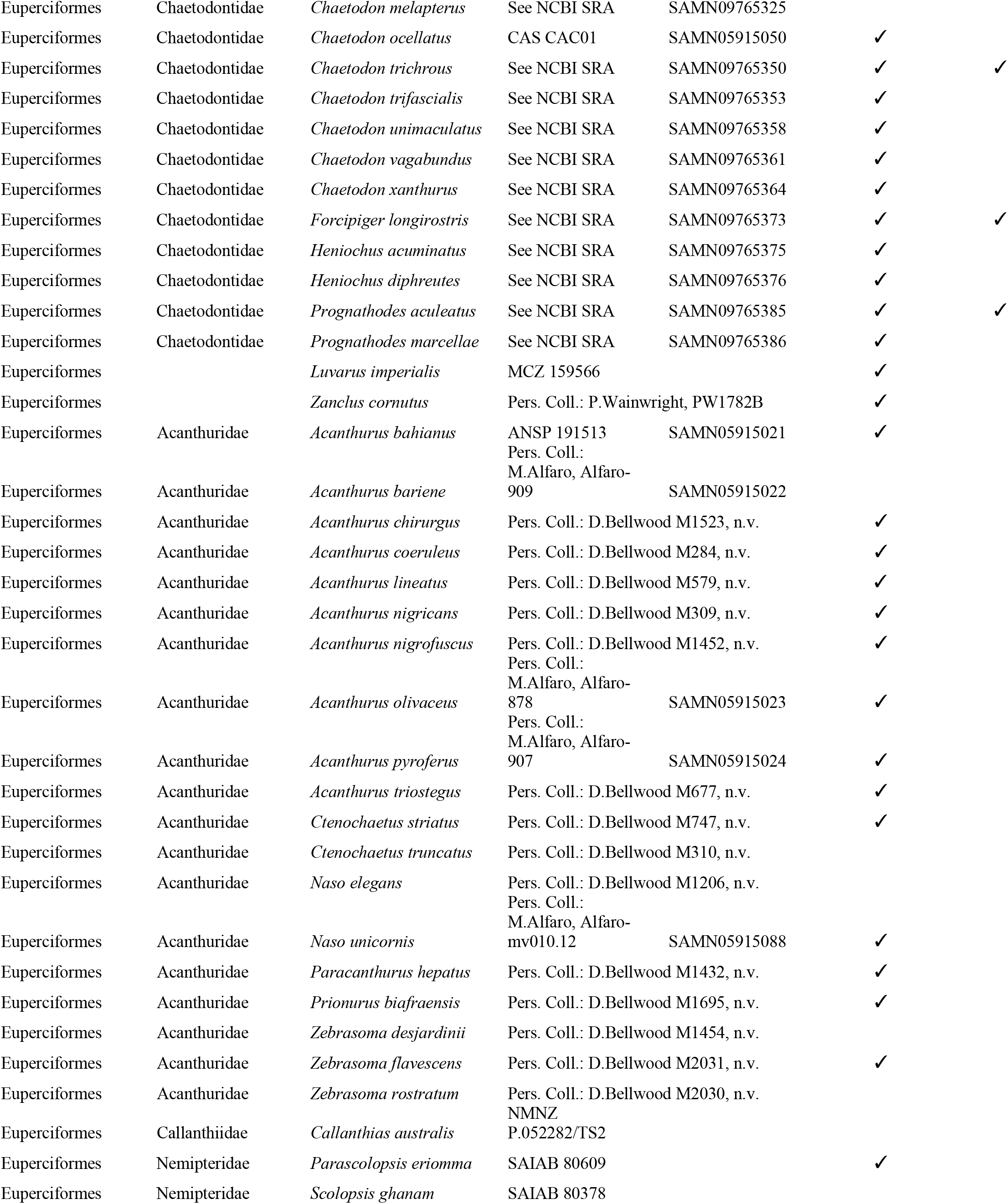
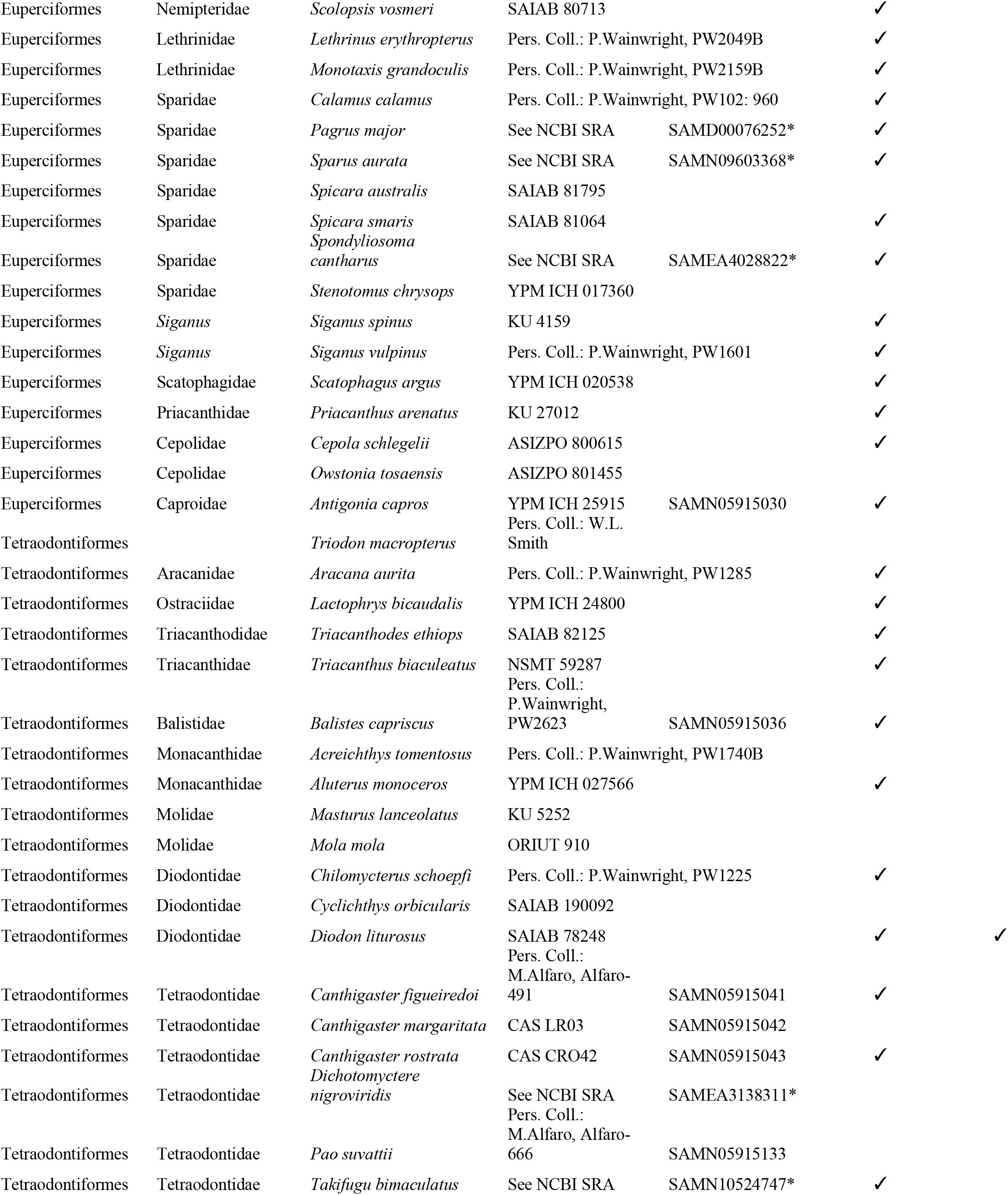
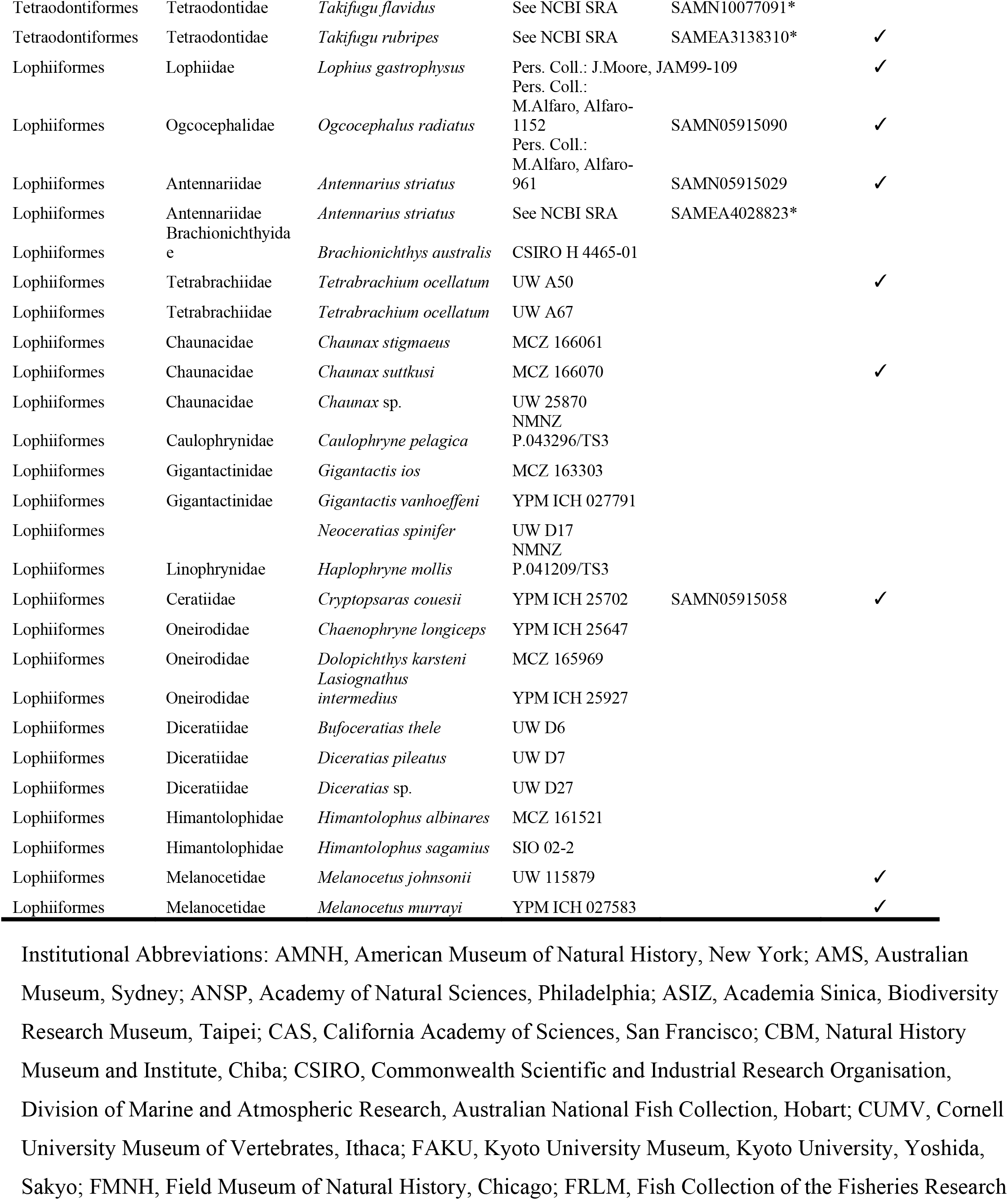
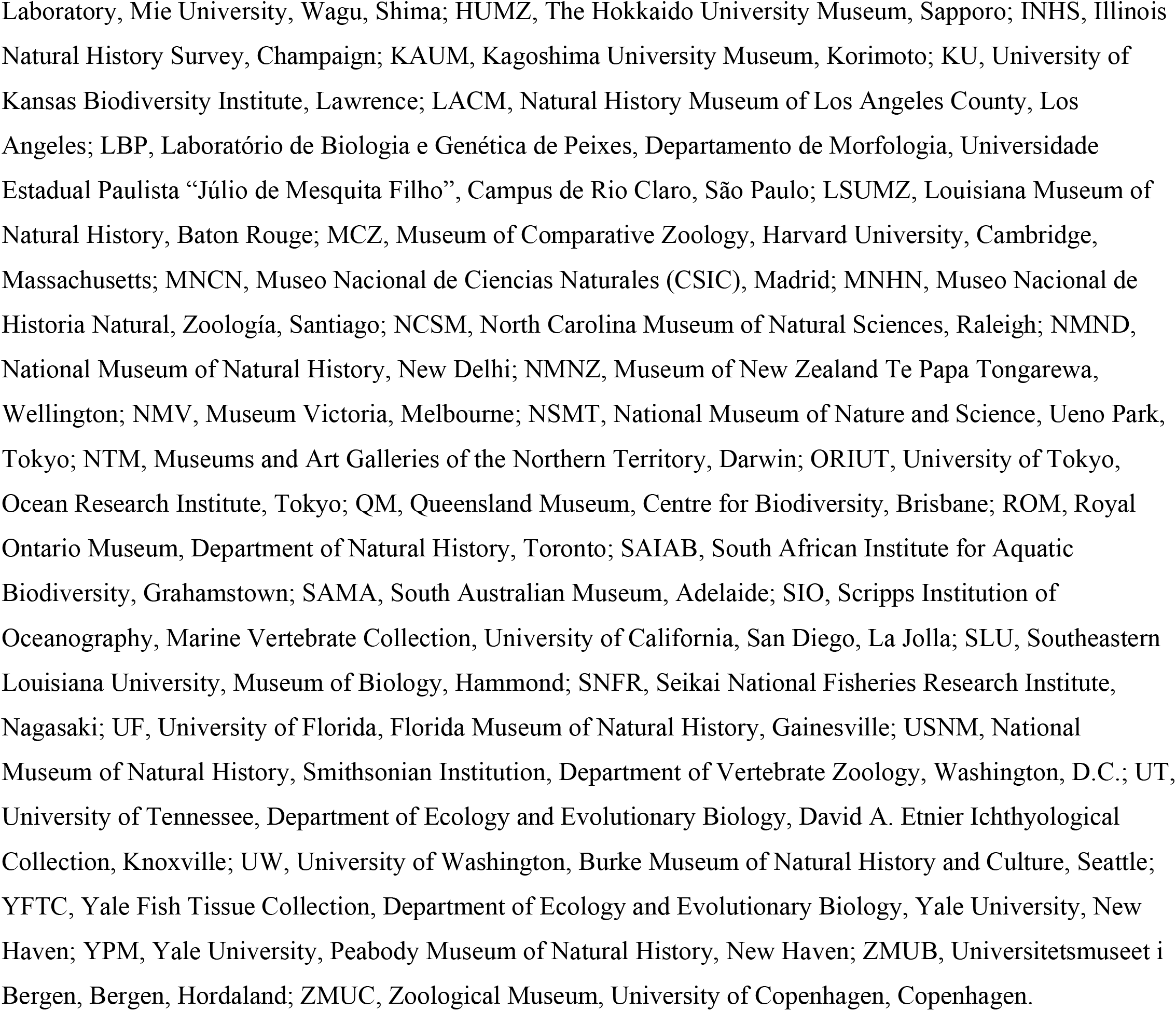
Binomial names and higher taxonomic classification of species included in the phylogeny, institutional voucher information for specimens (if applicable), and NCBI SRA BioSample accession numbers. The sixth column (‘Morphology Analyses’) denotes whether species were included in the body shape analyses, and the seventh column (‘Concordance Analyses’) denotes if samples were included in the BUCKy concordance factor analyses. NCBI BioSample numbers denoted with an asterisk (*) indicate samples for which UCE data were extracted from whole genome sequence data.

**Supplementary Table 2:** Summary of TOPD pairwise comparisons of tree topologies inferred using different methods.

|  | 702-taxon trees | 1,084-taxon trees |  |  |
| --- | --- | --- | --- | --- |
|  | Multiple-partition vs<br>IQ-TREEsingle* | IQ-TREEsingle* vs<br>IQ-TREEmulti† | IQ-TREEsingle* vs<br>RAxML | IQ-TREEmulti† vs<br>RAxML |
|  | *Acanthistius cinctus* | *Acanthistius cinctus* | *Acanthistius cinctus* | *Apogon affinis* |
|  | *Arripis trutta* | *Acanthurus triostegus* | *Acanthurus triostegus* | *Astrapogon stellatus* |
|  | *Chloroscombrus orqueta* | *Icosteus aenigmaticus* | *Apogon affinis* | *Cryptopsaras couesii* |
|  | *Icosteus aenigmaticus* |  | *Astrapogon stellatus* | *Ectodus descampsii* |
|  | *Lycodes concolor* |  | *Cryptopsaras couesii* | *Haplotaxodon microlepis* |
|  | *Pomatomus saltatrix* |  | *Ectodus descampsii* | *Nomorhamphus celebensis* |
|  | *Pseudanthias squamipinnis* |  | *Haplotaxodon microlepis* | *Phaeoptyx pigmentaria* |
|  | *Selaroides leptolepis* |  | *Icosteus aenigmaticus* | *Xenentodon cancila* |
|  | *Sillago sihama* |  | *Nomorhamphus celebensis* | *Zenarchopterus dispar* |
|  |  |  | *Phaeoptyx pigmentaria* |  |
|  |  |  | *Xenentodon cancila* |  |
|  |  |  | *Zenarchopterus dispar* |  |
| **Robinson-Foulds split**<br>**distances** | 0.033 | 0.006 | 0.012 | 0.007 |
| **# Disagreeing Splits / Total** | 46 / 1398 | 14 / 2212 | 26 / 2212 | 16 / 2212 |
| **Nodal distance** | 1.05 | 0.49 | 0.61 | 0.51 |

\*IQ-TREEsingle = tree resulting from an IQ-TREE analysis using an unpartitioned alignment † “IQ-TREEmulti = tree resulting from an IQ-TREE analysis using the ModelFinder Plus option Columns list taxa with different phylogenetic positions among the compared trees. Robinson-Foulds split distances, the number of disagreeing splits, and nodal distances are provided at the bottom of each column. All RAxML-ng analyses were performed on multiple-partition alignments that were partitioned with PartionFinder2. The topologies of the 702-taxon trees inferred using multiple-partition alignments in IQ-TREE and RAxML-ng were identical.

**Supplementary Table 3:**
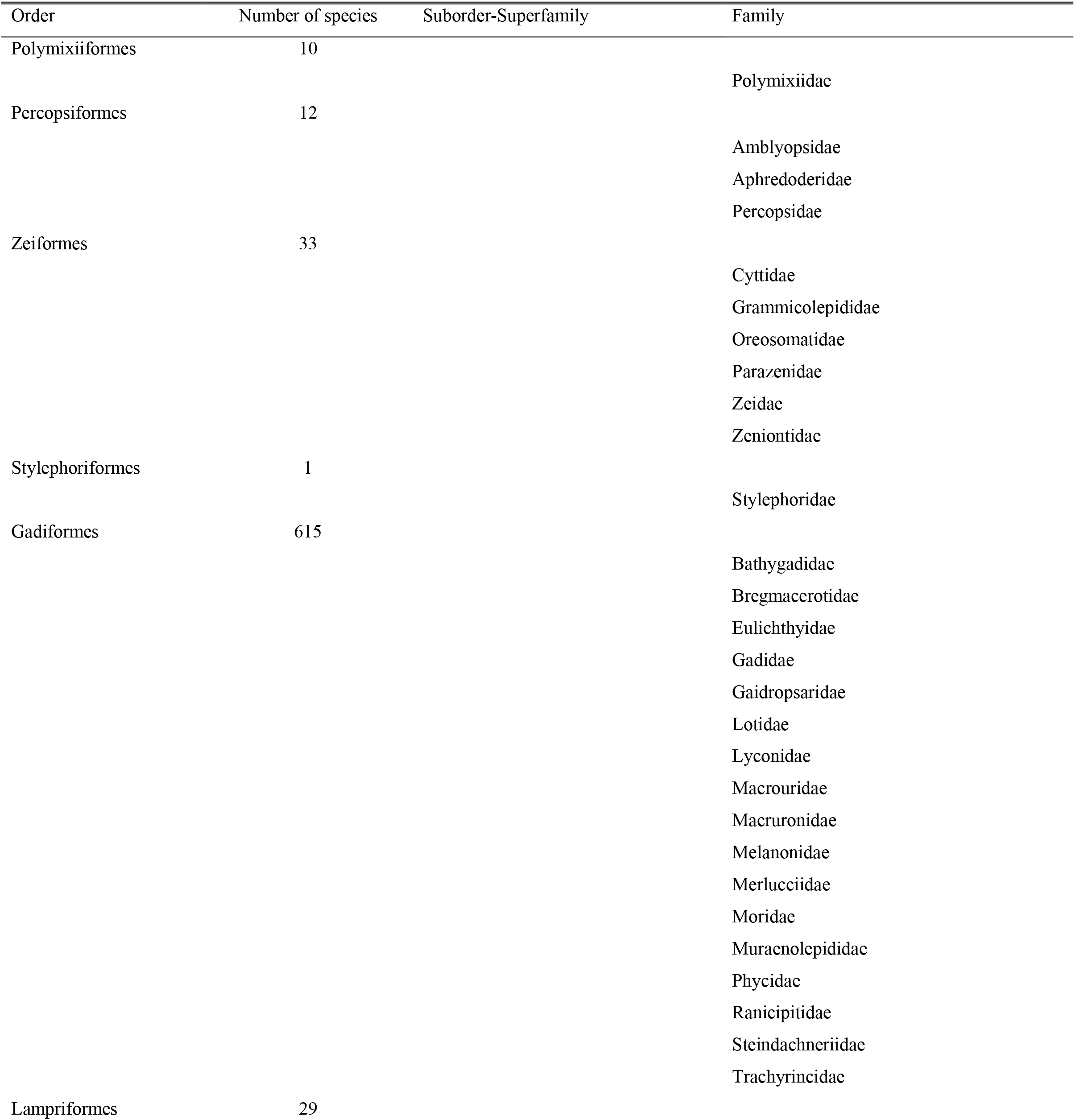
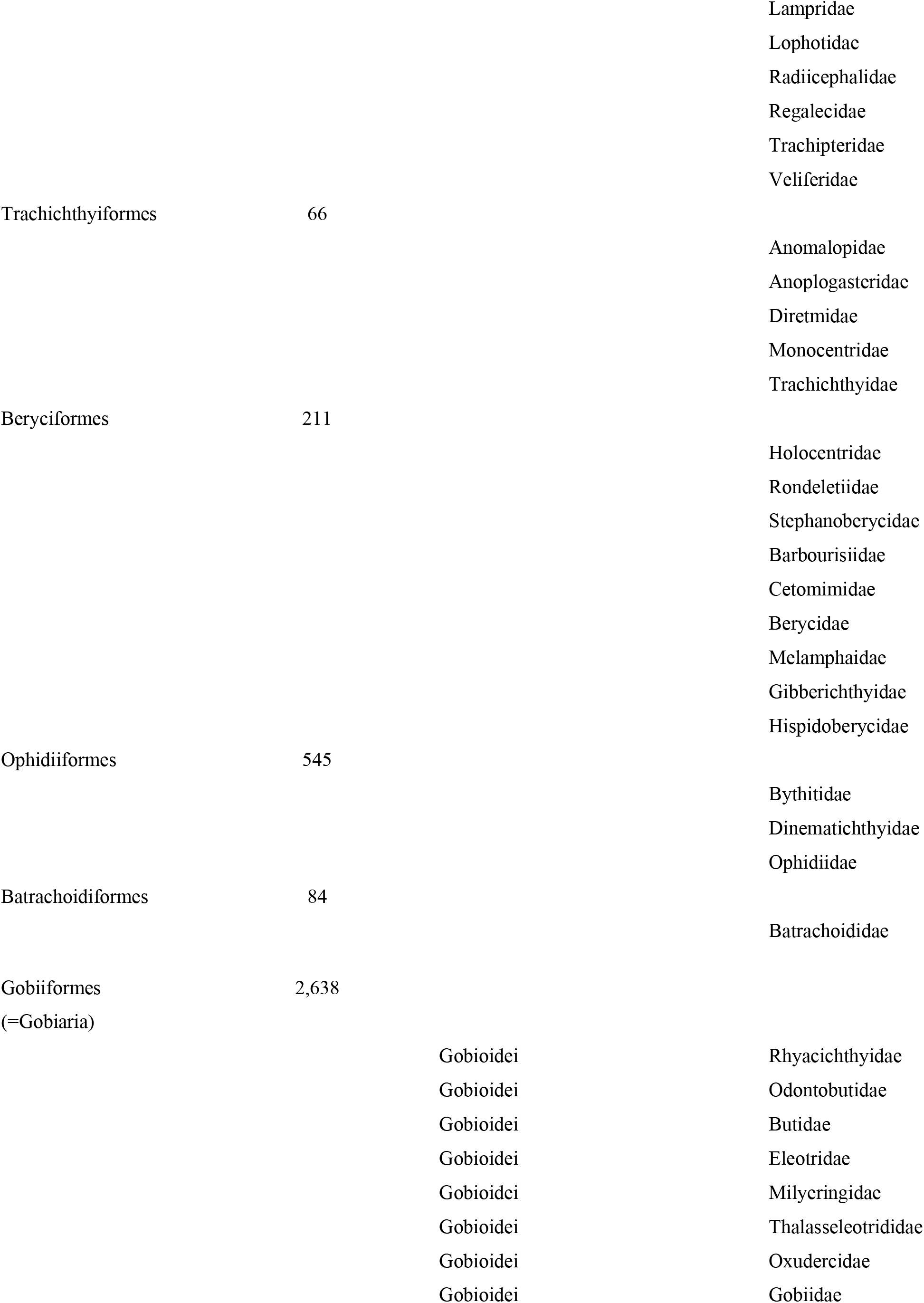
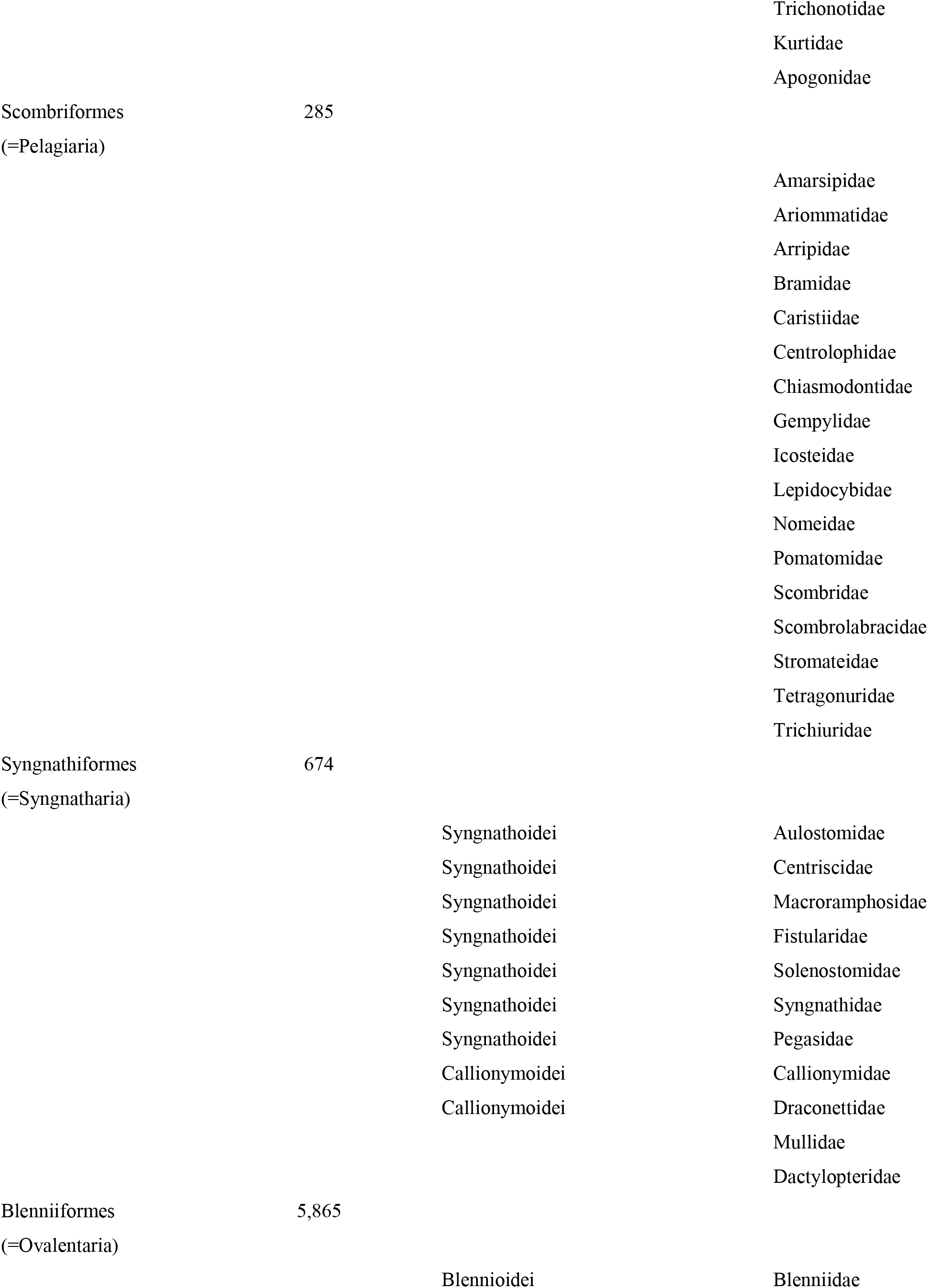
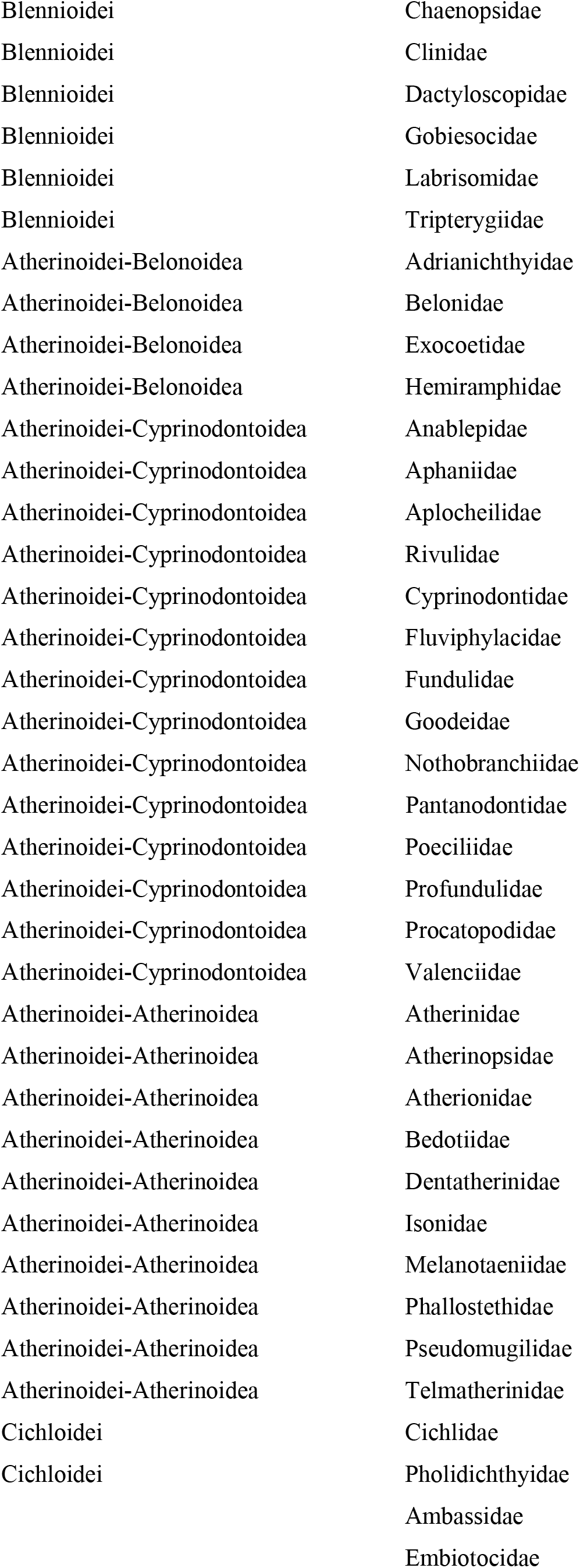
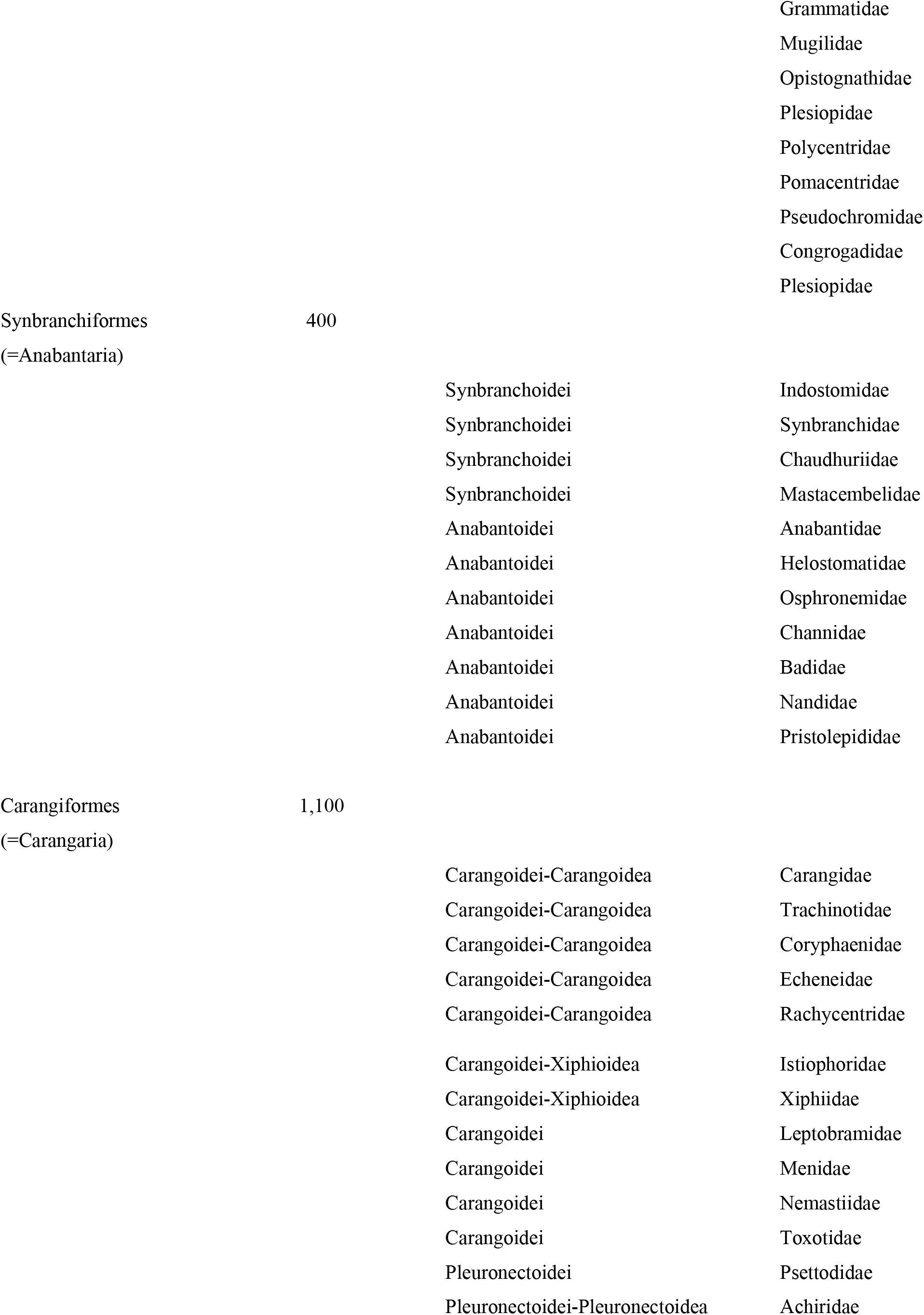
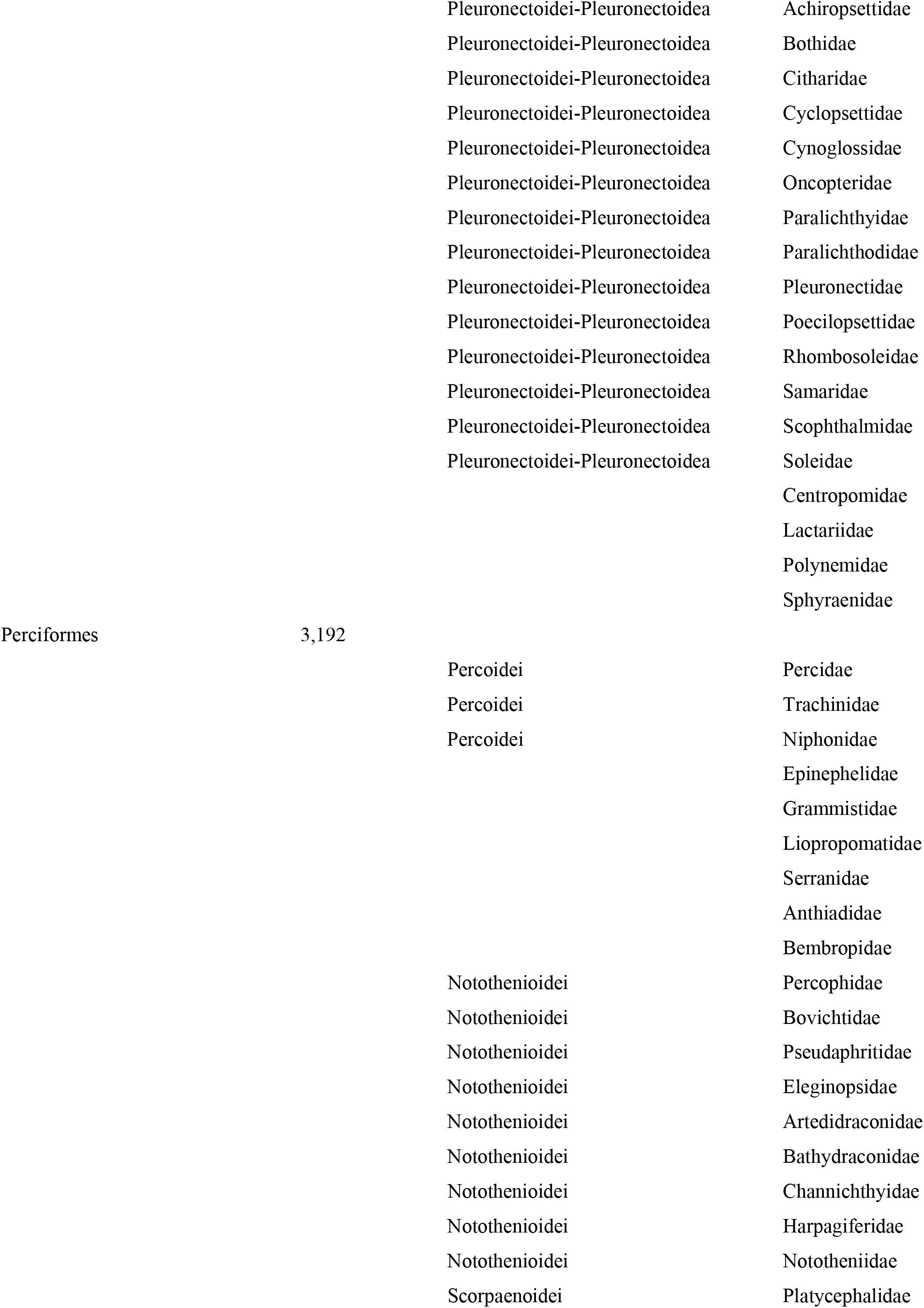
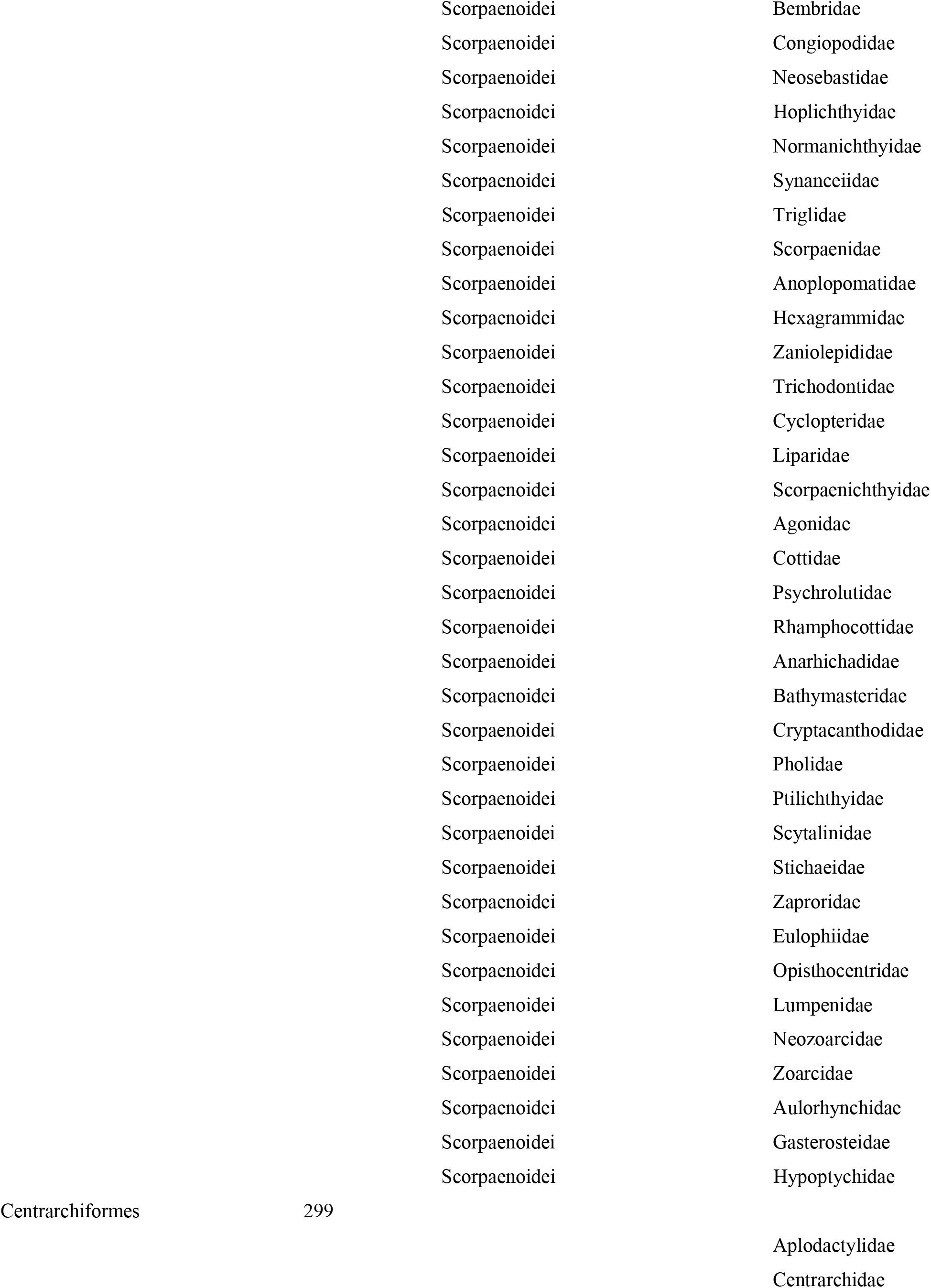
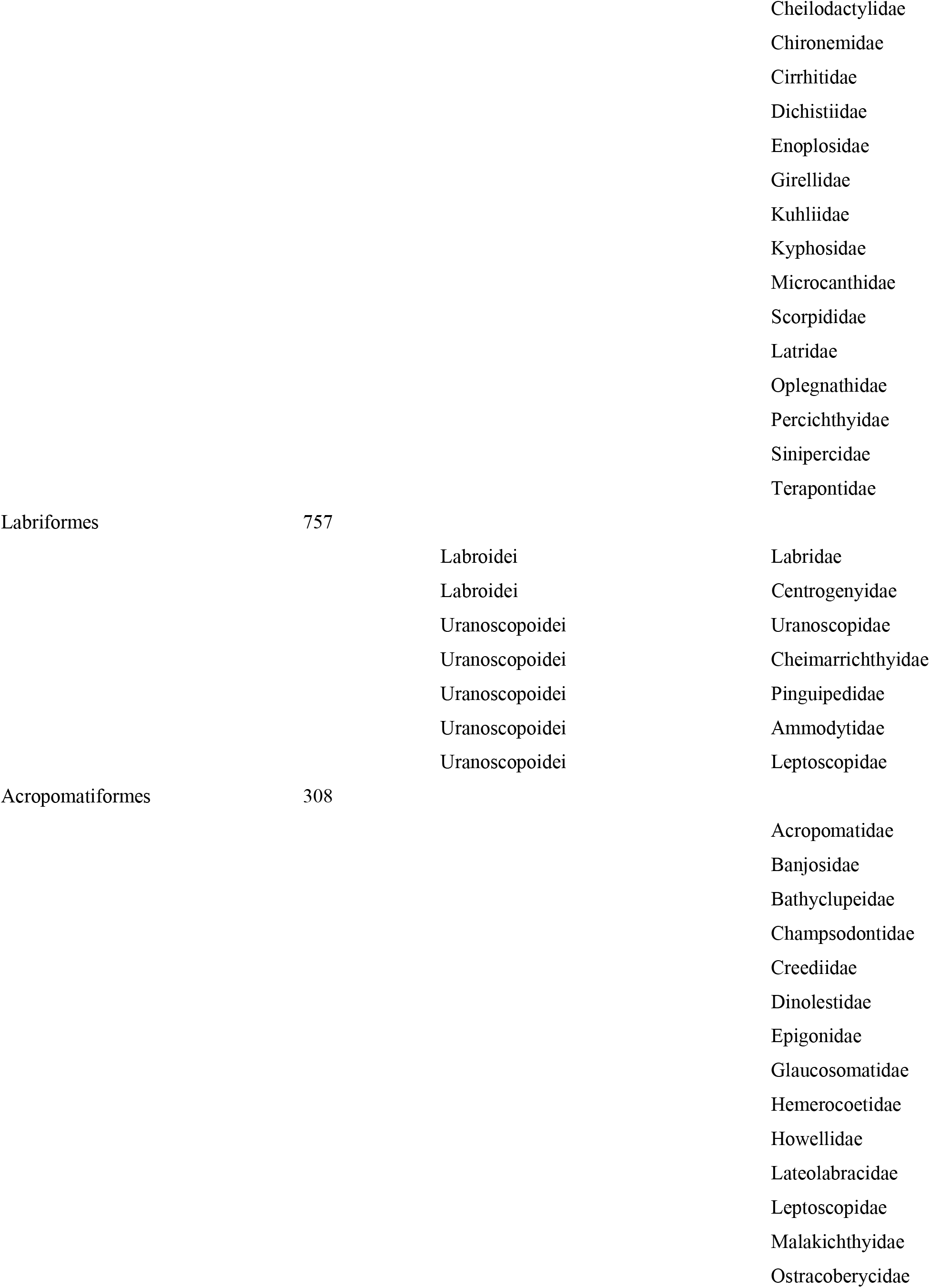
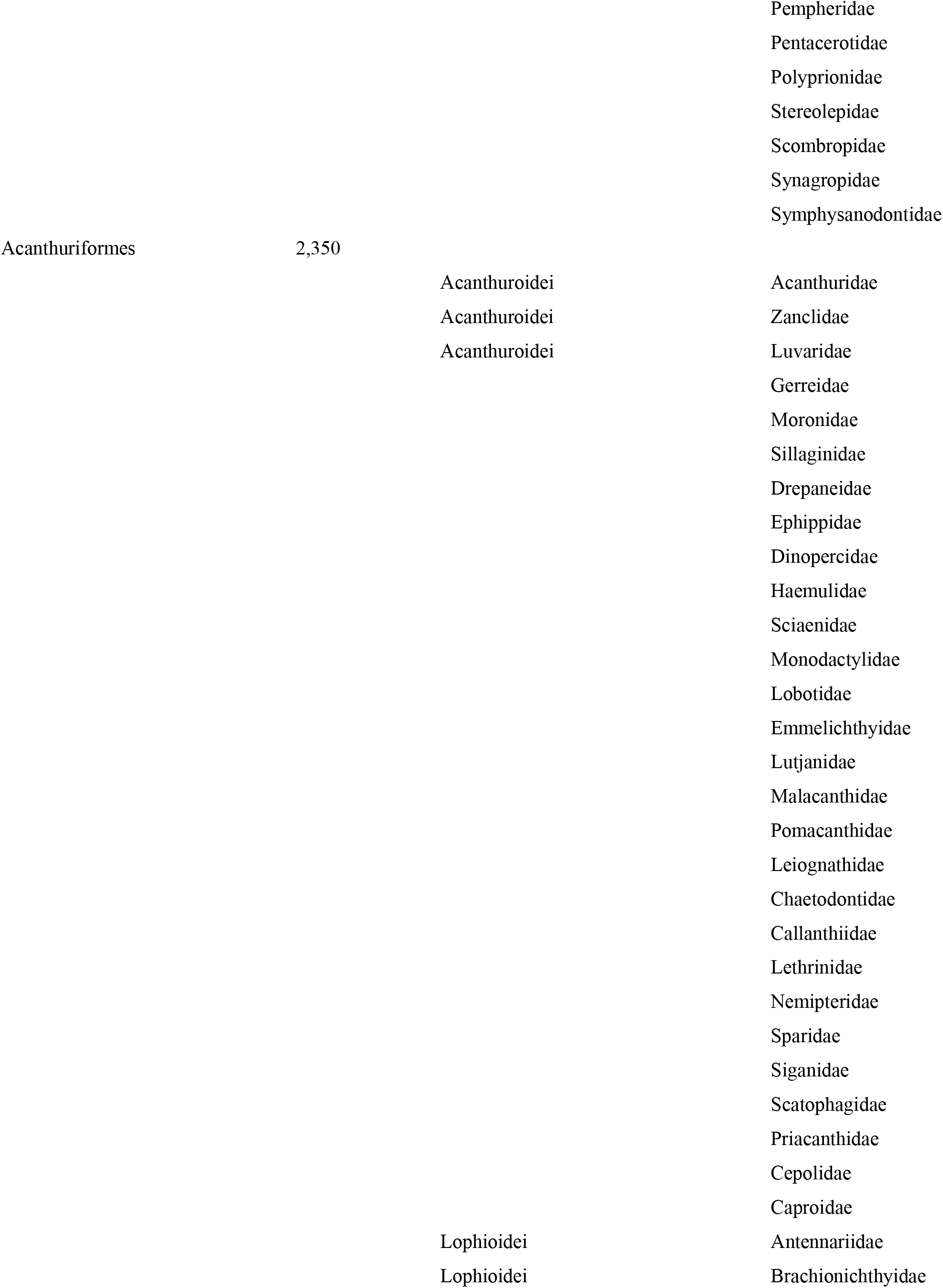
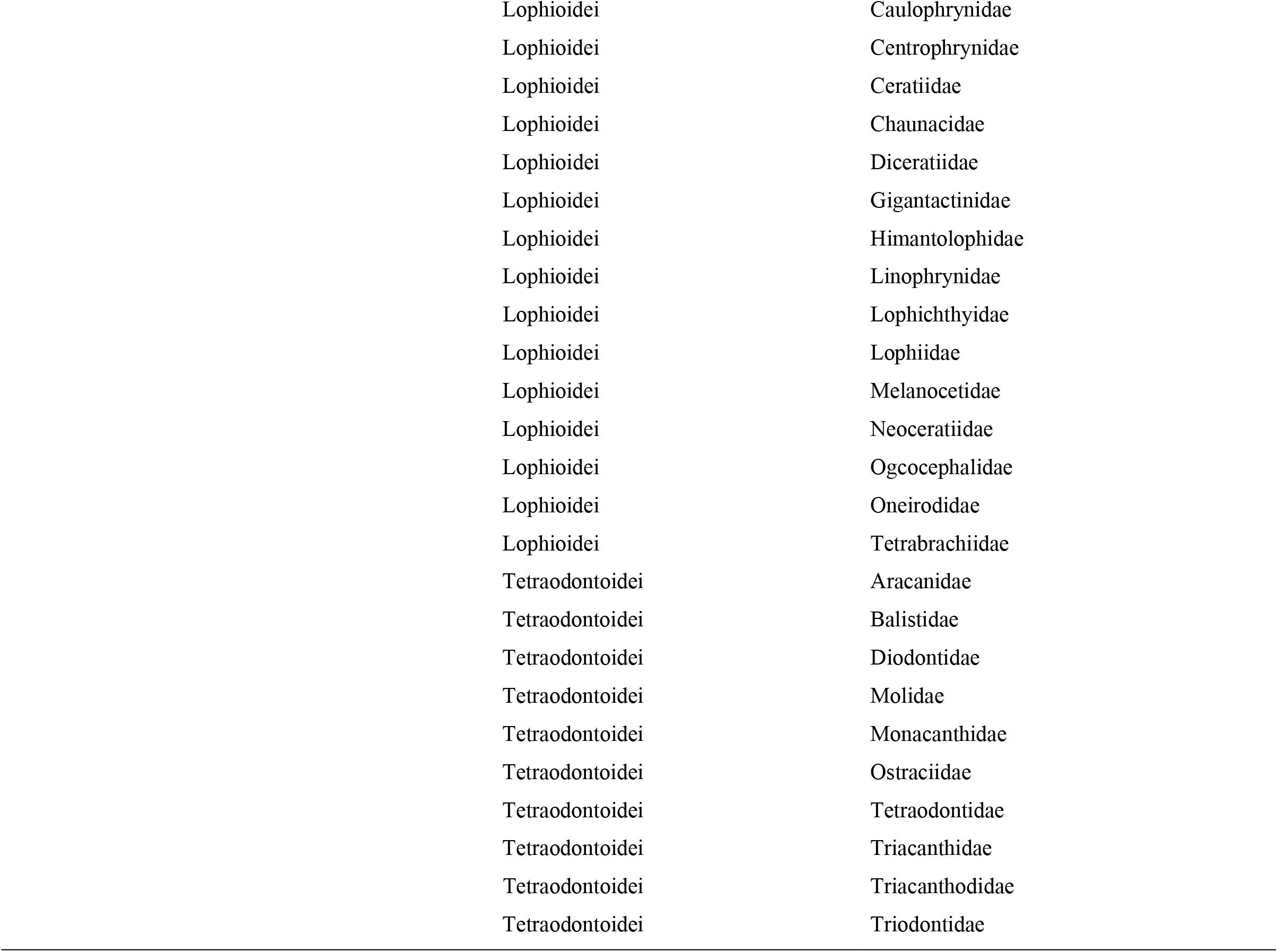
Classification of Acanthomorpha. The number of species in each taxonomic order is offered in the second column. Though every family is classified in an order, not every family is classified in a suborder or superfamily.

**Supplementary Table 4:**
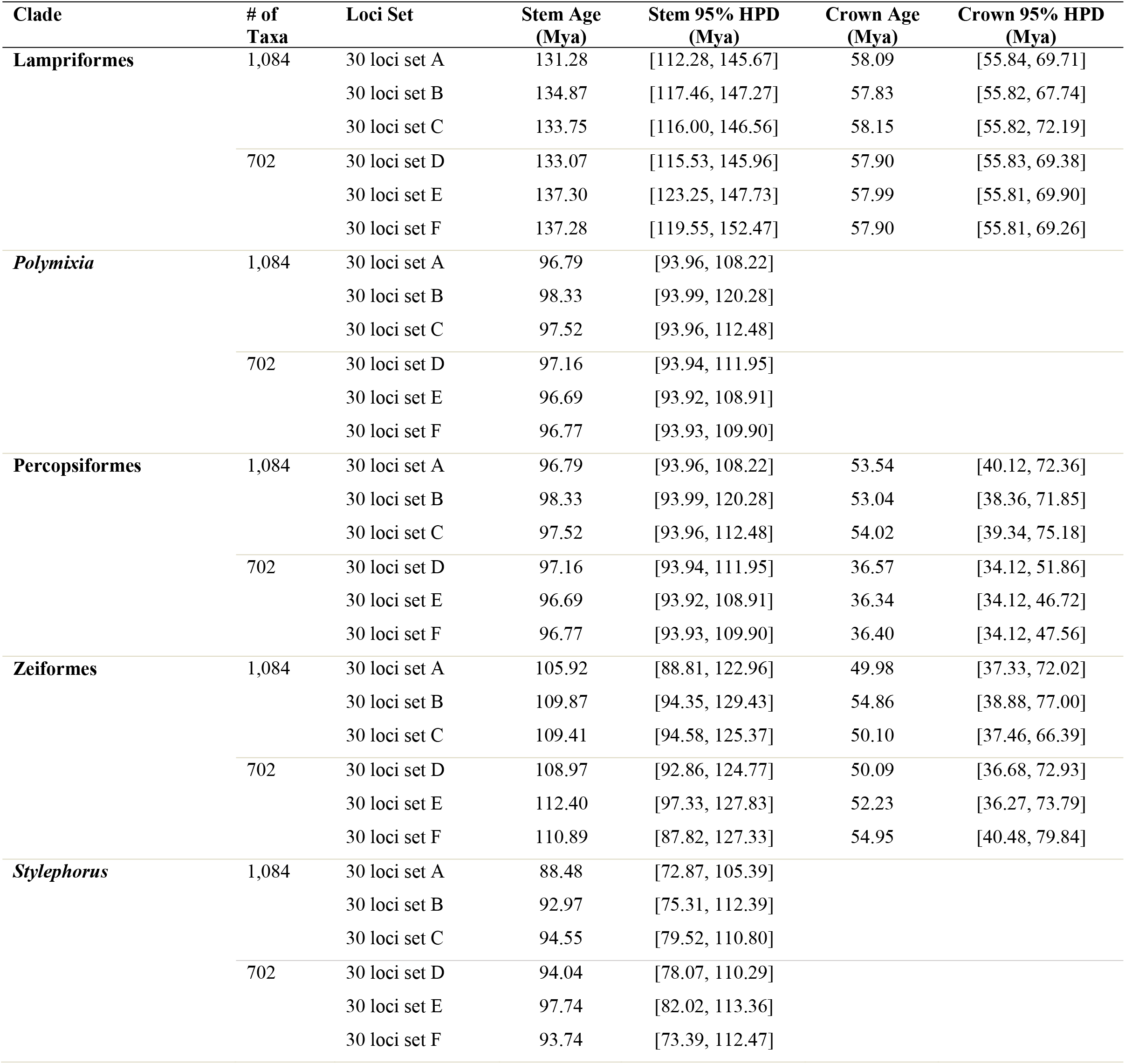
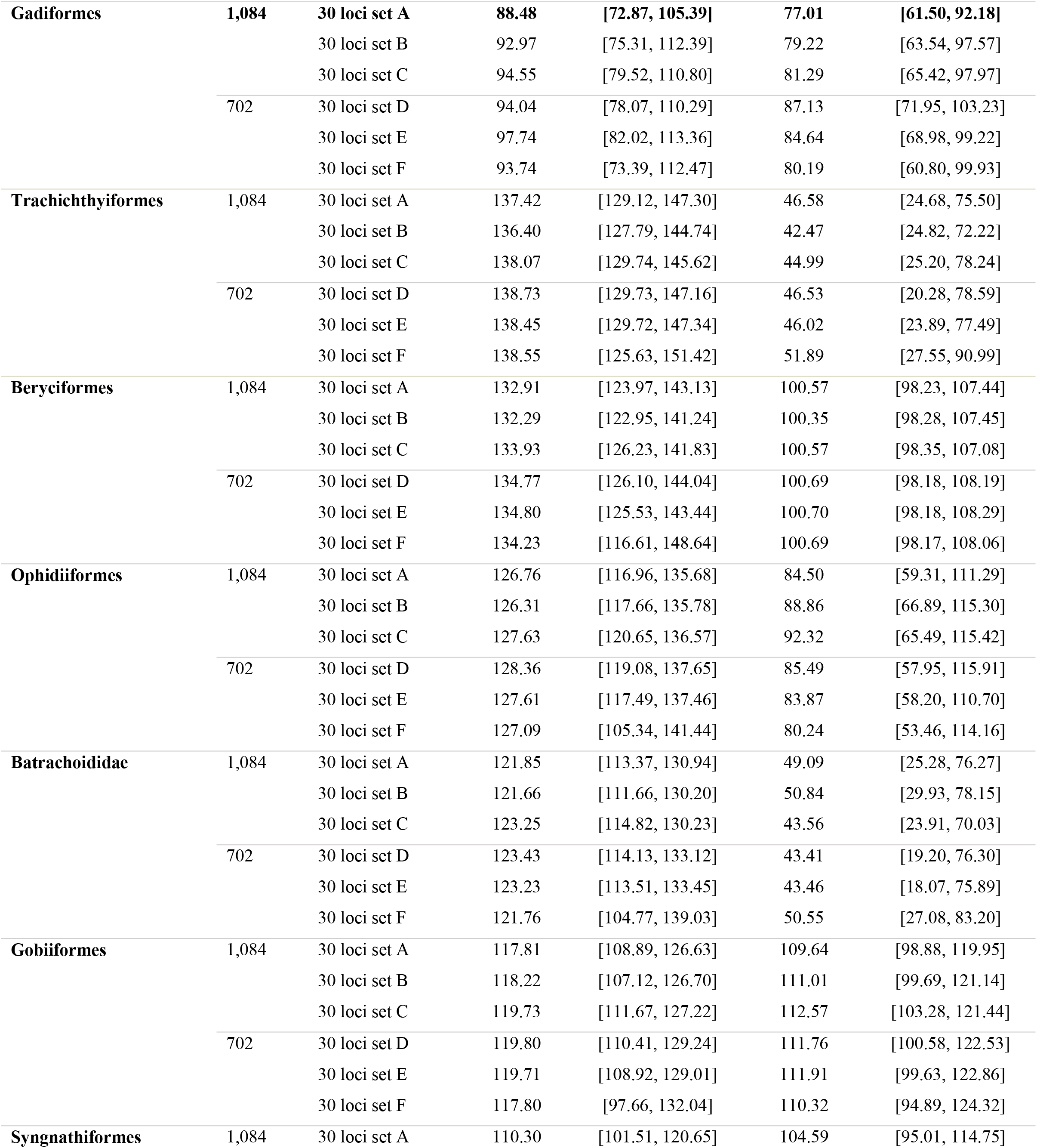
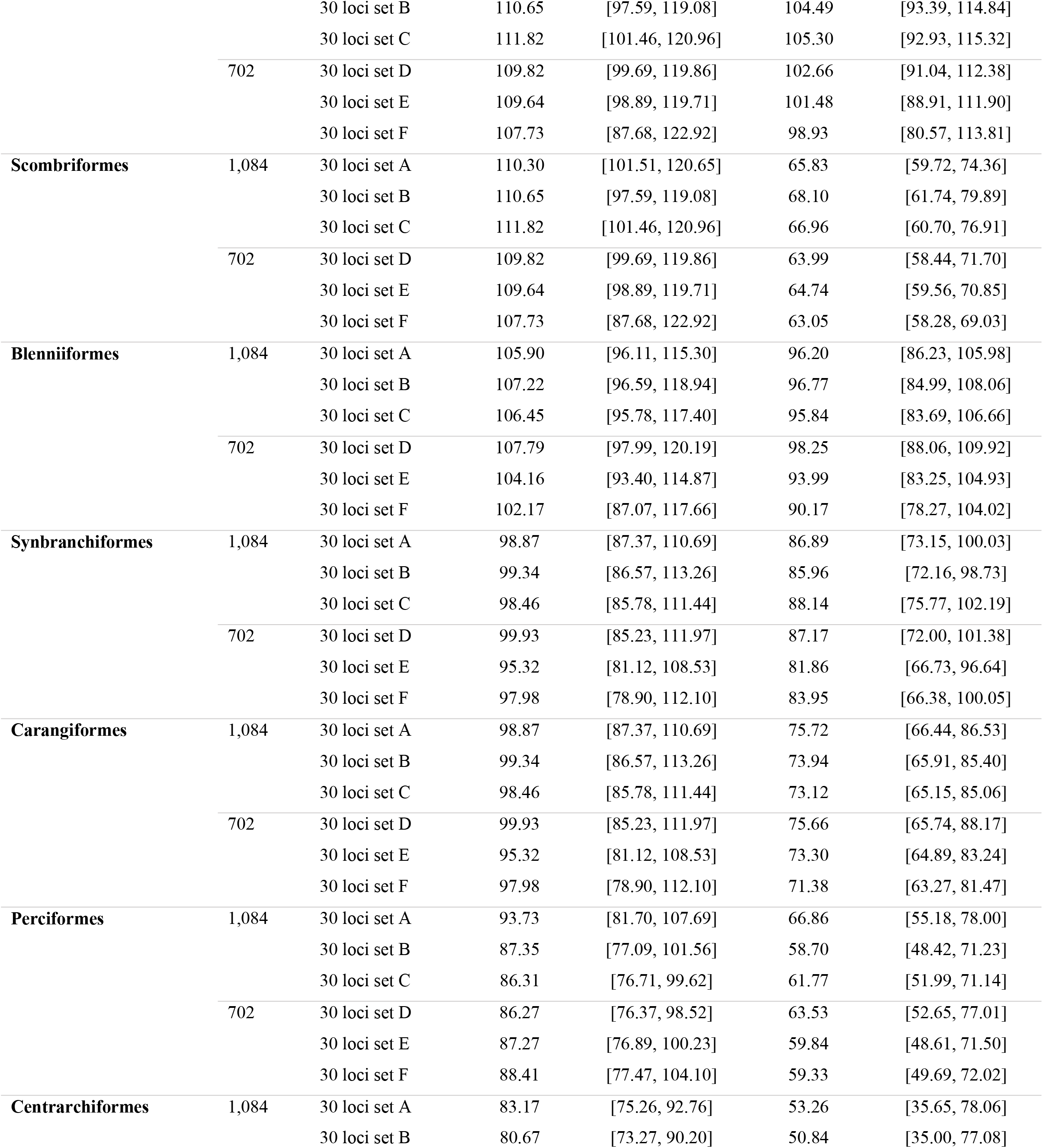
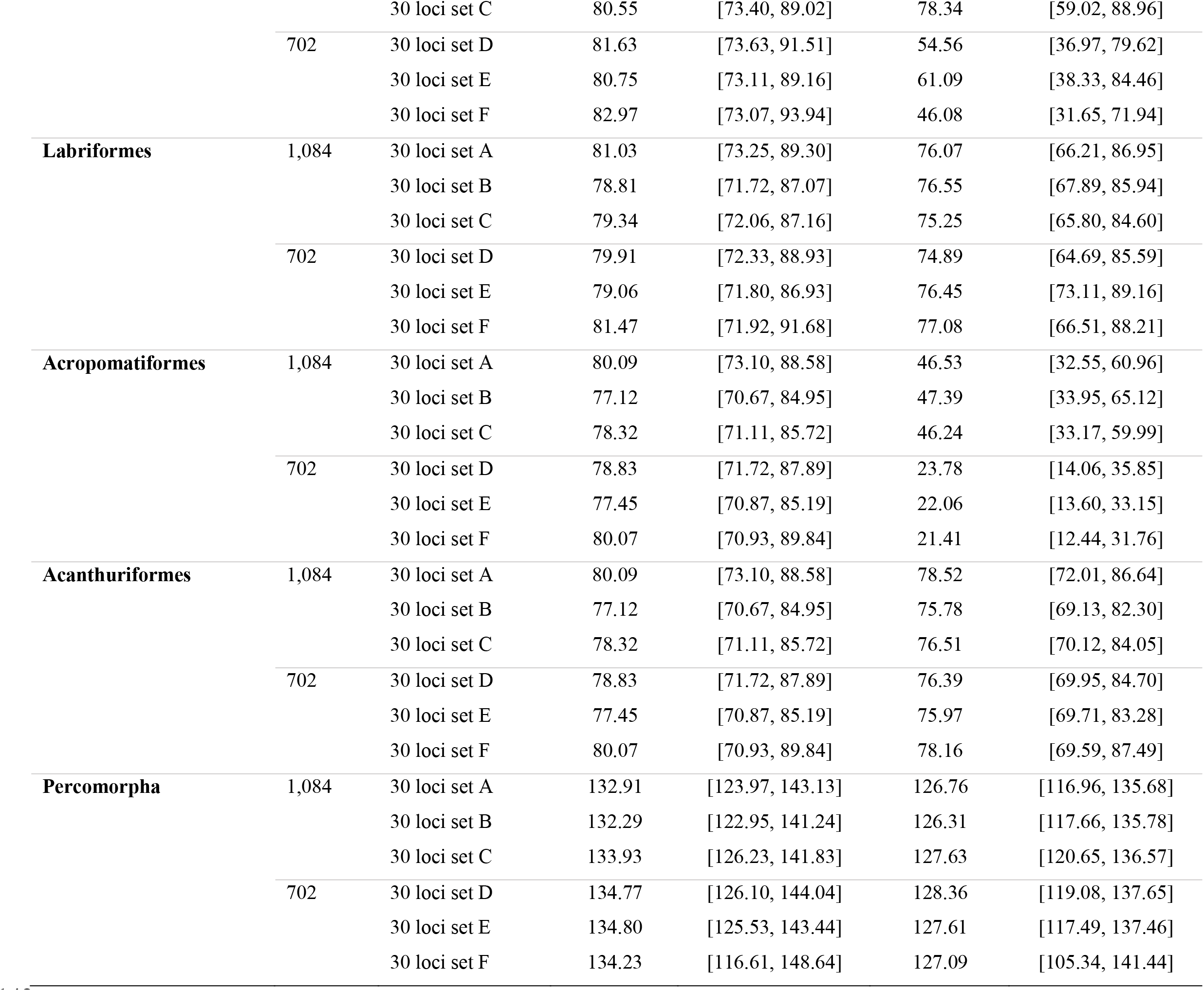
Divergence date estimates in millions of years (Mya) and 95% Highest Posterior Densities (HPD) for major acanthomorph clades, generated using six alignments of 30 UCE loci in BEAST. Estimates from the 702-taxon and 1,084-taxon time trees are both represented. All stem and crown ages are calculated from median node heights.

| Clade | # of Taxa | Loci Set | Stem Age (Mya) | Stem 95% HPD (Mya) | Crown Age (Mya) | Crown 95% HPD (Mya) |
| --- | --- | --- | --- | --- | --- | --- |
| **Lampriformes** | 1,084 | 30 loci set A | 131.28 | [112.28, 145.67] | 58.09 | [55.84, 69.71] |
|  |  | 30 loci set B | 134.87 | [117.46, 147.27] | 57.83 | [55.82, 67.74] |
|  |  | 30 loci set C | 133.75 | [116.00, 146.56] | 58.15 | [55.82, 72.19] |
|  | 702 | 30 loci set D | 133.07 | [115.53, 145.96] | 57.90 | [55.83, 69.38] |
|  |  | 30 loci set E | 137.30 | [123.25, 147.73] | 57.99 | [55.81, 69.90] |
|  |  | 30 loci set F | 137.28 | [119.55, 152.47] | 57.90 | [55.81, 69.26] |
| **Polymixia** | 1,084 | 30 loci set A | 96.79 | [93.96, 108.22] |  |  |
|  |  | 30 loci set B | 98.33 | [93.99, 120.28] |  |  |
|  |  | 30 loci set C | 97.52 | [93.96, 112.48] |  |  |
|  | 702 | 30 loci set D | 97.16 | [93.94, 111.95] |  |  |
|  |  | 30 loci set E | 96.69 | [93.92, 108.91] |  |  |
|  |  | 30 loci set F | 96.77 | [93.93, 109.90] |  |  |
| **Percopsiformes** | 1,084 | 30 loci set A | 96.79 | [93.96, 108.22] | 53.54 | [40.12, 72.36] |
|  |  | 30 loci set B | 98.33 | [93.99, 120.28] | 53.04 | [38.36, 71.85] |
|  |  | 30 loci set C | 97.52 | [93.96, 112.48] | 54.02 | [39.34, 75.18] |
|  | 702 | 30 loci set D | 97.16 | [93.94, 111.95] | 36.57 | [34.12, 51.86] |
|  |  | 30 loci set E | 96.69 | [93.92, 108.91] | 36.34 | [34.12, 46.72] |
|  |  | 30 loci set F | 96.77 | [93.93, 109.90] | 36.40 | [34.12, 47.56] |
| **Zeiformes** | 1,084 | 30 loci set A | 105.92 | [88.81, 122.96] | 49.98 | [37.33, 72.02] |
|  |  | 30 loci set B | 109.87 | [94.35, 129.43] | 54.86 | [38.88, 77.00] |
|  |  | 30 loci set C | 109.41 | [94.58, 125.37] | 50.10 | [37.46, 66.39] |
|  | 702 | 30 loci set D | 108.97 | [92.86, 124.77] | 50.09 | [36.68, 72.93] |
|  |  | 30 loci set E | 112.40 | [97.33, 127.83] | 52.23 | [36.27, 73.79] |
|  |  | 30 loci set F | 110.89 | [87.82, 127.33] | 54.95 | [40.48, 79.84] |
| **Stylephorus** | 1,084 | 30 loci set A | 88.48 | [72.87, 105.39] |  |  |
|  |  | 30 loci set B | 92.97 | [75.31, 112.39] |  |  |
|  |  | 30 loci set C | 94.55 | [79.52, 110.80] |  |  |
|  | 702 | 30 loci set D | 94.04 | [78.07, 110.29] |  |  |
|  |  | 30 loci set E | 97.74 | [82.02, 113.36] |  |  |
|  |  | 30 loci set F | 93.74 | [73.39, 112.47] |  |  |
| **Gadiiformes** | 1,084 | **30 loci set A** | **88.48** | **[72.87, 105.39]** | **77.01** | **[61.50, 92.18]** |
|  |  | 30 loci set B | 92.97 | [75.31, 112.39] | 79.22 | [63.54, 97.57] |
|  |  | 30 loci set C | 94.55 | [79.52, 110.80] | 81.29 | [65.42, 97.97] |
|  | 702 | 30 loci set D | 94.04 | [78.07, 110.29] | 87.13 | [71.95, 103.23] |
|  |  | 30 loci set E | 97.74 | [82.02, 113.36] | 84.64 | [68.98, 99.22] |
|  |  | 30 loci set F | 93.74 | [73.39, 112.47] | 80.19 | [60.80, 99.93] |
| **Trachichthyiformes** | 1,084 | 30 loci set A | 137.42 | [129.12, 147.30] | 46.58 | [24.68, 75.50] |
|  |  | 30 loci set B | 136.40 | [127.79, 144.74] | 42.47 | [24.82, 72.22] |
|  |  | 30 loci set C | 138.07 | [129.74, 145.62] | 44.99 | [25.20, 78.24] |
|  | 702 | 30 loci set D | 138.73 | [129.73, 147.16] | 46.53 | [20.28, 78.59] |
|  |  | 30 loci set E | 138.45 | [129.72, 147.34] | 46.02 | [23.89, 77.49] |
|  |  | 30 loci set F | 138.55 | [125.63, 151.42] | 51.89 | [27.55, 90.99] |
| **Beryciformes** | 1,084 | 30 loci set A | 132.91 | [123.97, 143.13] | 100.57 | [98.23, 107.44] |
|  |  | 30 loci set B | 132.29 | [122.95, 141.24] | 100.35 | [98.28, 107.45] |
|  |  | 30 loci set C | 133.93 | [126.23, 141.83] | 100.57 | [98.35, 107.08] |
|  | 702 | 30 loci set D | 134.77 | [126.10, 144.04] | 100.69 | [98.18, 108.19] |
|  |  | 30 loci set E | 134.80 | [125.53, 143.44] | 100.70 | [98.18, 108.29] |
|  |  | 30 loci set F | 134.23 | [116.61, 148.64] | 100.69 | [98.17, 108.06] |
| **Ophidiiformes** | 1,084 | 30 loci set A | 126.76 | [116.96, 135.68] | 84.50 | [59.31, 111.29] |
|  |  | 30 loci set B | 126.31 | [117.66, 135.78] | 88.86 | [66.89, 115.30] |
|  |  | 30 loci set C | 127.63 | [120.65, 136.57] | 92.32 | [65.49, 115.42] |
|  | 702 | 30 loci set D | 128.36 | [119.08, 137.65] | 85.49 | [57.95, 115.91] |
|  |  | 30 loci set E | 127.61 | [117.49, 137.46] | 83.87 | [58.20, 110.70] |
|  |  | 30 loci set F | 127.09 | [105.34, 141.44] | 80.24 | [53.46, 114.16] |
| **Batrachoididae** | 1,084 | 30 loci set A | 121.85 | [113.37, 130.94] | 49.09 | [25.28, 76.27] |
|  |  | 30 loci set B | 121.66 | [111.66, 130.20] | 50.84 | [29.93, 78.15] |
|  |  | 30 loci set C | 123.25 | [114.82, 130.23] | 43.56 | [23.91, 70.03] |
|  | 702 | 30 loci set D | 123.43 | [114.13, 133.12] | 43.41 | [19.20, 76.30] |
|  |  | 30 loci set E | 123.23 | [113.51, 133.45] | 43.46 | [18.07, 75.89] |
|  |  | 30 loci set F | 121.76 | [104.77, 139.03] | 50.55 | [27.08, 83.20] |
| **Gobiiformes** | 1,084 | 30 loci set A | 117.81 | [108.89, 126.63] | 109.64 | [98.88, 119.95] |
|  |  | 30 loci set B | 118.22 | [107.12, 126.70] | 111.01 | [99.69, 121.14] |
|  |  | 30 loci set C | 119.73 | [111.67, 127.22] | 112.57 | [103.28, 121.44] |
|  | 702 | 30 loci set D | 119.80 | [110.41, 129.24] | 111.76 | [100.58, 122.53] |
|  |  | 30 loci set E | 119.71 | [108.92, 129.01] | 111.91 | [99.63, 122.86] |
|  |  | 30 loci set F | 117.80 | [97.66, 132.04] | 110.32 | [94.89, 124.32] |
| **Syngnathiformes** | 1,084 | 30 loci set A | 110.30 | [101.51, 120.65] | 104.59 | [95.01, 114.75] |
|  |  | 30 loci set B | 110.65 | [97.59, 119.08] | 104.49 | [93.39, 114.84] |
|  |  | 30 loci set C | 111.82 | [101.46, 120.96] | 105.30 | [92.93, 115.32] |
|  | 702 | 30 loci set D | 109.82 | [99.69, 119.86] | 102.66 | [91.04, 112.38] |
|  |  | 30 loci set E | 109.64 | [98.89, 119.71] | 101.48 | [88.91, 111.90] |
|  |  | 30 loci set F | 107.73 | [87.68, 122.92] | 98.93 | [80.57, 113.81] |
| **Scombriformes** | 1,084 | 30 loci set A | 110.30 | [101.51, 120.65] | 65.83 | [59.72, 74.36] |
|  |  | 30 loci set B | 110.65 | [97.59, 119.08] | 68.10 | [61.74, 79.89] |
|  |  | 30 loci set C | 111.82 | [101.46, 120.96] | 66.96 | [60.70, 76.91] |
|  | 702 | 30 loci set D | 109.82 | [99.69, 119.86] | 63.99 | [58.44, 71.70] |
|  |  | 30 loci set E | 109.64 | [98.89, 119.71] | 64.74 | [59.56, 70.85] |
|  |  | 30 loci set F | 107.73 | [87.68, 122.92] | 63.05 | [58.28, 69.03] |
| **Blenniiformes** | 1,084 | 30 loci set A | 105.90 | [96.11, 115.30] | 96.20 | [86.23, 105.98] |
|  |  | 30 loci set B | 107.22 | [96.59, 118.94] | 96.77 | [84.99, 108.06] |
|  |  | 30 loci set C | 106.45 | [95.78, 117.40] | 95.84 | [83.69, 106.66] |
|  | 702 | 30 loci set D | 107.79 | [97.99, 120.19] | 98.25 | [88.06, 109.92] |
|  |  | 30 loci set E | 104.16 | [93.40, 114.87] | 93.99 | [83.25, 104.93] |
|  |  | 30 loci set F | 102.17 | [87.07, 117.66] | 90.17 | [78.27, 104.02] |
| **Synbranchiiformes** | 1,084 | 30 loci set A | 98.87 | [87.37, 110.69] | 86.89 | [73.15, 100.03] |
|  |  | 30 loci set B | 99.34 | [86.57, 113.26] | 85.96 | [72.16, 98.73] |
|  |  | 30 loci set C | 98.46 | [85.78, 111.44] | 88.14 | [75.77, 102.19] |
|  | 702 | 30 loci set D | 99.93 | [85.23, 111.97] | 87.17 | [72.00, 101.38] |
|  |  | 30 loci set E | 95.32 | [81.12, 108.53] | 81.86 | [66.73, 96.64] |
|  |  | 30 loci set F | 97.98 | [78.90, 112.10] | 83.95 | [66.38, 100.05] |
| **Carangiformes** | 1,084 | 30 loci set A | 98.87 | [87.37, 110.69] | 75.72 | [66.44, 86.53] |
|  |  | 30 loci set B | 99.34 | [86.57, 113.26] | 73.94 | [65.91, 85.40] |
|  |  | 30 loci set C | 98.46 | [85.78, 111.44] | 73.12 | [65.15, 85.06] |
|  | 702 | 30 loci set D | 99.93 | [85.23, 111.97] | 75.66 | [65.74, 88.17] |
|  |  | 30 loci set E | 95.32 | [81.12, 108.53] | 73.30 | [64.89, 83.24] |
|  |  | 30 loci set F | 97.98 | [78.90, 112.10] | 71.38 | [63.27, 81.47] |
| **Perciformes** | 1,084 | 30 loci set A | 93.73 | [81.70, 107.69] | 66.86 | [55.18, 78.00] |
|  |  | 30 loci set B | 87.35 | [77.09, 101.56] | 58.70 | [48.42, 71.23] |
|  |  | 30 loci set C | 86.31 | [76.71, 99.62] | 61.77 | [51.99, 71.14] |
|  | 702 | 30 loci set D | 86.27 | [76.37, 98.52] | 63.53 | [52.65, 77.01] |
|  |  | 30 loci set E | 87.27 | [76.89, 100.23] | 59.84 | [48.61, 71.50] |
|  |  | 30 loci set F | 88.41 | [77.47, 104.10] | 59.33 | [49.69, 72.02] |
| **Centrarchiformes** | 1,084 | 30 loci set A | 83.17 | [75.26, 92.76] | 53.26 | [35.65, 78.06] |
|  |  | 30 loci set B | 80.67 | [73.27, 90.20] | 50.84 | [35.00, 77.08] |
|  |  | 30 loci set C | 80.55 | [73.40, 89.02] | 78.34 | [59.02, 88.96] |
|  | 702 | 30 loci set D | 81.63 | [73.63, 91.51] | 54.56 | [36.97, 79.62] |
|  |  | 30 loci set E | 80.75 | [73.11, 89.16] | 61.09 | [38.33, 84.46] |
|  |  | 30 loci set F | 82.97 | [73.07, 93.94] | 46.08 | [31.65, 71.94] |
| **Labriformes** | 1,084 | 30 loci set A | 81.03 | [73.25, 89.30] | 76.07 | [66.21, 86.95] |
|  |  | 30 loci set B | 78.81 | [71.72, 87.07] | 76.55 | [67.89, 85.94] |
|  |  | 30 loci set C | 79.34 | [72.06, 87.16] | 75.25 | [65.80, 84.60] |
|  | 702 | 30 loci set D | 79.91 | [72.33, 88.93] | 74.89 | [64.69, 85.59] |
|  |  | 30 loci set E | 79.06 | [71.80, 86.93] | 76.45 | [73.11, 89.16] |
|  |  | 30 loci set F | 81.47 | [71.92, 91.68] | 77.08 | [66.51, 88.21] |
| **Acropomatiformes** | 1,084 | 30 loci set A | 80.09 | [73.10, 88.58] | 46.53 | [32.55, 60.96] |
|  |  | 30 loci set B | 77.12 | [70.67, 84.95] | 47.39 | [33.95, 65.12] |
|  |  | 30 loci set C | 78.32 | [71.11, 85.72] | 46.24 | [33.17, 59.99] |
|  | 702 | 30 loci set D | 78.83 | [71.72, 87.89] | 23.78 | [14.06, 35.85] |
|  |  | 30 loci set E | 77.45 | [70.87, 85.19] | 22.06 | [13.60, 33.15] |
|  |  | 30 loci set F | 80.07 | [70.93, 89.84] | 21.41 | [12.44, 31.76] |
| **Acanthuriformes** | 1,084 | 30 loci set A | 80.09 | [73.10, 88.58] | 78.52 | [72.01, 86.64] |
|  |  | 30 loci set B | 77.12 | [70.67, 84.95] | 75.78 | [69.13, 82.30] |
|  |  | 30 loci set C | 78.32 | [71.11, 85.72] | 76.51 | [70.12, 84.05] |
|  | 702 | 30 loci set D | 78.83 | [71.72, 87.89] | 76.39 | [69.95, 84.70] |
|  |  | 30 loci set E | 77.45 | [70.87, 85.19] | 75.97 | [69.71, 83.28] |
|  |  | 30 loci set F | 80.07 | [70.93, 89.84] | 78.16 | [69.59, 87.49] |
| **Percomorpha** | 1,084 | 30 loci set A | 132.91 | [123.97, 143.13] | 126.76 | [116.96, 135.68] |
|  |  | 30 loci set B | 132.29 | [122.95, 141.24] | 126.31 | [117.66, 135.78] |
|  |  | 30 loci set C | 133.93 | [126.23, 141.83] | 127.63 | [120.65, 136.57] |
|  | 702 | 30 loci set D | 134.77 | [126.10, 144.04] | 128.36 | [119.08, 137.65] |
|  |  | 30 loci set E | 134.80 | [125.53, 143.44] | 127.61 | [117.49, 137.46] |
|  |  | 30 loci set F | 134.23 | [116.61, 148.64] | 127.09 | [105.34, 141.44] |

**Supplementary Fig. 1:**
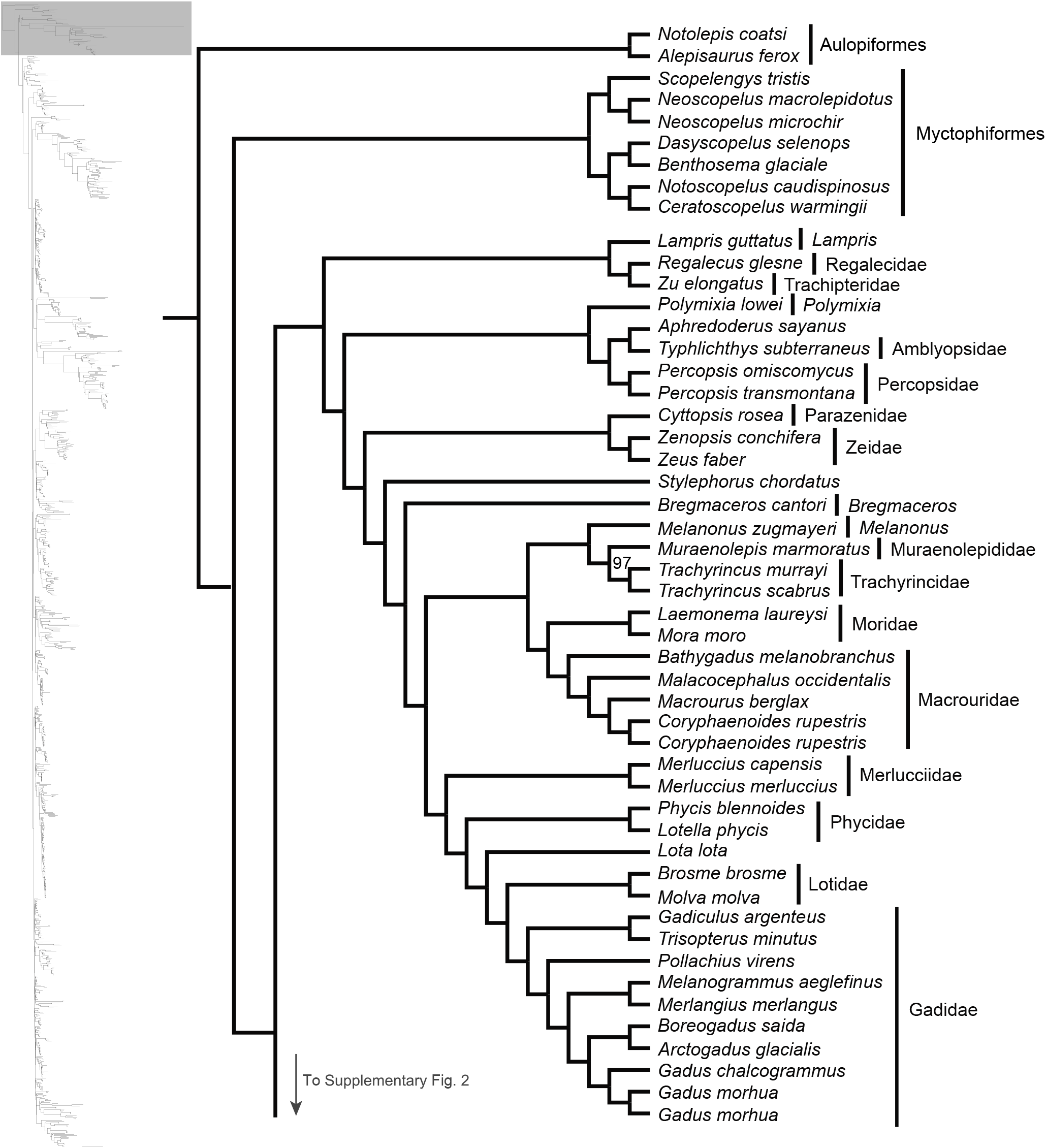
Maximum likelihood phylogeny inferred in IQ-TREE. A guide tree on the left marks (with a gray rectangle), the region of the acanthomorph tree represented in the figure. Numbers at nodes reflect bootstrap support values. All nodes without a numerical annotation have 100% bootstrap support. Orders, families or genera for taxa are listed to the right of the vertical black bars. Any tips left unassigned to a higher taxon are monotypic or monogeneric taxa.

**Supplementary Fig. 2:**
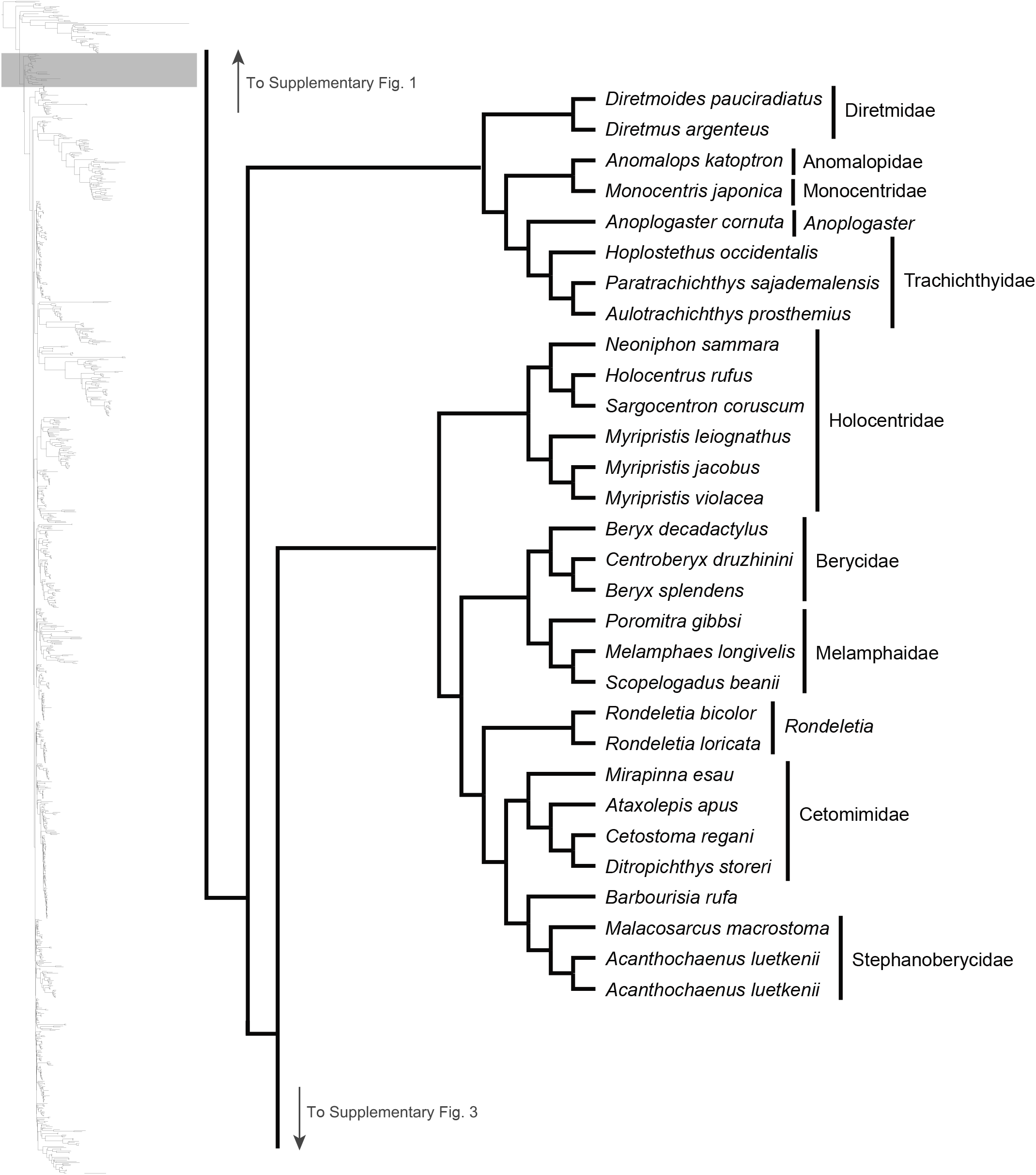
Maximum likelihood phylogeny inferred in IQ-TREE. A guide tree on the left marks (with a gray rectangle), the region of the acanthomorph tree represented in the figure. All nodes have 100% bootstrap support. Orders, families or genera for taxa are listed to the right of the vertical black bars. Any tips left unassigned to a higher taxon are monotypic or monogeneric taxa.

**Supplementary Fig. 3:**
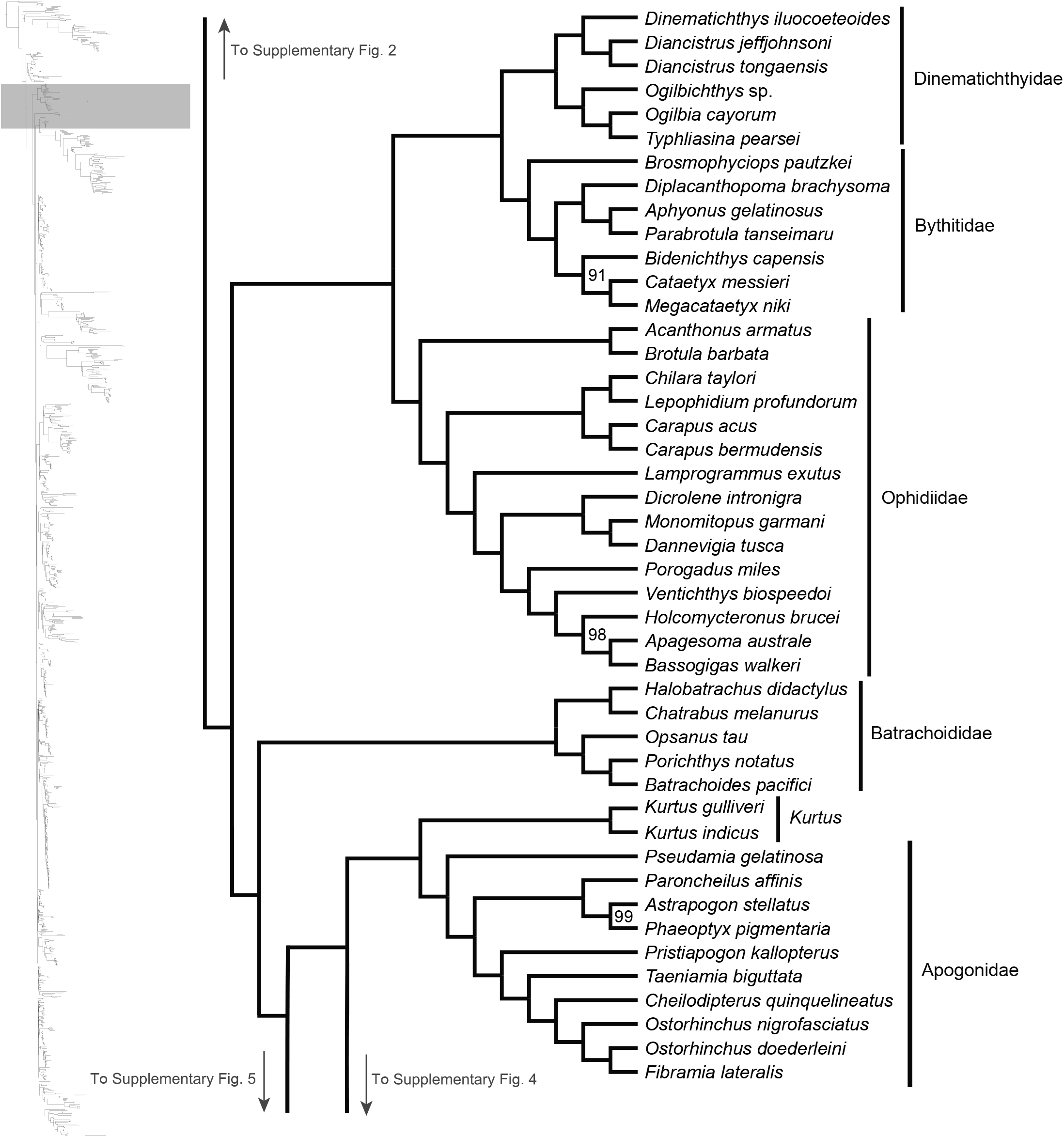
Maximum likelihood phylogeny inferred in IQ-TREE. A guide tree on the left marks (with a gray rectangle), the region of the acanthomorph tree represented in the figure. Numbers at nodes reflect bootstrap support values. All nodes without a numerical annotation have 100% bootstrap support. Orders, families or genera for taxa are listed to the right of the vertical black bars.

**Supplementary Fig. 4:**
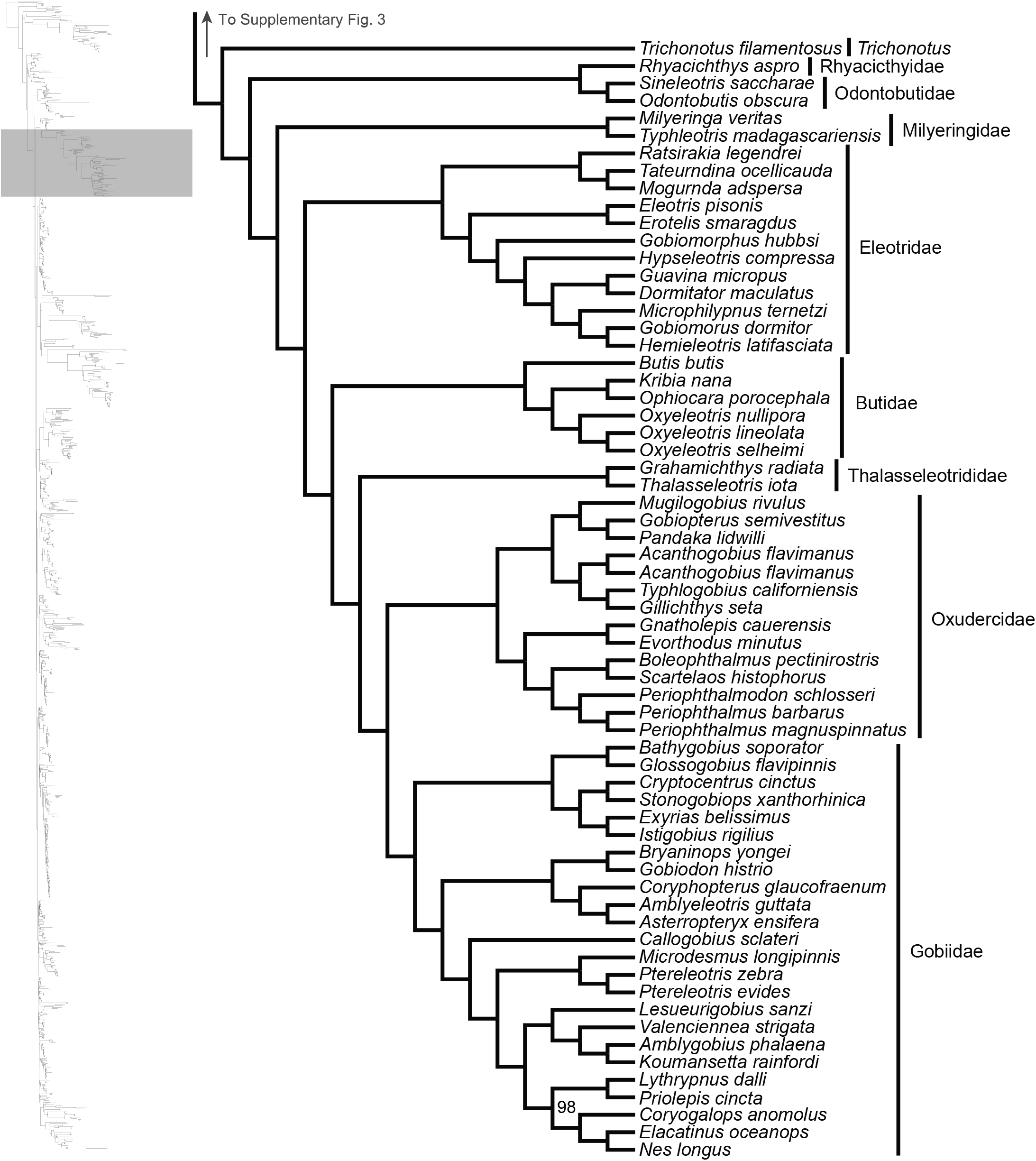
Maximum likelihood phylogeny inferred in IQ-TREE. A guide tree on the left marks (with a gray rectangle), the region of the acanthomorph tree represented in the figure. Numbers at nodes reflect bootstrap support values. All nodes without a numerical annotation have 100% bootstrap support. Orders, families or genera for taxa are listed to the right of the vertical black bars.

**Supplementary Fig. 5:**
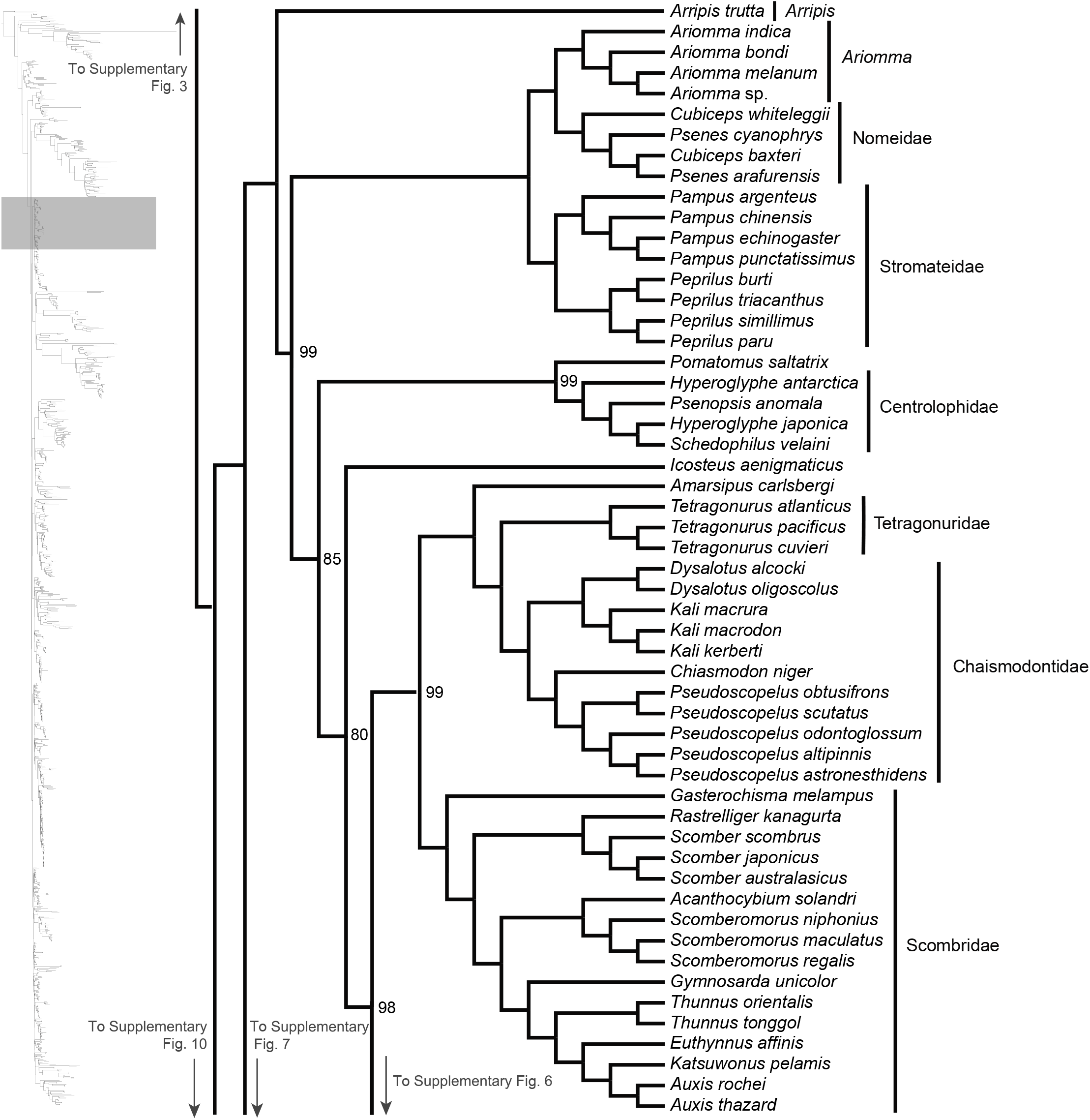
Maximum likelihood phylogeny inferred in IQ-TREE. A guide tree on the left marks (with a gray rectangle), the region of the acanthomorph tree represented in the figure. Numbers at nodes reflect bootstrap support values. All nodes without a numerical annotation have 100% bootstrap support. Orders, families or genera for taxa are listed to the right of the vertical black bars. Any tips left unassigned to a higher taxon are monotypic or monogeneric taxa.

**Supplementary Fig. 6:**
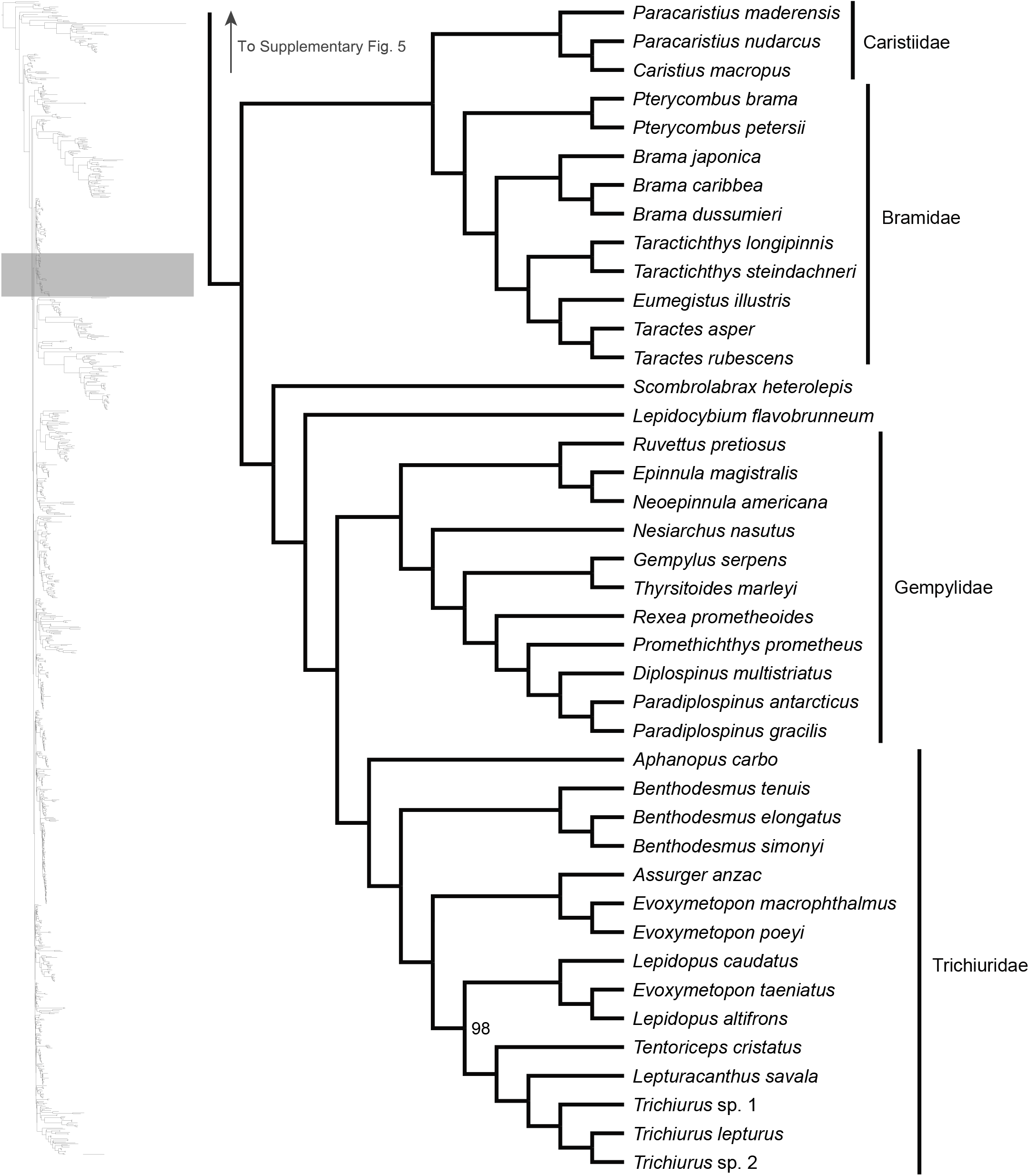
Maximum likelihood phylogeny inferred in IQ-TREE. A guide tree on the left marks (with a gray rectangle), the region of the acanthomorph tree represented in the figure. Numbers at nodes reflect bootstrap support values. All nodes without a numerical annotation have 100% bootstrap support. Orders, families or genera for taxa are listed to the right of the vertical black bars. Any tips left unassigned to a higher taxon are monotypic or monogeneric taxa.

**Supplementary Fig. 7:**
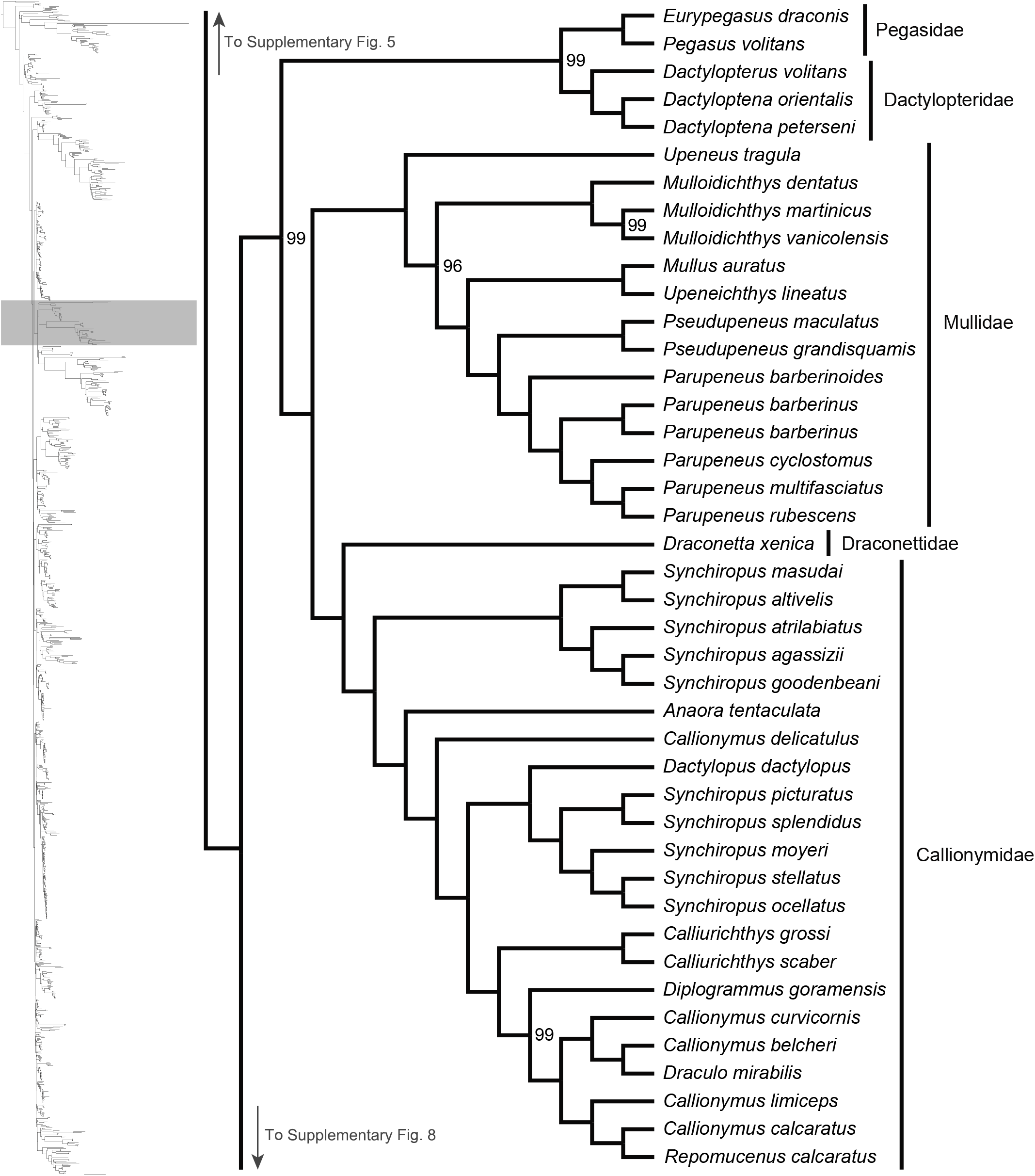
Maximum likelihood phylogeny inferred in IQ-TREE. A guide tree on the left marks (with a gray rectangle), the region of the acanthomorph tree represented in the figure. Numbers at nodes reflect bootstrap support values. All nodes without a numerical annotation have 100% bootstrap support. Orders, families or genera for taxa are listed to the right of the vertical black bars.

**Supplementary Fig. 8:**
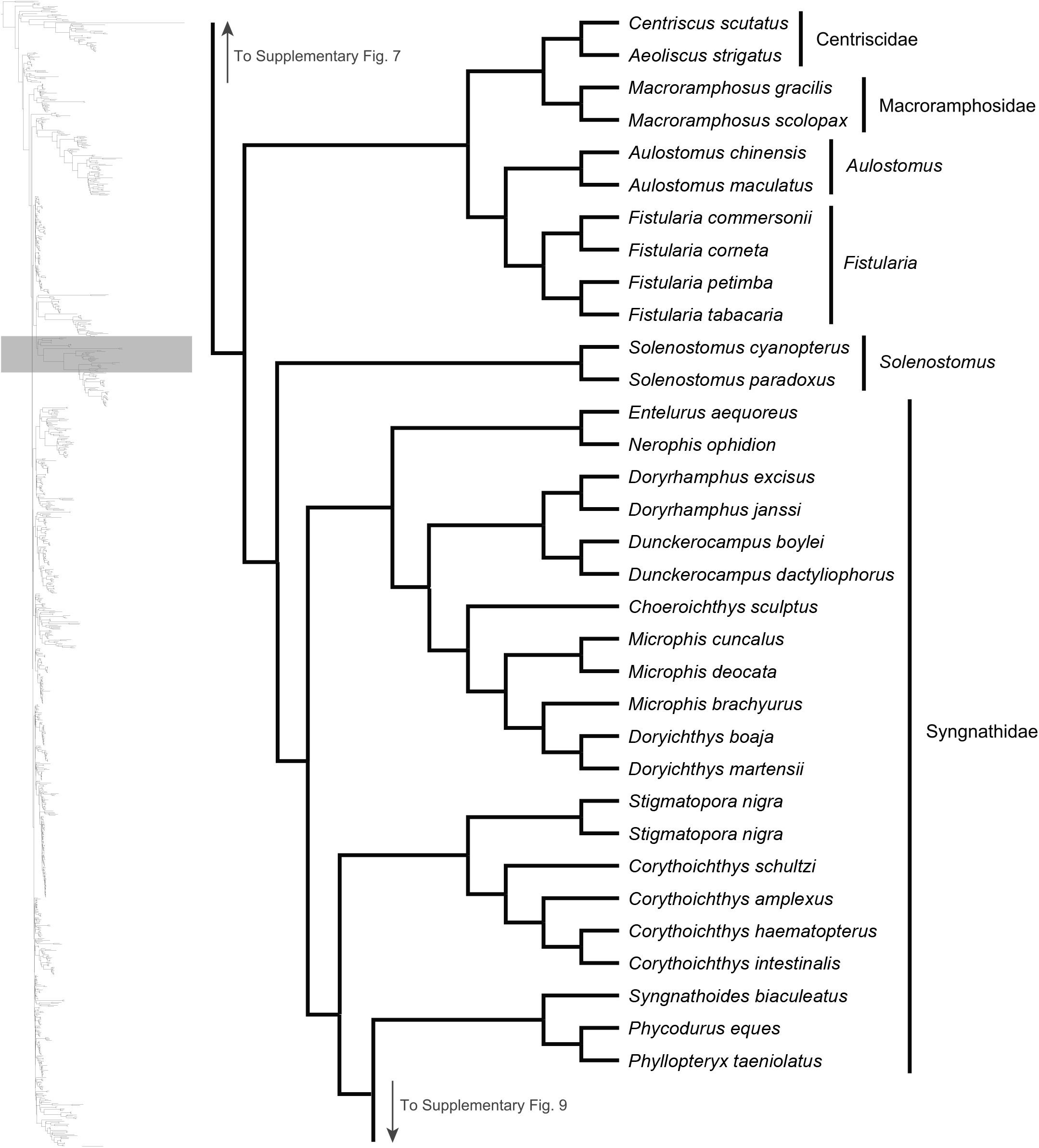
Maximum likelihood phylogeny inferred in IQ-TREE. A guide tree on the left marks (with a gray rectangle), the region of the acanthomorph tree represented in the figure. All nodes have 100% bootstrap support. Orders, families or genera for taxa are listed to the right of the vertical black bars.

**Supplementary Fig. 9:**
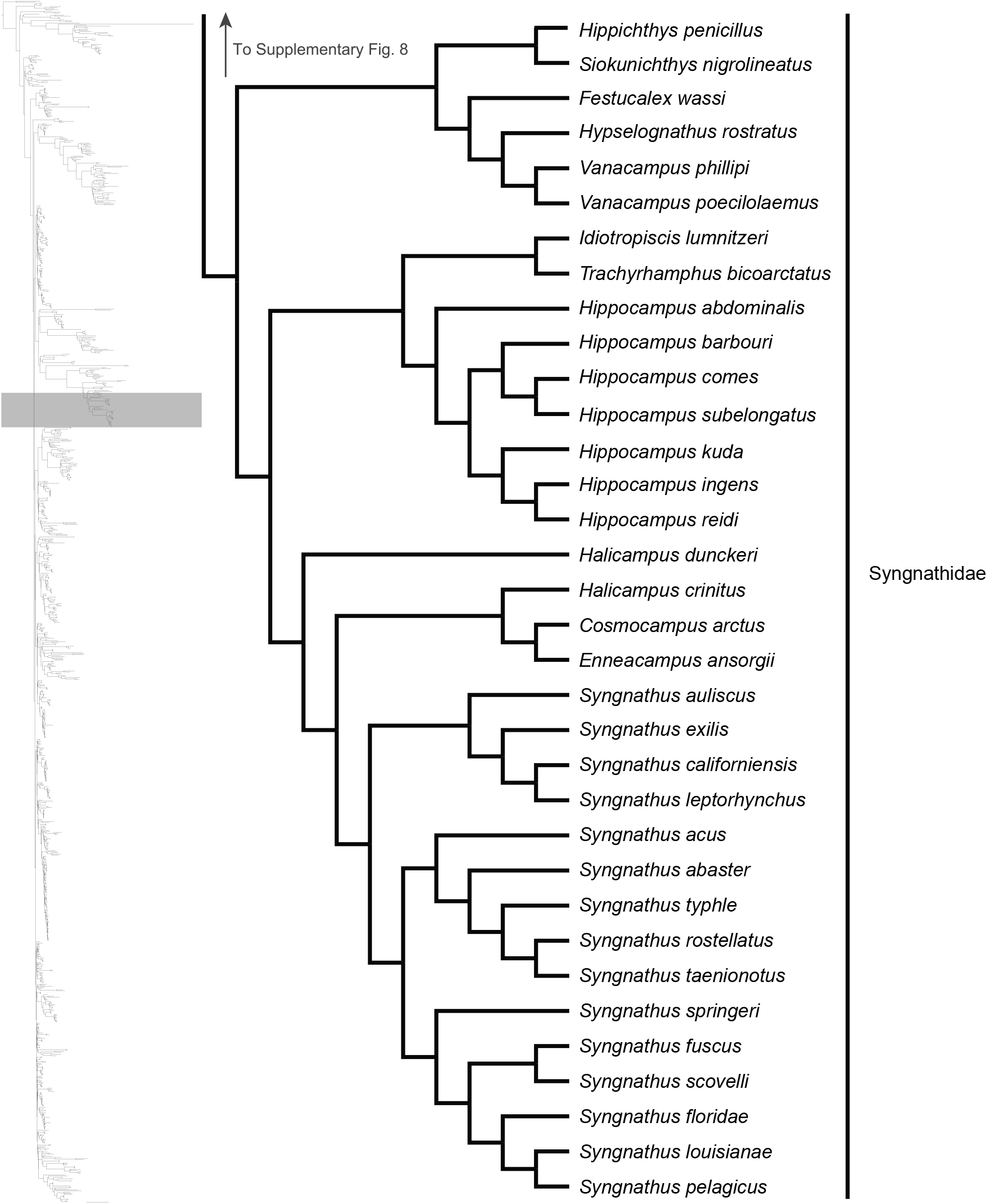
Maximum likelihood phylogeny inferred in IQ-TREE. A guide tree on the left marks (with a gray rectangle), the region of the acanthomorph tree represented in the figure. All nodes have 100% bootstrap support. Orders, families or genera for taxa are listed to the right of the vertical black bars.

**Supplementary Fig. 10:**
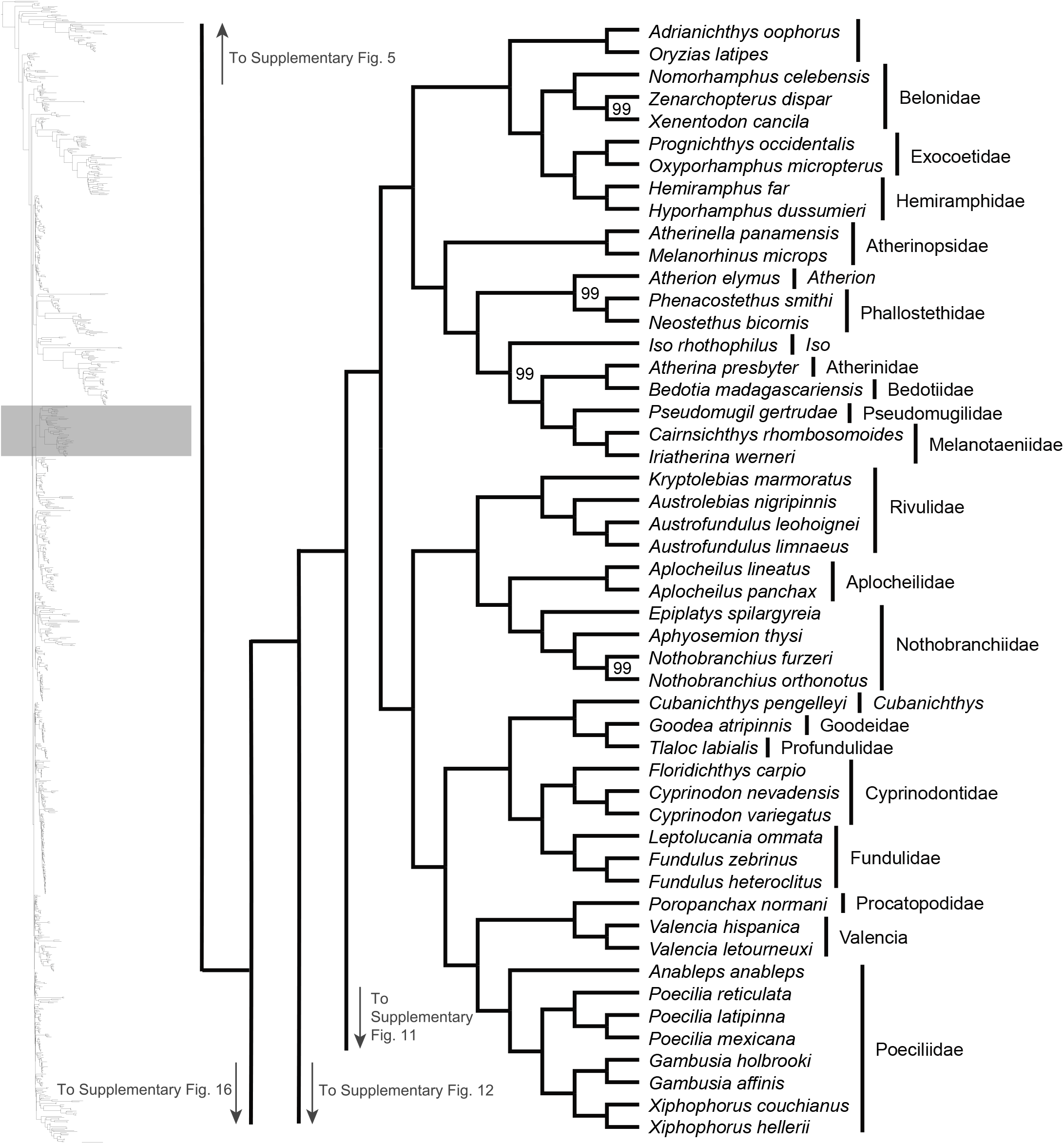
Maximum likelihood phylogeny inferred in IQ-TREE. A guide tree on the left marks (with a gray rectangle), the region of the acanthomorph tree represented in the figure. Numbers at nodes reflect bootstrap support values. All nodes without a numerical annotation have 100% bootstrap support. Orders, families or genera for taxa are listed to the right of the vertical black bars.

**Supplementary Fig. 11:**
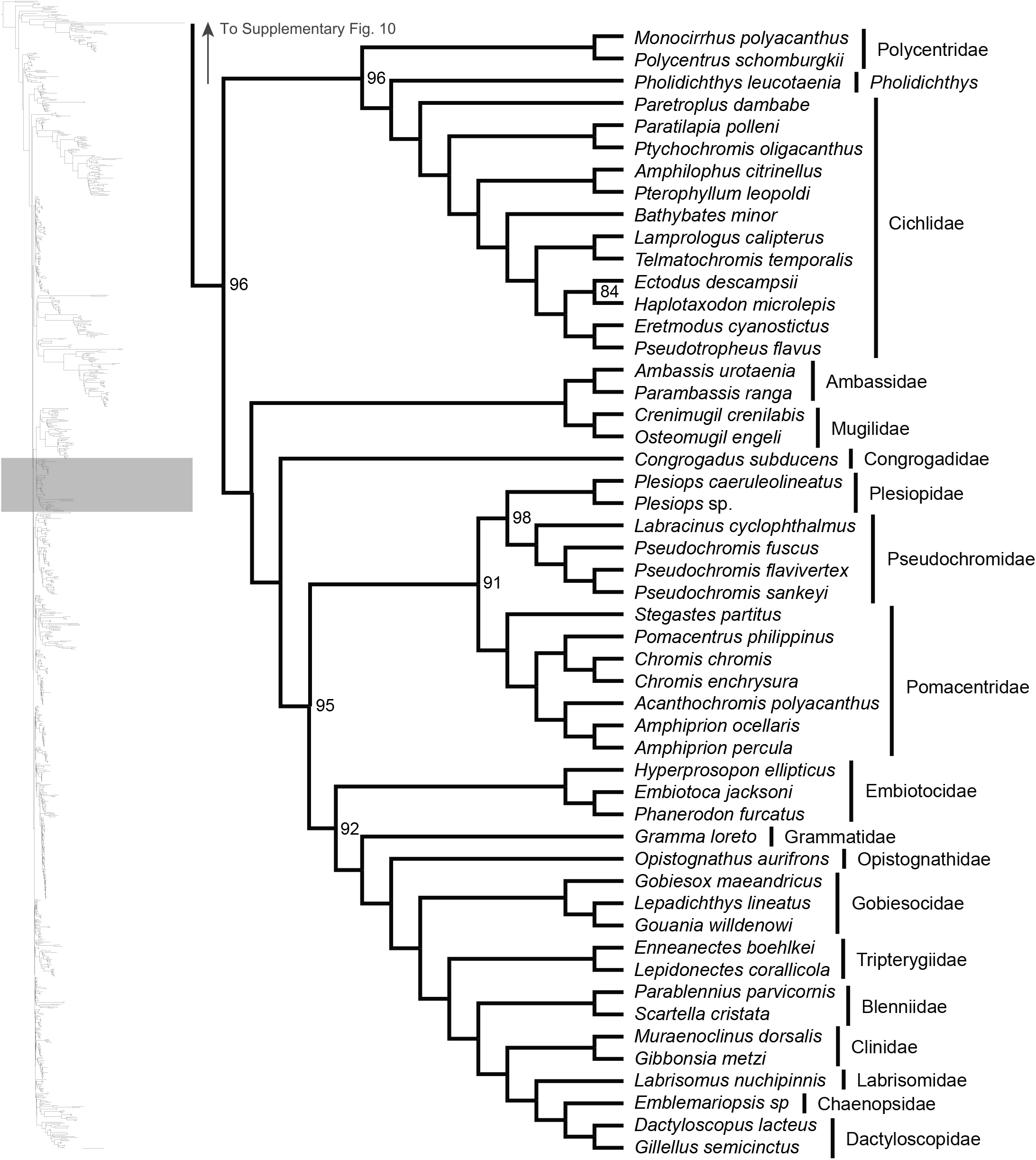
Maximum likelihood phylogeny inferred in IQ-TREE. A guide tree on the left marks (with a gray rectangle), the region of the acanthomorph tree represented in the figure. Numbers at nodes reflect bootstrap support values. All nodes without a numerical annotation have 100% bootstrap support. Orders, families or genera for taxa are listed to the right of the vertical black bars.

**Supplementary Fig. 12:**
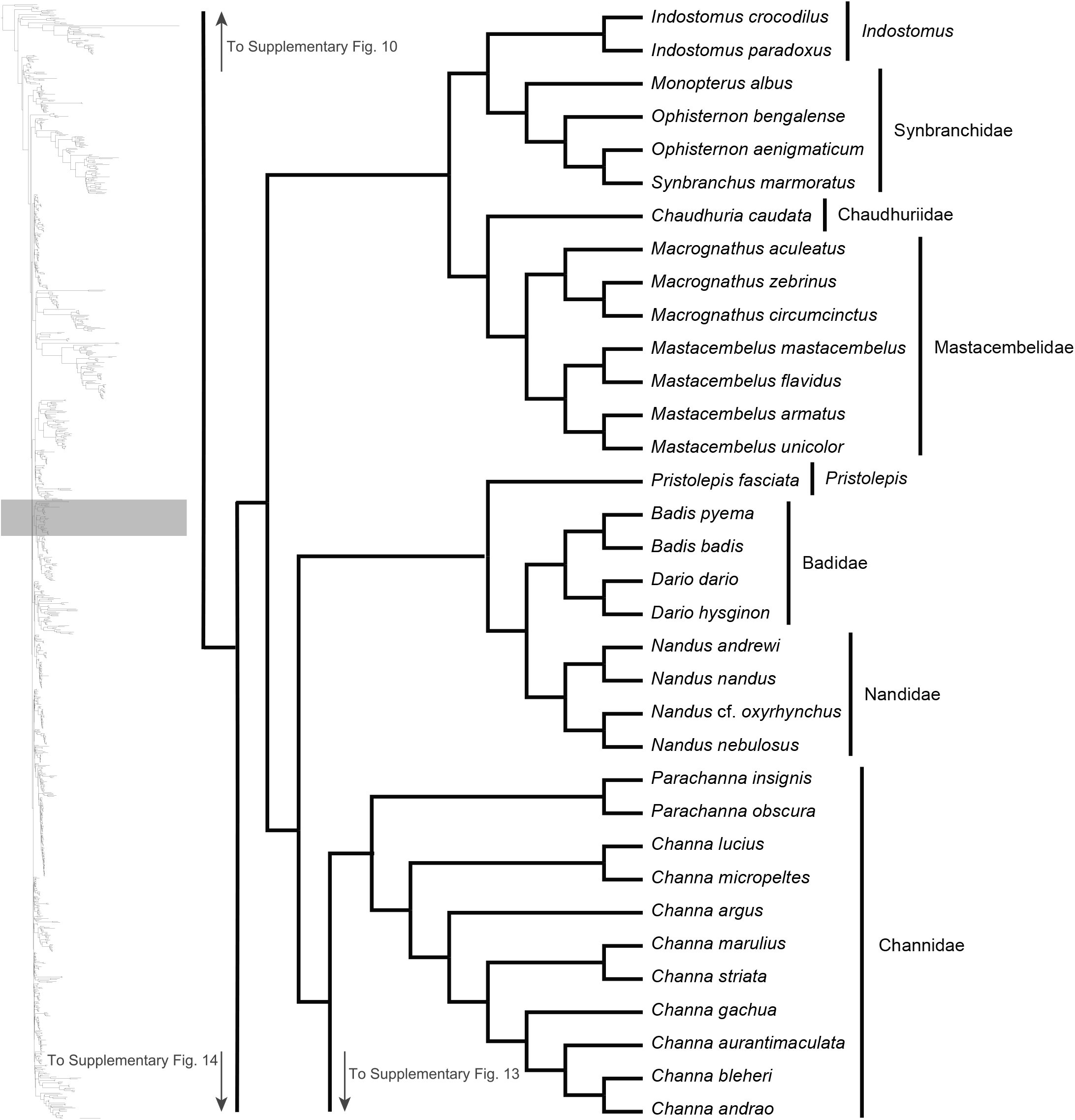
Maximum likelihood phylogeny inferred in IQ-TREE. A guide tree on the left marks (with a gray rectangle), the region of the acanthomorph tree represented in the figure. All nodes have 100% bootstrap support. Orders, families or genera for taxa are listed to the right of the vertical black bars.

**Supplementary Fig. 13:**
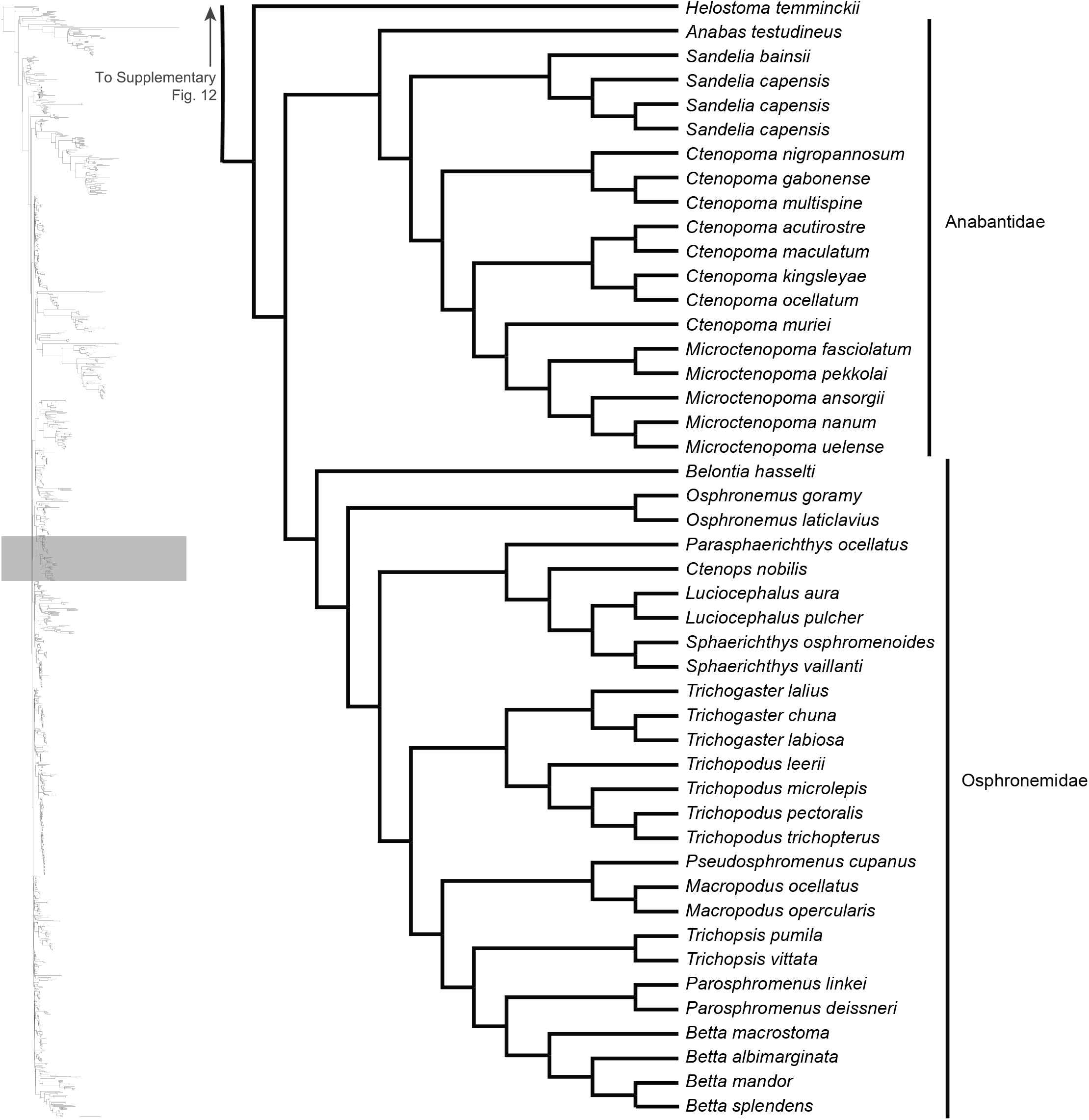
Maximum likelihood phylogeny inferred in IQ-TREE. A guide tree on the left marks (with a gray rectangle), the region of the acanthomorph tree represented in the figure. All nodes have 100% bootstrap support. Orders, families or genera for taxa are listed to the right of the vertical black bars. Any tips left unassigned to a higher taxon are monotypic or monogeneric taxa.

**Supplementary Fig. 14:**
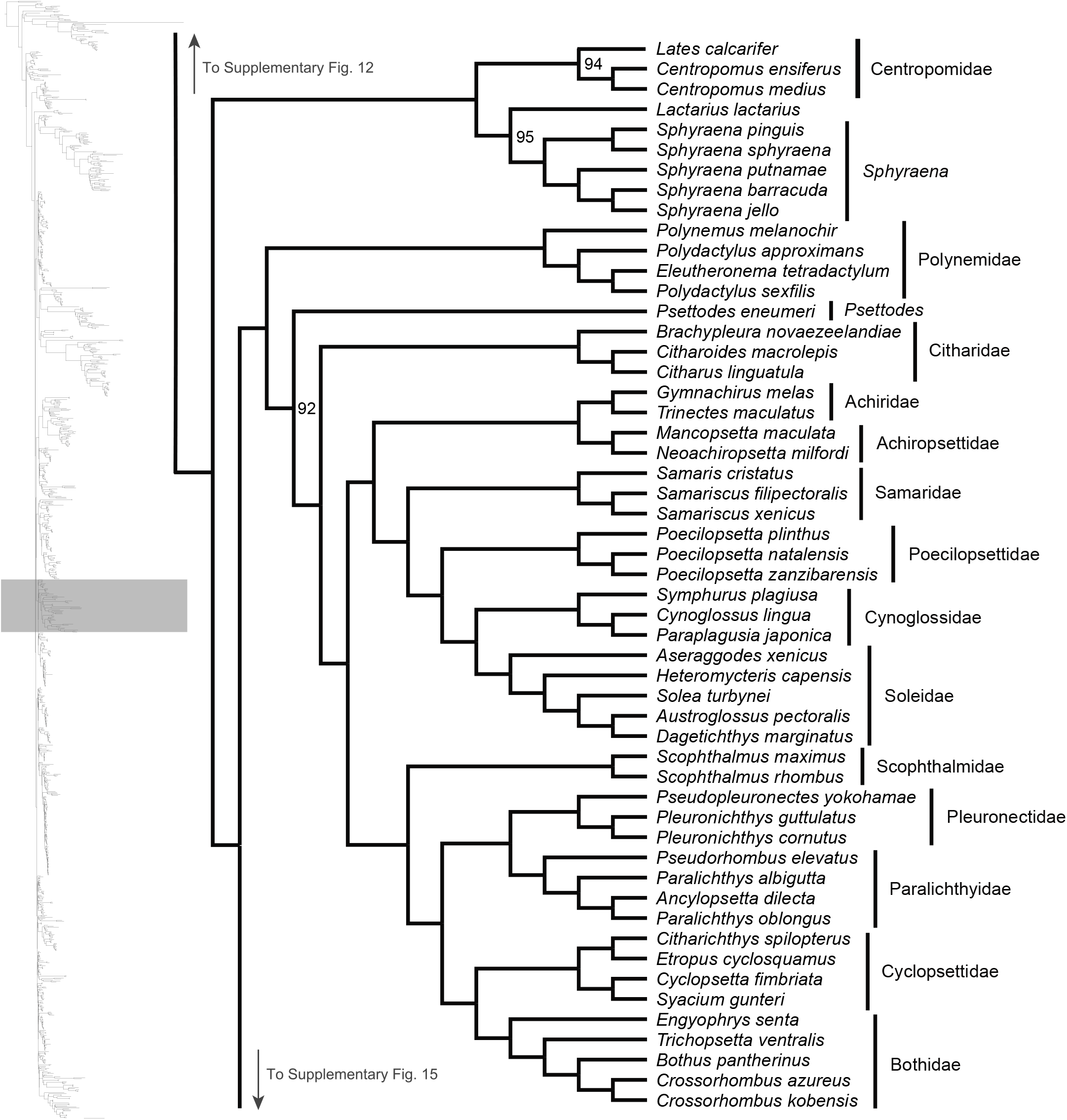
Maximum likelihood phylogeny inferred in IQ-TREE. A guide tree on the left marks (with a gray rectangle), the region of the acanthomorph tree represented in the figure. Numbers at nodes reflect bootstrap support values. All nodes without a numerical annotation have 100% bootstrap support. Orders, families or genera for taxa are listed to the right of the vertical black bars. Any tips left unassigned to a higher taxon are monotypic or monogeneric taxa.

**Supplementary Fig. 15:**
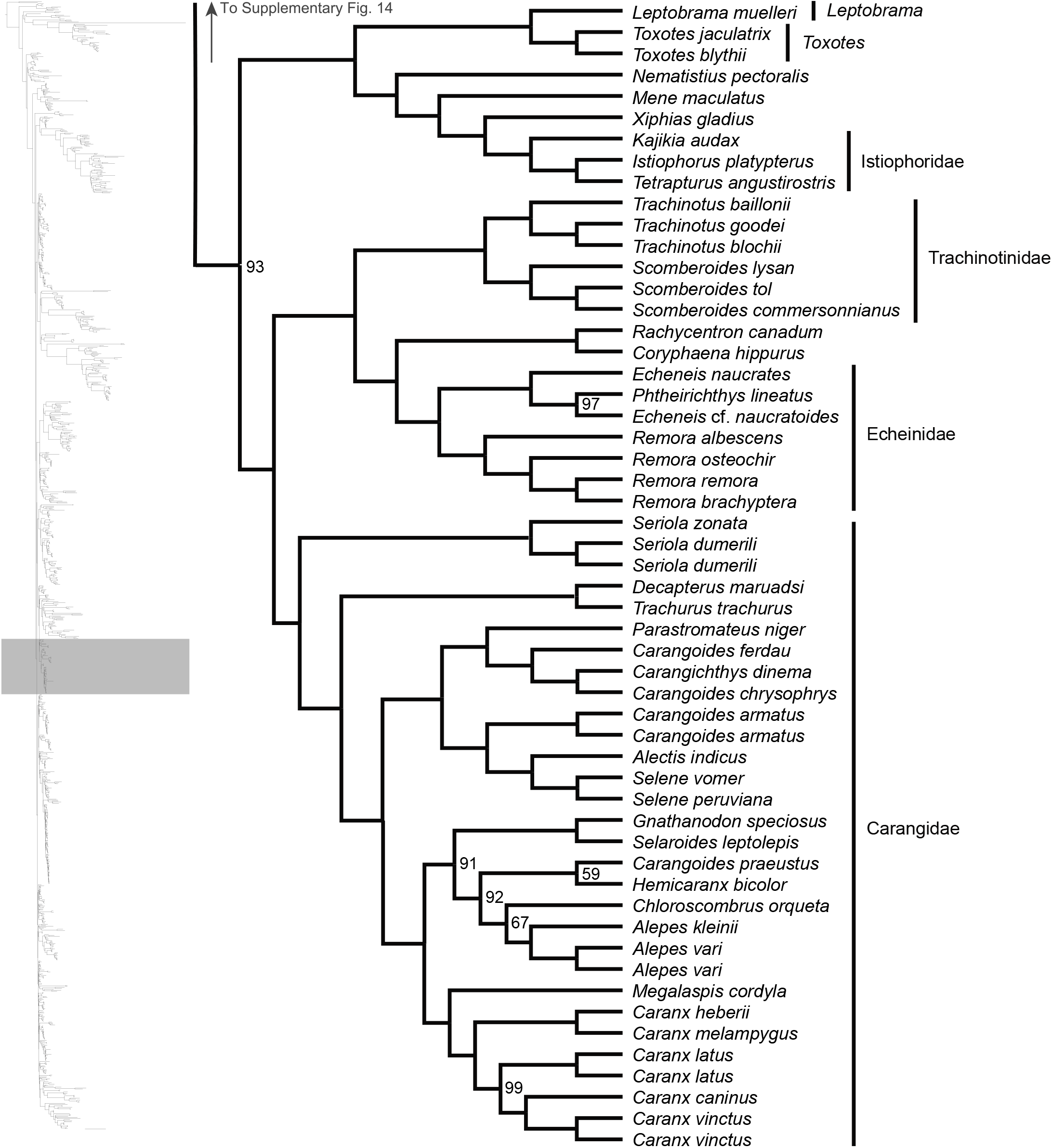
Maximum likelihood phylogeny inferred in IQ-TREE. A guide tree on the left marks (with a gray rectangle), the region of the acanthomorph tree represented in the figure. Numbers at nodes reflect bootstrap support values. All nodes without a numerical annotation have 100% bootstrap support. Orders, families or genera for taxa are listed to the right of the vertical black bars. Any tips left unassigned to a higher taxon are monotypic or monogeneric taxa.

**Supplementary Fig. 16:**
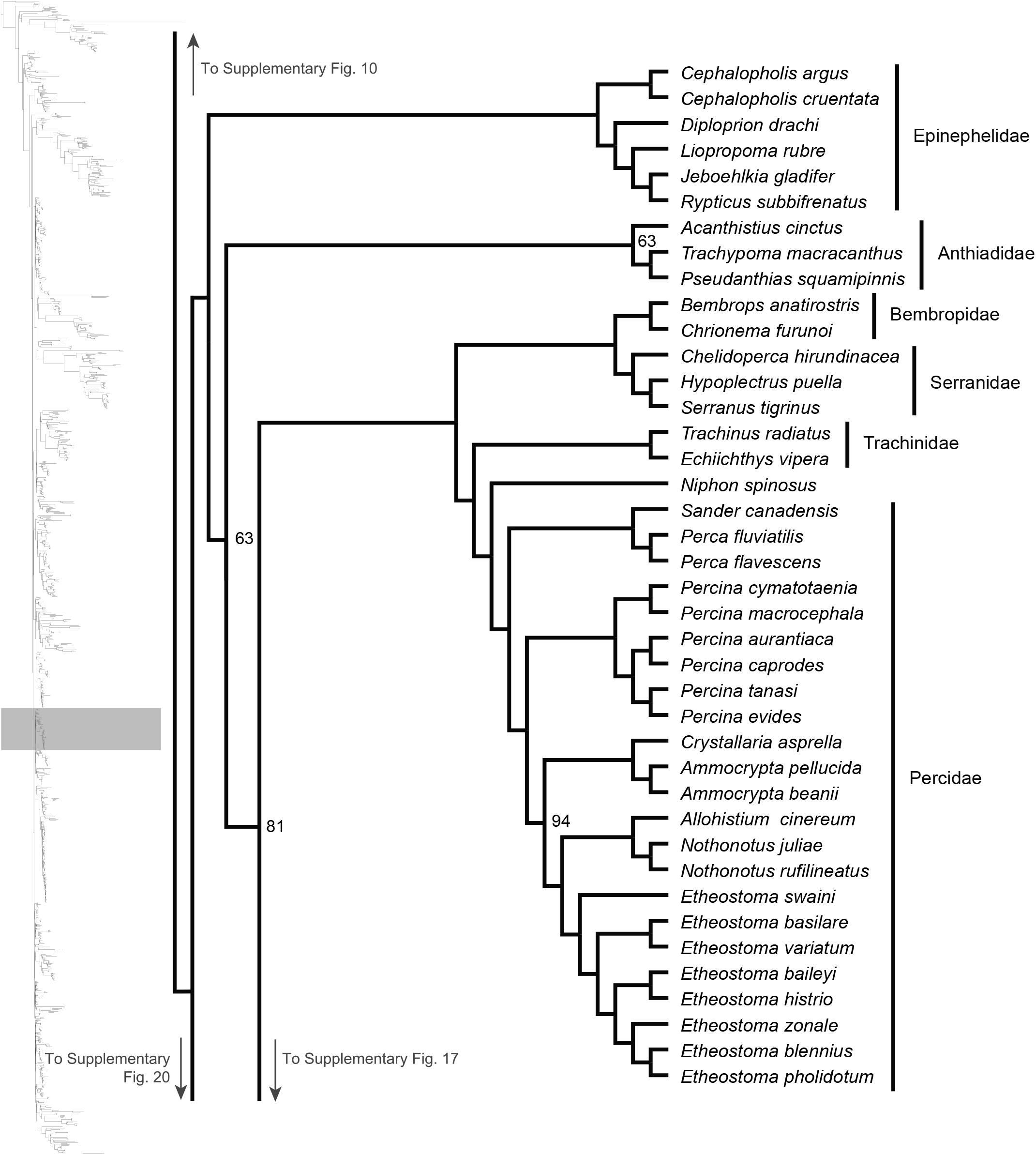
Maximum likelihood phylogeny inferred in IQ-TREE. A guide tree on the left marks (with a gray rectangle), the region of the acanthomorph tree represented in the figure. Numbers at nodes reflect bootstrap support values. All nodes without a numerical annotation have 100% bootstrap support. Orders, families or genera for taxa are listed to the right of the vertical black bars. Any tips left unassigned to a higher taxon are monotypic or monogeneric taxa.

**Supplementary Fig. 17:**
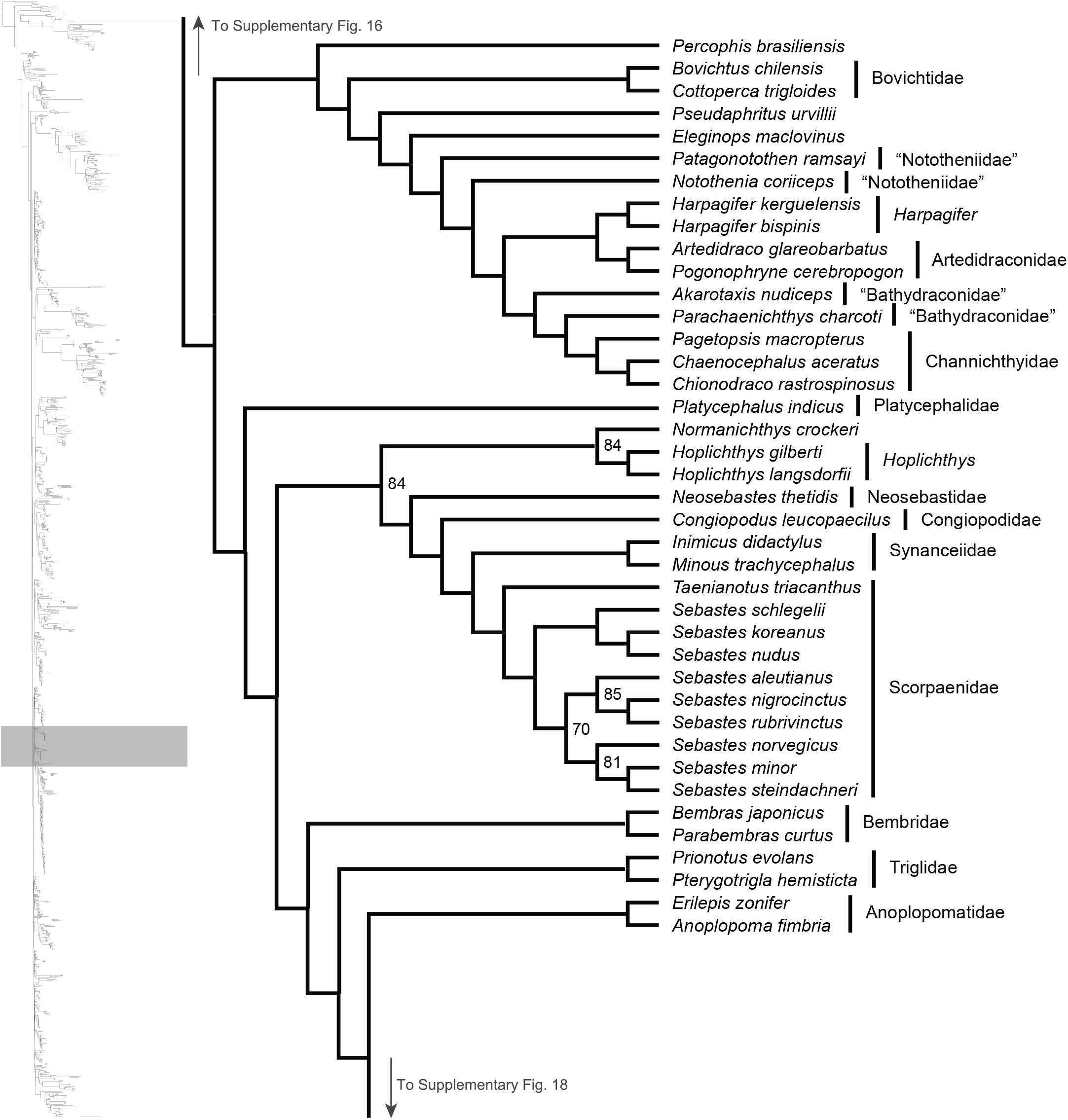
Maximum likelihood phylogeny inferred in IQ-TREE. A guide tree on the left marks (with a gray rectangle), the region of the acanthomorph tree represented in the figure. Numbers at nodes reflect bootstrap support values. All nodes without a numerical annotation have 100% bootstrap support. Orders, families or genera for taxa are listed to the right of the vertical black bars. Any tips left unassigned to a higher taxon are monotypic or monogeneric taxa.

**Supplementary Fig. 18:**
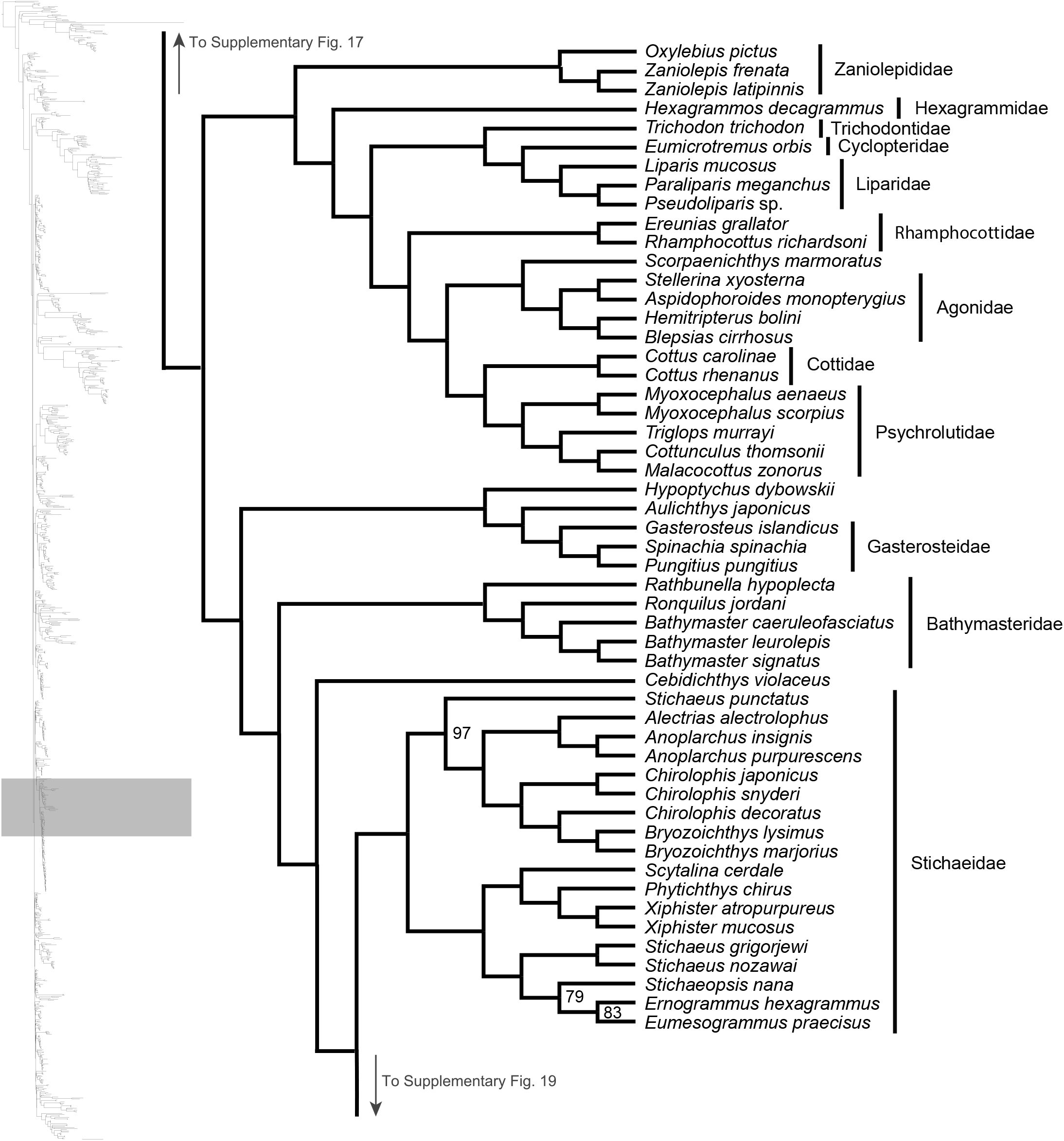
Maximum likelihood phylogeny inferred in IQ-TREE. A guide tree on the left marks (with a gray rectangle), the region of the acanthomorph tree represented in the figure. Numbers at nodes reflect bootstrap support values. All nodes without a numerical annotation have 100% bootstrap support. Orders, families or genera for taxa are listed to the right of the vertical black bars. Any tips left unassigned to a higher taxon are monotypic or monogeneric taxa.

**Supplementary Fig. 19:**
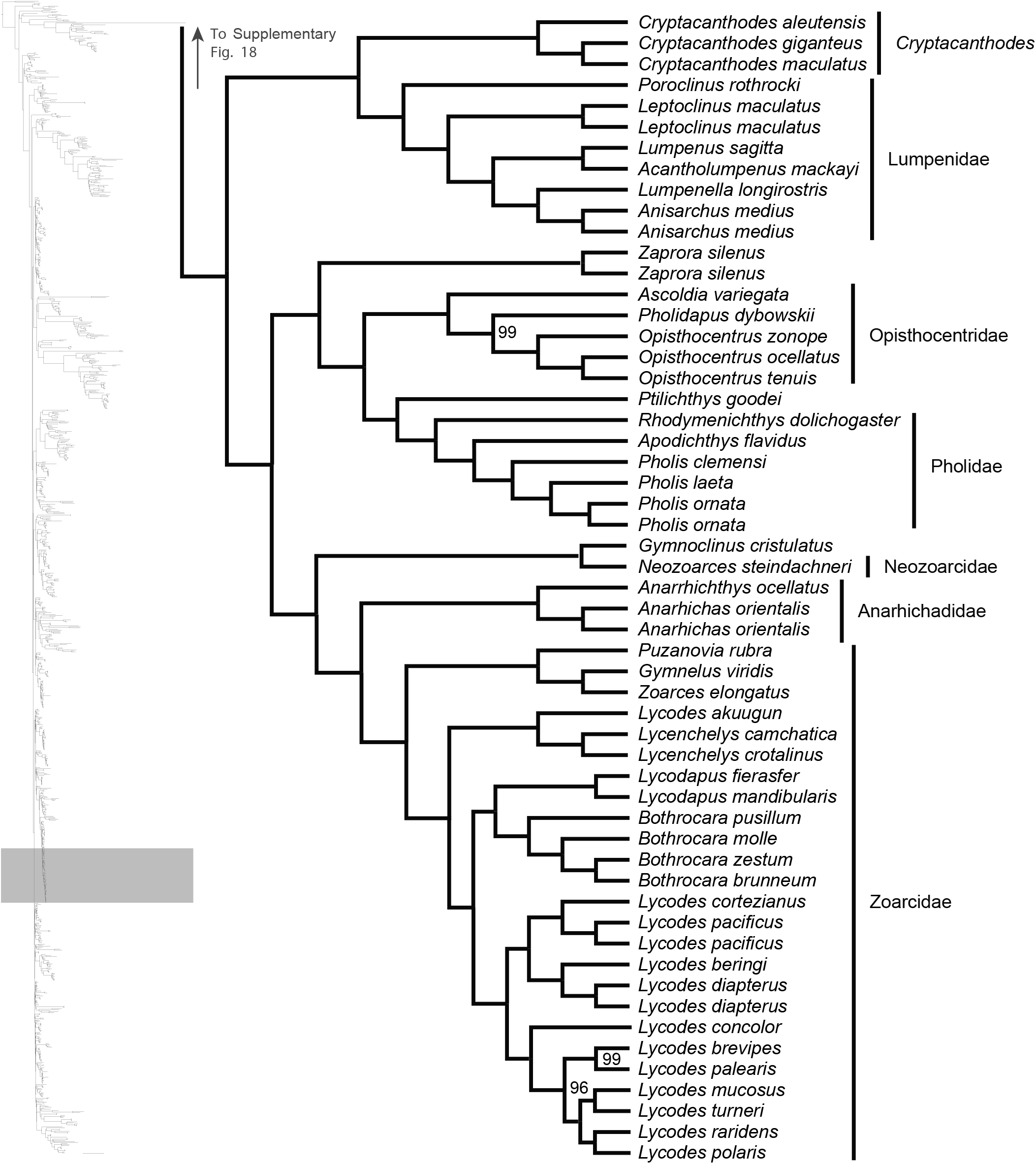
Maximum likelihood phylogeny inferred in IQ-TREE. A guide tree on the left marks (with a gray rectangle), the region of the acanthomorph tree represented in the figure. Numbers at nodes reflect bootstrap support values. All nodes without a numerical annotation have 100% bootstrap support. Orders, families or genera for taxa are listed to the right of the vertical black bars. Any tips left unassigned to a higher taxon are monotypic or monogeneric taxa.

**Supplementary Fig. 20:**
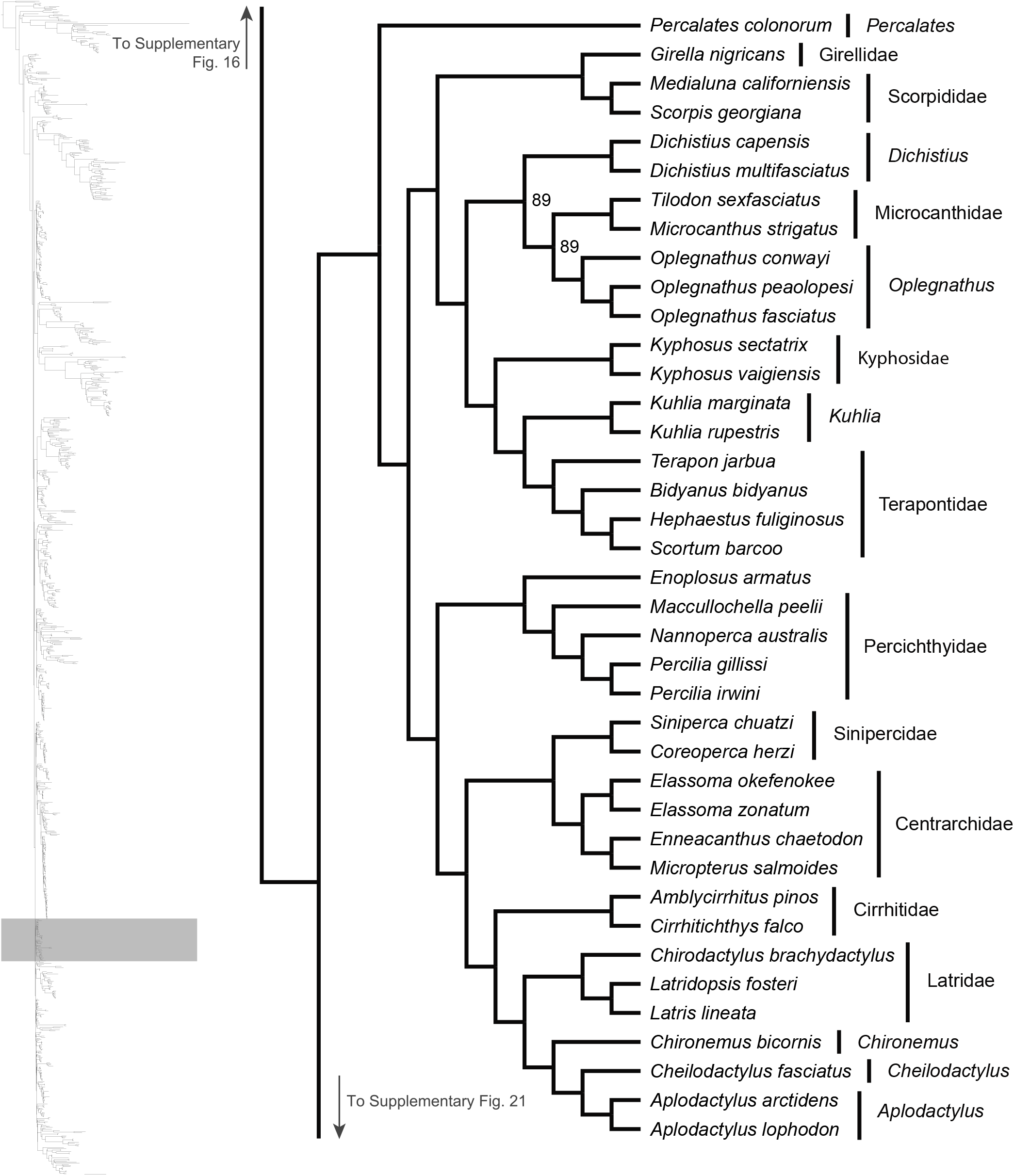
Maximum likelihood phylogeny inferred in IQ-TREE. A guide tree on the left marks (with a gray rectangle), the region of the acanthomorph tree represented in the figure. Numbers at nodes reflect bootstrap support values. All nodes without a numerical annotation have 100% bootstrap support. Orders, families or genera for taxa are listed to the right of the vertical black bars. Any tips left unassigned to a higher taxon are monotypic or monogeneric taxa.

**Supplementary Fig. 21:**
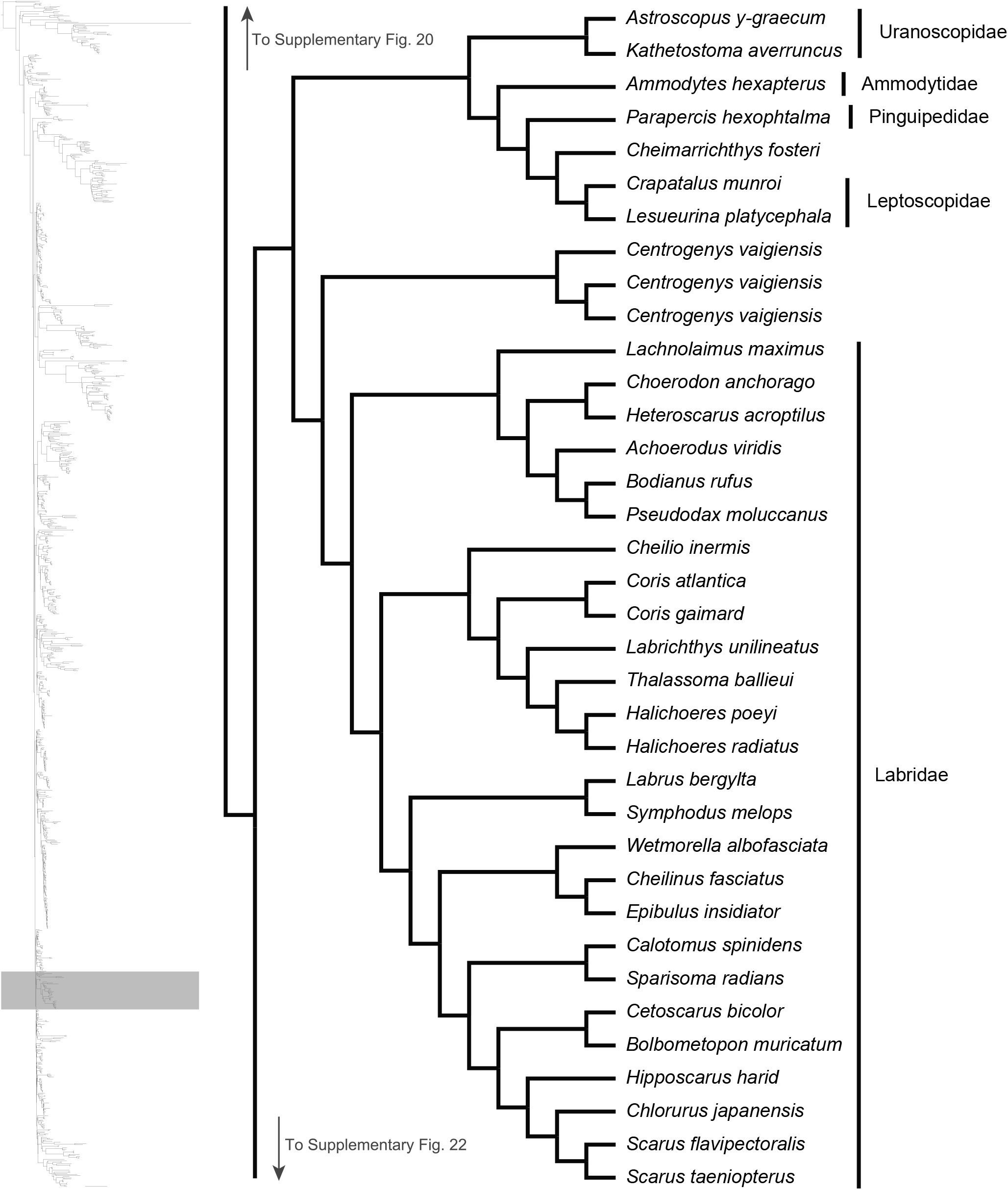
Maximum likelihood phylogeny inferred in IQ-TREE. A guide tree on the left marks (with a gray rectangle), the region of the acanthomorph tree represented in the figure. All nodes have 100% bootstrap support. Orders, families or genera for taxa are listed to the right of the vertical black bars. Any tips left unassigned to a higher taxon are monotypic or monogeneric taxa.

**Supplementary Fig. 22:**
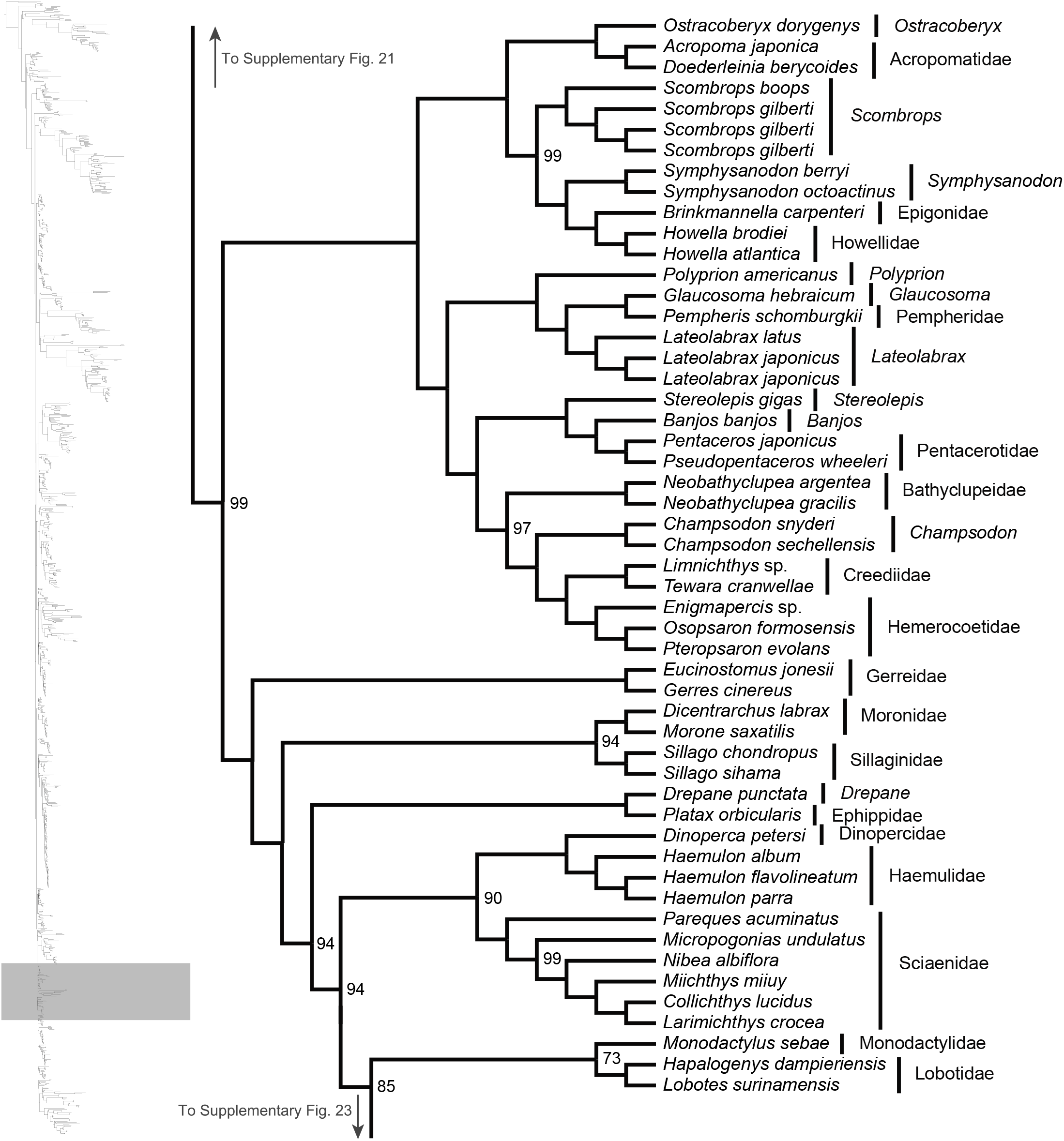
Maximum likelihood phylogeny inferred in IQ-TREE. A guide tree on the left marks (with a gray rectangle), the region of the acanthomorph tree represented in the figure. Numbers at nodes reflect bootstrap support values. All nodes without a numerical annotation have 100% bootstrap support. Orders, families or genera for taxa are listed to the right of the vertical black bars.

**Supplementary Fig. 23:**
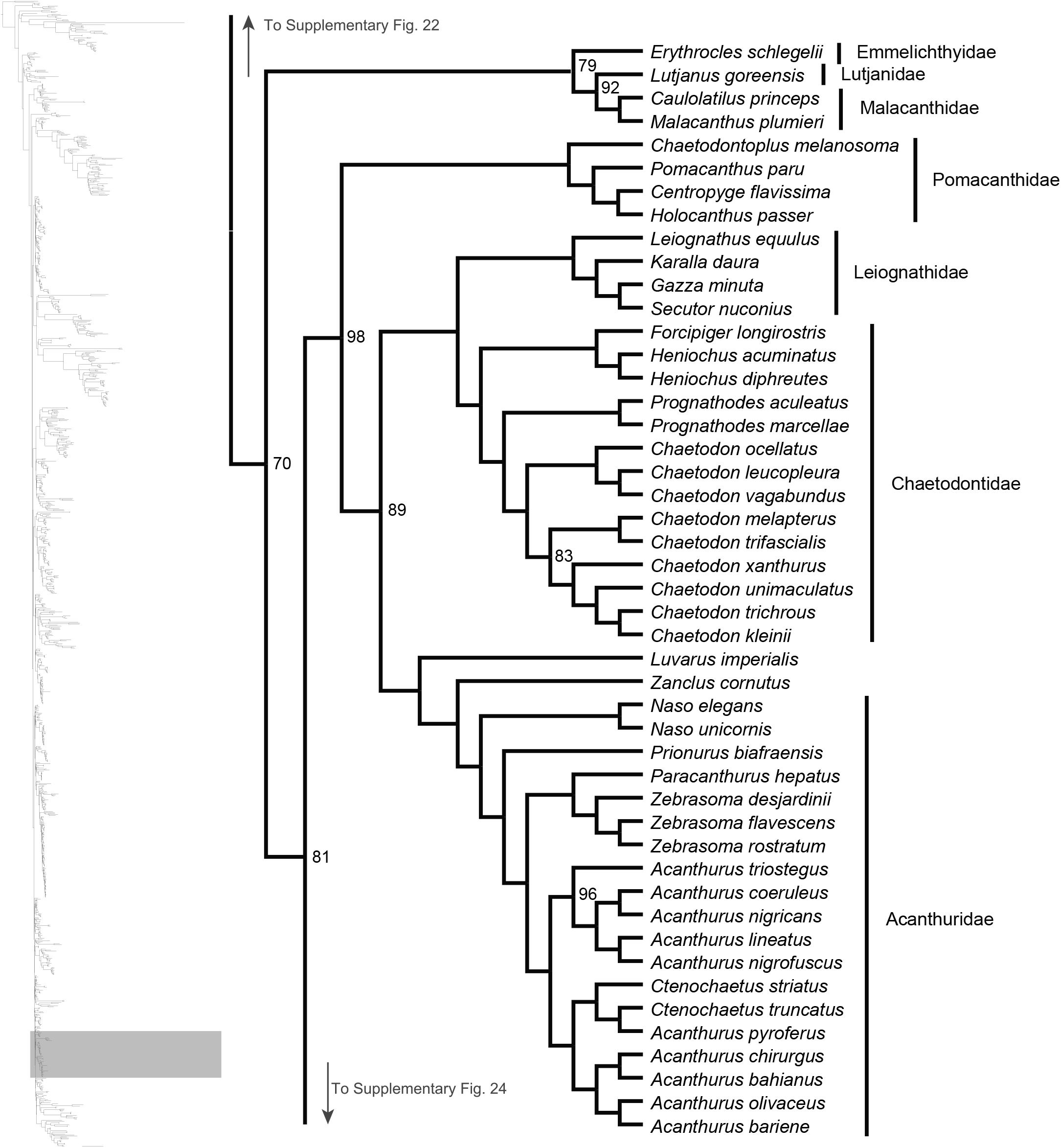
Maximum likelihood phylogeny inferred in IQ-TREE. A guide tree on the left marks (with a gray rectangle), the region of the acanthomorph tree represented in the figure. Numbers at nodes reflect bootstrap support values. All nodes without a numerical annotation have 100% bootstrap support. Orders, families or genera for taxa are listed to the right of the vertical black bars. Any tips left unassigned to a higher taxon are monotypic or monogeneric taxa.

**Supplementary Fig. 24:**
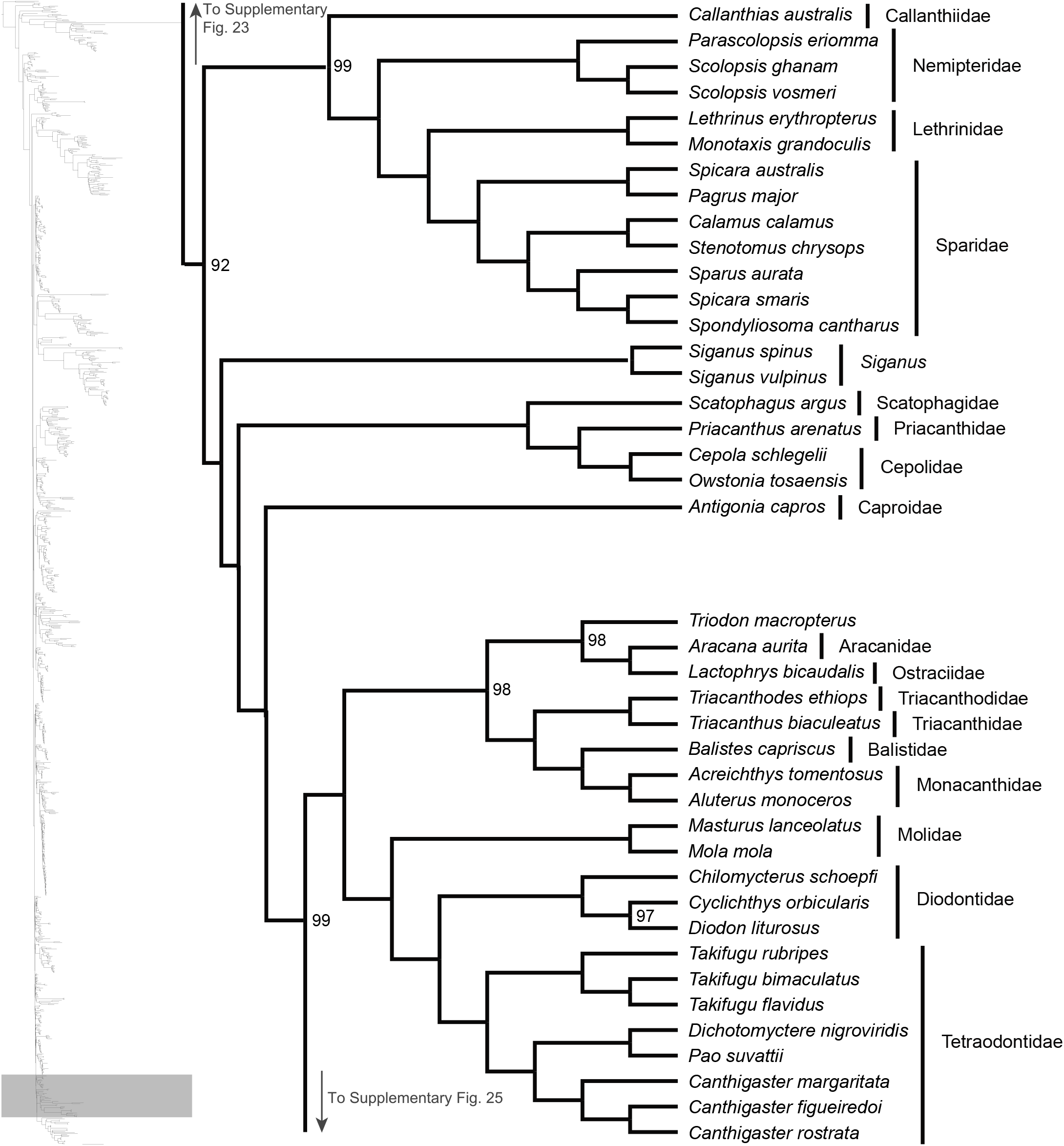
Maximum likelihood phylogeny inferred in IQ-TREE. A guide tree on the left marks (with a gray rectangle), the region of the acanthomorph tree represented in the figure. Numbers at nodes reflect bootstrap support values. All nodes without a numerical annotation have 100% bootstrap support. Orders, families or genera for taxa are listed to the right of the vertical black bars. Any tips left unassigned to a higher taxon are monospecific taxa.

**Supplementary Fig. 25:**
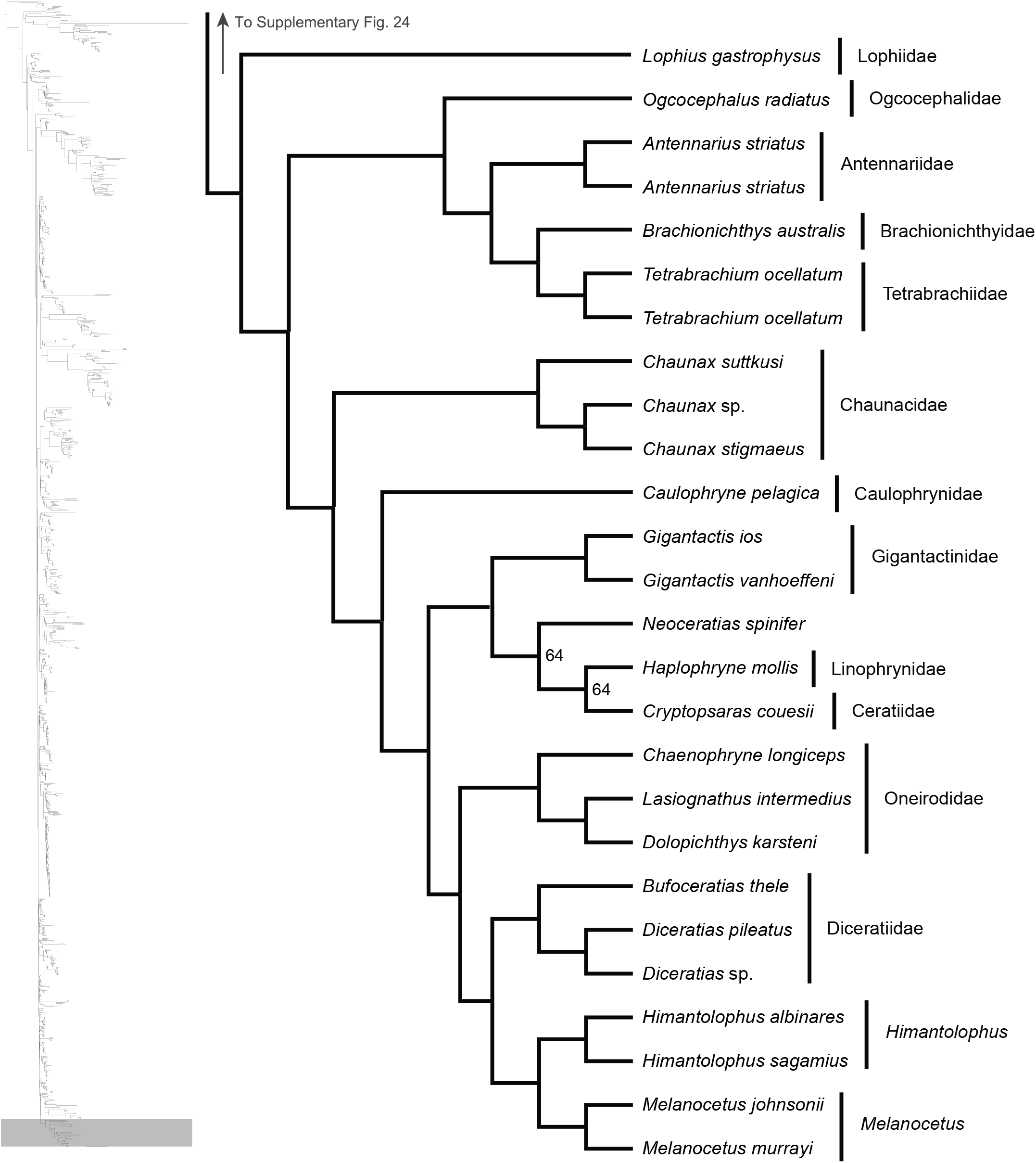
Maximum likelihood phylogeny inferred in IQ-TREE. A guide tree on the left marks (with a gray rectangle), the region of the acanthomorph tree represented in the figure. Numbers at nodes reflect bootstrap support values. All nodes without a numerical annotation have 100% bootstrap support. Orders, families or genera for taxa are listed to the right of the vertical black bars. Any tips left unassigned to a higher taxon are monotypic or monogeneric taxa.

**Supplementary Fig. 26:**
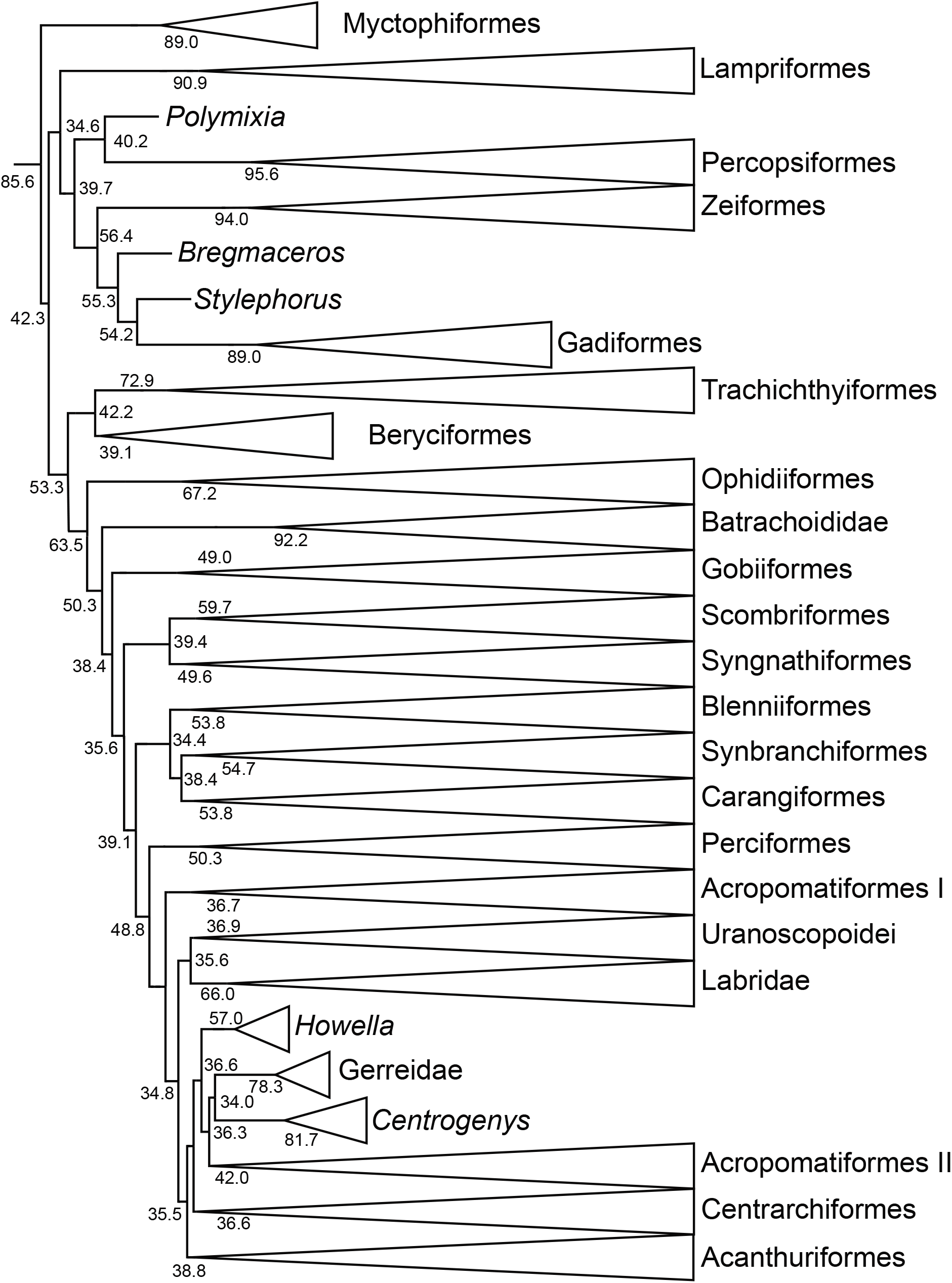
ASTRAL-III inferred species tree. Collapsed species tree inferred under the multi-species coalescent model. Local posterior probability values at nodes do not measure support for bipartitions, but rather are a function of the frequencies of the shown quartet topologies among all gene trees.

**Supplementary Fig. 27:**
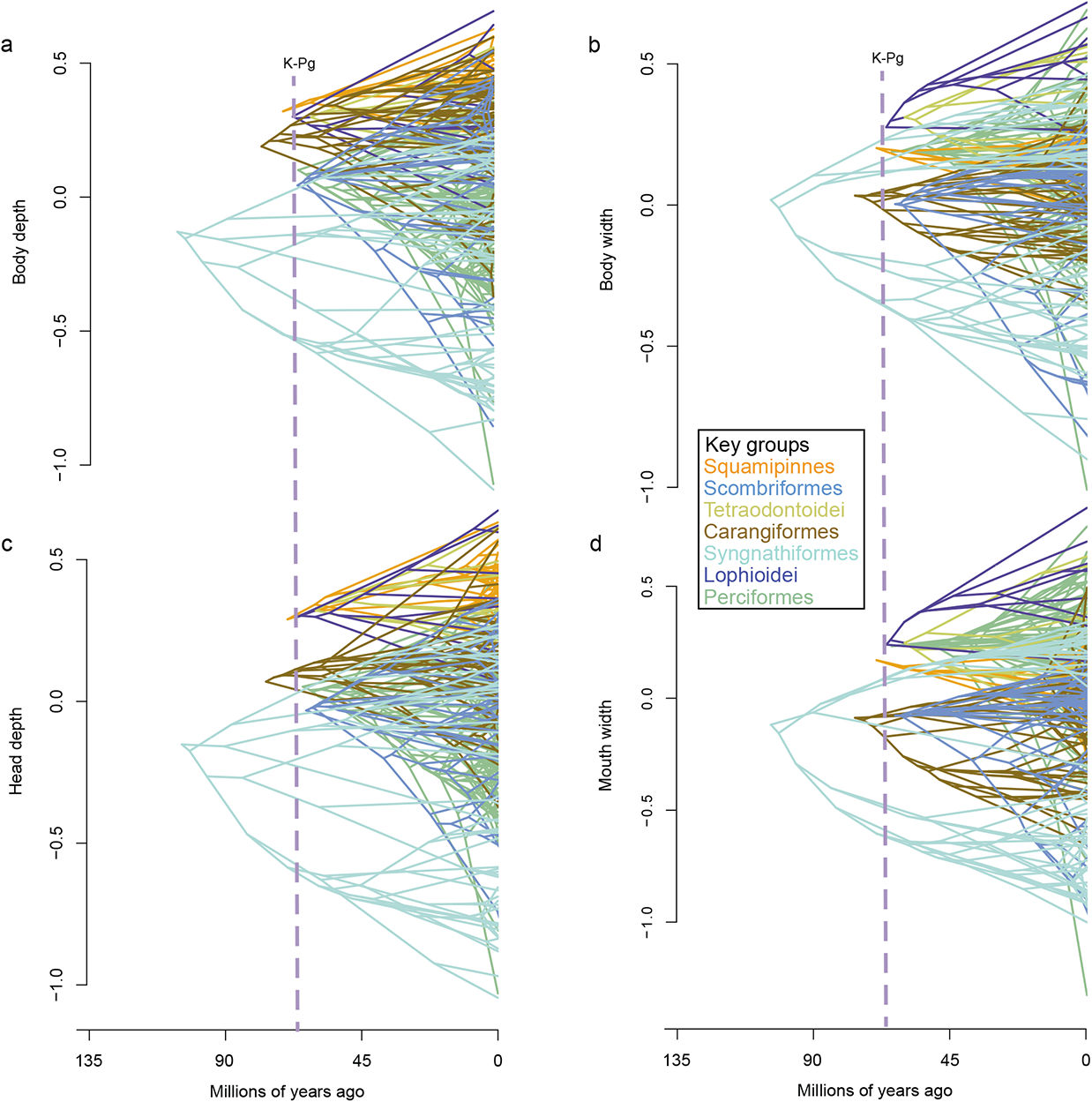
Phenograms depicting the evolutionary history of four phenotypic traits (body depth and width, and head depth and width) across seven major lineages that arose around the K-Pg. The vertical dashed line marks the K-Pg boundary.

**Supplementary Fig. 28:**
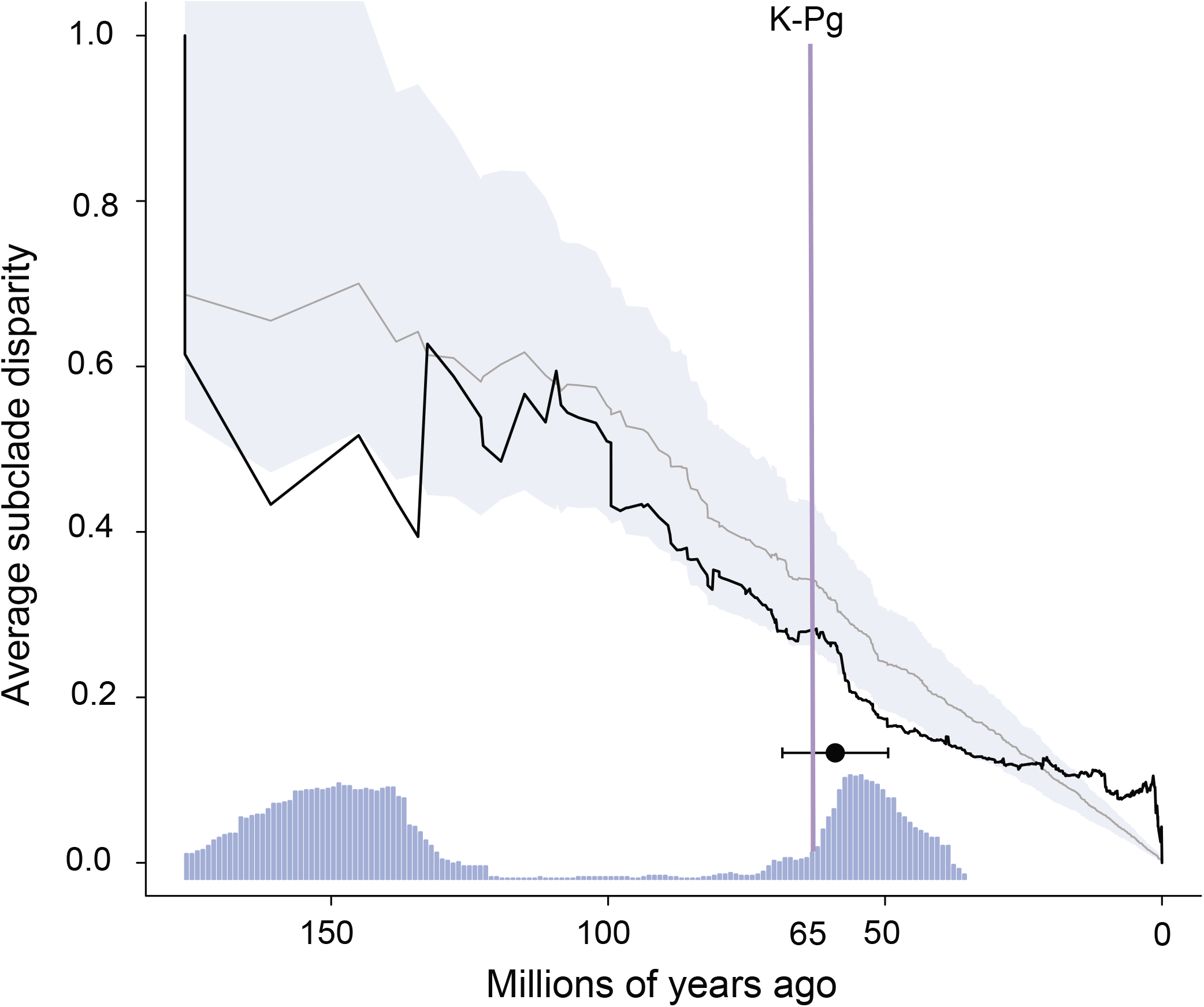
Disparity through time (DTT) for the combined morphological data, repeated on a sample of 100 trees from the posterior distribution of time-trees. The gray line and blue shaded region indicate the median and 95% confidence interval (CI) expected under a Brownian motion model (BM) of evolution, respectively, and solid black line indicates observed pattern of disparity. This is the same plot visualized in Fig. 2, but note that the early portion of the DTT plot that includes the outgroup is not shown in Fig. 2. Acanthomorph body shapes radiated for approximately 20 million years in the aftermath of the K-Pg, followed by within-lineage phenotypic diversification. The blue histogram along the x-axis shows the proportion of time-calibrated trees for which observed disparity falls outside of the BM model’s 95% CI in each one-million year interval. The inset, black box and whisker plot depicts the mean (±95% CI) of the initial time point at which the observed disparity dropped below that expected from BM following the K-Pg.

**Supplementary Fig. 29:**
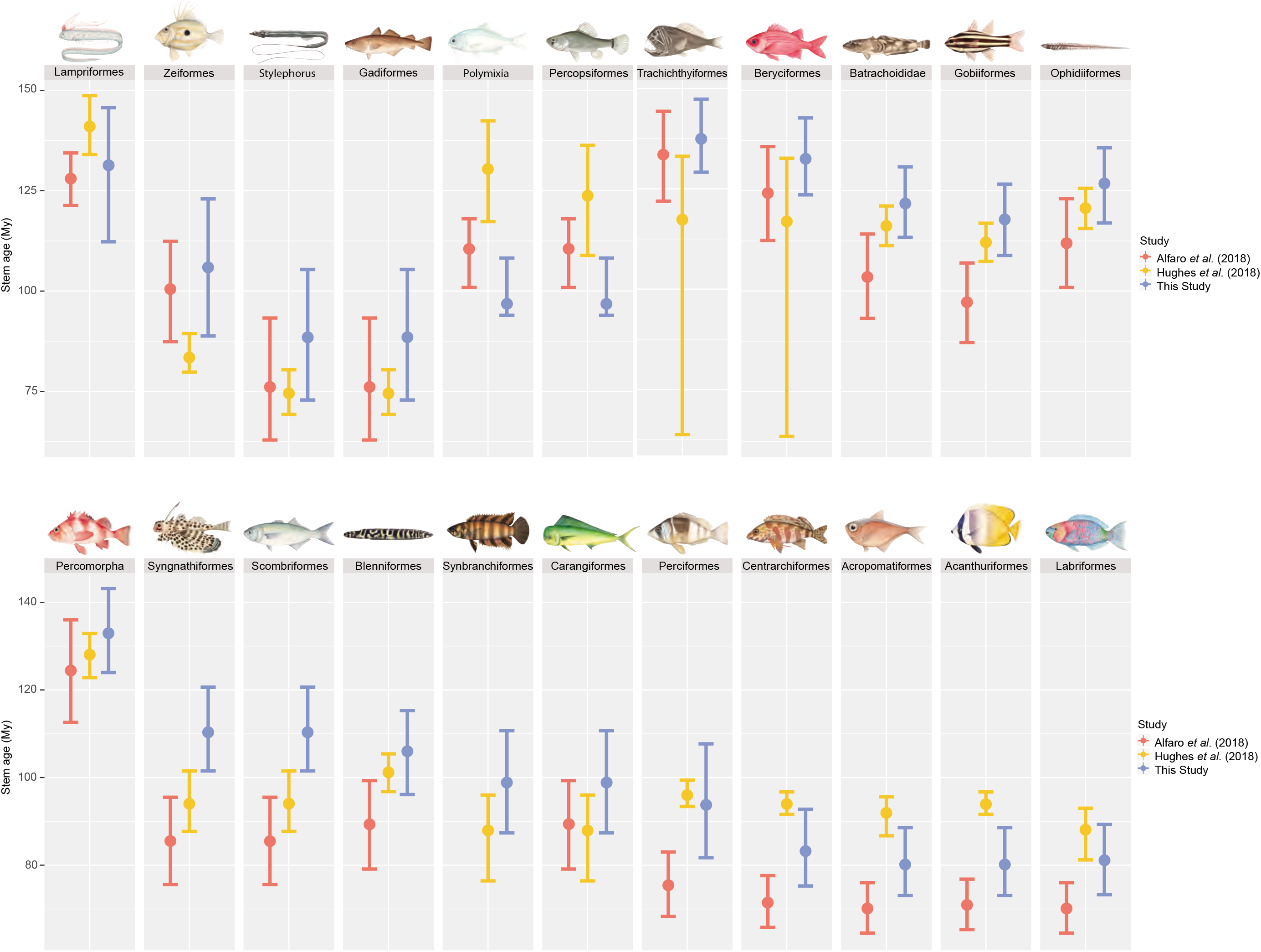
Median stem age estimates and 95% Highest Posterior Densities (HPD) for 22 major acanthomorph clades, as reported in the following 3 phylogenomic studies: Alfaro et al. (2018), Hughes et al. (2018) and this study. Estimates for this study are the raw node heights reported in the 1,084-taxa time tree from Figs. 1 and 2. The 95% HPD of stem ages for most of the represented clades overlap with previous estimates, but we observe some major discrepancies, likely due to differences in tree topologies and taxon sampling.

**Supplementary Fig. 30:**
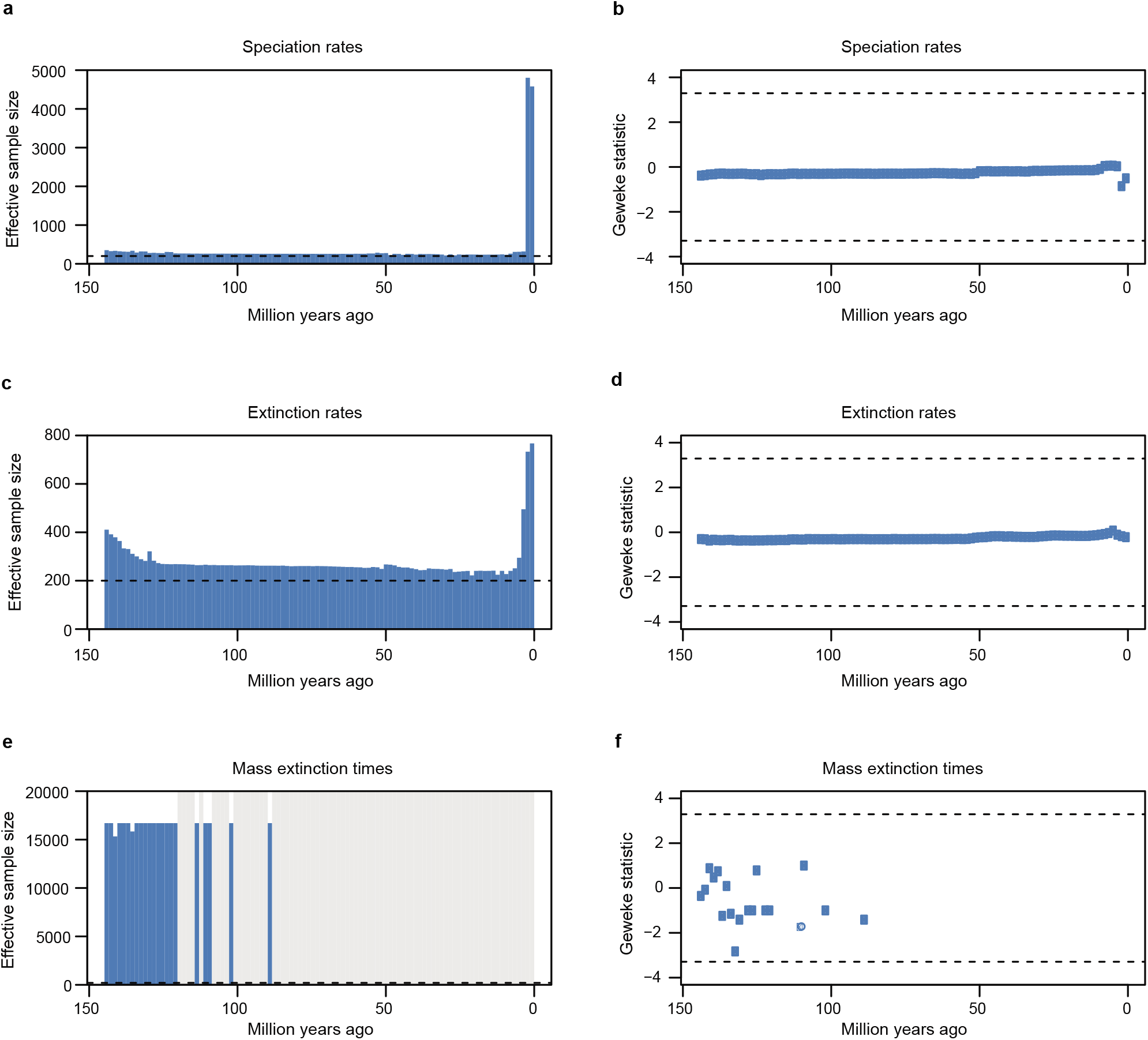
Effective sample sizes (ESS) of parameters and Geweke diagnostics reflect within-chain convergence in the TESS-CoMET analysis depicted in Extended Data Fig.3. Blue bars and dots reflect that the run successfully converged for the parameter at a given time interval. Horizontal dashed lines reflect canonical, acceptable threshold values or 95% confidence intervals for the two diagnostics (ESS ≥ 200 for a, c, and e; P > 0.05 for b, d, and f). a, Effective sample size for speciation rate estimates. b, Geweke statistic for post-burn-in speciation rate estimates. c, Effective sample size for extinction rate estimates. d, Geweke statistic for post-burn-in extinction rate estimates. e, Effective sample size for mass extinction times. f, Geweke statistic for post-burn-in mass extinction times.

**Supplementary Fig. 31:**
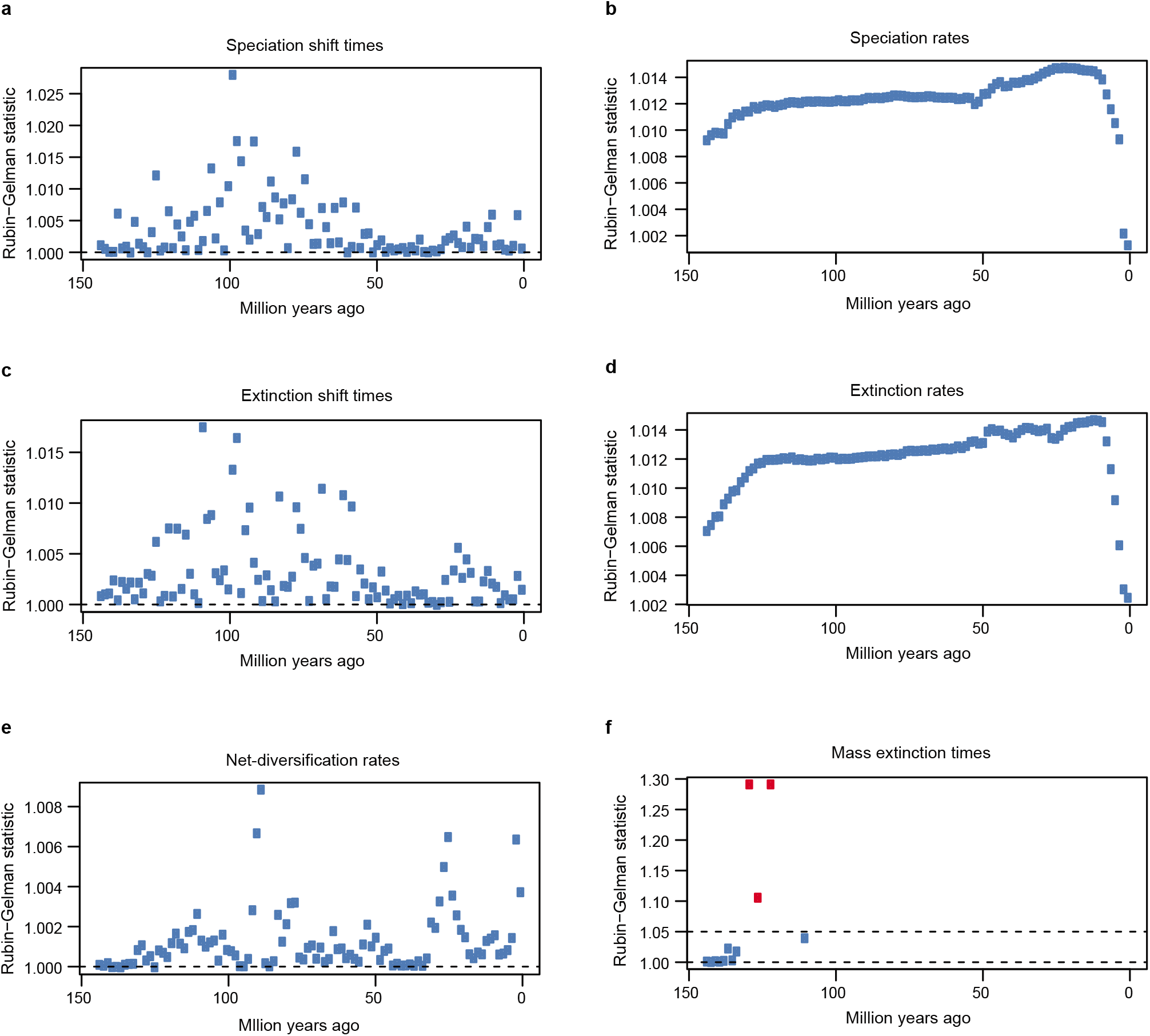
Results of the Gelman-Rubin test show convergence of three independent replicates of the TESS-CoMET analysis represented in Extended Data Fig.3. Blue dots indicate that the ratio of within-chain variance to between-chain variance is <1.05, suggesting the independent MCMC simulations have converged on the same distribution of parameter values. Red dots indicate failed convergence of runs for those time interval-specific parameter estimates. Horizontal dashed lines represent the ideal value (1.00) for the diagnostic. a, Rubin-Gelman statistic for speciation shift times. b, Rubin-Gelman statistic for speciation rates. c, Rubin-Gelman statistic for extinction shift times. d, Rubin-Gelman statistic for extinction rates. e, Rubin-Gelman statistic for net-diversification rates. f, Rubin-Gelman statistic for mass extinction time estimates. A second horizontal dashed line marks the critical value (1.05) above which the test is considered to have failed for that estimate.

**Supplementary Fig. 32:**
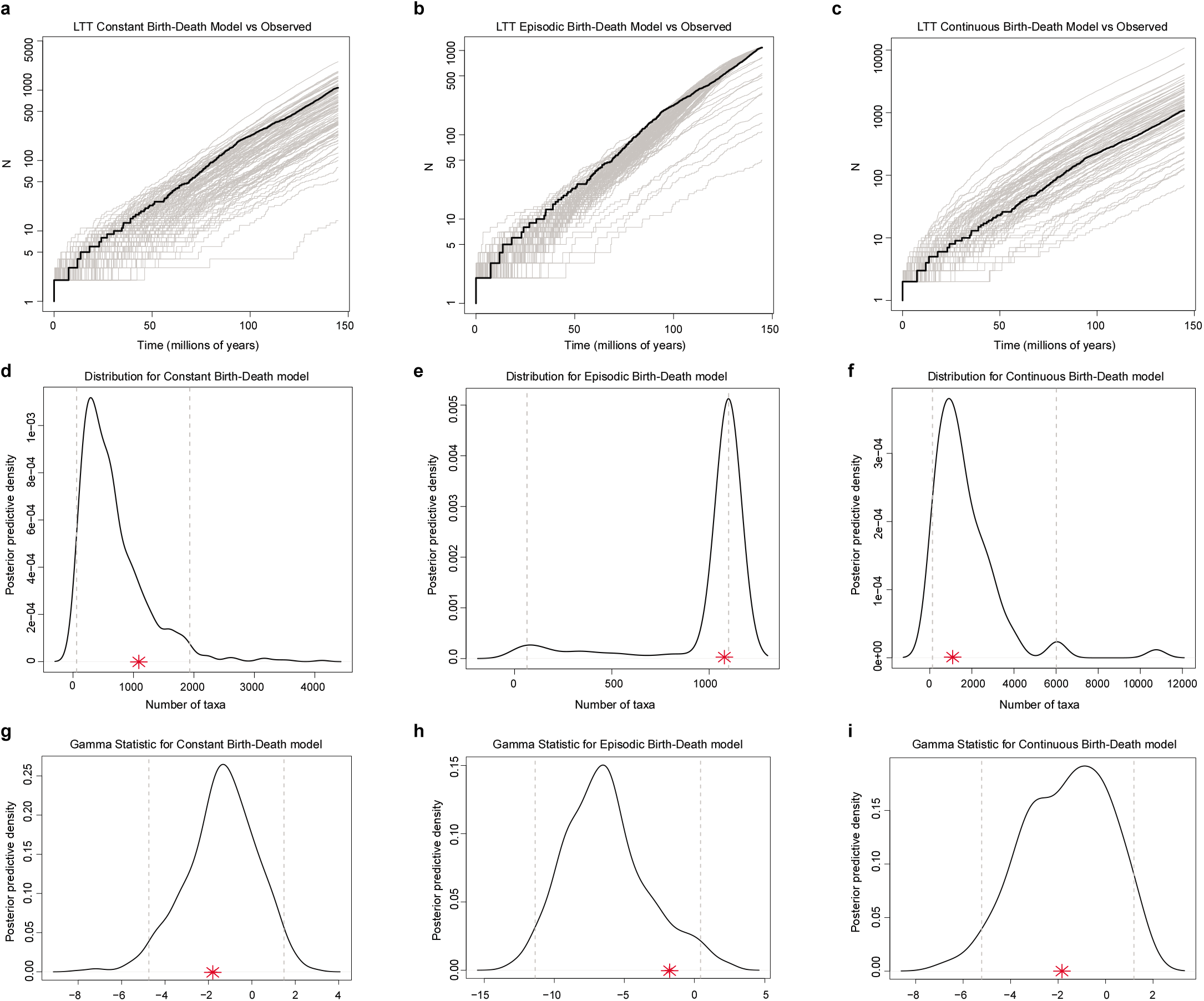
Assessment of the absolute fit of three candidate birth-death (BD) models to the time-calibrated phylogeny of Acanthomorpha. Figures a-c compare lineage-through-time (LTT) curves for simulated trees under a given model (grey lines) to the LTT curve observed for the acanthomorph phylogeny (black line). Figures d-e plot the posterior-predictive distributions for the total number of species estimated to exist in the tree under each model. Dashed lines represent the 95% credible interval for the distribution and red asterisks indicate the total number of acanthomorph species (1,075) in the empirical data. Figures g-i plot the posterior-predictive distributions for the gamma statistic. Dashed lines represent the 95% credible interval for the distribution and red asterisks indicate the value of the gamma statistic empirically calculated for the acanthomorph phylogeny. Candidate models include: 1.) a constant rate birth-death model, 2.) an episodic birth-death model with a rate shift 50 Mya and 3.) a birth-death model with decreasing speciation rates through time. All three candidate models assume uniform (random) incomplete taxon sampling.

**Supplementary Fig. 33:**
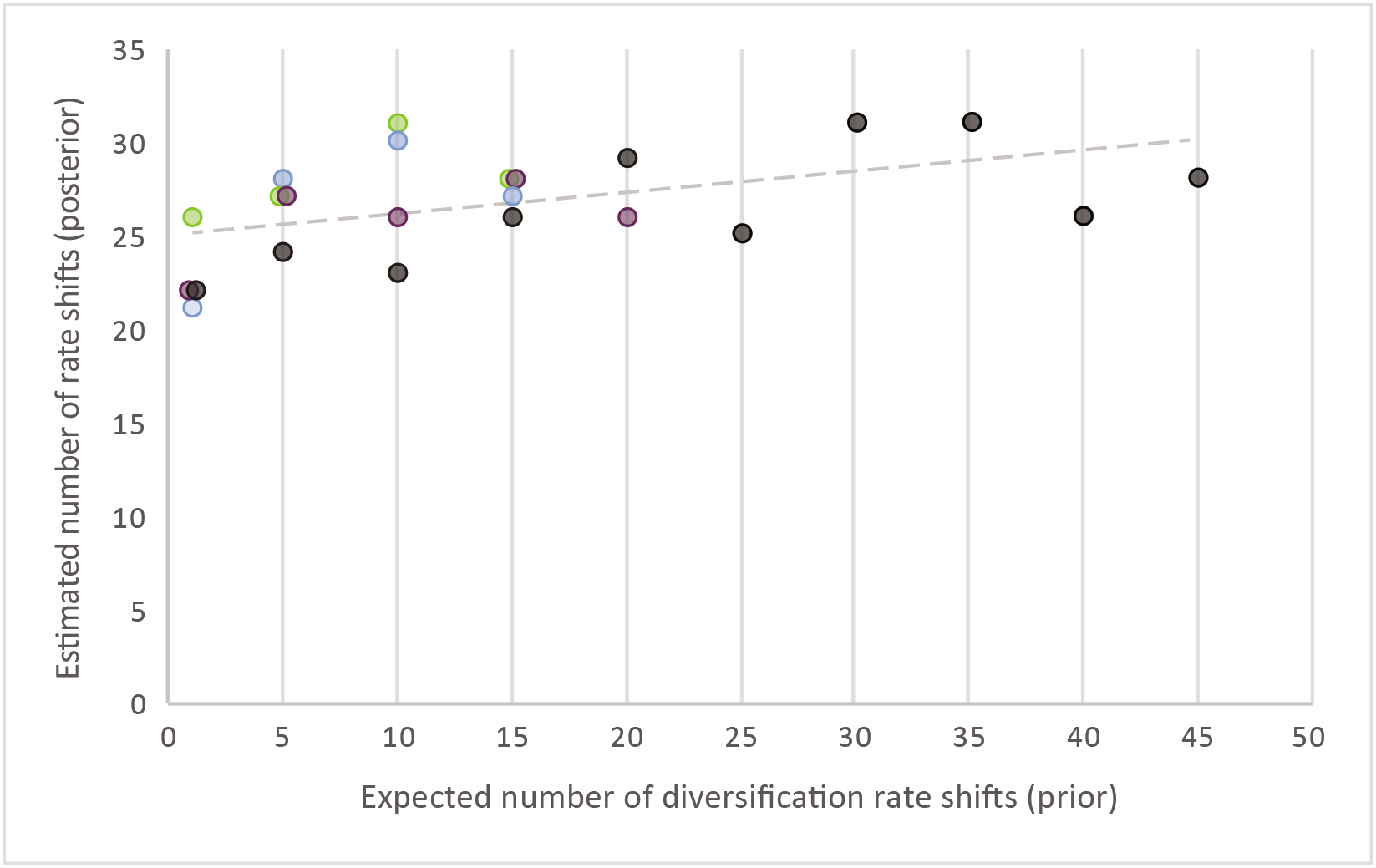
The posterior number of shifts in speciation rates inferred in BAMM is weakly predicted by the prior. Our BAMM analyses used varying numbers of expected speciation rate shifts, and so the resulting maximum shift credibility (MSC) configurations displayed different numbers of estimated rate shifts, shown here as a scatterplot. The slope of the dashed, regression line (0.1049) suggests that changing the inputted prior parameter does not greatly affect the posterior prediction (y = 0.1049x + 25.063; R-squared= 0.217). The standard error of the prior coefficient is 0.0436 and of the intercept is 0.8393. Colors of data points correspond to the settings used when running Markov Chain Monte Carlo (MCMC) simulations in BAMM: black corresponds to 4 MCMC chains in the analysis with a deltaT of 0.05, purple corresponds to 4 MCMC chains in the analysis with a deltaT of 0.1, blue corresponds to 2 MCMC chains in the analysis with a deltaT of 0.05, and green corresponds to 2 MCMC chains in the analysis with a deltaT of 0.1.

## Main references

1 Near, T. J. et al. Phylogeny and tempo of diversification in the superradiation of spiny-rayed fishes. Proc. Nat. Acad. Sci. USA 110, 12738–12743, doi:10.1073/pnas.1304661110 (2013).

2 Wainwright, P. C. & Longo, S. J. Functional innovations and the conquest of the oceans by acanthomorph fishes. Curr. Biol. 27, R550–R557, doi:https://doi.org/10.1016/j.cub.2017.03.044 (2017).

3 Eschmeyer, W. N. & Fricke, R. (California Academy of Sciences ( http://research.calacademy.org/research/ichthyology/catalog/fishcatmain.asp), San Francisco, 2021).

4 Alfaro, M. E. et al. Explosive diversification of marine fishes at the Cretaceous– Palaeogene boundary. Nature Ecol. & Evol. 2, 688–696, doi:10.1038/s41559-018-0494-6 (2018).

5 Meredith, R. W. et al. Impacts of the Cretaceous terrestrial revolution and KPg extinction on mammal diversification. Science 334, 521–524, doi:10.1126/science.1211028 (2011).

6 Stadler, T. Mammalian phylogeny reveals recent diversification rate shifts. Proc. Nat. Acad. Sci. USA 108, 6187–6192, doi:10.1073/pnas.1016876108 (2011).

7 Venditti, C., Meade, A. & Pagel, M. Multiple routes to mammalian diversity. Nature 479, 393–396, doi:Doi 10.1038/Nature10516 (2011).

8 Slater, G. J. Phylogenetic evidence for a shift in the mode of mammalian body size evolution at the Cretaceous-Palaeogene boundary. Methods Ecol Evol 4, 734–744, doi:10.1111/2041-210X.12084 (2013).

9 Liu, L. et al. Genomic evidence reveals a radiation of placental mammals uninterrupted by the KPg boundary. Proceedings of the National Academy of Sciences 114, E7282–E7290, doi:10.1073/pnas.1616744114 (2017).

10 Jetz, W. & Pyron, R. A. The interplay of past diversification and evolutionary isolation with present imperilment across the amphibian tree of life. Nature Ecol. & Evol. 2, 850–858, doi:10.1038/s41559-018-0515-5 (2018).

11 Longrich, N. R., Bhullar, B.-A. S. & Gauthier, J. Mass extinction of lizards and snakes at the Cretaceous-Paleogene boundary. Proc. Nat. Acad. Sci. USA 109, 21396–21401, doi:10.1073/pnas.1211526110 (2012).

12 Jarvis, E. D. et al. Whole-genome analyses resolve early branches in the tree of life of modern birds. Science 346, 1320, doi:10.1126/science.1253451 (2014).

13 Sibert, E. C. & Norris, R. D. New Age of Fishes initiated by the Cretaceous−Paleogene mass extinction. Proceedings of the National Academy of Sciences 112, 8537, doi:10.1073/pnas.1504985112 (2015).

14 Friedman, M. Explosive morphological diversification of spiny-finned teleost fishes in the aftermath of the end-Cretaceous extinction. Proc. R. Soc. B 277, 1675–1683, doi:10.1098/rspb.2009.2177 (2010).

15 Hughes, L. C. et al. Comprehensive phylogeny of ray-finned fishes (Actinopterygii) based on transcriptomic and genomic data. Proc. Nat. Acad. Sci. USA, doi:10.1073/pnas.1719358115 (2018).

16 Patterson, C. An overview of the early fossil record of acanthomorphs. Bull. Mar. Sci. 52, 29–59 (1993).

17 Schluter, D. The ecology of adaptive radiation. (Oxford University Press, 2000).

18 Johnson, G. D. & Patterson, C. Percomorph phylogeny: a survey of acanthomorphs and a new proposal. Bull. Mar. Sci. 52, 554–626 (1993).

19 Miya, M. et al. Major patterns of higher teleostean phylogenies: a new perspective based on 100 complete mitochondrial DNA sequences. Mol. Phylogenet. Evol. 26, 121–138, doi:10.1016/S1055-7903(02)00332-9 (2003).

20 Near, T. J. et al. Resolution of ray-finned fish phylogeny and timing of diversification. Proc. Nat. Acad. Sci. USA 109, 13698–13703, doi:10.1073/Pnas.1206625109 (2012).

21 Betancur-R, R. et al. The tree of life and a new classification of bony fishes. PLOS Cur. Tree of Life 2013 doi:10.1371/currents.tol.53ba26640df0ccaee75bb165c8c26288 (2013).

22 Wainwright, P. C. et al. The evolution of pharyngognathy: a phylogenetic and functional appraisal of the pharyngeal jaw key innovation in labroid fishes and beyond. Syst. Biol. 61, 1001–1027, doi:10.1093/sysbio/sys060 (2012).

23 Ribeiro, E., Davis, A. M., Rivero-Vega, R. A., Ortí, G. & Betancur-R, R. Post-Cretaceous bursts of evolution along the benthic-pelagic axis in marine fishes. Proceedings of the Royal Society B: Biological Sciences 285, 20182010, doi:10.1098/rspb.2018.2010 (2018).

24 Price, S. A. et al. Building a body shape morphospace of teleostean fishes. Integ. Comp. Biol. 59, 716–730, doi:10.1093/icb/icz115 (2019).

25 Smith, W. L., Stern, J. H., Girard, M. G. & Davis, M. P. Evolution of venomous cartilaginous and ray-finned fishes. Integ. Comp. Biol. 56, 950–961, doi:10.1093/icb/icw070 (2016).

26 Schwarzhans, W. & Stringer, G. Fish otoliths from the Late Maastrichtian Kemp Clay (Texas, USA) and the Early Dannian Clayton Formation (Arkansas, USA) and an assessment of extinction and survival of teleost lineages across the K-Pg boundary based on otoliths. Riv. Ital. Paleontol. Strat. 126, 395–446, doi:10.13130/2039-4942/13425 (2020).

27 Claverie, T. & Wainwright, P. C. A morphospace for reef fishes: elongation Is the dominant axis of body shape evolution. PloS one 9, e112732, doi:10.1371/journal.pone.0112732 (2014).

28 Friedman, S. T. et al. Body shape diversification along the benthic–pelagic axis in marine fishes. Proceedings of the Royal Society B: Biological Sciences 287, 20201053, doi:10.1098/rspb.2020.1053 (2020).

29 Friedman, M. et al. A phylogenomic framework for pelagiarian fishes (Acanthomorpha: Percomorpha) highlights mosaic radiation in the open ocean. Proceedings of the Royal Society B: Biological Sciences 286, 20191502, doi:10.1098/rspb.2019.1502 (2019).

30 Near, T. J. et al. Ancient climate change, antifreeze, and the evolutionary diversification of Antarctic fishes. Proc. Nat. Acad. Sci. USA 109, 3434–3439, doi:10.1073/pnas.1115169109 (2012).

31 Price, S. A., Holzman, R., Near, T. J. & Wainwright, P. C. Coral reefs promote the evolution of morphological diversity and ecological novelty in labrid fishes. Ecol Lett 14, 462–469, doi:10.1111/j.1461-0248.2011.01607.x (2011).

32 McGee, M. D. et al. The ecological and genomic basis of explosive adaptive radiation. Nature, doi:10.1038/s41586-020-2652-7 (2020).

33 Jetz, W., Thomas, G. H., Joy, J. B., Hartmann, K. & Mooers, A. O. The global diversity of birds in space and time. Nature 491, 444–448, doi:10.1038/nature11631 (2012).

34 Slater, G. J., Price, S. A., Santini, F. & Alfaro, M. E. Diversity versus disparity and the radiation of modern cetaceans. Proc. R. Soc. B 277, 3097–3104 (2010).

## Methods references

35 Faircloth, B. C. PHYLUCE is a software package for the analysis of conserved genomic loci. Bioinformatics 32, 786–788, doi:10.1093/bioinformatics/btv646 (2016).

36 Faircloth, B. C. et al. Ultraconserved elements anchor thousands of genetic markers spanning multiple evolutionary timescales. Syst. Biol. 61, 717–726, doi:10.1093/sysbio/sys004 (2012).

37 Nguyen, L. T., Schmidt, H. A., von Haeseler, A. & Minh, B. Q. IQ-TREE: A fast and effective stochastic algorithm for estimating maximum likelihood phylogenies. Mol. Biol. Evol. 32, 268–274, doi:10.1093/molbev/msu300 (2015).

38 Kozlov, A. M., Darriba, D., Flouri, T., Morel, B. & Stamatakis, A. RAxML-NG: a fast, scalable and user-friendly tool for maximum likelihood phylogenetic inference. Bioinformatics 35, 4453–4455, doi:10.1093/bioinformatics/btz305 (2019).

39 Puigbò, P., Garcia-Vallvé, S. & McInerney, J. O. TOPD/FMTS: a new software to compare phylogenetic trees. Bioinformatics 23, 1556–1558, doi:10.1093/bioinformatics/btm135 (2007).

40 Mai, U. & Mirarab, S. TreeShrink: fast and accurate detection of outlier long branches in collections of phylogenetic trees. Bmc Genomics 19, 272, doi:10.1186/s12864-018-4620-2 (2018).

41 Katoh, K. & Standley, D. M. MAFFT multiple sequence alignment software Version 7: improvements in performance and usability. Mol. Biol. Evol. 30, 772–780, doi:10.1093/molbev/mst010 (2013).

42 Zhang, C., Rabiee, M., Sayyari, E. & Mirarab, S. ASTRAL-III: polynomial time species tree reconstruction from partially resolved gene trees. BMC Bioinf. 19, 153, doi:10.1186/s12859-018-2129-y (2018).

43 Ronquist, F. et al. MrBayes 3.2: efficient Bayesian phylogenetic inference and model choice across a large model space. Syst. Biol. 61, 539–542, doi:10.1093/sysbio/sys029 (2012).

44 Ane, C., Larget, B., Baum, D. A., Smith, S. D. & Rokas, A. Bayesian estimation of concordance among gene trees. Mol. Biol. Evol. 24, 412–426, doi:10.1093/molbev/msl170 (2007).

45 Bouckaert, R. et al. BEAST 2: a software platform for Bayesian evolutionary analysis. Plos Comput Biol 10, e1003537, doi:10.1371/journal.pcbi.1003537 (2014).

46 Gernhard, T. The conditioned reconstructed process. Journal of Theoretical Biology 253, 769–778, doi:10.1016/j.jtbi.2008.04.005 (2008).

47 Drummond, A. J., Ho, S. Y. W., Phillips, M. J. & Rambaut, A. Relaxed phylogenetics and dating with confidence. PLOS Biol. 4, 699–710, doi:10.1371/journal.pbio.0040088 (2006).

48 Harrington, R. C. et al. Phylogenomic analysis of carangimorph fishes reveals flatfish asymmetry arose in a blink of the evolutionary eye. BMC Evol. Biol. 16, 224, doi:10.1186/s12862-016-0786-x (2016).

49 Lanfear, R., Frandsen, P. B., Wright, A. M., Senfeld, T. & Calcott, B. PartitionFinder 2: New methods for selecting partitioned models of evolution for molecular and morphological phylogenetic analyses. Mol. Biol. Evol. 34, 772–773, doi:10.1093/molbev/msw260 (2017).

50 Rambaut, A., Drummond, A. J., Xie, D., Baele, G. & Suchard, M. A. Posterior summarization in Bayesian phylogenetics using Tracer 1.7. Syst. Biol. 67, 901–904, doi:10.1093/sysbio/syy032 (2018).

51 Drummond, A. J., Suchard, M. A., Xie, D. & Rambaut, A. Bayesian phylogenetics with BEAUti and the BEAST 1.7. Mol. Biol. Evol. 29, 1969–1973, doi:10.1093/molbev/mss075 (2012).

52 Höhna, S., May, M. R. & Moore, B. R. TESS: an R package for efficiently simulating phylogenetic trees and performing Bayesian inference of lineage diversification rates. Bioinformatics 32, 789–791, doi:10.1093/bioinformatics/btv651 (2016).

53 Rabosky, D. L. Automatic detection of key innovations, rate shifts, and diversity-dependence on phylogenetic trees. PLOS One 9, e89543, doi:10.1371/journal.pone.0089543 (2014).

54 Rabosky, D. L. et al. BAMMtools: an R package for the analysis of evolutionary dynamics on phylogenetic trees. Methods Ecol Evol 5, 701–707, doi:10.1111/2041-210X.12199 (2014).

55 Revell, L. J. phytools: an R package for phylogenetic comparative biology (and other things). Methods Ecol Evol 3, 217–223, doi:10.1111/j.2041-210X.2011.00169.x (2012).

56 Harmon, L. J., Weir, J. T., Brock, C. D., Glor, R. E. & Challenger, W. GEIGER: investigating evolutionary radiations. Bioinformatics 24, 129–131, doi:10.1093/bioinformatics/btm538 (2008).

## Supplementary References

57 Longo, S. J. et al. Phylogenomic analysis of a rapid radiation of misfit fishes (Syngnathiformes) using ultraconserved elements. Mol. Phylogenet. Evol. 113, 33–48, doi:https://doi.org/10.1016/j.ympev.2017.05.002 (2017).

58 DiBattista, J. D. et al. Ice ages and butterflyfishes: Phylogenomics elucidates the ecological and evolutionary history of reef fishes in an endemism hotspot. Ecol. & Evol. 8, 10989–11008, doi:10.1002/ece3.4566 (2018).

59 Glenn, T. C. et al. Adapterama I: universal stubs and primers for 384 unique dual-indexed or 147,456 combinatorially-indexed Illumina libraries (iTru & iNext). PeerJ 7, e7755, doi:10.7717/peerj.7755 (2019).

60 Bolger, A. M., Lohse, M. & Usadel, B. Trimmomatic: a flexible trimmer for Illumina sequence data. Bioinformatics 30, 2114–2120 (2014).

61 Grabherr, M. G. et al. Full-length transcriptome assembly from RNA-Seq data without a reference genome. Nat Biotechnol 29, 644–652, doi:10.1038/nbt.1883 (2011).

62 Capella-Gutiérrez, S., Silla-Martínez, J. M. & Gabaldón, T. trimAl: a tool for automated alignment trimming in large-scale phylogenetic analyses. Bioinformatics 25, 1972–1973, doi:10.1093/bioinformatics/btp348 (2009).

63 Minh, B. Q., Nguyen, M. A. T. & von Haeseler, A. Ultrafast Approximation for Phylogenetic Bootstrap. Mol. Biol. Evol. 30, 1188–1195, doi:10.1093/molbev/mst024 (2013).

64 Kalyaanamoorthy, S., Minh, B. Q., Wong, T. K. F., von Haeseler, A. & Jermiin, L. S. ModelFinder: fast model selection for accurate phylogenetic estimates. Nat. Methods 14, 587–589 (2017).

65 Robinson, D. F. & Foulds, L. R. Comparison of phylogenetic trees. Math. Biosci. 53, 131–147 (1981).

66 Steel, M. A. & Penny, D. Distributions of tree comparison metrics: Some new results. Syst. Biol. 42, 126–141. (1993).

67 Oliveros, C. H. et al. Earth history and the passerine superradiation. Proceedings of the National Academy of Sciences 116, 7916, doi:10.1073/pnas.1813206116 (2019).

68 Drummond, A. J. & Rambaut, A. BEAST: Bayesian evolutionary analysis by sampling trees. BMC Evol. Biol. 7, 214 (2007).

69 Rabosky, D. L., Donnellan, S. C., Grundler, M. & Lovette, I. J. Analysis and visualization of complex macroevolutionary dynamics: an example from Australian scincid lizards. Syst. Biol. 63, 610–627 (2014).

70 Davesne, D. et al. Early fossils illuminate character evolution and interrelationships of Lampridiformes (Teleostei, Acanthomorpha). Zool. J. Linn. Soc. 172, 475–498 (2014).

71 Delbarre, D. J., Davesne, D. & Friedman, M. Anatomy and relationships of *Aipichthys pretiosus* and *‘Aipichthys’ nuchalis* (Acanthomorpha: Lampridomorpha), with a review of Late Cretaceous relatives of oarfishes and their allies. J Syst Palaeontol 14, 545–567 (2016).

72 Ogg, J. G. & Hinnov, L. A. in The Geologic Time Scale 2012 Vol. 2 (eds F. Gradstein, J. Ogg, M.D. Schmitz, & G.M. Ogg) 793–853 (Elsevier, 2012).

73 Patterson, C. A review of Mesozoic acanthopterygian fishes, with special reference to those of the English Chalk. Phil. Trans. R. Soc. B 247, 213–482 (1964).

74 Owen, E. in Fossils of the Chalk. Palaeontological Association Field Guides to Fossils: Number 2 (ed A.B. Smith) 9–14 (Oxford University Press, 1987).

75 Friedman, M., Beckett, H. T., Close, R. A. & Johanson, Z. The English Chalk and London Clay: two remarkable British bony fish *Lagerstätten*. Geological Society, London, Special Publications 430, 165, doi:10.1144/SP430.18 (2016).

76 Rosen, D. E. & Patterson, C. The structure and relationships of the paracanthopterygian fishes. Bull. Amer. Mus. Nat. Hist. 141, 357–474 (1969).

77 Murray, A. M. & Wilson, M. V. H. in Mesozoic fishes 2-systematics and fossil record (eds G. Arratia & H.-P. Schultze) 397–411 (Verlag Dr. Friedrich Pfeil, 1999).

78 Armbruster, J. W., Niemiller, M. L. & Hart, P. B. Morphological evolution of the Cave-, Spring-, and Swampfishes of the Amblyopsidae (Percopsiformes). Copeia 104, 763–777, doi:10.1643/CI-15-339 (2016).

79 Evanoff, E., McIntosh, W. C. & Murphey, P. C. Stratigraphic summary and ^40^Ar/^39^Ar geochronology of the Florissant Formation, Colorado. Proc. Denver Mus. Nature Sci. Series 4, no. 1, 1–16 (2001).

80 Danil’chenko, P. G. Kostistye ryby Maikopskikh otlozhenii Kavkaza [Bony fishes of the Maikop deposits of the Caucasus]. Trudy Paleon. Inst. 78, 1–208. [In Russian] (1960).

81 Baciu, D.-S., Bannikov, A. F. & Tyler, J. C. Revision of the fossil fishes of the family Zeidae (Zeiformes). Boll. Mus. Storia Nat. Verona, Geol. Paleon. Preist. 29, 95–128 (2005).

82 Santini, F., Tyler, J. C., Bannikov, A. F. & Baciu, D. S. A phylogeny of extant and fossil buckler dory fishes, family Zeidae (Zeiformes, Acanthomorpha). Cybium 30, 99–107 (2006).

83 Jones, R. W. & Simmons, M. D. A review of the stratigraphy of Eastern Paratethys (Oligocene-Holocene), with particular emphasis on the Black Sea. AAPG Mem. 68, 39–52 (1997).

84 Luterbacher, H. P. et al. in A geologic time scale 2004 (eds F. Gradstein, J. Ogg, & A. Smith) 384–408 (Cambridge University Press, 2004).

85 Danil’chenko, P. G. in Ocherki po filogenii i sistematike iskopayemykh ryb i beschelyustnykh [Outlines on the phylogeny and systematics of fossil fishes and agnathans] (ed D.V. Obruchev) 113–156. [In Russian] (Nauka, 1968).

86 Bannikov, A. F. Review of fossil Lampridiformes (Teleostei) finds with a description of a new Lophotidae genus and species from the Oligocene of the northern Caucasus. Paleontol. J. 33, 68–76 (1999).

87 Bannikov, A. F. & Parin, N. N. The list of marine fishes from Cenozoic (Upper Paleocene-middle Miocene) localities in southern European Russia and adjacent countries. J. Ichthyol. 37, 150–155 (1997).

88 Patterson, C. New Cretaceous berycoid fishes from the Lebanon. Bull. Brit. Mus. (Nat. Hist.) Geol. 14, 69–109 (1967).

89 Zehren, S. J. The comparative osteology and phylogeny of the Beryciformes. Evolutionary Monogr. 1, 1–389 (1979).

90 Sorbini, L. Gli Holocentridae di Monte Bolca. I: *Eoholocentrum, nov. gen*., Eoholocentrum macrocephalum (de Blainville) (Pisces-Actinopterygii). Stud. Ric. Giaciam. Terz. Bolca 2, 205–228 (1975).

91 Sorbini, L. Gli Holocentridae di Monte Bolca. II: *Tenuicentrum pattersoni* nov. gen. nov. sp. Nuovi dati a favoure dell’origine monofiletica dei beryciformi (Pisces). Stud. Ric. Giaciam. Terz. Bolca 2, 456–472 (1975).

92 Sorbini, L. Gli Holocentridae di Monte Bolca. III. *Berybolcensis leptacanthus* (Agassiz). Stud. Ric. Giaciam. Terz. Bolca 4, 19–35 (1979).

93 Stewart, J. D. Taxonomy, paleoecology, and stratigraphy of the halecostome-inoceramid associations of the North American Upper Cretaceous epicontinental seaway, University of Kansas, (1984).

94 Papazzoni, C., Carnevale, G., Fornaciari, E., Giusberti, L. & Trevisani, E. 29–36 (2014).

95 Papazzoni, C. A. & Trevisani, E. Facies analysis, palaeoenvironmental reconstruction, and biostratigraphy of the “Pesciara di Bolca” (Verona, northern Italy): An early Eocene *Fossil-Lagerstätte*. Palaeogeogr. Palaeoclimat. Palaecol. 242, 21–35 (2006).

96 Sorbini, L. The Cretaceous fishes of Nardò. I°. Order Gasterosteiformes (Pisces). Boll. Mus. Civ. Stor. Nat. Verona 8, 1–27 (1981).

97 Orr, J. W. Phylogenetic relationships of gasterosteiform fishes (Teleostei: Acanthomorpha), University of Washington, (1995).

98 Pietsch, T. W. Evolutionary relationships of the sea moths (Teleostei: Pegasidae) with a classification of the gasterosteiform families. Copeia 1978, 517–529 (1978).

99 Medizza, F. & Sorbini, L. in I vertebrati fossili italiani—Catalogo dell Mostra 131–134 (Museo Civico di Storia Naturale, 1980).

100 Blot, J. La faune ichthyologique des gisements du Monte Bolca (Province de Vérone, Italie). Catalogue systématique présentat l’état actuel des recherches concernant cette faune. Bull. Mus. nation. d’Hist. nat., Paris, 4e serie, sec. C 2, 339-396 (1980).

101 Doiuchi, R. & Nakabo, T. Molecular phylogeny of the stromateoid fishes (Teleostei : Perciformes) inferred from mitochondrial DNA sequences and compared with morphology-based hypotheses. Mol. Phylogenet. Evol. 39, 111–123 (2006).

102 Carnevale, G. & Bannikov, A. F. Description of a new stromateoid fish from the Miocene of St. Eugene, Algeria. Acta Palaeontol Pol 51, 489–497 (2006).

103 Horn, M. H. Systematic Status and Aspects of the Ecology of the Elongate Ariommid Fishes (Suborder Stromateoidei) in the Atlantic. Bull. Mar. Sci. 22, 537–558 (1972).

104 Bannikov, A. F. An new species of stromateoid fishes (Perciformes) the Lower Oligocene of the Caucasus. Paleontol. J. 22, 107–112 (1988).

105 Bannikov, A. Morphology and phylogeny of fossil stromateoid fishes (Perciformes). Geobios 28, Supplement 2, 177–181 (1995).

106 Doiuchi, R., Sato, T. & Nakabo, T. Phylogenetic relationships of the stromateoid fishes (Perciformes). Ichthyol. Res. 51, 202–212 (2004).

107 Carnevale, G. Fossil fishes from the Serravallian (Middle Miocene) of Torricella Peligna, Italy. Palaeontographia Italica 91, 1–67 (2007).

108 Santini, F., Carnevale, G. & Sorenson, L. First molecular scombrid timetree (Percomorpha: Scombridae) shows recent radiation of tunas following invasion of pelagic habitat. Ital. J. Zool. 80, 210–221, doi:Doi 10.1080/11250003.2013.775366 (2013).

109 Arambourg, C. Resultats scientifiques de la mission C. Arambourg en Syrie et en Iran (1938–1939). II. Les poissons Oligocène de l’Iran. Notes Mém. Moyen-Orient 8, 1-210 (1967).

110 Matsui, T. Review of mackerel genera *Scomber* and *Rastrelliger* with description of a new species of *Rastrelliger*. Copeia 1967, 71–83 (1967).

111 Monsch, K. A. Revision of the scombroid fishes from the Cenozoic of England. T Roy Soc Edin-Earth 95, 445–489 (2005).

112 Monsch, K. A. The PhyloCode, or alternative nomenclature: Why it is not beneficial to palaeontology, either. Acta Palaeontol Pol 51, 521–524 (2006).

113 Collette, B. B. & Russo, J. L. Morphology, systematics, and biology of the Spanish Mackerels (*Scomberomorus*, Scombridae). Fish. Bull. 82, 545–692 (1984).

114 Leriche, M. Les poissons éocènes de la Belgique. Memoirs of the Royal Belgian Museum of Natural Sciences 3, 49–228 (1905).

115 Monsch, K. A. & Bannikov, A. F. New taxonomic synopses and revision of the scombroid fishes (Scombroidei, Perciformes), including billfishes, from the Cenozoic of territories of the former USSR. Earth Env Sci T R So 102, 253–300, doi:Doi 10.1017/S1755691011010085 (2011).

116 Bannikov, A. F. Fossil scombrids of the USSR. Trudy Paleon. Inst. 210, 1–111 (1985).

117 Murray, A. M. & Thewissen, J. G. M. Eocene actinopterygian fishes from Pakistan, with the description of a new genus and species of channid (Channiformes). J. Vert. Paleo. 28, 41–52 (2008).

118 Gingerich, P. D. Stratigraphic and micropaleontological constraints on the middle Eocene age of the mammal-bearing Kuldana Formation of Pakistan. J. Vert. Paleo. 23, 643–651, doi:Doi 10.1671/2409 (2003).

119 Rabosky, D. L. et al. An inverse latitudinal gradient in speciation rate for marine fishes. Nature, doi:10.1038/s41586-018-0273-1 (2018).

120 Bannikov, A. The systematic composition of the Eocene actinopterygian fish fauna from Monte Bolca, northern Italy, as known to date. St. Ric. Giac. Terz. Bolca 15 (2014).

121 Carnevale, G., Bannikov, A., Marramà, G., Tyler, J. C. & Zorzin, R. in The Bolca Fossil-Lagerstatten: A window into the Eocene World (eds C. Andrea Papazzoni et al.) 37–63 (Rendiconti della Società Paleontologica Italiana, 2014).

122 Johnson, G. D. Scombroid phylogeny: an alternative hypothesis. Bull. Mar. Sci. 39, 1–41 (1986).

123 Friedman, M. & Johnson, G. D. A new species of *Mene* (Perciformes: Menidae) from the Paleocene of South America, with notes on paleoenvironment. J. Vert. Paleo. 25, 770–783 (2005).

124 Bonde, N. A distinct fish fauna in the basal ashseries of the Fur/Ølst Formation (U. Paleocene, Denmark). Aarhus Geoscience 6, 33–48 (1997).

125 Anthonissen, D. E. & Ogg, J. G. in The Geologic Time Scale 2012 (eds F. Gradstein, J. Ogg, M.D. Schmitz, & G.M. Ogg) 1083–1127 (Elsevier, 2012).

126 Smith-Vaniz, W. F. in Ontogeny and systematics of fishes (eds H.G. Moser et al.) 522–530 (Allen Press, 1984).

127 Eastman, C. R. Descriptions of Bolca fishes. Bull. Mus. Comp. Zool. 46, 1–36 (1904).

128 Gushiken, S. Phylogenetic relationships of the perciform genera of the family Carangidae. Jap. J. Ich. 34, 443–461 (1988).

129 Friedman, M., Johanson, Z., Harrington, R. C., Near, T. J. & Graham, M. R. An early fossil remora (Echeneoidea) reveals the evolutionary assembly of the adhesion disc. Proc. R. Soc. B 280, 20131200, doi:10.1098/rspb.2013.1200 (2013).

130 Micklich, N. R. New information on the fish fauna of the Frauenweiler fossil site. Ital. J. Zool. 65, 169–184 (1998).

131 O’Toole, B. Phylogeny of the species of the superfamily Echeneoidea (Perciformes : Carangoidei: Echeneidae, Rachycentridae, and Coryphaenidae), with an interpretation of echeneid hitchhiking behaviour. Can. J. Zool. 80, 596–623 (2002).

132 Bannikov, A. F. Fossil carangids and apolectids of the USSR. Trudy Paleon. Inst. 244, 1–106 (1990).

133 Friedman, M. The evolutionary origin of flatfish asymmetry. Nature 454, 209–212 (2008).

134 Chanet, B. A cladistic reappraisal of the fossil flatfishes record consequences on the phylogeny of the Pleuronectiformes (Osteichthyes: Teleostei). Ann. Sci. nat., Zool., Paris 13e Sér. 18 (1997).

135 Chanet, B. Eubuglossus eocenicus (Woodward 1910) from the Upper Lutetian of Egypt, one of the oldest soleids (Teleostei, Pleuronectiformes). Neu. Jahrbuch Geolo. Paläon., Monatshefte 1994, 391–398 (1994).

136 Chapleau, F. & Keast, A. A phylogenetic reassessment of the monophyletic status of the family Soleidae, with comments on the suborder Soleoidei (Pisces, Pleuronectiformes). Can. J. Zool. 66, 2797–2810, doi:DOI 10.1139/z88-408 (1988).

137 Baciu, D. S. & Chanet, B. Les poissons plats fossils (Teleostei: Pleuronectiformes) de l’Oligocène de Piatra Neamt (Roumanie). Oryctos 4, 17–38 (2002).

138 Sakamoto, K., Uyeno, T. & Micklich, N. *Oligopleuronectes germanicus* gen. et sp. nov., an Oligocene pleuronectid flatfish from Frauenweiler, S-Germany. Bulletin of the National Science Museum Series C (Geology & Paleontology) 30, 89–94 (2004).

139 Cooper, J. A. & Chapleau, F. Phylogenetic status of Paralichthodes algoensis (Pleuronectiformes: Paralichthodidae). Copeia 1998, 477–481. (1998).

140 Carnevale, G., Bannikov, A. F., Landini, W. & Sorbini, C. Volhynian (early Sarmatian sensu lato) fishes from Tsurevsky, North Caucasus, Russia. J. Paleontol. 80, 684–699, doi:Doi 10.1666/0022-3360(2006)80[684:Vesslf]2.0.Co;2 (2006).

141 Chanet, B. & Sorbini, C. A male fish *Bothus podas* (Delaroche, 1809) (Pleuronectiformes: Bothidae) in the Pliocene of the Marecchia river (Italy). Bollettino della Societa Paleontologica Italiana 40, 345-350 (2001).

142 Murray, A. M. Eocene cichlid fishes from Tanzania, East Africa. J. Vert. Paleo. 20, 651–664 (2000).

143 Murray, A. M. The oldest fossil cichlids (Teleostei: Perciformes): Indication of a 45 million-year-old species flock. Proc. R. Soc. B 268, 679–684. (2001).

144 Murray, A. M. The fossil record and biogeography of the Cichlidae (Actinopterygii: Labroidei). Biol. J. Linn. Soc. 74, 517–532 (2001).

145 Harrison, T. et al. in Eocene biodiversity: Unusual occurrences and rarely sampled habitats (ed G.F. Gunnell) 39–74 (Kluwer Academic/Plenum Publishers, 2001).

146 Benton, M. J. et al. Constraints on the timescale of animal evolutionary history. Palaeontologia Electronica 18 (2015).

147 Bannikov, A., Parin, N. V. & Pinna, J. *Rhamphexocoetus volans*, gen. et sp. nov. a new beloniform fish (Beloniformes, Exocoetidei) from the lower Eocene of Italy. J. Ichthyol. 25(2), 150–155 (1985).

148 Rosen, D. E. The relationships and taxonomic position of the halfbeaks, killifishes, silversides and their relatives. Bull. Amer. Mus. Nat. Hist. 127, 217–268 (1964).

149 Rosen, D. E. & Parenti, L. R. Relationships of *Oryzias* and the groups of atherinomorph fishes. Amer. Mus. Novit. 2719, 1–25 (1981).

150 Collette, B. B., McGowen, G. E., Parin, N. V. & Mito, S. in Ontogeny and systematics of fishes (eds H.G. Moser et al.) 335–354 (American Society of Ichthyologists and Herpetologists, 1984).

151 Bellwood, D. R. & Schultz, O. A review of the fossil record of parrotfishes (Labroidei: Scaridae) with a description of a new *Calatomus* species from the Middle Miocene (Badenian) of Austria. Ann. Naturhist. Mus. Wien 92, 55–71 (1991).

152 Bannikov, A. F. & Tyler, J. C. Phylogenetic revision of the fish families Luvaridae and †Kushlukiidae (Acanthuroidei), with a new genus and two new species of Eocene luvarids. Smithson. Contrib. Paleobiol. 81, 1–45 (1995).

153 Blot, J. & Tyler, J. C. New genera and species of fossil surgeon fishes and their relatives (Acanthuroidei, Teleostei) from the Eocene of Monte Bolca, Italy, with application of the Blot Formula to both fossil and recent forms. St. Ric. Giac. Terz. Bolca 6, 13–92 (1990).

154 Tyler, J. C. & Bannikov, A. F. Relationships of the fossil and recent genera of rabbitfishes (Acanthuroidei: Siganidae). Smithson. Contrib. Paleobiol. 84, 1–35 (1997).

155 Micklich, N. R., Tyler, J. C., Johnson, G. D., Swidnicka, E. & Bannikov, A. F. First fossil records of the tholichthys larval stage of butterfly fishes (Perciformes, Chaetodontidae), from the Oligocene of Europe. Palaontol. Z. 83, 479–497 (2009).

156 Carnevale, G. Morphology and biology of the Miocene butterflyfish *Chaetodon ficheuri* (Teleostei: Chaetodontidae). Zool. J. Linn. Soc. 146, 251–267 (2006).

157 Blum, S. D. The osteology and phylogeny of the Chaetodontidae (Teleostei: Perciformes), University of Hawaii, (1988).

158 Krijgsman, W., Hilgen, F. J., Raffi, I., Sierro, F. J. & Wilson, D. S. Chronology, causes and progression of the Messinian salinity crisis. Nature 400, 652–655 (1999).

159 Hilgen, F. J. et al. Extending the astronomical (polarity) time scale into the Miocene. Earth Planet. Sci. Lett. 136, 495–510 (1995).

160 Carnevale, G. The first fossil ribbonfish (Teleostei, Lampridiformes, Trachipteridae). Geol. Mag. 141, 573–582 (2004).

161 Yabumoto, Y. & Uyeno, T. A new Miocene ponyfish of the genus *Leiognathus* (Pisces, Leiognathidae). Bull. Nat. Sci. Mus., Tokyo Ser. C 20, 67–77 (1994).

162 Yabumoto, Y. & Uyeno, T. *Euleiognathus*, a new genus proposed for the Miocene ponyfish, Leiognathus tottori Yabumoto and Uyeno 1994 (Perciformes: Leiognathidae) from Japan. Ichthyol. Res. 58, 19-23 (2011).

163 Chakrabarty, P. & Sparks, J. S. Diagnoses for *Leiognathus* Lacepede 1802, *Equula* Cuvier 1815, *Equulites* Fowler 1904, *Eubleekeria* Fowler 1904, and a new ponyfish genus (Teleostei: Leiognathidae). Amer. Mus. Novit., 1-11 (2008).

164 Yamashita, T. & Kimura, S. A new species, *Gazza squamiventralis*, from the East Coast of Africa (Perciformes: Leiognathidae). Ichthyol. Res. 48, 161–166 (2001).

165 Sparks, J. S. & Chakrabarty, P. Description of a new genus of ponyfishes (Teleostei: Leiognathidae), with a review of the current generic-level composition of the family. Zootaxa 3947, 181–190 (2015).

166 Sparks, J. S., Dunlap, P. V. & Smith, W. L. Evolution and diversification of a sexually dimorphic luminescent system in ponyfishes (Teleostei: Leiognathidae), including diagnoses for two new genera. Cladistics 21, 305–327 (2005).

167 Gill, A. C. & Michalski, S. Osteological evidence for monophyly of the Leiognathidae (Teleostei: Acanthomorpha: Acanthuriformes). Zootaxa; Vol 4732, No 3: 13 Feb. 2020 DO - 10.11646/zootaxa.4732.3.4 (2020).

168 Lourens, L., Hilgen, F., Shackleton, N. J., Laskar, J. & Wilson, D. in A geologic time scale 2004 (eds F. Gradstein, J. Ogg, & A. Smith) 409–440 (Cambridge University Press, 2004).

169 Santini, F. & Tyler, J. C. A phylogeny of the families of fossil and extant tetraodontiform fishes (Acanthomorpha, Tetraodontiformes), Upper Cretaceous to recent. Zool. J. Linn. Soc. 139, 565–617 (2003).

170 Bannikov, A. F. & Tyler, J. C. A new genus and species of triggerfish from the Middle Eocene of the northern Caucasus, the earliest member of the Balistidae (Tetraodontiformes). Paleontol. J. 42, 615–620 (2008).

171 Carnevale, G. & Pietsch, T. W. An Eocene frogfish from Monte Bolca, Italy: The earliest known skeletal record for the family. Palaeontology 52, 745–752 (2009).

172 Pietsch, T. W. The osteology and relationships of the anglerfish genus *Tetrabrachium* with comments on lophiiform classification. Fish. Bull. 79, 387–419 (1981).

173 Pietsch, T. W. in Ontogeny and systematics of fishes (eds H.G. Moser, et al.) 320–325 (American Society of Ichthyologists and Herpetologists, 1984).

174 Betancur-R, R. et al. Phylogenetic classification of bony fishes. BMC Evol. Biol. 17, 162, doi:10.1186/s12862-017-0958-3 (2017).

175 Greenwood, P. H., Rosen, D. E., Weitzman, S. H. & Myers, G. S. Phyletic studies of teleostean fishes, with a provisional classification of living forms. Bull. Amer. Mus. Nat. Hist. 131, 341–455 (1966).

176 Nelson, J. S., Grande, T. C. & Wilson, M. V. H. Fishes of the world. 5th edn, (John Wiley & Sons, Inc., 2016).

177 Li, B. et al. RNF213, a new nuclear marker for acanthomorph phylogeny. Mol. Phylogenet. Evol. 50, 345–363 (2009).

178 Malmstrøm, M. et al. Evolution of the immune system influences speciation rates in teleost fishes. Nat Genet 48, 1204–1210, doi:10.1038/ng.3645 http://www.nature.com/ng/journal/v48/n10/abs/ng.3645.html - supplementary-information (2016).

179 Roa-Varón, A. et al. Confronting sources of systematic error to resolve historically contentious relationships: a case study using gadiform fishes (Teleostei, Paracanthopterygii, Gadiformes). Syst. Biol., doi:10.1093/sysbio/syaa095 (In press).

180 Dornburg, A. et al. New insights on the sister lineage of percomorph fishes with an anchored hybrid enrichment dataset. Mol. Phylogenet. Evol. 110, 27–38, doi:https://doi.org/10.1016/j.ympev.2017.02.017 (2017).

181 Fricke, R., Eschmeyer, W. N. & Fong, J. D. Eschmeyer’s Catalog of Fishes: species by family/subfamily, <http://researcharchive.calacademy.org/research/ichthyology/catalog/SpeciesByFamily.asp> (2021).

182 Gill, F., Donsker, D. & Rasmussen, P. IOC World Bird List (v10.2), 2020).

183 Burgin, C. J., Colella, J. P., Kahn, P. L. & Upham, N. S. How many species of mammals are there? J. Mammal. 99, 1–14, doi:10.1093/jmammal/gyx147 (2018).

184 Uetz, P., Freed, P. & Hošek, J. The Reptile Database, http://www.reptile-database.org accessed December 2020, 2020).

185 Jordan, D. S. A classification of fishes including families and genera as far as known. Stanford U. Publ. Univ. Ser. Biol. Sci. 3, 77–243 (1923).

186 Nelson, J. S. Fishes of the world, 4th edition. (John Wiley, 2006).

187 Smith, W. L. & Craig, M. T. Casting the percomorph net widely: the importance of broad taxonomic sampling in the search for the placement of serranid and percid fishes. Copeia 2007, 35–55 (2007).

188 Nelson, G. in The hierarchy of life: molecules and morphology in phylogenetic analysis (eds B. Fernholm, K. Bremer, & H. Jôrnvall) 325–336 (Elsevier, 1989).

189 Møller, P. R., Knudsen, S. W., Schwarzhans, W. & Nielsen, J. G. A new classification of viviparous brotulas (Bythitidae) – with family status for Dinematichthyidae – based on molecular, morphological and fossil data. Mol. Phylogenet. Evol. 100, 391–408, doi:http://dx.doi.org/10.1016/j.ympev.2016.04.008 (2016).

190 Campbell, M. A. et al. Evolutionary affinities of the unfathomable Parabrotulidae: molecular data indicate placement of *Parabrotula* within the family Bythitidae, Ophidiiformes. Mol. Phylogenet. Evol. 109, 337–342, doi:https://doi.org/10.1016/j.ympev.2017.02.004 (2017).

191 Evseenko, S. A., Gordeeva, N. V., Bolshakova, Y. Y. & Kobyliansky, S. H. Morphology and molecular phylogenetic relationships of *Barathronus multidens* (Ophidiiformes: Bythitidae). Cybium: international journal of ichthyology 42, 137–141 (2018).

192 Miya, M., Satoh, T. R. & Nishida, M. The phylogenetic position of toadfishes (order Batrachoidiformes) in the higher ray-finned fish as inferred from partitioned Bayesian analysis of 102 whole mitochondrial genome sequences. Biol. J. Linn. Soc. 85, 289–306 (2005).

193 Smith, W. L. & Wheeler, W. C. Venom evolution widespread in fishes: a phylogenetic road map for the bioprospecting of piscine venoms. J. Hered. 97, 206–217 (2006).

194 Thacker, C. E. Phylogeny of Gobioidei and placement within Acanthomorpha, with a new classification and investigation of diversification and character evolution. Copeia 2009, 93–104 (2009).

195 Chakrabarty, P., Davis, M. P. & Sparks, J. S. The first record of a trans-oceanic sister-group relationship between obligate vertebrate troglobites. Plos One 7 (2012).

196 Kuang, T. et al. Phylogenomic analysis on the exceptionally diverse fish clade Gobioidei (Actinopterygii: Gobiiformes) and data-filtering based on molecular clocklikeness. Mol. Phylogenet. Evol. 128, 192–202, doi:https://doi.org/10.1016/j.ympev.2018.07.018 (2018).

197 McCraney, W. T., Thacker, C. E. & Alfaro, M. E. Supermatrix phylogeny resolves goby lineages and reveals unstable root of Gobiaria. Mol. Phylogenet. Evol. 151, 106862, doi:https://doi.org/10.1016/j.ympev.2020.106862 (2020).

198 Thacker, C. E. et al. Molecular phylogeny of Percomorpha resolves *Trichonotus* as the sister lineage to Gobioidei (Teleostei: Gobiiformes) and confirms the polyphyly of Trachinoidei. Mol. Phylogenet. Evol. 93, 172–179, doi:https://doi.org/10.1016/j.ympev.2015.08.001 (2015).

199 Orrell, T. M., Collette, B. B. & Johnson, G. D. Molecular data support separate scombroid and xiphioid clades. Bull. Mar. Sci. 79, 505–519 (2006).

200 Song, H. Y. et al. Mitogenomic circumscription of a novel percomorph fish clade mainly comprising “Syngnathoidei” (Teleostei). Gene 542, 146–155, doi:10.1016/j.gene.2014.03.040 (2014).

201 Gosline, W. A. A reinterpretation of the teleostean fish order Gobiesociformes. Proc. California Acad. Sci. 38, 363–381 (1970).

202 Nelson, J. S. Fishes of the world, 3rd edition. 3rd edn, (Wiley, 1994).

203 Springer, V. G. & Johnson, G. D. Study of the dorsal gill-arch musculature of teleostome fishes, with special reference to the Actinopterygii. Bull. Biol. Soc. Wash. 11, 1–260 (2004).

204 Wiley, E. O. & Johnson, G. D. in Origin and phylogenetic interrelationships of teleosts (eds J.S. Nelson, H.-P. Schultze, & M.V.H. Wilson) 123–182 (Verlag Dr. Friedrich Pfeil, 2010).

205 Johnson, G. D. Percomorph phylogeny: progress and problems. Bull. Mar. Sci. 52, 3–28 (1993).

206 Kim, B.-J. Comparative anatomy and phylogeny of the family Mullidae (Teleostei: Perciformes). Memoirs of the Graduate School of Fisheries Sciences, Hokkaido University 49, 1–74 (2002).

207 Parenti, L. R. Relationships of atherinomorph fishes (Teleostei). Bull. Mar. Sci. 52, 170–196. (1993).

208 Reznick, D. N., Furness, A. I., Meredith, R. W. & Springer, M. S. The origin and biogeographic diversification of fishes in the family Poeciliidae. PLOS ONE 12, e0172546, doi:10.1371/journal.pone.0172546 (2017).

209 Bragança, P. H. N., Amorim, P. F. & Costa, W. J. E. M. Pantanodontidae (Teleostei, Cyprinodontiformes), the sister group to all other cyprinodontoid killifishes as inferred by molecular data. Zoosystematics and Evolution 94, 137–145 (2018).

210 Pohl, M., Milvertz, F., Meyer, A. & Vences, M. Multigene phylogeny of cyprinodontiform fishes suggests continental radiations and a rogue taxon position of Pantanodon. Vertebrate Zoology 65, 37–44 (2015).

211 Parenti, L. R. A phylogenetic and biogeographic analysis of cyprinodontiform fishes (Teleostei: Atherinomorpha). Bull. Amer. Mus. Nat. Hist. 168, 335–557 (1981).

212 Costa, W. J. E. M. in Phylogeny and classification of Neotropical fishes (eds L.R. Malabarba et al.) 537–560 (EDIPUCRS, 1998).

213 Lovejoy, N. R. Reinterpreting recapitulation: Systematics of needlefishes and their allies (Teleostei : Beloniformes). Evolution 54, 1349-1362 (2000).

214 Lovejoy, N. R., Iranpour, M. & Collette, B. B. Phylogeny and jaw ontogeny of beloniform fishes. Integ. Comp. Biol. 44, 366–377, doi:Doi 10.1093/Icb/44.5.366 (2004).

215 Dasilao, J. C. & Sasaki, K. Phylogeny of the flyingfish family Exocoetidae (Teleostei, Beloniformes). Ichthyol. Res. 45, 347–353, doi:10.1007/BF02725187 (1998).

216 Bloom, D. D., Unmack, P. J., Gosztonyi, A. E., Piller, K. R. & Lovejoy, N. R. It’s a family matter: molecular phylogenetics of Atheriniformes and the polyphyly of the surf silversides (Family: Notocheiridae). Mol. Phylogenet. Evol. 62, 1025–1030 (2012).

217 Campanella, D. et al. Multi-locus fossil-calibrated phylogeny of Atheriniformes (Teleostei, Ovalentaria). Mol. Phylogenet. Evol. 86, 8–23, doi:https://doi.org/10.1016/j.ympev.2015.03.001 (2015).

218 Dyer, B. S. & Chernoff, B. Phylogenetic relationships among atheriniform fishes (Teleostei: Atherinomorpha). Zool. J. Linn. Soc. 117, 1–69 (1996).

219 Liem, K. F. & Greenwood, P. H. A functional-approach to the phylogeny of the pharyngognath teleosts. Am. Zool. 21, 83–101 (1981).

220 Kaufman, L. & Liem, K. F. Fishes of the suborder Labroidei (Pisces: Perciformes): phylogeny, ecology, and evolutionary significance. Breviora 472, 1–19 (1982).

221 Springer, V. G. & Orrell, T. M. Appendix: phylogenetic analysis of 147 families of acanthomorph fishes based primarily on dorsal gill-arch muscles and skeleton. - In: Springer, V.G. & Johnson, G.D. Study of the dorsal gill-arch musculature of teleostome fishes, with special reference to the Actinopterygii. Bull. Biol. Soc. Wash. 11, 236–260 (2004).

222 Rosen, D. E. & Patterson, C. On Müller’s and Cuvier’s concepts of pharyngognath and labyrinth fishes and the classification of percomorph fishes, with an atlas of percomorph dorsal gill arches. Amer. Mus. Novit. 2983, 1–57 (1990).

223 Collins, R. A., Britz, R. & Rüber, L. Phylogenetic systematics of leaffishes (Teleostei: Polycentridae, Nandidae). J. Zool. Syst. Evol. Res. 53, 259–272, doi:10.1111/jzs.12103 (2015).

224 Lundberg, J. G. in Biological Relationships between Africa and South America (ed P. Goldblatt) 156–199 (Yale University Press, 1993).

225 Lavoué, S. Origins of Afrotropical freshwater fishes. Zool. J. Linn. Soc. 188, 345–411, doi:10.1093/zoolinnean/zlz039 (2020).

226 Stiassny, M. L. J. What are grey mullets? Bull. Mar. Sci. 52, 197–219. (1993).

227 Stiassny, M. L. J. Notes on the anatomy and relationships of the bedotiid fishes of Madagascar, with a taxonomic revision of the genus *Rheocles* (Atherinomorpha: Bedotiidae). Amer. Mus. Novit. 2979, 1–33 (1990).

228 Lin, H. C. & Hastings, P. A. Phylogeny and biogeography of a shallow water fish clade (Teleostei: Blenniiformes). BMC Evol. Biol. 13, doi:Unsp 210 Doi 10.1186/1471-2148-13-210 (2013).

229 Kawahara, R. et al. Interrelationships of the 11 gasterosteiform families (sticklebacks, pipefishes, and their relatives): a new perspective based on whole mitogenome sequences from 75 higher teleosts. Mol. Phylogenet. Evol. 46, 224–236 (2008).

230 Mabuchi, K., Miya, M., Azuma, Y. & Nishida, M. Independent evolution of the specialized pharyngeal jaw apparatus in cichlid and labrid fishes. BMC Evol. Biol. 7, -(2007).

231 Britz, R. & Johnson, G. D. “Paradox Lost”: Skeletal ontogeny of *Indostomus paradoxus* and its signficance for the phylogenetic relationships of Indostomidae (Teleostei, Gasterosteiformes). Amer. Mus. Novit. 3383, 1–43 (2002).

232 Travers, R. A. A review of the Mastacembeloidei, a suborder of synbranchiform teleost fishes. Part II: Phylogenetic analysis. Bull. Br. Mus. Nat. Hist. (Zool.) 47, 83–150 (1984).

233 Gosline, W. A. The suborders of perciform fishes. Proc. U.S. Nat. Mus. 124, 1–78 (1968).

234 Datovo, A., de Pinna, M. C. C. & Johnson, G. D. The infrabranchial musculature and Its bearing on the phylogeny of percomorph fishes (Osteichthyes: Teleostei). Plos One 9, e110129, doi:10.1371/journal.pone.0112600 (2014).

235 Li, C. H., Ricardo, B. R., Smith, W. L. & Ortí, G. Monophyly and interrelationships of snook and barramundi (Centropomidae *sensu* Greenwood) and five new markers for fish phylogenetics. Mol. Phylogenet. Evol. 60, 463–471 (2011).

236 Sanciangco, M. D., Carpenter, K. E. & Betancur-R, R. Phylogenetic placement of enigmatic percomorph families (Teleostei: Percomorphaceae). Mol. Phylogenet. Evol. 94, Part B, 565-576, doi:http://dx.doi.org/10.1016/j.ympev.2015.10.006 (2016).

237 Girard, M. G., Davis, M. P. & Smith, W. L. The phylogeny of carangiform fishes: morphological and genomic investigations of a new fish clade. Copeia 108, 265–298, doi:10.1643/CI-19-320 (2020).

238 Chen, W.-J., Bonillo, C. & Lecointre, G. Repeatability of clades as a criterion of reliability: a case study for molecular phylogeny of Acanthomorpha (Teleostei) with larger number of taxa. Mol. Phylogenet. Evol. 26, 262–288 (2003).

239 Dettaï, A. & Lecointre, G. In search of notothenioid (Teleostei) relatives. Antarctic Sci. 16, 71–85 (2004).

240 Matschiner, M., Hanel, R. & Salzburger, W. On the origin and trigger of the notothenioid adaptive radiation. Plos One 6, e18911 (2011).

241 Lautredou, A.-C. et al. New nuclear markers and exploration of the relationships among Serraniformes (Acanthomorpha, Teleostei): the importance of working at multiple scales. Mol. Phylogenet. Evol. 67, 140–155 (2013).

242 Johnson, G. D. *Niphon spinosus*: A primitive epinepheline serranid, with comments on the monophyly and intrarelationships of the Serranidae. Copeia, 777–787 (1983).

243 Johnson, G. D. *Niphon spinosus*, a primitive epinepheline serranid: corroborative evidence from the larvae. Jap. J. Ich. 35, 7–18 (1988).

244 Johnson, G. D. in Ontogeny and systematics of fishes (eds H.G. Moser et al.) 464–498 (American Society of Ichthyologists and Herpetologists, 1984).

245 Near, T. J. et al. Identification of the notothenioid sister lineage illuminates the biogeographic history of an Antarctic adaptive radiation. BMC Evol. Biol. 15, 109, doi:10.1186/s12862-015-0362-9 (2015).

246 Smith, W. L., Elizabeth, E. & Clara, R. Phylogeny and taxonomy of flatheads, scorpionfishes, sea robins, and stonefishes (Percomorpha: Scorpaeniformes) and the evolution of the lachrymal saber. Copeia 106, 94–119, doi:10.1643/CG-17-669 (2018).

247 Clardy, T. R. Phylogenetic systematics of the prickleback family Stichaeidae (Cottiformes: Zoarcidae) using morphological data Ph.D. thesis, The College of William & Mary, (2014).

248 Kwun, H. J. & Kim, J.-K. Molecular phylogeny and new classification of the genera *Eulophias* and *Zoarchias* (PISCES, Zoarcoidei). Mol. Phylogenet. Evol. 69, 787–795, doi:https://doi.org/10.1016/j.ympev.2013.06.025 (2013).

249 Radchenko, O. A. The system of the suborder Zoarcoidei (Pisces, Perciformes) as inferred from molecular genetic data. Russian Journal of Genetics 51, 1096–1112, doi:10.1134/S1022795415100130 (2015).

250 Near, T. J. et al. Nuclear gene-inferred phylogenies resolve the relationships of the enigmatic Pygmy Sunfishes, *Elassoma* (Teleostei: Percomorpha). Mol. Phylogenet. Evol. 63, 388–395, doi:10.1016/j.ympev.2012.01.011 (2012).

251 Chen, W.-J., Lavoué, S., Beheregaray, L. B. & Mayden, R. L. Historical biogeography of a new antitropical clade of temperate freshwater fishes. J. Biogeogr. 41, 1806–1818, doi:10.1111/jbi.12333 (2014).

252 Greenwood, P. H. A revised familial classification for certain cirrhitoid genera (Teleostei, Percoidei Cirrhitoidea), with comments on the group’s monophyly and taxonomic ranking. Bull. Nat. Hist. Mus. Lond. (Zool.) 61, 1–10 (1995).

253 Burridge, C. P. & Smolenski, A. J. Molecular phylogeny of the Cheilodactylidae and Latridae (Perciformes: Cirrhitoidea) with notes on taxonomy and biogeography. Mol. Phylogenet. Evol. 30, 118–127 (2004).

254 Kimura, K., Imamura, H. & Kawai, T. Comparative morphology and phylogenetic systematics of the families Cheilodactylidae and Latridae (Perciformes: Cirrhitoidea), and proposal of a new classification. Zootaxa 4536, doi:10.11646/zootaxa.4536.1.1 (2018).

255 Ludt, W. B., Burridge, C. P. & Chakrabarty, P. A taxonomic revision of Cheilodactylidae and Latridae (Centrarchiformes: Cirrhitoidei) using morphological and genomic characters. Zootaxa 4585, doi:10.11646/zootaxa.4585.1.7 (2019).

256 McDowall, R. M. Relationships and taxonomy of the New Zealand torrent fish, *Cheimarrichthys fosteri* Haast (Pisces: Mugiloididae). Journal of the Royal Soc N Z 3, 199–217 (1973).

257 Pietsch, T. W. Phylogenetic relationships of trachinoid fishes of the family Uranoscopidae. Copeia 1989, 253–303 (1989).

258 Pietsch, T. W. & Zabatien, C. P. Osteology and interrelationships of the sand lances (Teleostei: Ammodytidae). Copeia 1990, 78–100 (1990).

259 Imamura, H. & Matsuura, K. Redefinition and phylogenetic relationships of the family Pinguipedidae (Teleostei : Perciformes). Ichthyol. Res. 50, 259–269 (2003).

260 McDowall, R. M. Biogeography of the New Zealand torrentfish, *Cheimarrichthys fosteri* (Teleostei : Pinguipedidae): a distribution driven mostly by ecology and behaviour. Envir. Biol. Fishes 58, 119–131 (2000).

261 Last, P. R. in FAO species identification guide for fishery purposes: The living marine resources of the Western Central Pacific, Volume 6. Bony fishes part 4 (Labridae to Latimeriidae) (eds K.E. Carpenter & V.H. Niem) 3517 (FAO, 2001).

262 Davis, M. P., Sparks, J. S. & Smith, W. L. Repeated and widespread evolution of bioluminescence in marine fishes. PLoS ONE 11, e0155154, doi:10.1371/journal.pone.0155154 (2016).

263 Prokofiev, A. M. Osteology and some other morphological characters of *Howella sherborni*, with a discussion of the systematic position of the genus (Perciformes, Percoidei). J. Ichthyol. 47, 413–426 (2007).

264 Patterson, C. & Rosen, D. E. in Papers on the systematics of gadiform fishes Vol. 32 Natural History Museum of Los Angeles County Science Series (ed D.M. Cohen) 5–36 (Natural History Museum of Los Angeles County, 1989).

265 Chanet, B. et al. Evidence for a close phylogenetic relationship between the teleost orders Tetraodontiformes and Lophiiformes based on an analysis of soft anatomy. Cybium 37, 179–198 (2013).

266 Baldwin, C. C. The phylogenetic significance of colour patterns in marine teleost larvae. Zool. J. Linn. Soc. 168, 496–563 (2013).

267 Winterbottom, R. The familial phylogeny of the Tetraodontiformes (Acanthopterygii: Pisces) as evidenced by their myology. Smithson. Contrib. Zool. No. 155, 1–201 (1974).

268 Leis, J. M. in Ontogeny and systematics of fishes (eds H.G. Moser et al.) 459–463 (American Society of Ichthyologists and Herpetologists, 1984).

269 Holcroft, N. I. A molecular analysis of the interrelationships of tetraodontiform fishes (Acanthomorpha : Tetraodontiformes). Mol. Phylogenet. Evol. 34, 525–544 (2005).

270 Alfaro, M. E., Santini, F. & Brock, C. D. Do reefs drive diversification in marine teleosts? Evidence from the pufferfish and their allies (Order Tetraodontiformes). Evolution 61, 2104–2126 (2007).

271 Yamanoue, Y. et al. Phylogenetic position of tetraodontiform fishes within the higher teleosts: Bayesian inferences based on 44 whole mitochondrial genome sequences. Mol. Phylogenet. Evol. 45, 89–101 (2007).

272 Santini, F., Sorenson, L. & Alfaro, M. E. A new phylogeny of tetraodontiform fishes (Tetraodontiformes, Acanthomorpha) based on 22 loci. Mol. Phylogenet. Evol. 69, 177-187, doi:Doi 10.1016/J.Ympev.2013.05.014 (2013).

273 Arcila, D., Pyron, R. A., Tyler, J. C., Orti, G. & Betancur-R, R. An evaluation of fossil tip-dating versus node-age calibrations in tetraodontiform fishes (Teleostei: Percomorphaceae). Mol. Phylogenet. Evol. 82, 131–145 (2015).

274 Arcila, D. & Tyler, J. C. Mass extinction in tetraodontiform fishes linked to the Palaeocene-Eocene thermal maximum. Proc. R. Soc. B 284 (2017).

275 Miya, M. et al. Evolutionary history of anglerfishes (Teleostei: Lophiiformes): a mitogenomic perspective. BMC Evol. Biol. 10 (2010).

276 Pietsch, T. W. & Orr, J. W. Phylogenetic relationships of deep-sea anglerfishes of the suborder Ceratioidei (Teleostei: Lophiiformes) based on morphology. Copeia 2007, 1–34 (2007).

277 Derouen, V., Ludt, W. B., Ho, H.-C. & Chakrabarty, P. Examining evolutionary relationships and shifts in depth preferences in batfishes (Lophiiformes: Ogcocephalidae). Mol. Phylogenet. Evol. 84, 27–33, doi:https://doi.org/10.1016/j.ympev.2014.12.011 (2015).

278 Swann, J. B., Holland, S. J., Petersen, M., Pietsch, T. W. & Boehm, T. The immunogenetics of sexual parasitism. Science 369, 1608, doi:10.1126/science.aaz9445 (2020).

279 Pietsch, T. W. Oceanic anglerfishes extraordinary diversity in the deep sea. (University of California Press, 2009).

280 de la Estrella, M., Forest, F., Wieringa, J. J., Fougère-Danezan, M. & Bruneau, A. Insights on the evolutionary origin of Detarioideae, a clade of ecologically dominant tropical African trees. New Phytologist 214, 1722–1735, doi:https://doi.org/10.1111/nph.14523 (2017).

281 Thomson, R. C., Spinks, P. Q. & Shaffer, H. B. A global phylogeny of turtles reveals a burst of climate-associated diversification on continental margins. Proceedings of the National Academy of Sciences 118, e2012215118, doi:10.1073/pnas.2012215118 (2021).

282 Condamine, F. L., Nel, A., Grandcolas, P. & Legendre, F. Fossil and phylogenetic analyses reveal recurrent periods of diversification and extinction in dictyopteran insects. Cladistics 36, 394–412, doi:https://doi.org/10.1111/cla.12412 (2020).

283 May, M. R., Höhna, S. & Moore, B. R. A Bayesian approach for detecting the impact of mass-extinction events on molecular phylogenies when rates of lineage diversification may vary. Methods Ecol Evol 7, 947–959, doi:https://doi.org/10.1111/2041-210X.12563 (2016).

284 Moore, B. R., Höhna, S., May, M. R., Rannala, B. & Huelsenbeck, J. P. Critically evaluating the theory and performance of Bayesian analysis of macroevolutionary mixtures. Proceedings of the National Academy of Sciences 113, 9569–9574 (2016).

285 Rabosky, D. L., Mitchell, J. S. & Chang, J. Is BAMM Flawed? Theoretical and Practical Concerns in the Analysis of Multi-Rate Diversification Models. Syst. Biol. 66, 477–498 (2017).

286 Chang, J., Rabosky, D. L. & Alfaro, M. E. Estimating Diversification Rates on Incompletely Sampled Phylogenies: Theoretical Concerns and Practical Solutions. Syst. Biol. 69, 602–611, doi:10.1093/sysbio/syz081 (2020).

287 Louca, S. & Pennell, M. W. Extant timetrees are consistent with a myriad of diversification histories. Nature 580, 502–505, doi:10.1038/s41586-020-2176-1 (2020).

